# Damaged goods? Evaluating the impact of X-ray damage on conformational heterogeneity in room temperature and cryo-cooled protein crystals

**DOI:** 10.1101/2021.06.27.450091

**Authors:** Filip Yabukarski, Tzanko Doukov, Daniel A Mokhtari, Siyuan Du, Daniel Herschlag

**Author notes:** Current address: Chan Zuckerberg Biohub, San Francisco, California 94518, United States (Filip Yabukarski). **Classification:** Biophysics. **Author contributions:** F.Y., T.D., and D.H, designed research; F.Y. and T.D. performed research; D.A.M. contributed analytic tools; F.Y., T.D., S.D., and D.H. analyzed data; F.Y. and D.H. wrote the paper.

## Abstract

X-ray crystallography is a cornerstone of biochemistry. Traditional freezing of protein crystals to cryo-temperatures mitigates X-ray damage and facilitates crystal handling but provides an incomplete window into the ensemble of conformations at the heart of protein function and energetics. Room temperature (RT) X-ray crystallography provides more extensive ensemble information, and recent developments allow conformational heterogeneity, the experimental manifestation of ensembles, to be extracted from single crystal data. However, high sensitivity to X-ray damage at RT raises concerns about data reliability. To systematically address this critical question, we obtained increasingly X-ray-damaged high-resolution datasets (1.02–1.52 Å) from single thaumatin, proteinase K, and lysozyme crystals. Heterogeneity analyses indicated a modest increase in conformational disorder with X-ray damage. Nevertheless, these effects do not alter overall conclusions and can be minimized by limiting the extent of X-ray damage or eliminated by extrapolation to obtain heterogeneity information free from X-ray damage effects. To compare these effects to damage at cryo temperature and to learn more about damage and heterogeneity in cryo-cooled crystals, we carried out an analogous analysis of increasingly damaged proteinase K cryo datasets (0.9–1.16 Å). We found X-ray damage-associated heterogeneity changes that were not observed at RT. This observation and the scarcity of reported X-ray doses and damage extent render it difficult to distinguish real from artifactual conformations, including those occurring as a function of temperature. The ability to aquire reliable heterogeneity information from single crystals at RT provides strong motivation for further development and routine implementation of RT X-ray crystallography to obtain conformational ensemble information.

**Significance:** X-ray crystallography has allowed biologists to visualize the proteins that carry out complex biological processes and has provided powerful insights into how these molecules function. Our next level of understanding requires information about the ensemble of conformations that is at the heart of protein function and energetics. Prior results have shown that room temperature (RT) X-ray crystallography provides extensive ensemble information, but are subject to extenstive X-ray damage. We found that ensemble information with little or no effects from X-ray damage can be collected at RT. We also found that damage effects may be more prevalent than recognized in structures obtained under current standard cryogenic conditions. RT X-ray crystallography can be routinely implemented to obtain needed information about conformational ensembles.

## Introduction

Crystallographic structural information has been the cornerstone for the interpretation of functional studies of proteins and for understanding biological processes and their regulation. It is now routine to visualize the fold, interactions, and functional sites of proteins via X-ray crystallography. Tens of thousands of protein crystal structures in the Protein Data Bank (PDB, (1)) have been leveraged, together with simplified energetic or empirical rules, to predict protein structures from sequence and to design new proteins (2–4).

Nevertheless, the advances in predicting *structure* have not been paralleled by equal progress in connecting structure to *energetics*—in predicting folding free energies or binding and catalytic constants. This limitation arises, at least in part, because the free energies associated with processes such as folding, binding, and catalysis are defined by the ensemble of conformational states sampled by the protein, with the probability of adopting each state being determined by its relative free energy via Boltzmann distributions; in other words, biological function is dictated by conformational landscapes, with each landscape defining a conformational ensemble (5–9). Ensemble information is required to relate structure to free energies and thus to relate structural states to how fast and efficiently proteins carry out their functions.

Traditional X-ray crystallography provides limited ensemble information (10–12). Information about the conformational heterogeneity present in the crystal, the experimental manifestation of ensembles, has sometimes been extracted from crystallographic B-factors, but these values are incomplete reporters of conformational heterogeneity (13, 14). Further, the traditional practice of cooling crystals to cryogenic temperatures can alter conformational distributions, including at functional sites (15–19).

Ensemble information can be extracted from “high-sequence similarity PDB” (HSP) ensembles obtained from dozens of traditional X-ray crystallography models of different protein variants and in different crystal forms (20). The overall ensemble information obtained was shown to generally agree with measurements at room temperature and has provided invaluable insights into conformational landscapes, though there are also temperature-induced conformational effects (20, 21). Perhaps the largest limitation to this approach is that tens of new cryo X-ray structures are required, rendering it impractical to obtain ensemble information for new variants as is needed to test models and explore new systems. Thus, additional approaches for obtaining ensemble information are needed.

Accurate ensemble information can be obtained at room temperature from a single protein crystal using novel modeling methods. This approach eliminates potential artifacts from cryo-cooling and alleviates the need for dozens of crystal structures to obtain an HSP ensemble (15, 18, 22–25). In addition, recent computational methods have provided more accurate representations of conformational heterogeneity from X-ray data. The program *Ringer* evaluates electron density to uncover low-population alternative side-chain rotameric states (26), while the program *qFit*, combined with manual modeling, delivers multi-conformer models with explicit alternative conformations and orientations (24, 25, 27). Crystallographic disorder parameters (1-S^2^)^1^ quantify the conformational heterogeneity within multi-conformer models and accurately represent solution behavior (22). These tools enable the detection and quantification of changes in conformational heterogeneity between different protein states (21, 29, 30).

Collecting high-quality and complete X-ray diffraction datasets from protein crystals at room temperature is challenging because of the increased crystal X-ray damage sensitivity relative to cryo-temperature (31–37). Cryo-crystallography became overwhelmingly popular after the realization that cryo-cooling increased the amount and quality of diffraction data obtainable from a single crystal (38, 39). Nevertheless, the need for accurate ensemble information and the limitations of cryo-crystallography in delivering such information reignited interest in room temperature X-ray crystallography (18, 23, 40).

A definitive return to room temperature X-ray crystallography requires methods to routinely collect high quality data at widely accessible synchrotron facilities and minimizing X-ray damage to protein crystals (34, 41–44) (see *supplementary text 1*). In prior work we described a widely applicable approach for collecting high-quality room temperature X-ray diffraction datasets from single crystals (41) which now allows us to address the second major challenge, the impact of X-ray damage on conformational heterogeneity determination.

X-ray damage inevitably occurs during any synchrotron data collection, but to date only one study that we are aware of has directly addressed the effects of X-ray damage on conformational heterogeneity in protein crystals at room temperature, concluding that the overall heterogeneity is not damage-dominated (28). Following from these encouraging findings, there is a clear need for in-depth, quantitative, and systematic analysis of the effects of X-ray damage on conformational heterogeneity to determine the capabilities of room temperature crystallography to deliver atomic-level conformational ensemble information and to learn how to minimize or eliminate damage-based artifacts, if possible. In addition, studies of X-ray damage under cryo conditions have focused on chemical damage (32, 45–47), but there is little information about, and thus a need for analysis of, damage effects on apparent conformational heterogeneity.

To determine the reliability of conformational heterogeneity information obtained from X-ray diffraction at room temperature, we obtained increasingly damaged datasets from single crystals of thaumatin, proteinase K, and lysozyme. We obtained high-resolution data (room temperature datasets of 1.02-1.52 Å in this work), facilitating the detection and quantification of heterogeneity and the ability to isolate X-ray damage effects. Our results indicated lack of major X-ray damage-induced changes in protein side chain rotameric states. Our analyses suggested a modest and unevenly distributed increase in conformational disorder with X-ray damage and suggested that limiting the overall diffraction intensity decay of the data to about 70% of its initial value can provide datasets largely devoid of X-ray-induced conformational heterogeneity effects. Importantly, we provide an analysis pipeline to ensure that this is the case and demonstrate that X-ray damage-associated heterogeneity effects can be corrected for, via a generally-applicable procedure. Complementary analysis of atomic-resolution cryo temperature X-ray datasets (0.9 – 1.16 Å) revealed X-ray damage-associated conformational heterogeneity changes that were not observed at room temperature. These observations caution that, in practice, structures obtained from cryo-cooled crystals for which the effects of X-ray damage have not been carefully evaluated may not always represent distinct states from the conformational landscape. Overall, our results, combined with prior findings (28, 48), suggest that room temperature X-ray data from single crystals can and should be used to obtain accurate conformational heterogeneity information as is needed to connect structure to energetics and to provide deeper understanding of protein function.

## Results

### Obtaining increasingly X-ray damaged datasets from single crystals at room temperature

X-ray damage to protein crystals occurs at all temperatures but protein crystal are equisitely sensitive to X-ray damage at room temperature, rendering it critical to assess damage under these conditions (see *supplementary text 2*). We employed recent data collection approach that optimized the room temperature X-ray diffraction experiment and allowed us to obtain high-resolution datasets of increasing X-ray damage from thaumatin, proteinase K, and lysozyme single crystals ((41) **Figure 1, Tables S1-S4**). For each crystal, we struck a compromise between the extent of X-ray-damage and the resolution of the most X-ray damaged dataset (see Materials and Methods). We collected diffraction data until the total diffraction intensity dropped to about half of its initial value, which represents significant X-ray damage, but the most damaged datasets were still of high-resolution (≤1.5 Å, see Materials and Methods (**Figure 1A–B**)). To assess the effects of X-ray damage on heterogeneity, we obtained four complete increasingly X-ray damaged datasets from the thaumatin and proteinase K crystals and three from the lysozyme crystal (**Figure 1**, **Tables S1-S4**, see Mateirals and Methods). Overall, all diffraction datasets were of sufficiently high resolution to allow accurate and quantitative modeling of conformational heterogeneity using the *Ringer* and *qFit* multi-conformer approaches and to quantify conformational heterogeneity via crystallographic disorder parameters (1-S^2^) (22, 25, 26).

**Figure 1.**
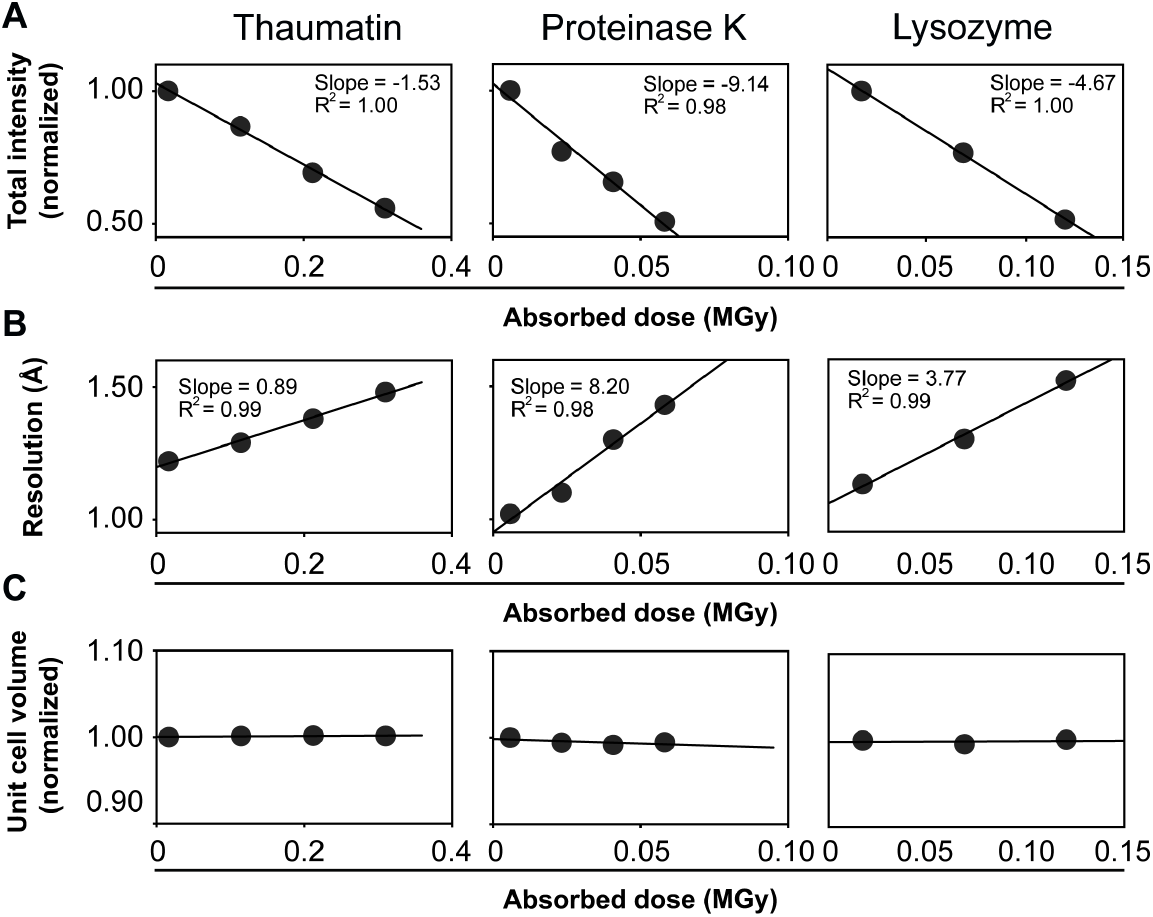
Increasingly X-ray damaged room temperature datasets from single crystals of thaumatin, proteinase K, and lysozyme. Each plot point represents a complete dataset of 120 images that was collected from the same crystal orientation (see Materials and Methods). (**A**) Normalized dataset intensity, (**B**) dataset resolution, and (**C**) unit cell volume as a function of the absorbed X-ray dose (average diffraction weighted dose (DWD), see Materials and Methods).

### X-ray damage effects at room temperature: *Ringer* and electron density analysis of side-chain rotameric distributions

#### Lack of major X-ray damage effects on side-chain rotameric distributions at room temperature

To evaluate the effect of X-ray damage on conformational heterogeneity, we first looked for changes in side-chain rotameric distributions with increasing damage^2^. We used the program *Ringer*, which systematically samples electron density around side chain dihedral angles and can identify low-occupancy alternative rotameric states at low electron density levels, thereby enabling the detection of potential changes in side chain rotamer distributions with X-ray damage (26).

We obtained and compared normalized *Ringer* profiles for each residue in each protein from the increasingly X-ray damaged datasets, and we quantified the similarity between profiles by calculating the Pearson correlation coefficient (P_CC_) between the differentially damaged datasets (see Materials and Methods). Analysis of the normalized profiles is illustrated in **Figure 2**.

**Figure 2.**
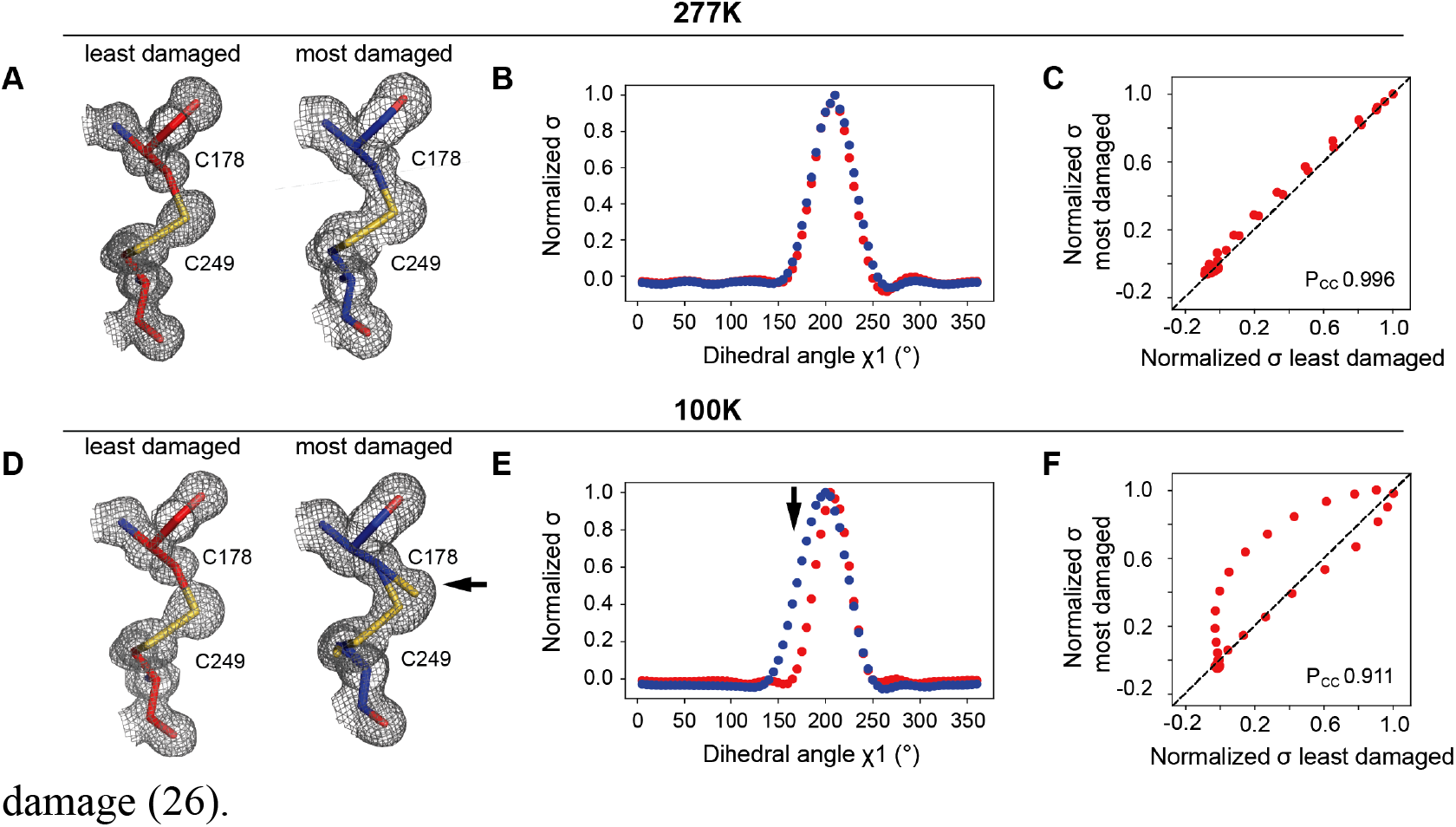
Illustration of *Ringer* analysis to quantitatively evaluate the effects of X-ray damage on side chain rotameric distributions. These panels depict examples from the proteinase K datasets obtained in this work. Data from the least (red, left) and most (blue, right) X-ray damaged datasets collected at room temperature (**A**) or cryo temperature (**D**). Visually, it appears that there is partial X-ray-induced breakage of the disulfide bond in (**D**) but not in (**A**). The arrow points to the appearance of a new population for C178 that manifests as a broadened distribution. The electron density (1 σ, in grey mesh) and model (in sticks) of the C178-C249 disulfide bond are shown. (**B, E**) Normalized *Ringer* profiles. Plots of electron density (σ) as a function of dihedral angle χ^1^ for C178 (electron density from the least and most damaged datasets in red and blue, respectively). Each point in a *Ringer* profile represents the electron density for a specified dihedral angle (see Materials and Methods). The arrow indicates a difference between the least and most damaged datasets corresponding to the appearance of a new state for C178 in the damaged cryo dataset (**D**). (**C, F**) Correlation plots between electron density values (σ) from B and E, respectively, of least (x-axis) and most (y-axis) damaged datasets. The Pearson correlation coefficient (P_CC_) represents the agreement between least and most damaged *Ringer* profiles with P_CC_ = 1 for a perfect correlation (dashed line).

When the least and most X-ray damaged room temperature datasets are compared for all residues of thaumatin, proteinase K, and lysozyme, 98% of the residues had P_CC_ ≥ 0.95 (**Figure 3A, Table S5**); mean square errors (MSEs) of the distributions yielded similar agreement (**Figure S1** and **Table S5**). Further, the majority of the 2% outliers exhibited *Ringer* profiles with little change in their rotameric distributions (**Figure S2**). Only three residues out of 485 side chains across all three protein gave a clear change in rotameric distribution (**Figure 3B,** see also **Figure S2**). These results indicate that X-ray damage at room temperature does not induce changes in side-chain rotameric distributions for the vast majority of residues.

**Figure 3.**
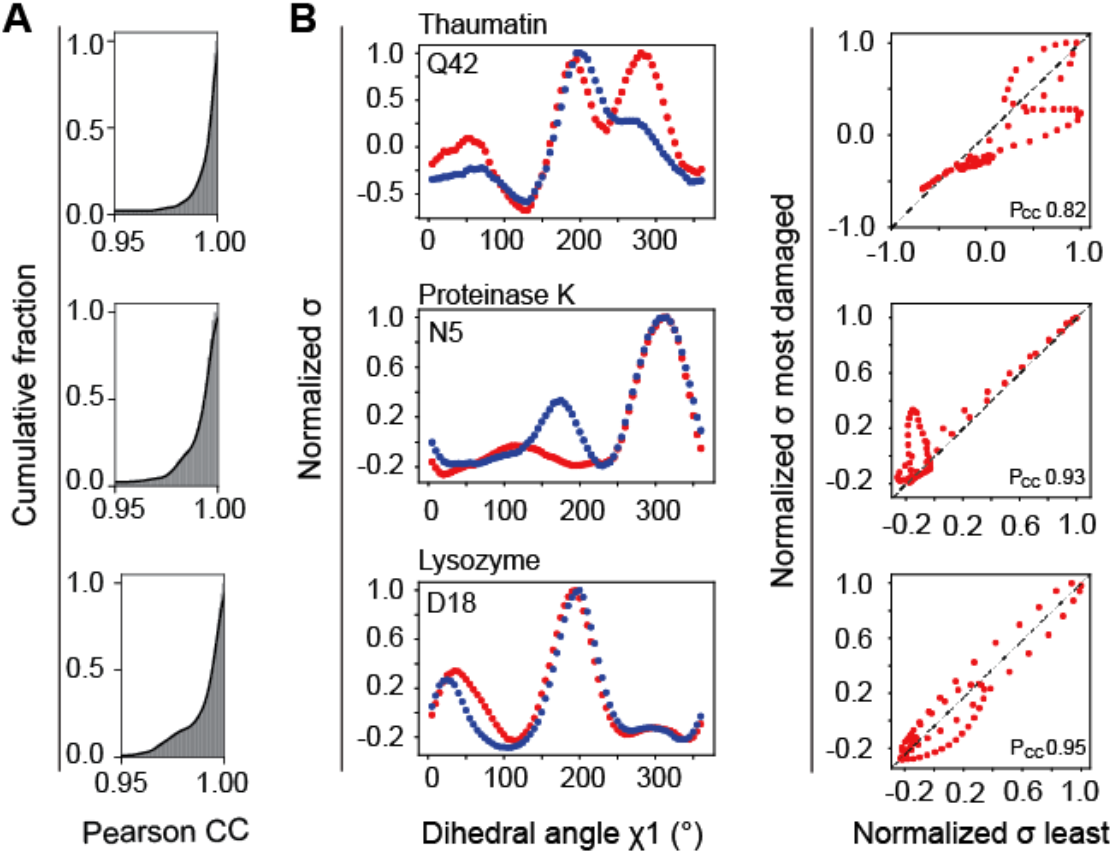
Limited effects of X-ray damage on side-chain rotameric distributions at room temperature. (**A**) Cumulative fraction of Pearson correlation coefficients (P_CC_) for the dihedral angle χ^1^ of each residue in thaumatin (top), proteinase K (middle), and lysozyme (bottom) (see **Table S5** for Pcc values for all residues). P_CC_ were obtained as described in **Figure 2** and Materials and Methods. (**B**) Most significant outliers with P_CC_ ≤ 0.95 for thaumatin (top), proteinase K (middle), and lysozyme (bottom). **Figure S2** shows the *Ringer* profiles and correlation plots for all residues with Pcc ≤ 0.95.

We next focused on disulfide bonds and functional active site residues, all found to be exquisitely sensitive to X-ray damage at cryo temeperatures (45–47, 49–55). For both disulfide bonds in all three proteins and functional active site residues in lysozyme and proteinase K, quantitative comparison of normalized *Ringer* profiles showed no evidence for changes in rotameric states (see **Figures S3-S5** and *supplementary texts* 3 and 4 for additional analyses, including electron density analyses, and discussion), consistent with the above observations of the lack of X-ray damage-associated rotameric state changes for the vast majority of residues in all three proteins. Thus, X-ray damage at room temperature does not alter conclusions about local conformational preferences and distributions of disulfide bonds and functional active site groups.

### The effect of X-ray damage on conformational heterogeneity determined from multi-conformer models and disorder parameters

In the previous section we determined if changes in rotameric states appear as a result of X-ray damage at room temperature. In this section we carry out complementary analyses to further evaluate the effects of X-ray damage at room temperature by obtaining and comparing multi-conformer models and crystallographic disorder parameters (1-S^2^) from the increasingly damaged datasets for each protein (see Materials and Methods, **Figure 4A, Tables S8-S10**).While the *Ringer* analysis above allowed to evaluate potential changes in rotameric states, the disorder parameter (1-S^2^) analysis can identify changes in heterogeneity regardless if they are associated with changes in rotameric populations or changes within the same rotameric state (22). Further, quantification of heterogeneity from multi-conformer models is most often achived via (1-S^2^).

**Figure 4.**
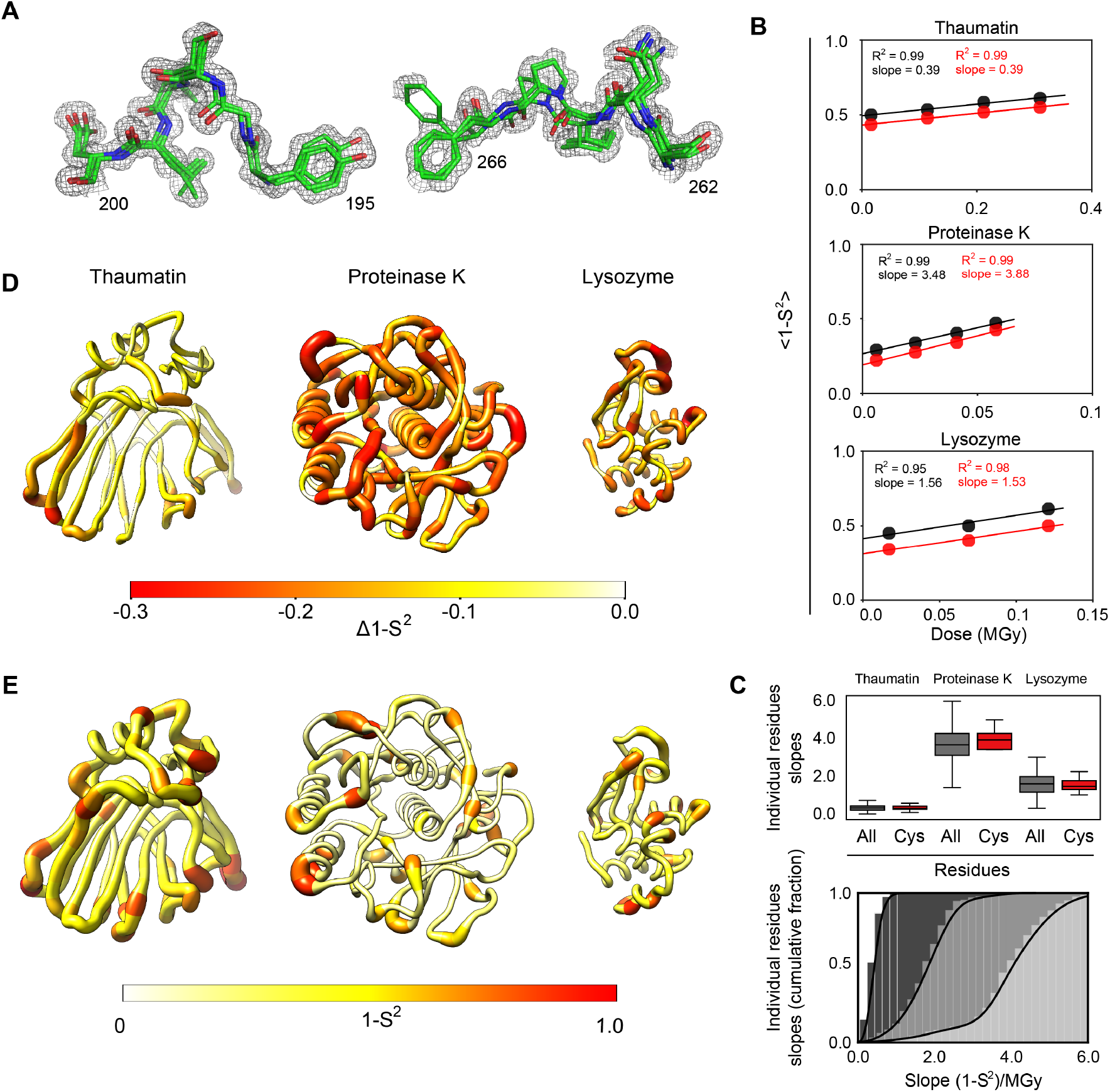
Evaluating the effects of X-ray damage on conformational heterogeneity at room temperature using multi-conformer models and disorder parameters (1-S^2^). (**A**) Illustration of multi-conformer models; shown are two regions of the proteinase K 277 K multi-conformer model from the least damaged dataset (PDB 7LPU). (**B**) Average (1-S^2^) for all residues (black circles) and for disulfide bond-forming cysteine residues (red circles) as a function of the absorbed X-ray dose for each protein. (**C**) Top. Boxplot showing the distribution of slopes obtained from a plot of the (1-S^2^) value as a function of the absorbed X-ray dose calculated for each residue in each protein (see **Tables S12-S14** for individual values). The whiskers show the quartiles of the distributions. Bottom. The data for all residues from the top plot now represented as a cumulative fraction. Dark to light grey: thaumatin, lysozyme, proteinase K. (**D**) Δ(1-S^2^) values between the least and most damaged datasets plotted on the structure of each protein. The diameter of the worm representation and the color both correlate with the magnitude of the Δ(1-S^2^). (**E**) X-ray damage free (zero-dose) (1-S^2^) values plotted on the structure of each protein as in panel (**D**).

#### Evaluating the effect of X-ray damage on conformational heterogeneity from increasingly damaged datasets

We quantified the overall impact of X-ray damage on conformational heterogeneity by first comparing the average (1-S^2^) values, ⟨1 − *S*^2^⟩, from the increasingly damaged datasets (**Tables S12-S14**). The black symbols in **Figure 4B** show that ⟨1 − *S*^2^⟩ values gradually increase with increasing dose for all three proteins. To determine if heterogeneity increases equally for all residues or if the effects are heterogeneous, we plotted the (1-S^2^) value as a function of X-ray dose for each residue in each protein and we extracted the slopes of the corresponding plots (**Tables S12-S14**). The grey symbols in **Figure 4C** show a wide distribution of positive slopes, indicating that while the conformational heterogeneity increased for all residues, it increased to different extent for different residues. **Figures 4D** shows the changes in conformational heterogeneity throughout each protein, by plotting the difference (1-S^2^) values (Δ(1-S^2^)) for their least and most X-ray damaged datasets on the protein structure (also see **Figure S7**). There are regions with greater and lesser X-ray damage effects throughout each protein, with some apparent clustering of regions of higher X-ray damage.

We compared (1-S^2^) values for disulfide-bond forming cysteine residues and for all other residues (**Figures 4B–C**), which provided evidence for similar behavior of cysteine and other residues, consistent with the analyses from the previous section. As we were able to collect increasingly X-ray-damaged datasets from the same crystal, we could obtain (1-S^2^) values for each residue that are free of X-ray damage effects by extrapolating to zero dose (zero-dose (1-S^2^)); the extrapolation was done by fitting a linear equation to the plot of (1-S^2^) for each residue as a function of the absorbed X-ray dose so that the y-intercept represented the zero-dose values of (1-S^2^) (see Materials and Methods). **Figure 4E** shows the distribution of zero-dose (1-S^2^) values for each protein. While not trivial, these extrapolated data provide the highest accuracy conformational heterogeneity information.

#### Evaluating the effect of accumulating X-ray damage on apparent conformational heterogeneity

In the previous sections we addressed the fundamental question of how X-ray damage impacts conformational heterogeneity in protein crystals at room temperature using independent increasingly damaged datasets, and we showed how these modest effects could be eliminated by extrapolating (1-S^2^) values to zero dose (**Figure 4E**). Nevertheless, it may not be possible, or practical, in all cases to obtain a series of high-resolution and complete increasingly X-ray damaged room temperature datasets from the same crystal. As a result, in most cases all the collected X-ray diffraction data with increasing damage will likely be merged in a single dataset. Thus, the confident use of conformational heterogeneity information from protein crystals at room temperature requires an assessment of the extent to which merging increasingly X-ray damaged data impacts the observed heterogeneity. To emulate a typical data collection and analysis from single crystals, we analyzed datasets in which an increasing amount of increasingly X-ray damaged data were merged.

We obtained four increasingly X-ray damaged datasets for thaumatin and proteinase K and three such datasets for lysozyme by merging increasing amounts of diffraction data (**Tables S15-S17** for diffraction statistics, and see Materials and Methods). We refined multi-conformer models and calculated (1-S^2^) values for each dataset (see **Tables S15-S17** for refinement statistics). The values of ⟨1 − S^2^⟩ as a function of relative overall intensity decay (*I/I*_0_) for each protein are plotted in **Figure 5A**. The ⟨1 − S^2^⟩ values are essentially unchanged for datasets for which the diffraction intensity of the merged data has decayed to about 70% of its initial value (*I/I*_0_ ~ 0.7, average from all three proteins), whereas merging data decayed to more than *I/I*_0_ ~ 0.7 results in an increase in apparent ⟨1 − S^2^⟩ values (**Figure 5A**).

**Figure 5.**
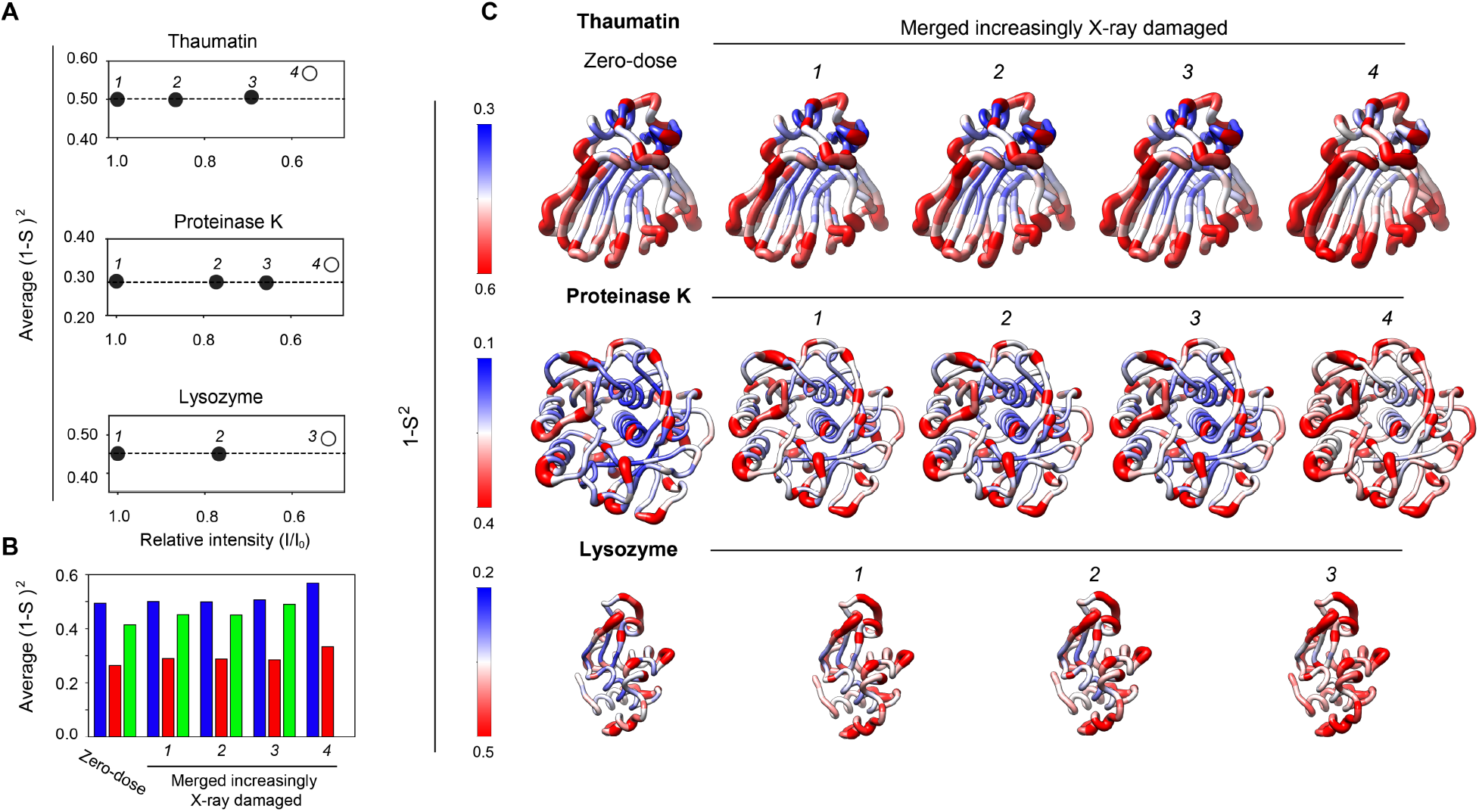
Evaluating the effects of X-ray damage on conformational heterogeneity at room temperature in datasets with increasingly damaged data merged together. (**A**) Analysis of datasets that have accumulated an increasing amount of damage to emulate typical room temperature data collection from single crystals. Average (1-S^2^) as a function of the relative intensity (I/I_0_) of the most damaged diffraction data merged together (see **Tables S18**-**S20**). *I/I*_0_ is the ratio of the total intensity of a diffraction dataset from the beginning of the data collection (*I*_0_) and the intensity of a diffraction dataset obtained from the same crystal orientation in later stages of data collection (*I*) (see Materials and Methods). Open symbols are used when (*I/I*_0_) is less than 0.6. (**B**) Bar plot of the average values for the zero-dose (1-S^2^) and for (1-S^2^) from the increasingly damaged (merged) thaumatin (blue), proteinase K (red) and lysozyme (green) datasets. (**C**) Comparison of zero-dose (1-S^2^) (far left) and (1-S^2^) from the least to most X-ray damaged merged datasets (left to right) with (1-S^2^) values plotted on the structure of each protein. Note that the scales differ to best visualize each protein. The diameter of the worm representation is correlated with the magnitude of the (1-S^2^) values. The zero-dose (1-S^2^) and the (1-S^2^) from the least damaged datasets are qualitatively and quantitatively similar (also see **Figure S10**).

**Figure 5B** shows a comparison of the average values for the zero-dose (1-S^2^) and for (1-S^2^) from the increasingly X-ray damaged datasets and **Figure 5C** shows the per residue (1-S^2^) values for the same datasets plotted on the structure of each protein. This analysis indicates that the average and per residue values for the zero-dose (1-S^2^) and for all (1-S^2^) from increasingly X-ray damaged datasets are of the same magnitude. Detailed analysis indicates that the values of the zero-dose (1-S^2^) and for the (1-S^2^) from the datasets that have decayed not more than *I/I*_0_ ~ 0.7 are higly similar and that the (1-S^2^) for the most X-ray damaged datasets (dataset 4 for thaumatin and proteinase K and dataset 3 for lysozyme, all of which have decayed to *I/I_0_* < 0.6) are the most different when compared to all others. **Figure 5C** indicates that the (1-S^2^) values increase most strikingly in the central region of thaumatin’s β-sandwich and the solvent exposed turns, while the most notable increase in (1-S^2^) values in proteinase K is predominantly observed within the protein’s core, in particular in the two central α-helices. Overall, these results suggest that, on average, protein diffraction data decayed to not more than about *I/I*_0_ ~ 0.7 can be merged and that the extracted conformational heterogeneity information can be used with reasonable confidence.

### Evaluating the effects of X-ray damage on conformational heterogeneity at cryo temperature

The importance of X-ray structural models in biomedical research is reflected in the tremendous developments in X-ray sources of increasing brilliance which, together with the extended lifetimes of protein crystals at cryo-temperatures (38, 39), have resulted in the determination of more than 10^5^ X-ray structures to date. The primary goals of X-ray crystallography has been to obtain information about the folded state of proteins, and how they interact with one another and with small molecules. In keeping with this goal, since its introduction in X-ray crystallography, cryo-cooling has enabled the facile collection of diffraction data that can be used to obtain high-resolution models for a very large number of proteins.

Nevertheless, cryo-cooling can alter protein conformations in the crystal (15–19). Now, with increasing focus on how proteins function as dynamic entities, accuracy in structural details and information that extends beyond static states are imperative (6, 7, 15, 22, 23, 56–58). Much prior work has demonstrated specific X-ray damage to side chains at cryo temperatures, but our current knowledge of the extent and impact of X-ray damage on the distribution of rotameric states under these conditions is limited (33, 46, 47, 51, 53). Russi and coworkers observed an X-ray-induced increase in average (1-S^2^) values under cryo-conditions (28), suggesting potential impact on rotamer distributions and underscoring the need for further evaluation, which we have carried out herein for proteinase K.

To assess the effects of X-ray damage on side chain rotameric distributions at cryo temperature, we collected increasingly X-ray damaged cryo temperature proteinase K datasets until the total diffraction intensity dropped to about half of its initial value (**Figure 6A** and **Table S21**), paralleling the procedure we used in evaluating room temperature X-ray damage (**Figure 1**). We carried out our comparisons in this way because diffraction intensity decay is often used as a practical measure of damage in X-ray data collection (59). However, as the X-ray dose required to decrease the overall diffraction intensity is typically 50-100 times lower at room temperature than at cryo temperature (see *suppelemntary text 2*), the total X-ray doses absorbed by the proteinase K crystal were substantially higher at cryo temperature (**Tables S2** and **S21**).

**Figure 6.**
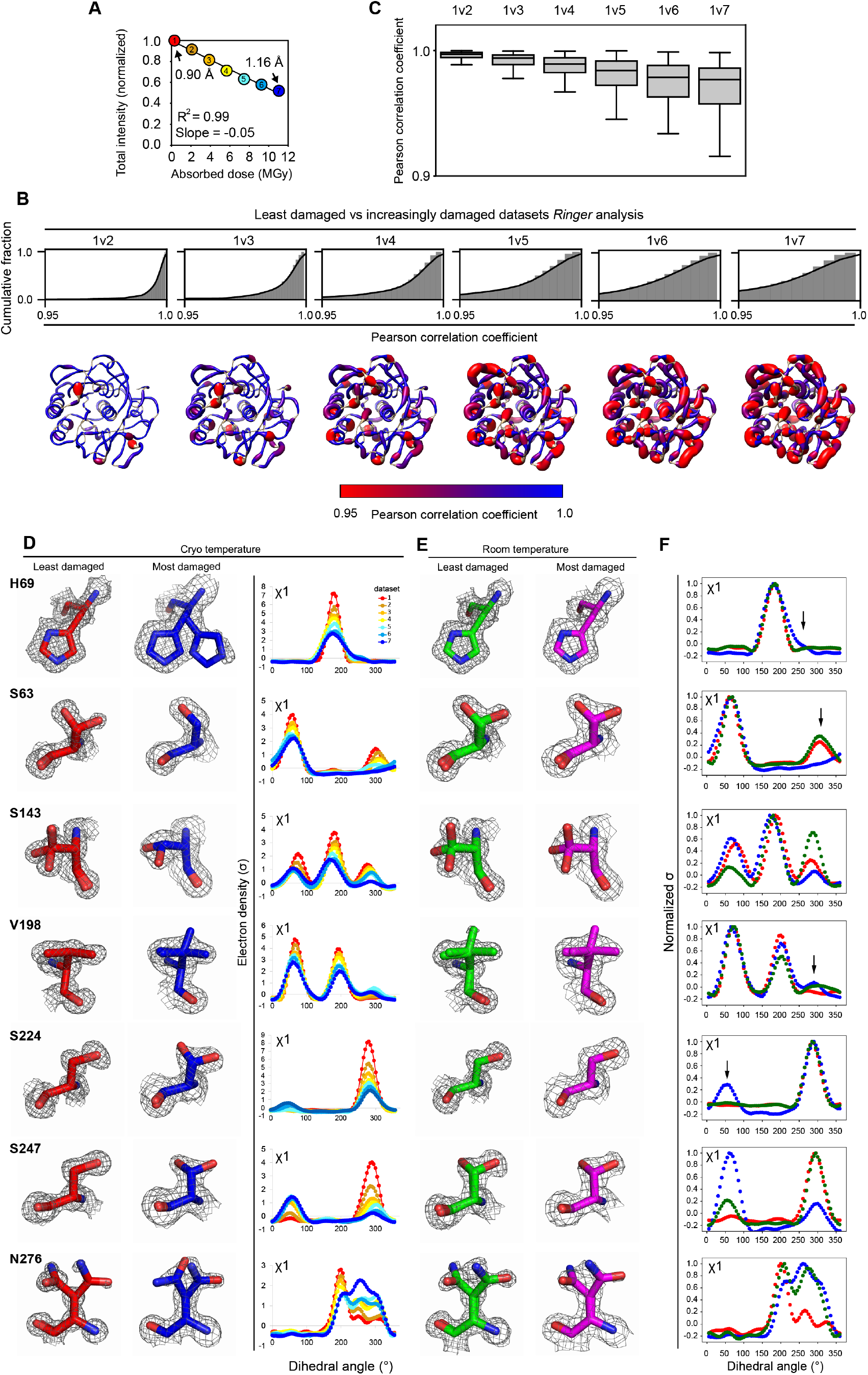
X-ray damage to cryo cooled crystals can alter protein side chain rotameric distributions and can impact conclusions about changes in conformational heterogeneity with temperature. (**A**) Plot of the normalized total intensity *vs.* absorbed dose for increasingly damaged datasets collected at 100 K from the same orientation of a single proteinase K crystal (**Table S21**). The resolutions of the least damaged and most damaged cryo datasets are indicated. Cumulative fraction (**B**) and boxplots (**C**) of Pearson correlation coefficients between the *Ringer* profiles for all residues in the least and increasingly damaged cryo datasets. In (**B**) the lower row shows P_CC_ values plotted on the proteinase K structure. The diameter of the worm representation is inversely correlated with the magnitude of the P_CC_. (**Figures S12-S17** show the Ringer plots for all residues with PCC ≤ 0.95, see also **Table S22**). (**D-F**) A subset of residues with P_CC_ of ≤0.95. (**D**) Left and middle columns. The least (dataset 1) and most (dataset 7) damaged cryo datasets, residues are shown as sticks. Electron density is shown as grey mesh and is contoured at 1σ for all residues except H69, for which the contour is at 0.4 sigma. Right. Raw *Ringer* profiles for the residues in (**D**) from the increasingly damaged datasets; colors correspond to those in (**A**). (**F**) Same residues as in (**D**) but from the least (dataset 1) and most (dataset 4) damaged room temperature datasets. Electron density is shown as grey mesh as in (**D**). (**F**) Normalized *Ringer* profiles for the residues in (**D**) and (**E**): the least and most damaged 100 K datasets are shown in red and blue, respectively, with the least damaged room temperature dataset shown in green. Arrows show the appearance or disappearance of peaks. The room temperature *Ringer* profiles were obtained from electron density maps with resolution matched to the resolution of the 100 K datasets (i.e. 1.16 Å).

As carried out with our room temperature data, we refined structural models against each of these increasingly X-ray–damaged datasets. **Figure S11** shows the final model and electron density for the least X-ray damaged dataset (0.90 Å resolution), and **Table S21** gives data collection and refinement statistics for all models. Diffraction resolution decayed as expected, although the most damaged dataset was still of relatively high resolution (1.16 Å; **Figure 6A** and **Table S21)**, allowing damage effects to be readily analyzed. The 11 MGy X-ray dose absorbed by the most damaged proteinase K dataset (dataset 7, **Figure 6A**) is significantly below the Henderson and Garman dose limits of 20 MGy and 30 MGy, respectively, which have been used as a rule of thumb in cryo-crystallography X-ray data collection (59, 60) and is consistent with previous estimates of diffraction half-doses of about 10 MGy at 100 K (61). In addition, the widespread use in the recent years of micron-sized X-ray beams with increasingly brilliant X-ray sources at many synchrotrons presumably leads to depositing large X-ray doses a general practice, and makes it likely that a significant fraction of high-resolution cryo structures on the PDB have been obtained from datasets that have absorbed an X-ray dose of at least 10 MGy. Nevertheless, for the vast majority of high-resolution structures in the PDB, the X-ray dose and extent of diffraction decay are not reported and the extent of X-ray damage present is difficult or not possible to estimate. Thus, an in-depth analysis of cryo datasets with increasing X-ray damage for a model protein is important to begin to calibrate these possible effects, and we have used proteinase K for this analysis.

#### Evaluating X-ray damage effects on proteinase K side chain rotameric distributions at cryo temperatures

To assess any potential changes in side-chain rotameric distributions with X-ray damage and determine whether this could be an issue of concern in practice, we obtained *Ringer* profiles for each side chain in each of the increasingly X-ray damaged datasets and calculated P_CC_ between the *Ringer* profiles of the least damaged dataset and each of the increasingly damaged datasets, for every residue. The fraction of residues for which P_CC_ ≥ 0.95 decreased with increasing X-ray damage (**Figures 6B** and **6C**), suggesting that X-ray damage alters multiple side chain rotameric distributions for proteinase K.

Inspection and comparison of the *Ringer* profiles and electron density maps for residues with P_CC_ ≤ 0.95 revealed changes in rotameric state distributions for a number of residues, with the most striking changes shown in **Figure 6D**. X-ray damage had diverse effects: disappearance of rotameric states (S63), redistribution of side chain rotameric distributions (S143, V198, S224, S247, N276), and appearance of new rotameric states (H69). Inspection of the electron density for surrounding residues indicated that none of these changes appeared to be directly coupled to breaking any of proteinase K’s two disulfide bonds. The changes in rotameric distributions appeared gradually (**Figure 6D,** right panel), and at least in some instances were already present at *I/I*_0_ of about 0.9 (e.g., S247 and N276 in **Figure 6D**, right panel, compare *Ringer* profiles in red (dataset 1) and tan (dataset 2)). Thus, even with seemingly high-quality diffraction data, X-ray damage can be present and can affect certain interpretations and conclusions.

#### Comparing X-ray damage effects on side chain rotameric distribution at cryo and room temperatures

When the least and most X-ray damaged cryo datasets are compared, only about 80% of all residues have P_CC_ ≥ 0.95, compared to 98% of residues with P_CC_ ≥ 0.95 at room temperature (**Figure 6B** 1v7, compare with **Figure 3B**, middle), suggesting that when datasets the diffraction intensity of which has decayed to a similar extent are compared (e.g. to about 50% of its initial value for the proteinase K and comparing cryo and room temperature datasets of similar initial resolutions (0.90 Å (100 K dataset) and 1.02 Å (277 K)) X-ray damage has a more profound impact on rotameric state distributions at cryo that at room temperature. **Figure 6E** shows the electron density of the least and most X-ray damaged room temperature proteinase K datasets for the same residues as highlighted in **Figure 6D** as subject to damage effects under cryo conditions. The rotameric distributions do not change with X-ray damage at room temperature, consistent with the analyses in previous sections (compare **Figure 6D** left and middle columns and **Figure 6E** left and right columns, respectively).

The above results suggest that X-ray damage present in a cryo dataset may impact structural conclusions and also conclusions made about temperature effects on the conformational landscape. Take as an example the residues from **Figure 6D**. Comparing the least damaged cryo dataset with the room temperature dataset (**Figure 6D**, left and **6E**, left, respectively) identifies new rotameric states for V198, S247, and N276 at room temperature, no changes for S63 and H69, and a redistribution of rotameric states for S143 and S224. However, comparing the most damaged cryo dataset with the room temperature dataset (**Figure 6D**, middle and **6E**, left, respectively) leads to different conclusions–H69 and S224 as residues for which rotameric states disappear, S63 as a residue for which a new rotameric state appears, and V198 and S247 as unchanged. In other words, comparing the room temperature dataset with the most X-ray damaged cryo dataset would identify *false positives* changes to residues S63 and H69 and as *false negative* effects at V198 and S247. In addition, for S143, S224, S247, and N276, comparing the room temperature dataset with either the least or most damaged cryo dataset would lead to different estimates of the change of relative rotameric state distributions with temperature (see also *supplementary text* 5 and **Figure S20**).

#### Proteinase K side chain rotameric distributions at cryo versus room temperature

Previous work identified temperature-associated differences between a XFEL room temperature dataset and a synchrotron cryo-temperature dataset for proteinase K, including an increase in the number of alternative conformations at room temperature (62). While XFEL data are X-ray damage free (see *suplementary text 1*), synchrotron X-ray data are not, and inspection of the structural model from the cryo-cooled synchrotron data from previous work (PDB code 5KXV) and comparison with the least and most X-ray damaged cryo-cooled models from this study suggest that the 5KXV cryo-cooled structure had suffered X-ray damage effects (**Figure S21**), suggesting that rotameric distributions could have been altered. The previous work focused on assessing changes of rotameric states across temperature; our analysis allowed us to extend these comparisons to identify changes in rotameric distributions in the absence of apparent changes in the rotameric states. Indeed, the side chains of residues S216 and S219 change rotameric distributions without changing rotameric states, extending the number of serine residues for which temperature-associated rotameric changes were previously observed (**Figure S18**) (62). The rotameric changes that we observed across temperature (26 residues out of 279, about 10%, **Figure S20**, left, **Figure S18**) provide a highly accurate window into temperature-associated molecular behaviors central to proteins and may provide a valuable resource to evaluate and improve the ability of computational models to predict temperature-dependent changes.

The cryo and room temperature proteinase K datasets obtained in this work, because of their high accuracy and resolution, allow for analyses and comparisons that might not have been previously possible. For example, our room temperature dataset allowed us to uncover a dynamic network comprising side chains and water molecules extending from the solvent-exposed high-affinity Ca^2+^ binding site in proteinase K about 17 Å into the hydrophobic core. Cryo-cooling appears to quench these motions (**Figure 7** and **Figure S22**).

**Figure 7.**
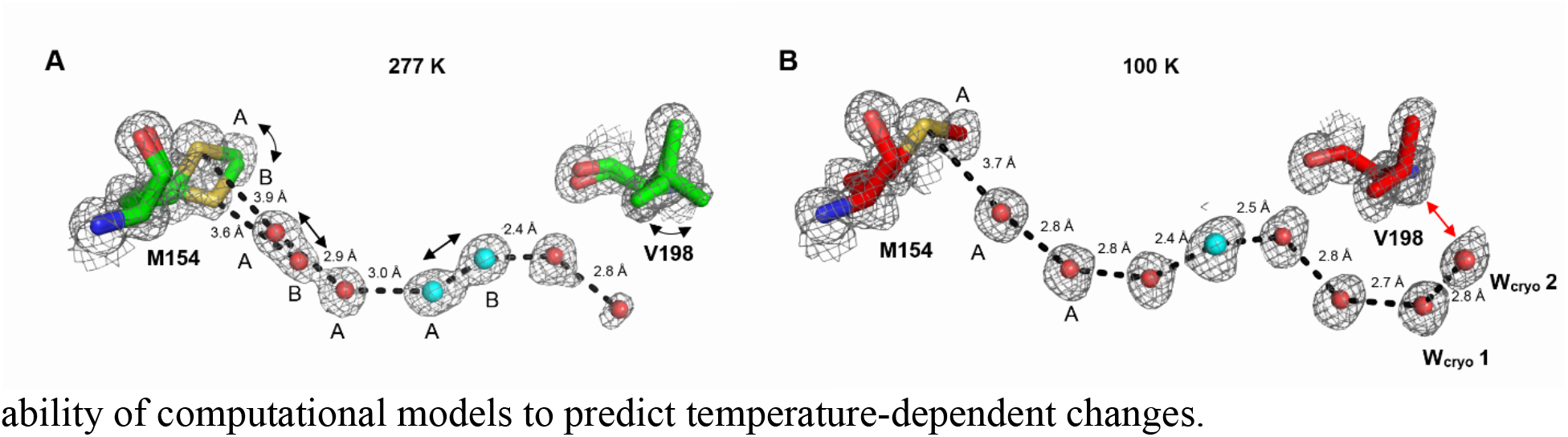
Extended network of interactions couples motions of distant residues M154 and V198 in proteinase K. M154 and V198 motions are quenched at cryo (right) but not at room temperature (left). Water molecules form an extended network upon freezing that quenches V198 motion. Waters W_cryo_ 1 and W_cryo_ 2 are too close to V198 to allow the existence of the alternative rotameric state observed at room temperature (also see **Figure S22**).

#### Comparison of hydrogen bonds from room temperature and cryo X-ray data

Hydrogen bonds are ubiquitous and central to protein folding, molecular recognition, and enzyme catalysis and have been studied broadly using high-resolution protein crystal structures, with the vast majority obtained via cryo X-ray crystallography (63–72). Given the importance of hydrogen bonds, we used our high-resolution structural data to appraise the effects of X-ray damage on hydrogen bond lengths. We identified all hydrogen bond pairs involving backbone and side chains within each protein and then compared the hydrogen bond lengths between the structural models obtained from the least and most X-ray damaged datasets (see Materials and Methods). The correlation between hydrogen bond lengths obtained from the least damaged (dataset 1) and most damaged (dataset 7) 100 K proteinase K structures has a slope of 1.03 and R^2^ value of 0.92 (**Figure S23A**), and **Figures S23B** and **S24** show that R^2^ decreases with increasing X-ray damage. The same analysis using the least (dataset 1) and most (dataset 4) damaged room temperature proteinase K datasets yielded a correlation with a slope of 1.03 and R^2^ value of 0.98 (**Figure S23C**, left), and we found similar highly consistent correlations for thaumatin and lysozyme least vs most X-ray damaged room temperature datasets (slope = 1.00 and R^2^ = 0.97 and slope = 1.04 and R^2^ = 0.95, respectively, **Figure S23C**, middle and right, respectively). Our analyses suggest that X-ray damage is more likely to impact hydrogen bond lengths obtained from cryo-cooled crystals compared to room temperature crystals, underscoring the importance of tracing X-ray damage and its impact whenever possible.

## Discussion

As we move to a new era of mechanistic understanding in biology, focus has shifted from static protein structures to dynamic ensembles of states that are determined by free energy landscapes (5–7, 23, 30, 56, 73, 74). To understand the complex functions carried out by biomolecules and their control we need to determine how and why these ensembles and their underlying energy landscapes change as biomolecules carry out their function, and how they change in response to interacting with other molecules and to mutation.

In principle, the limitations of cryo-crystallography and HSP ensembles (see Introduction) are overcome by room temperature X-ray crystallography (23–26, 30), but room temperature X-ray crystallography itself poses several challenges: crystals without cryo-cooling generally diffract to lower resolution, are highly sensitive to X-ray damage, and present practical challenges for crystal handling and data collection (34–37, 41). We previously described approaches to make room temperature X-ray crystallography more practical and robust and to reduce X-ray damage (41). Here we carry out the next step needed to test and establish the broad applicability of room temperature X-ray crystallography, building on and extending prior studies to systematically assess the effects of X-ray damage (28, 48, 75).

### The impact of X-ray damage at room temperature

Our analyses show that very high-resolution data, 1.02 – 1.22 Å in this work, can be obtained without X-ray damage causing substantial distortion of the conformations present. We found no evidence for X-ray damage-associated changes in rotameric distributions of functional active site residues, local conformational heterogeneity, or hydrogen bond lengths. We did find evidence for some X-ray damage occuring to disulfide bonds, in agreement with prior work (28, 48, 76), but no evidence for X-ray associated changes in cysteine rotameric distributions and no extreme sensitivity of these bonds.

From a practical perspective, Blundell and Johnson in their foundational room temperature work recommended discarding crystals when the relative diffraction intensity (*I/I*_0_) of a given reflection dropped to 0.85 of its original value (*I*_0_), but suggested that a limit of 0.7 (*I/I*_0_) could be acceptable if the crystal supply was limited (77). The analyses in this work allowed us to further evaluate the proposed decay limit. We found that merging data with overall intensity decreases of up to about 70% of the initial value (*I/I*_0_ ~ 0.7) did not substantially impact the calculated heterogeneity. Thus, the Blundell and Johnson’s lower intensity decay limit is expected to provide conformational heterogeneity information devoid of major alterations from X-ray damage and can be further tested and refined following the approaches used herein.

Overall, the results herein and our prior work (41) provide the means to readily collect room temperature X-ray data and to test and ensure that the data collected and models obtained are of high integrity. In addition, we demonstrate the ablity to obtain “gold standard” zero-damage ensemble information and extract ensemble information free of X-ray damage for instances where the highest accuracy and precision are needed.^3^ While not every protein will be suitable for room temperature X-ray crystallography, due to the need for larger-than-average crystals to facilitate high-quality data collection (41) and for relatively high resolutions to robustly model and interpret heterogeneity (generally better than 1.7–2.0 Å, depending on the method (22, 24, 25)), many proteins and protein-complexes meet these criteria and can be used to obtain generalizable insights into the interplay of dynamics and function and provide ground truth measurements for testing and developing computational models.

### The impact of X-ray damage at cryo temperature

The effects of X-ray damage on protein structure at cryo temperatures has been extensively investigated, with a large number of studies revealing X-ray-induced disulfide bond reduction and side chain decarboxylation (45–47, 59). Here we evaluated the effects of X-ray damage within a cryo-cooled crystal on the apparent conformational landscape of a protein, as the appearance or disappearance of rotameric states can lead to erroneous mechanistic, functional, and energetic conclusions.

When comparing proteinase K cryogenic *versus* room temperature datasets of similar resolution and with similar loss in the overall diffraction intensity due to X-ray damage, we found fewer effects from X-ray damage at room temperature. We also found multiple changes and multiple types of changes in rotameric states for several individual side chains that were not observed at room temperature. These changes included states that appeared, disappeared, and redistributed. Subtle changes in rotameric states and hydrogen bond lengths were also more prevalent under the cryogenic conditions.

The X-ray dose required to halve the diffraction intensity of a protein crystal is typically 50-100 times higher at cryo temperature than at room temperature so that less diffraction decay is expected and observed at cryo temperature for a given X-ray dose (34–37). Nevertheless, in practice, data are often collected based on empirical measures of the amount of damage so that cryogenic data are typically collected with much higher X-ray doses than room temperature data and, importantly, these X-ray doses are often not known.

In their seminal work on cryo-cooled ferritin crystals, Owen and coworkers analyzed traditional structural models and electron density maps and proposed a practical diffraction intensity decay limit of *I/I*_0_ ~ 0.7; the authors cautioned, and very rightly so, that larger decays could compromise the biological information (59). Our observations that rotameric state changes are already present at *I/I*_0_ ~ 0.9, suggest that, in addition to following the recommended *I/I*_0_ ~ 0.7 limit, in-depth analyses may be required to evaluate and address X-ray damage effects at even lower decays when detailed ensemble comparisons are intended (also see *supplementary text* 6).

These findings have direct implications for HSP ensemble analyses. HSP ensembles obtained from multiple cryogenic X-ray structural models provide valuable depictions of overall protein ensemble properties (20, 21). However these ensembles are typically collections of structural models obtained for other purposes, often from crystals that have absorbed relatively high X-ray doses to obtain structural data to high resolution, so that, correspondingly, the amount of X-ray damage can be high and the effects on rotameric distributions generally unknown.

### X-ray damage and multitemperature X-ray crystallography

Cryo-cooling can alter conformational heterogeneity, and it has been found that over one-third of residues in protein crystals exhibit different and generally broader distribution of states at room temperture (15, 18, 23, 30, 78). Temperature-induced changes could be localized or could propagate throughout the protein, and our analysis of proteinase K provided an example of propagated temperature-induced changes where freezing appears to hinder a long-range rotameric side-chain rearrangement by hindering the ability of water molecules to reorient when frozen. In principle, the temperature dependence of a conformational ensemble can provide information about the forces underlying the energy landscape, and particular attention has been paid to changes arising from the so-called glass transition around 180–220 K (30, 79, 80). More broadly, an exciting new approach termed “Multitemperature Multiconformer X-ray crystallography” (MMX) was recently proposed in which a series of datasets obtained at various temperatures can be used to follow changes in rotameric distributions with temperature and could be exploited to uncover energetic coupling between states and provide testable hypothesis about allostery in proteins (30, 78). Our results show that effects from X-ray damage can alter these conclusions. X-ray damage effects can lead to both false positives and false negatives and to incorrect conclusions that particular changes do or do not happen. More generally, confident interpretation of conformational heterogeneity changes across temperature will require evaluation of the effects of X-ray damage on conformational heterogeneity at each temperature, following the approaches outlined in this work (also see *supplementary text 7*).

Fundamentally, conformational distributions of protein residues and their temperature-dependent changes can provide invaluable information about the forces that dictate protein behavior and thus function. Such information can in principle be used to tune and test force fields, and ensure their accuracy and general applicability. As we move toward an era with reliable information about protein ensembles, we can envision an understanding of protein function firmly grounded in the fundamental forces that dictate and define the energy landscapes that guide conformational states and function.

## Materials and methods

### Protein crystallization, X-ray diffraction data collection, and model refinment

All proteins were crystallized as previously described (41); single-crystal diffraction data were collected at SSRL, beamline BL9-2 at 277 K or at 100 K. Diffraction data processing, model building, and difference electron density maps were carried out using standard methods. Multi-conformer models were obtained and crystallographic disorder parameters were calculated as previously described (22, 25) (see also *SI Appendix*). All structural models were deposited on the PDB and all accession codes can be found in the *SI Appendix.*

### *Ringer* analysis

*Ringer* profiles were obtained for each residue in each protein using the *Ringer* program as implemented in phenix.refine and using a 5° sampling angle (26) (see also *SI Appendix*).

### Calculating Pearson correlation coefficients (P_CC_)

The Pearson correlation coefficient (Pcc) between normalized *Ringer* profiles was calculated using the scipy.stats.pearsonr function of the SciPy package in Python 3 (81) (see also *SI Appendix*).

### Hydrogen Bond Comparisons

Hydrogen atoms were written using the Reduce program (82). Only hydrogen bonds with nitrogen or oxygen donor/acceptor and those made between protein atoms were included in the analysis. A relative B-factor for the hydrogen bond is calculated using B-factors of the least damaged dataset (see also *SI Appendix*).

## Acknowledgments

Use of the Stanford Synchrotron Radiation Lightsource (SSRL), SLAC National Accelerator Laboratory, is supported by the U.S. Department of Energy, Office of Science, and Office of Basic Energy Sciences under Contract No. DE-AC02-76SF00515. The SSRL Structural Molecular Biology Program is supported by the DOE Office of Biological and Environmental Research and by the National Institute of Health (NIH), National Institute of General Medical Sciences (NIGMS, P41GM103393). The contents of this publication are solely the responsibility of the authors and do not necessarily represent the official views of NIH or NIGMS. This work was funded by a National Science Foundation (NSF) Grant (MCB-1714723) to DH. FY was supported in part by a long-term Human Frontiers Science Program postdoctoral fellowship. DAM acknowledges support from the Stanford Medical Scientist Training Program and a Stanford Interdisciplinary Graduate Fellowship (Anonymous Donor) affiliated with Stanford ChEM-H. We thank Dirk Zajonc, Antoine Royant, Lindsay Deis, James S. Fraser, William Weis, Ariana Peck, and Catherine Stark for feedback, Lisa Dunn (SSRL) for help with scheduling experimental beam time.

## Footnotes

1 The original work used order parameters (S^2^). Disorder parameters are simply 1-(S^2^) and were used in this and prior work (28).

2 Changes in rotameric distributions include appearance of a new rotameric state (e.g., 100% A → 50% A and 50% B), dissapearence of an existing rotameric state (e.g., 50% A and 50% B → 100% A), and/or redistribution of rotameric populations within existing rotameric states (e.g., 20% A and 80% B → 80% A and 20% B)

3 The extrapolated zero-damage disorder parameters (1-S^2^) do not define a unique ensemble of states as, in principle, different combinations of states, each of which can be more or less flexible, can give rise to similar (1-S^2^). Nevertheless, this ensemble information can be used to make direct comparisons to (1-S^2^) from other experimental approaches, such as NMR, and from computational models.

## Notes

### Competing Interest Statement

The authors have declared no competing interest.

