## Supplementary file 1 for "Damaged goods? Evaluating the impact of X-ray damage on conformational heterogeneity in room temperature and cryo-cooled protein crystals"

### Supplementary information

Supplementary text #1

***Room temperature X-ray diffraction data collection.*** Room temperature diffraction data from protein crystals can be obtained at either synchrotron X-ray beamlines or at X-ray free electron lasers (XFELs) facilities. (Although home X-ray sources are still in use, the vast majority of experiments are currently not conducted using home sources.) In particular, serial crystallography using XFELs enabled collecting diffraction data that is essentially X-ray damage-free (1–3). Nevertheless, this approach is not limitation-free: obtainable diffraction resolutions tend to be lower when compared to resolutions obtained at synchrotron beamlines, data collection often requires complex sample delivery instrumentation (e.g., for a crystal slurry), and currently instrument availability is limited. Recent technical and instrumental developments have led to the development of the serial synchrotron crystallography (SSX) technique, which combines both traditional synchrotron data collection with serial diffraction data collection from large number of small crystals, but these datasets are not X-ray damage-free, as is the case for any non-XFEL data collection approach (4).

Supplementary text #2

***X-ray damage in protein crystals.*** X-ray damage can be described as global, the overall decay of diffraction intensity, and specific, the chemical changes in proteins that are directly related to changes in structure (5–7). Global X-ray damage occurs at all temperatures, but the X-ray dose required to halve the diffraction intensity is typically 50-100 times lower at room temperature than at cryo temperatures (8–11). Commensurate with the pervasive use of cryo cooling in X-ray crystallography, there have been numerous studies of X-ray damage under these conditions (5, 12–15). At cryo temperatures, specific X-ray damage generally occurs before substantial overall diffraction intensity decay, and X-ray damage can alter structural details, the most commonly observed of which are disulfide bond reduction, decarboxylation of acidic side chains, and modifications at metallo-centers, with active site residues suggested to be particularly sensitive (5, 12–16). The effects of specific X-ray damage at room temperature have been less studied, but recent work generally points to less specific X-ray damage relative to cryo temperatures (8, 17, 18).

Supplementary text #3

***Assessing disulfide bond X-ray damage at room temperature.*** Disulfide bonds are exquisitely sensitive to X-ray damage. At cryo temperatures specific damage to disulfide bonds, such as breaking the disulfide bond as illustrated in **Figure 2D** often appears before other X-ray damage effects can be observed in electron density maps (12, 16, 19). To evaluate the extent to which disulfide bonds are X-ray damaged with increasing absorbed X-ray doses at room temperature, we compared the *Ringer* profiles and the electron density maps for each of the 26 disulfide bond-forming cysteine residues (13 disulfides) in all three proteins (**Figure S3**). The 2Fo-Fc electron density maps (1σ, grey mesh) for the least *vs.* the most X-ray damaged datasets were highly similar in all cases, the associated Fo-Fc electron density maps (3σ, green and red mesh for positive and negative density peaks, respectively) were featureless, and the corresponding *Ringer* profiles were highly similar in all cases, with $P_{CC} \geq 0.997 \pm 0.002$ in all cases (average and standard deviation, respectively, also see **Table S6**). To more thoroughly assess potential differences due to X-ray damage, we calculated difference Fo-Fo electron density maps between the least and most damaged datasets for each protein (**Figure S4**). Inspection of these maps reveals a stronger than average negative feature and thus unambiguous damage in one case, C121-C193 in thaumatin (**Figure S4**), a disulfide bond within a region that was previously identified as highly sensitive to X-ray damage (20). However, even in this case, the damage is modest; it could not be identified via inspection of conventional 2Fo-Fc and Fo-Fc maps alone and did not lead to the detectable appearance of new rotameric states. Focusing on the C6-C127 and C76-C94 disulfide bonds in lysozyme, previously identified as the two most susceptible to X-ray damage disulfide bonds in cryo-cooled lysozyme crystals (16), we observed no indication of damage for the former and minor damage features (Fo-Fo difference electron density, **Figure S4**) present for the later that could originate from minor direct damage, consistent with spectroscopic evidence for X-ray damage-induced changes in lysozyme disulfide bond vibration at room temperature, presumably due to S-S bond lengthening (16, 17, 21). The X-ray damage effects to disulfides that we observed were generally of the same magnitude as features observed at other random sites, again consistent with prior observations suggesting that at room temperature specific chemical damage could appear as randomly as global damage (17). On the other hand, in thaumatin the Fo-Fo difference electron density map features for the C56-C66 disulfide are indicative of an elongation of the bond (**Figure S4**), consistent with previous observations of disulfide bond elongation caused by X-ray damage (16, 17, 21). For the other residues, peaks around disulfide bonds in difference Fo-Fo maps were weak and similar in magnitude to peaks throughout the entire map. Overall, our results provide evidence that some X-ray damage occurred to disulfide bonds in our datasets but that the damage was generally not more prominent than damage observed at non-disulfide sites (**Figures S3-S5**). These results are consistent with recent studies at room temperature that found X-ray damage to disulfides (4, 17, 18) and provide evidence for similar dose scales for specific and global X-ray damage evolution. Nevertheless, despite evidence for minor X-ray damage in the difference electron density maps, the *Ringer* analysis showed no evidence for changes in rotameric populations for disulfide bond-forming cysteine residues, including for the C121-C193 bond which presented the strongest difference Fo-Fo features (**Figure S4**), within the limits of X-ray damage explored in this work.

Previous studies provided evidence both for and against X-ray associated damage at disulfide bonds at room temperature (4, 8, 9, 17, 18, 22), but the origin of these conflicting results remains unknown. Why do disulfide bonds appear more or less damaged in different room temperature studies? Based on multi-crystal averaged increasingly X-ray damaged datasets Russi et al. proposed that the amount of damage to a specific site might be different at different X-ray dose rates and that data collected at high dose rates might result in more extensive site-specific damage (18). Thus, differing reports about X-ray damage to disulfide bonds at room temperature could be due to different extent of disulfide bond breaking being caused by the range of X-ray dose rates employed in different studies. Alternatively or in addition, it is possible that the extent of free radical formation and disulfide bond reduction are directly related to the type and concentration of buffer components present in the crystal. Future work should systematically evaluate the effect of X-ray dose and dose rates and the impact of buffer components and their concentration on the extent to which disulfide bonds are X-ray damaged at room temperature.

Supplementary text #4

***Assesing X-ray damage to functional active site residues at room temperature.*** Functional active site residues that are often of most interest, in particular for enzymology studies in which cryo-structures and functional data are combined in structure-function analyses, have consistently been identified among the most susceptible to X-ray damage in cryo-cooled crystals (14, 15, 23–27). While some of this tendency may result from more careful analysis of active sites, there is evidence that aspartate and glutamate residues, which are common at active sites, are more sensitive to X-ray damage than residues other than disulfide bonds (12, 14–16), including the lysozyme E35 and D52 active site residues previously described as highly-sensitive to damage (15)

As little is known about the impact of X-ray damage on active site residues at room temperature, we evaluated the effects on lysozyme catalytic residues E35 and D52 and proteinase K catalytic residues D39, H69, and S224 (catalytic triad) and N161 (oxyanion hole). We focused on evaluating potential changes in rotameric distributions. As above, we compared *Ringer* profiles and electron density maps from the least and most X-ray damaged room temperature datasets. These comparisons revealed nearly identical *Ringer* profiles with average $P_{CC} \geq 0.989 \pm 0.012$ (average and standard deviation, respectively, see **Table S7** for individual values) (**Figure S5A**), highly-similar 2Fo-Fc electron density maps, and featureless Fo-Fc and Fo-Fo difference electron density maps, the later also indicated lack of major decarboxylation occurring due to X-ray damage (**Figure S5B-D**). In particular, while decarboxylation of acidic residues has been described as one of the most pervasive X-ray damage effects in cryo-cooled protein crystals (12, 14), including the lysozyme E35 and D52 active site residues previously described as highly-sensitive to damage (15), we found no evidence for significant X-ray damage to these active site residues in our room temperature data (**Figure S5**). Similarly, we observe no significant damage to the catalytic triad His 69 in proteinase K (**Figure S5**), in contrast to the high X-ray damage sensitivity observed for the equivalent His 440 in cryo-cooled crystals of acetylcholine esterase (15). Our comparisons of the least and most X-ray damaged proteinase K and lysozyme room temperature datasets indicate lack of significant damage to functional active site residues and no changes in rotameric state populations (**Figures S3-S5**), consistent with observations of lack of X-ray damage effects on the conformational distributions of functional active site residues in the enzyme CypA (18). Thus, the results presented in this work extend previous findings and further increase the confidence in conformational heterogeneity information obtained from protein crystals at room temperature.

Supplementary text #5

***Comparing X-ray damage effects on side chain rotameric distribution at cryo and room temperatures.*** **Figure S20** shows the overall distribution of temperature-induced rotameric changes identified from *Ringer* profiles with $P_{CC} \leq 0.95$ as obtained from comparison of the least X-ray damaged room temperature proteinase K dataset with the least (left) and the most X-ray damaged proteinase K cryo dataset cryo dataset (right). While both comparisons point to structure-wide temperature-induced changes, the comparison with the more X-ray damaged cryo dataset shows nearly twice as many side chains that undergo apparent temperature-dependent conformational changes (43 vs 26, **Figures S18-S19**). Thus, X-ray damage under cryo conditions can lead to overestimates of temperature-induced changes, and these differences are observed throughout the structure (**Figure S20**).

Supplementary text #6

***Modeling X-ray damage effects using B-factors.*** With respect to modeling X-ray damage effects in cryo-cooled crystals, previous work suggested that the dominant X-ray damage effects could be adequately modelled by B-factors (18). Our results indicate that while B-factors may generally provide adequate proxies for overall X-ray damage trends, in-depth X-ray damage analyses benefits from more complex models that capture the appearance or disappearance of rotameric states, as B-factors do not adequately represent changes in rotameric states.

Supplementary text #7

***X-ray damage effects and conformational heterogeneity at intermediate temperatures.*** Global X-ray damage occurs at all temperatures, but the X-ray dose required to decrease the overall diffraction intensity (e.g. to halve the intensity) is lower at cryo temperature (see *supplemental text* #2), allowing the collection of higher resolution data and/or more data. However, cryo-cooling can quench and alter conformational heterogeneity in cryo-cooled protein crystals relative to crystals at room temperature (28–31). Thus, there are advantages and disadvantages associated with the collection of diffraction data from either cryo-cooled and room temperature crystals.

Recent work from multiple groups collectively suggested that collecting data at intermediate temperatures could have distinct advantages over data collected from either cryo-cooled or room temperatures crystals (11). First, foundational work has provided evidence for activation of both harmonic and anharmonic motions in protein crystals above the protein glass transition temperature generally occurring within the 180-220 K temperature range (30, 31). Building on this work, more recent studies provided evidence for a complex evolution of conformational heterogeneity in the 180-220 K temperature range, and also suggested that the vast majority of anharmonic motions, e.g., those responsible for the population of alternative side chain rotameric states, are activated at and above 250 K (28, 29). Thus, a body of work indicates that at 250 K, conformational heterogeneity should be similar to conformational heterogeneity at room temperature. Second, X-ray damage work by Warkentin and Thorne on thaumatin crystals indicated that at temperatures slightly below room temperature, crystals were more resistant to global X-ray damage than at room temperature (11). They found that much of the increased resilience of protein crystals to global X-ray damage observed at 100 K relative to room temperature can be achieved by cooling the crystals to temperatures below room temperature but above the protein glass transition where solvent remains fluid (11). Thus, prior work indicates that, while conformational heterogeneity in protein crystals is expected to be similar at 250 K and room temperature, more diffraction data can be collected at 250 K relative to room temperature from a given protein crystal, making 250 K a temperature of particular interest for protein X-ray crystallography.

Because X-ray damage has pervasive effects on protein structure at cryo temperatures, including alteration of conformational heterogeneity, while X-ray damage effects appear minimal at room temperature, it is currently unclear how X-ray damage would affect conformational heterogeneity at 250 K. Thus, to confidently model conformational heterogeneity using diffraction data collected at 250 K, there is a need to evaluate the effects of X-ray damage at this intermediate temperature.

### Materials and Methods

**Protein crystallization and X-ray diffraction data collection.** *Tritirachium album* proteinase K (catalog # P2308), *Thaumatococcus daniellii* thaumatin (catalog # T7638), and hen egg lysozyme (catalog #L4919) were purchased from Sigma. Proteins were crystallized at room temperature using standard literature conditions and as previously described (using a hanging drop (proteinase K and lysozyme) and sitting drop (thaumatin) setups (32)). Briefly, lysozyme dissolved in 0.1 M sodium acetate pH 4.6 was crystallized in 0.6 M sodium chloride; thaumatin dissolved in water was crystallized in 24% potassium sodium tartrate, 15% ethylene glycol (v/v), and 0.1 M BisTris Propane pH 6, and proteinase K dissolved in 0.05 M Tris HCl, pH 7.5, was crystallized in either 1 M ammonium sulfate or 0.5 M sodium nitrate.

Room temperature X-ray diffraction data from single crystals was collected using a recently described approach (32). Briefly, prior to data collection, crystals were transferred from the crystallization solution to paratone N oil (Hampton Research, Aliso Viejo, CA) where excess crystallization solution was stripped and crystals where then either frozen in liquid nitrogen for 100 K data collection (proteinase K) and then mounted on the goniometer or directly mounted on the goniometer for 277 K data collection. Data collection temperature was controlled using the beamline $N_2$ cryocooler/heater. Increasingly X-ray damaged single-crystal diffraction data were collected at SSRL, beamline BL9-2, using wavelength of 0.88557 Å. See **Tables 1-2, S1-S4** and **S21** for diffraction data collection statistics. For all crystals diffraction data were collected using the rotation method and collecting consecutive 360° datasets. Only the first 120° of each 360° rotation were used for subsequent analysis unless stated otherwise. Collecting increasingly X-ray damaged datasets from single crystals allowed to circumvent potential complications associated with merging partial diffraction datasets from multiple crystals. Because diffraction resolution inevitably decreases with increasing X-ray damage and high resolution data are required to reliably detect potential X-ray induced conformational heterogeneity changes (in particular the appearance or disappearance of alternative rotameric states (33, 34)), we struck a compromise between the extent of damage and the overall resolution of the most damaged dataset in each series (**Figure 1**); we collected diffraction data until the total diffraction intensity dropped to about half of its initial value ($I/I_0 \sim 0.5$), which represents a significant extent of damage, yet the most damaged datasets were still of high-resolution (~1.4 – 1.5 Å, resolution was cut at $CC_{1/2} \geq 0.30$). Any potential benefit of collecting more X-ray damaged datasets would have been offset by the increasingly lower resolution of datasets, which in turn would have reduced the accuracy of rotameric state analysis and multi-conformer modeling (for example, making minor side chain conformations undetectable due to the overall loss of electron density details with decreasing resolution) and would have made ($1\text{-}S^2$) analyses less reliable. Absorbed X-ray doses were calculated using RADDOSE 3D (35, 36) and average diffraction weighted doses (DWDs) are reported. DWDs for the increasingly damaged room temperature datasets (**Tables S1-S4**) were comparable to X-ray doses used in previous studies of the effects of X-ray damage on protein crystals at room temperature (17, 18).

***Absence of significant dehydration effects.*** In addition to the increased X-ray damage sensitivity, dehydration, which is not a significant issue under cryo conditions, can occur during room temperature data collection and alter conformational heterogeneity (37). As dehydration often leads to large decreases in unit cell volumes ($V_{\text{unit cell}}$), we assessed dehydration during our data collection by comparing $V_{\text{unit cell}}$ from the increasingly damaged datasets from each crystal for each protein. **Figure 1C** shows that $V_{\text{unit cell}}$ of increasingly damaged datasets are within ±1% of the initial values, providing evidence against significant dehydration during data collection (see also **Tables S1-S4** and (32)). Our data collection approach entails coating the crystals with oil, which limits possible dehydration effects and has advantages over other dehydration-protection approaches (32, 38).

**Crystallographic data processing and model building.** Data processing was carried out with in-house scripts: http://smb.slac.stanford.edu/facilities/software/xds/#autoxds_script. Briefly, data reduction was done using the XDS package (39), scaling and merging was done using *Aimless* (40, 41) and structure factor amplitudes were obtained using *Truncate* (40, 42). The overall diffraction intensity for a given dataset was obtained as reported by XSCALE from the the XDS package (39) by integrating the intensity over a consistent oscillation range (120°) and the same crystal orientation. Initial phases were obtained via molecular replacement using *PHASER* (43) and the PDB models 1RQW, 1IC6, 193L as search models for thaumatin, proteinase K, and lysozyme, respectively. Model building was carried out with the program *ARP/wARP* (44) and manually in *Coot* (45). The commercial thaumatin used in this work (Sigma, catalog # T7638) is a mixture of thaumatin I and thaumatin II, and the final refined models contained the residues that were the best fit to the electron density. Thaumatin I and thaumatin II sequences differ in four positions: 68, 85, 89, and 98 (precursor numbering) with the residues for thaumatin I/thaumatin II being N/K, S/R, K/R R/Q, respectively. The final refined models contain residues corresponding to thaumatin II except for position 68, which corresponds to thaumatin I as these side chains best fit the electron density. The residue at position 207 in final refined proteinase K models contained aspartate instead of serine (reference uniprot code P06373) as the electron density unambiguously supported modeling of an aspartate, consistent with other high-resolution proteinase K structures from the PDB.

Traditional, single conformation models, in which only major alternative side chain and backbone conformations were modeled, were refined manually after visual inspection with *Coot* and using *phenix.refine* (46, 47). Torsion-angle simulated annealing (as implemented in *phenix.refine*) was used during the initial stages of refinement. Riding hydrogens were added in the late stages of refinement and their scattering contribution was accounted for in the refinement. Ligand restraints were generated using the *elBOW* program from *phenix.refine*.

Model quality was assessed using *Molprobity* (48) as implemented in *phenix.refine* and via the PDB Validation server (https://validate-rcsb-2.wwpdb.org/). For each protein, a model was built and refined using the least damaged dataset (dataset 1). The same model was then refined independently against increasingly damaged datasets. Careful visual inspection of models and maps was used to adjust for any damage-related changes and edited models were again refined using *phenix.refine*. This procedure was used to reduce modeling inconsistencies that could originate if a new model was built and refined independently for each increasingly damaged dataset. See **Tables S1-S3** and **S21** for refinement statistics.

Multi-conformer models were obtained from the 277 K diffraction datasets, using previously described methods (28, 33, 49–51). Briefly, the program *qFit* was used to obtain multi-conformer models (49, 50) using as input the traditional single-conformation models obtained above after removal of the riding hydrogen atoms. Subsequent to the automated multi-conformer model building, ill-defined water molecules were deleted and alternative protein side and main chain conformations and orientations were edited manually after visual inspection in *Coot* and based on the fit to the electron density (52). Both alternative side chain rotameric states as well as alternative orientations within the same rotameric state were modeled (see **Figure 4A** for example regions from a typical refined multi-conformer model). Models were subsequently refined with *phenix.refine*, refining atomic isotropic B-factors and occupancies (46, 47). Riding hydrogen atoms were added in the late stages of refinement and their scattering contribution was accounted for in the refinement. Final multi-conformer model quality was checked by *MolProbity* (48) and via the PDB Validation server (https://validate-rcsb-2.wwpdb.org/).

Multi-conformer models from increasingly X-ray damaged datasets were obtained as follows. For each protein, an initial multi-conformer model was built and refined using the least damaged, highest resolution dataset (dataset 1). The refined multi-conformer model was then re-refined independently against increasingly damaged datasets. This procedure was used to reduce modeling inconsistencies that could originate when a new multi-conformer model is built and refined for each increasingly damaged dataset. In particular, the procedure was essential in eliminating subjectivity in modeling low population states as the increasingly damaged datasets were of decreasing resolution, which results in electron density features associated with low-population states to become visible at different electron density standard deviation (σ) levels. Using the refined multi-conformer model from the least damaged dataset for refinement against increasingly damaged datasets in which alternative states' occupancies are refined allows for the diffraction data and not subjectivity to decide on which alternative states disappear (occupancy refines to zero, in which case, after inspection, the alternative state is removed and the model re-refined), which rotameric states persist (non-zero occupancy), and how the persisting states' distributions change (changes in states' occupancies). Careful visual inspection of the resulting models and electron density maps allowed to identify and model any new rotameric states that would appear with increasing damage. Importantly, no evidence for the appearance of new rotameric states with damage was found for the room temperature datasets (and the lack of X-ray damage associated changes was supported by independent *qFit* runs). See **Tables S4, S8-S10** and **S15-S17** for multi-conformer models refinement statistics. The same set of reflections was used for $R_{free}$ calculation in the refinement of all models within an increasing damaged dataset series.

**Crystallographic disorder parameter (1-$S^2$) calculation.** Crystallographic and solution NMR order parameters ($S^2$) have been shown to correlate well and range between 1 for a completely rigid residue and 0 for a completely unrestrained residue (33). Here we used the opposite of order parameters and calculated disorder parameters (1-$S^2$). High resolution data (generally better than ~1.7 Å) is required for crystallographic (1-$S^2$) analysis, and the high resolution of the datasets obtained in this work (1.02 to 1.54 Å, **Tables S1-S4**) make the (1-$S^2$) analyses in this work reliable. Crystallographic disorder parameters, (1-$S^2$), were obtained from the 277 K multi-conformer models as previously described (18, 33). These disorder parameters include both harmonic and anharmonic contributions as captured by the crystallographic atomic displacement parameters (B-factors) and by the occupancies of alternative rotameric states. The analysis was applied to the bond most closely associated with the first side-chain dihedral angle ($\chi_1$), using Cβ—H for all amino acids other than Gly and Cα—H for Gly. For each residue, the extrapolation to zero-dose (1-$S^2$) was done by fitting a linear equation to the plot of (1-$S^2$) as a function of absorbed X-ray dose and extracting the y-intercept. All fits were of good quality with average $R^2$ and standard deviation of 0.91±0.09, 0.96±0.06, and 0.93±0.05 for thaumatin, proteinase K, and lysozyme, respectively (**Tables S12-S14**).

**Assessing the conformational heterogeneity within two distinct crystals of the same protein.** To assess potential variation from experimental and/or modeling factors, we evaluated the extent of similarity between conformational heterogeneity as captured by (1-$S^2$) obtained from two different lysozyme crystals. To eliminate potential (1-$S^2$) differences due to differences in diffraction resolution, quality, or X-ray dose effects, we identified a second crystal that was of similar size, exhibited similar diffraction statistics, and absorbed a similar X-ray dose as the lysozyme crystal and dataset used (crystal 1 *vs.* crystal 2, **Table S4**). The refined multi-conformer models from these two crystals yielded highly similar (1-$S^2$) which correlated with $R^2$ = 0.97 and slope = 1.01 and with an average per residue Δ(1-$S^2$) of 0.01 ± 0.02 (**Figure S6** and **Table S11**). Thus, the conformational heterogeneity information obtained from two different lysozyme crystals was highly similar, and these results lead to a general expectation for strong correlation between conformational heterogeneity from different crystals of the same protein.

All structural models refined in this work were deposited on the PDB with the following accession codes: 7LFG, 7LJV, 7LJW, 7LJZ, 7LK5, 7LK6, 7LNB, 7LNC, 7LND (thaumatin 277 K), 7LN7, 7LPT, 7LPU, 7LPV, 7LQ8, 7LQ9, 7LQA, 7LQB, 7LQC (proteinase K 277 K), 7LTD, 7LTI, 7LTV, 7LU0, 7LU1, 7LQC, 7LU3 (proteinase K 100 K), 7LLP, 7LN8, 7LN9,

7LOQ, 7LOR, 7LP6, 7LPL, 7LPM (lysozyme 277K). See **Tables S1-S4**, **S8-S10**, **S21** for details.

**Fo-Fo difference electron density maps.** Difference electron density maps were obtained via standard procedures using the *Isomorphous difference map* script from the phenix suite (47). For most fair comparisons, the resolution of the two datasets being compared were matched by adjusting the resolution of the less X-ray damaged dataset to match the resolution of the more X-ray damaged dataset. Fo-Fo are displayed at contour level of 3σ unless stated otherwise.

***Ringer* analysis.** *Ringer* profiles were obtained for each residue in each protein as follows. The final structural models and diffraction data were used to calculate composite omit maps to reduce potential model bias (47). To evaluate the effect of map resolution differences on Pearson correlation coefficients ($P_{CC}$), we calculated $P_{CC}$ values for combinations of *Ringer* profiles obtained from proteinase K 100 K dataset 1 refined at 0.9 Å (maximum resolution) or 1.16 Å (resolution of the most damaged dataset 7) and composite omit maps calculated from either model at 0.9 Å (maximum resolution) or 1.16 Å (resolution of the most damaged dataset 7). **Figure S9** shows that artefactual decreased $P_{CC}$ could emerge from differences in the resolution of the electron density maps but that when the resolution of the composite omit maps is adjusted, ≥99% of the $P_{CC}$ values are ≥0.99 (**Figure S9** far right). Thus, for comparison of *Ringer* profiles from the least and most damaged datasets, the composite omit maps for both the least and most damaged datasets were calculated at the resolution of the most damaged dataset. The resulting map and refined models were then submitted to *Ringer* as implemented in the *phenix* suite using a 5° sampling angle (34, 47). Because the absolute amount of electron density (σ) can vary between datasets irrespective of changes in rotameric distributions, we normalized all *Ringer* profiles prior to comparison. The normalized *Ringer* profiles were then used to calculate Pearson correlation coefficients ($P_{CC}$) (see below). Further, the *Ringer* analysis was predominantly focused on χ1 angles because i) most protein side chains have χ1 and ii) because most side chains with χ2 angles are surface exposed and the electron density required to calculate χ2 angles is weaker than the average, making accurate quantitative comparisons difficult. Because all analyzed proteinase K 100 K datasets are of atomic resolution (0.9 – 1.16 Å) and because we observe both appearance and disappearance of rotameric states (**Figure 6D**), the systematic decrease of agreement between *Ringer* profiles from increasingly damaged datasets observed in **Figure 6B-C** is unlikely to be due to poor data resolution that would preclude the detection of rotameric states.

**Calculating Pearson correlation coefficients ($P_{CC}$) and mean square errors (MSE).** The Pearson correlation coefficient (Pcc, also known as Pearson's *r*) between normalized *Ringer* profiles was calculated using the scipy.stats.pearsonr function of the SciPy package in Python 3

(53). Mean square errors were calculated between normalized *Ringer* profiles using the sklearn.metrics. mean_squared_error function of the scikit-learn package in Python 3, with the non-default parameter “squared = False" (54).

**Hydrogen Bond Comparisons.** Hydrogen atoms were written using the *Reduce* program (55). For each comparison, hydrogen bonds were identified in the least X-ray damaged structure using the MDanalysis Hydrogen Bond analysis module (56, 57) with a 3.5 Å heavy distance cutoff and a 120° bond angle cutoff. Only hydrogen bonds with nitrogen or oxygen donor/acceptor and those made between protein atoms were included in the analysis; any “hydrogen bonds” identified between backbone atoms of neighboring residues were also excluded. We omitted hydrogen bonds involving atoms with multiple conformations modeled (main text and **Figure S23-S24**) and confirmed our results by including hydrogen bonds involving backbone nitrogen or oxygen atoms that have multiple conformations and averaging all possible lengths (**Figure S25**). Despite the overall high resolution of all datasets, not all hydrogen bonding groups are similarly well-defined in the electron density and more hydrogen bond length variation is expected for groups with less well-defined electron density; these differences are reflected in the relative B-factor of a hydrogen bonding group and we calculated and used relative B-factors as a proxy for positional accuracy. A relative B-factor for the hydrogen bond is calculated using B-factors of the least damaged structure:

$$\text{relative B-factor}_i = (B_{i,\,donor} / \langle B \rangle + B_{i,\,acceptor} / \langle B \rangle) / 2$$

where $B_{i,\,donor}$ is the B-factor for the donor heavy atom for hydrogen bond i, $B_{i,\,acceptor}$ is the B-factor for the acceptor atom for the same hydrogen bond, and $\langle B \rangle$ is the average B-factor of all atoms in the structure.

**Figure generation.** PyMOL (58) and UCSF Chimera (59) were used for figure generation.

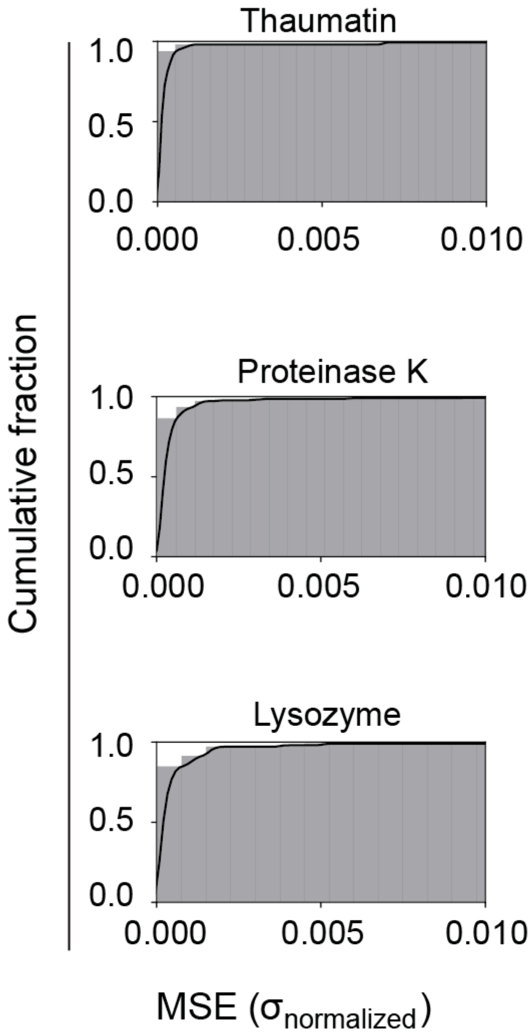

**Figure S1. Modest effects of X-ray damage on side-chain rotameric distributions at room temperature.** Mean square errors (MSE) were obtained in the same way as Pearson correlation coefficients ($P_{CC}$) in **Figure 3A** except that MSEs represent the agreement between the experimental correlation and a slope = 1 correlation line (diagonal) from correlation plots between electron density values ($\sigma$) of two datasets, each plotted on x-axis and y-axis as illustrated in **Figures 2C, F** and **3B**. The cumulative fractions of MSEs shown here are from correlation plots between electron density values ($\sigma$) of the least damaged (x-axis) and most damaged (y-axis) datasets for the dihedral angle $\chi^1$ of each residue in thaumatin (top), proteinase K (middle), and lysozyme (bottom) (see **Table S5** for MSEs for all residues).

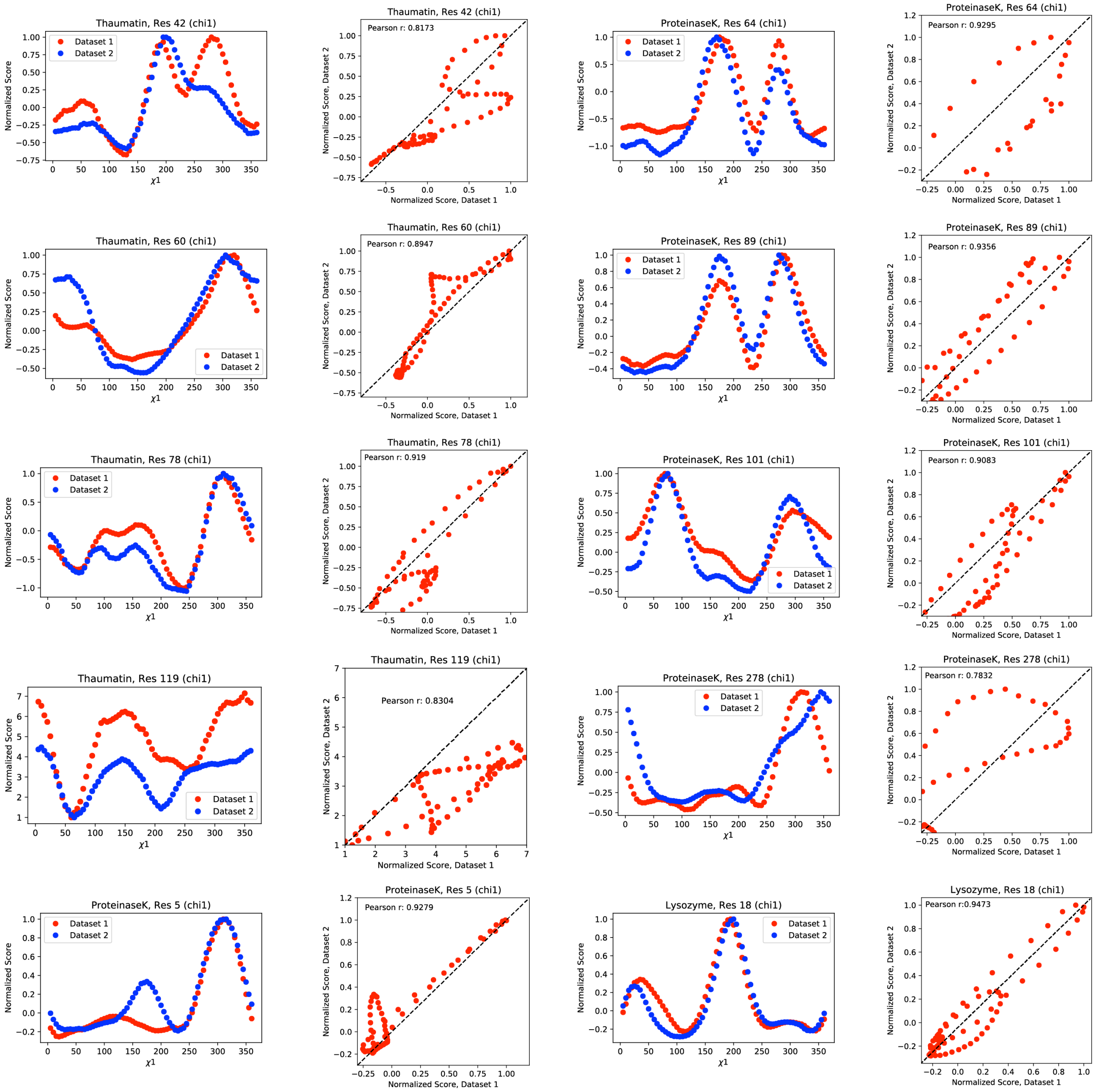

**Figure S2. X-ray damage does not lead to major changes in side chain rotameric states in protein crystals at room temperature.** Normalized *Ringer* profiles of the least (red) and most (blue) damaged datasets for thaumatin, proteinase K, and lysozyme for all residues with Pearson correlation coefficients ($P_{CC}$) ≤ 0.95 and the associated correlation plots and respective $P_{CC}$ values.

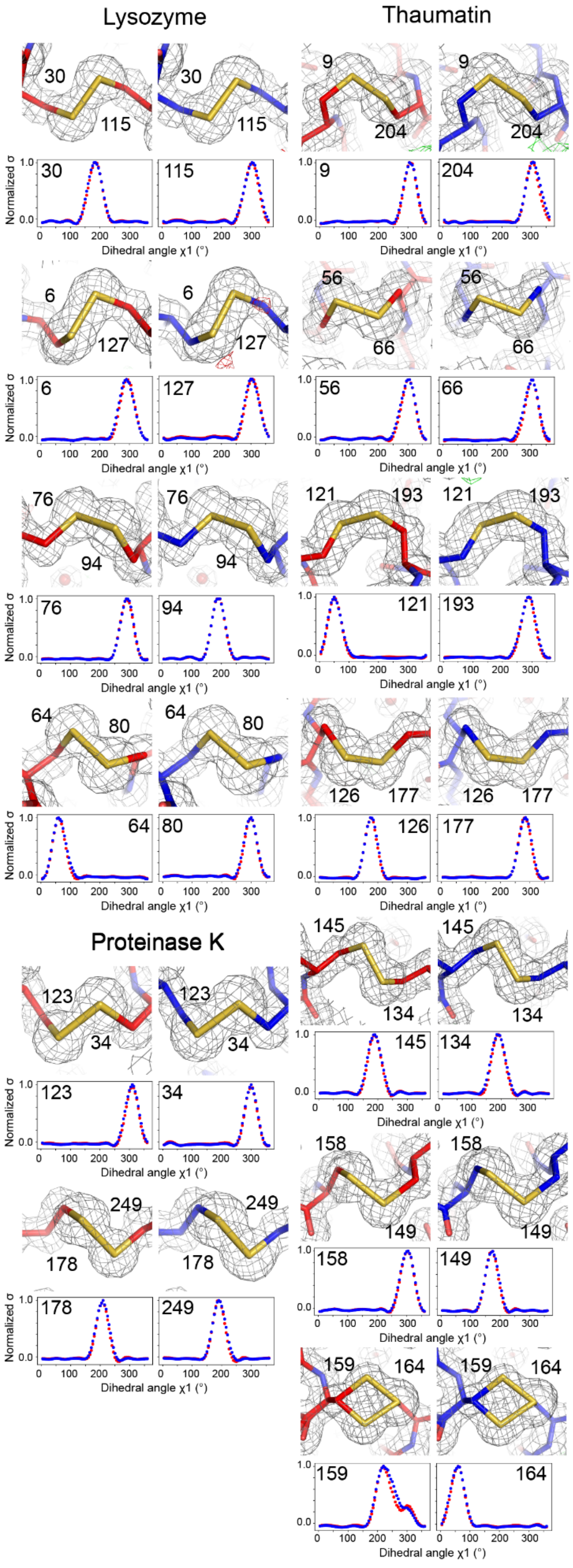

Lysozyme
Thaumatin
Proteinase K
Normalized σ
Dihedral angle χ1 (°)

**Figure S3**. ***Ringer* and electron density analyses indicate limited X-ray damage to disulfide bonds at room temperature**. Electron density maps (2Fo-Fc in grey mesh at a contour level of 1σ, negative and positive Fo-Fc peaks in red and green mesh, respectively, at a contour level of 3σ) and refined structural models (shown in sticks) for the least damaged (red) and most damaged (blue) datasets for lysozyme, proteinase K, and thaumatin (**Tables S1-S3**). The normalized *Ringer* profiles were obtained for each cysteine residue from the least damaged dataset (red) and most damaged dataset (blue) for each protein and $P_{CC}$ were calculated as described in Materials and Methods (see **Table S6** for individual values).

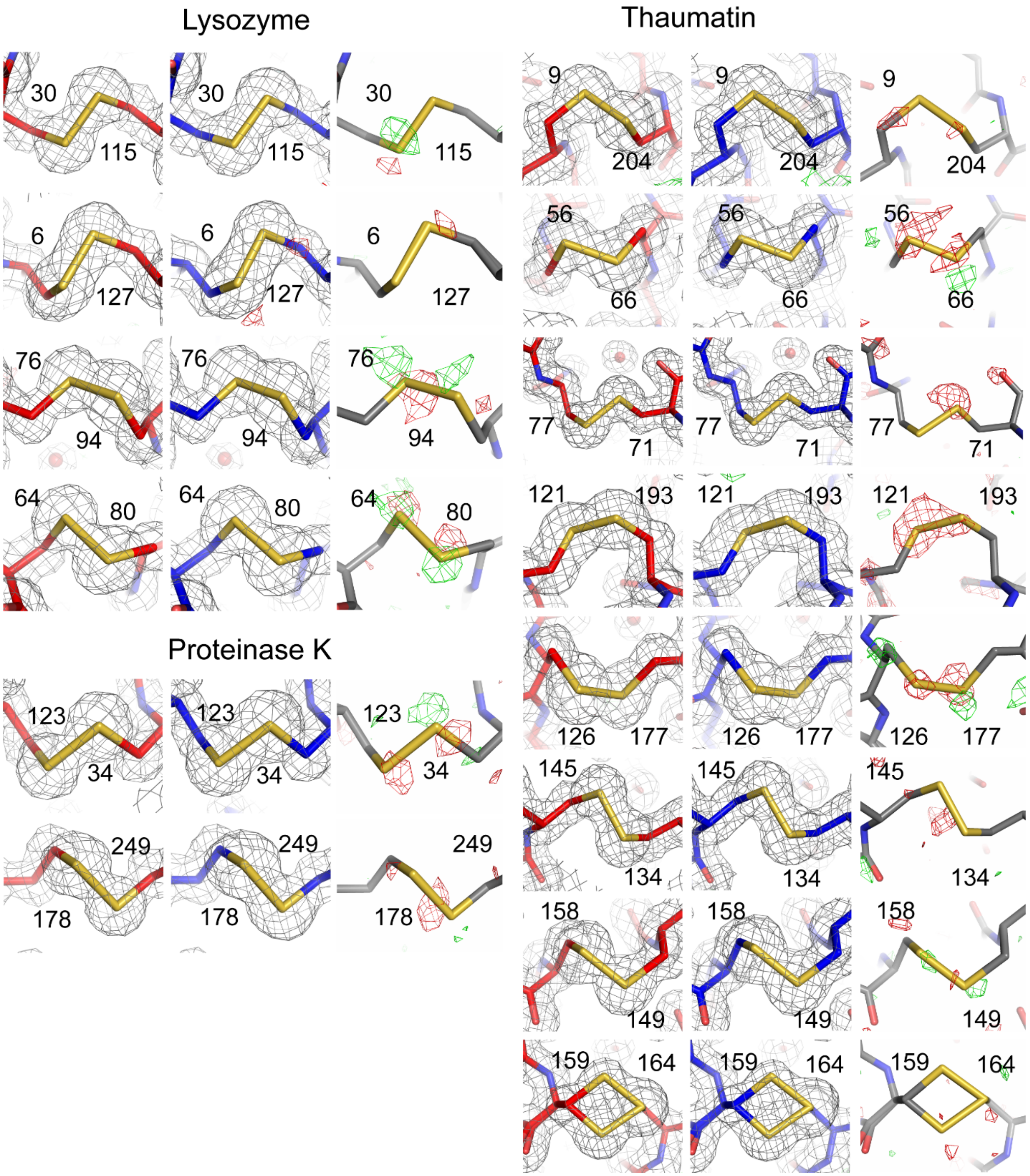

**Figure S4. Difference electron density (Fo-Fo) maps provide evidence for limited X-ray damage to disulfide bonds at room temperature**. Shown are the refined structural models (shown in sticks) for the least damaged (red) and most damaged (blue) datasets for lysozyme, proteinase K, and thaumatin, the associated electron density maps (2Fo-Fc, grey mesh at 1σ, left and middle), and difference electron density maps (Fo-Fo, negative and positive peaks in red and green mesh, respectively, at a contour level of 3σ, right).

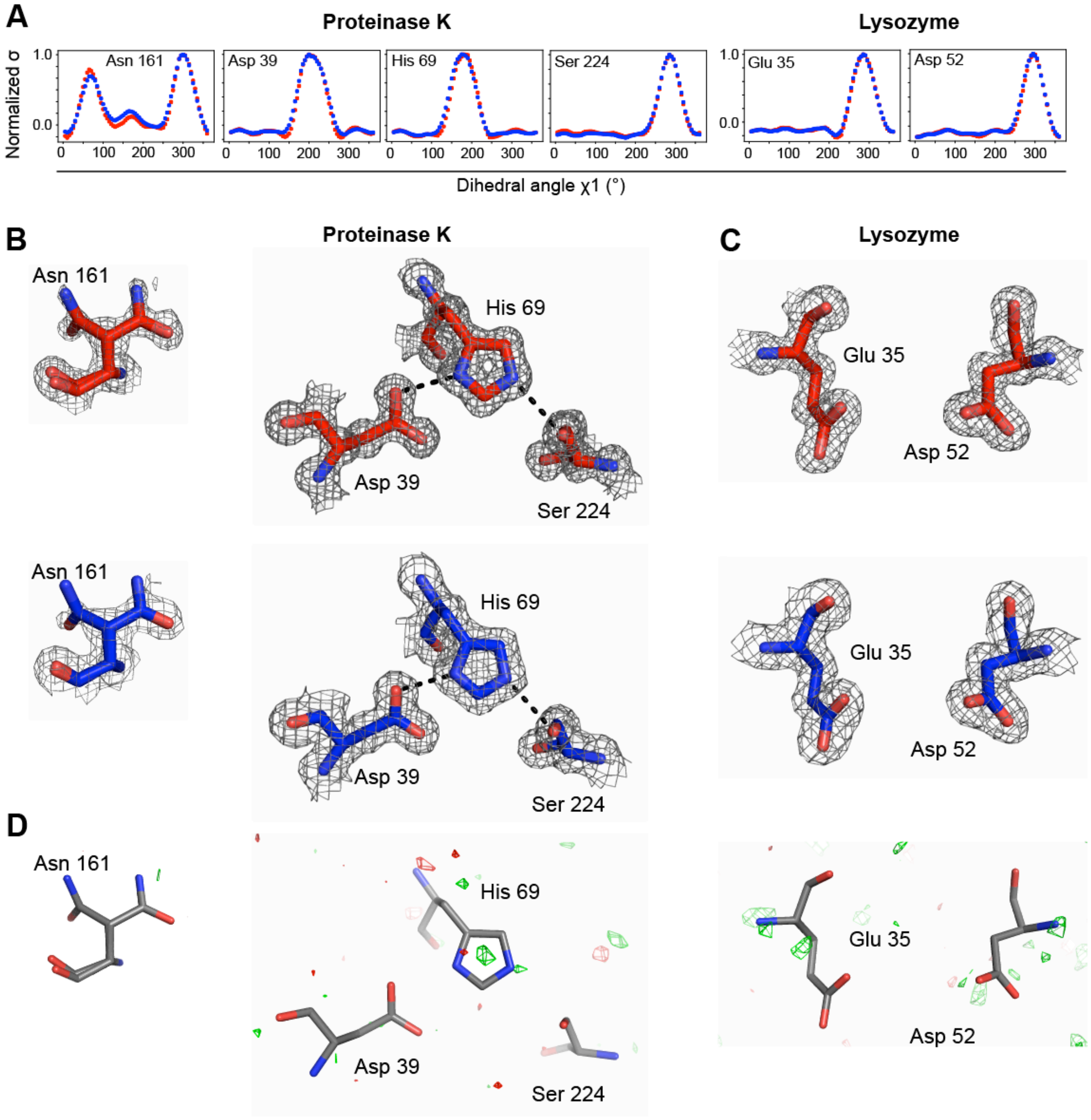

**Figure S5. Absence of substantial X-ray damage to active site residues at room temperature.** (**A**) Normalized *Ringer* profiles for dihedral angles $\chi^1$ obtained from the least and most damaged datasets (red and blue, respectively) (also see **Table S7** for analysis of dihedral angles $\chi^2$ and $P_{CC}$ values). (**B**) Electron density and stick models for the proteinase K oxyanion hole hydrogen bond donor N161 and catalytic triad S224, H69, and D39 and (**C**) the lysozyme catalytic residues E35 and D52. (**B**, **C**) The electron density map (2Fo-Fc in grey mesh at a contour level of 1σ) and refined structural models (sown in sticks) for the least damaged (red) and most damaged (blue) datasets for each protein. (**D**) The Fo-Fo difference electron density maps between the least and most damaged datasets contoured at 3σ (positive peaks in green mesh and negative peaks in red mesh) for proteinase K and lysozyme active site residues shown above in panels **B** and **C**.

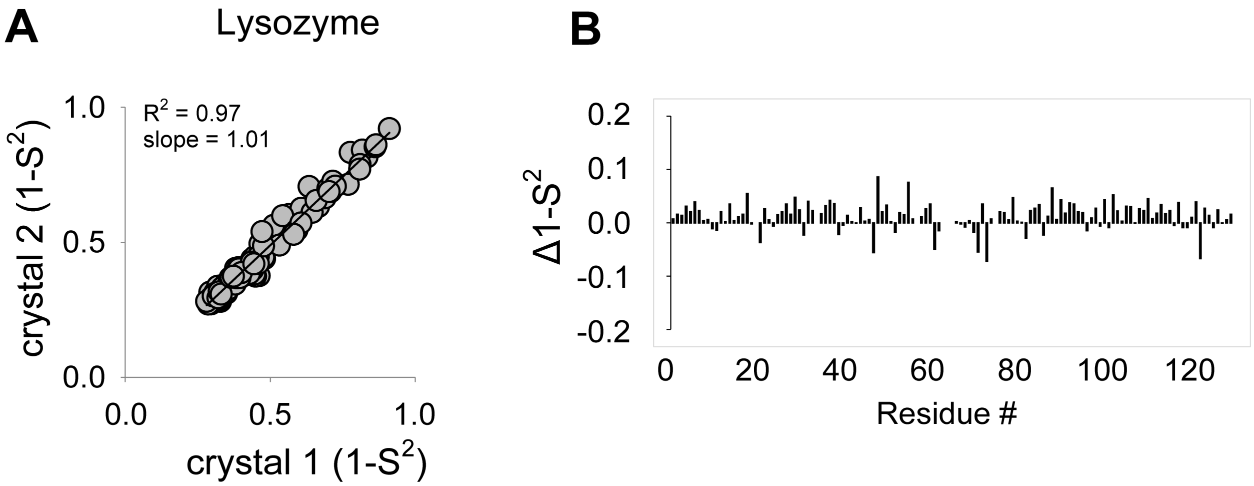

**Figure S6. Comparison of ($1$-$S^2$) values from two independent lysozyme crystals at room temperature indicates highly similar conformational heterogeneity**. (**A**) Correlation plot and (**B**) difference ($1$-$S^2$) as a function of residue number.

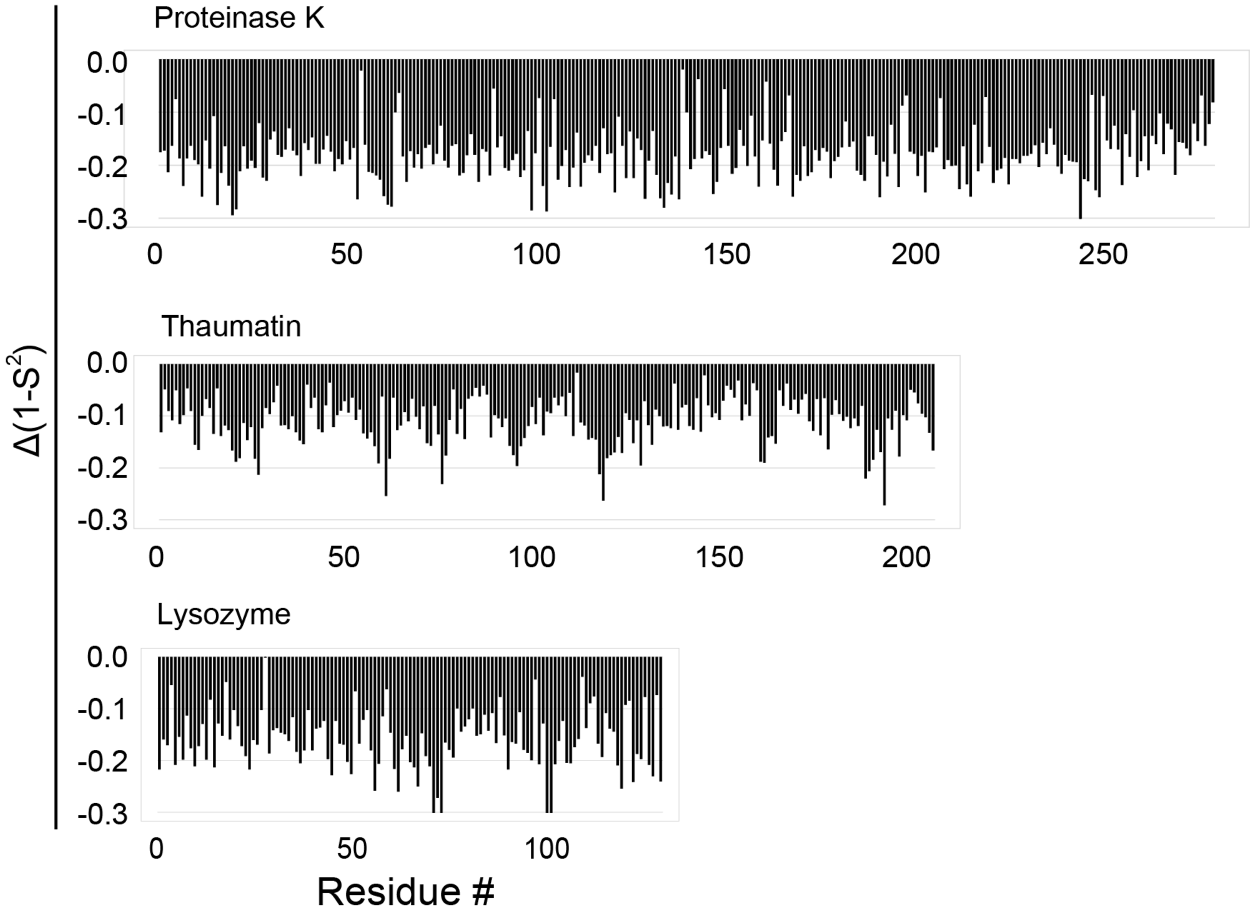

**Figure S7. X-ray damage leads to a modest but measurable increase in conformational heterogeneity as captured by disorder parameters (1-$S^2$) in protein crystals at room temperature.** $\Delta$(1-$S^2$) values between the least and most damaged room temperature datasets for proteinase K, thaumatin, and lysozyme were obtained by subtracting the most damaged from the least damaged (1-$S^2$) values for each residue; the negative (1-$S^2$) values thus indicate higher (1-$S^2$) values obtained from the most damaged dataset. The increased (1-$S^2$) values in the most damaged dataset are not due to resolution differences (see **Figure S8**).

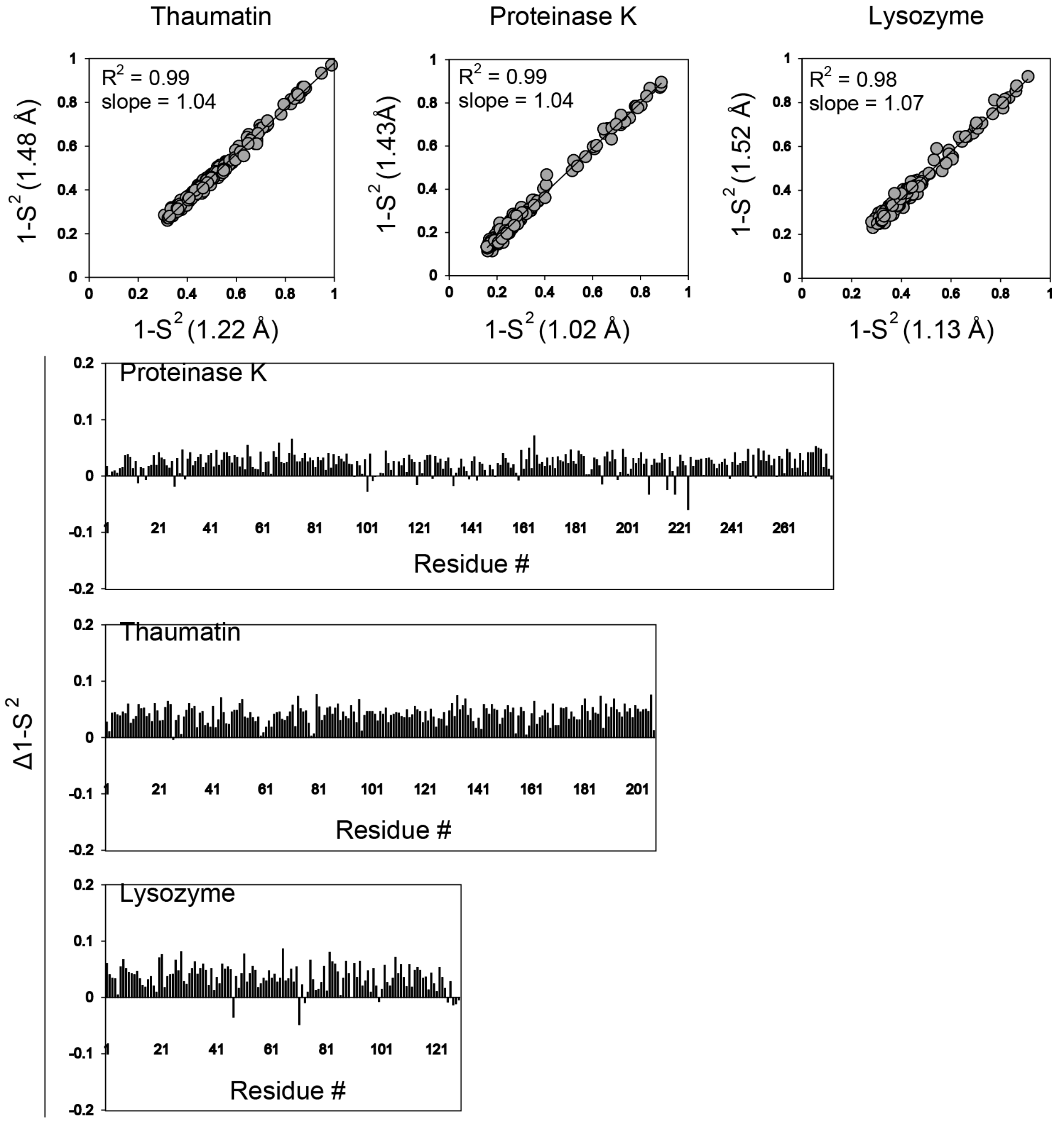

**Figure S8. Decreasing resolution does not lead to major changes in the calculated (1-$S^2$) values**. Comparison of (1-$S^2$) values from the multi-conformer models obtained from the least damaged dataset when the dataset is of optimal resolution (1.02 Å, 1.22 Å and 1.13 Å for proteinase K, thaumatin, and lysozyme, respectively) and when the resolution of the same dataset is cut to match the resolution of the most damaged dataset (1.43 Å, 1.48 Å, and 1.52 Å for proteinase K, thaumatin, and lysozyme, respectively; see Materials and Methods). Correlation plots (top) and difference (1-$S^2$) (bottom) as a function of residue number. The analysis indicates that the lower resolution of the most damaged datasets does not appear as a main factor contributing to the observed increase in (1-$S^2$) values with X-ray damage in Figure S7. The Δ(1-$S^2$) are small and positive in contrast to the larger negative Δ(1-$S^2$) observed in Figure S7.

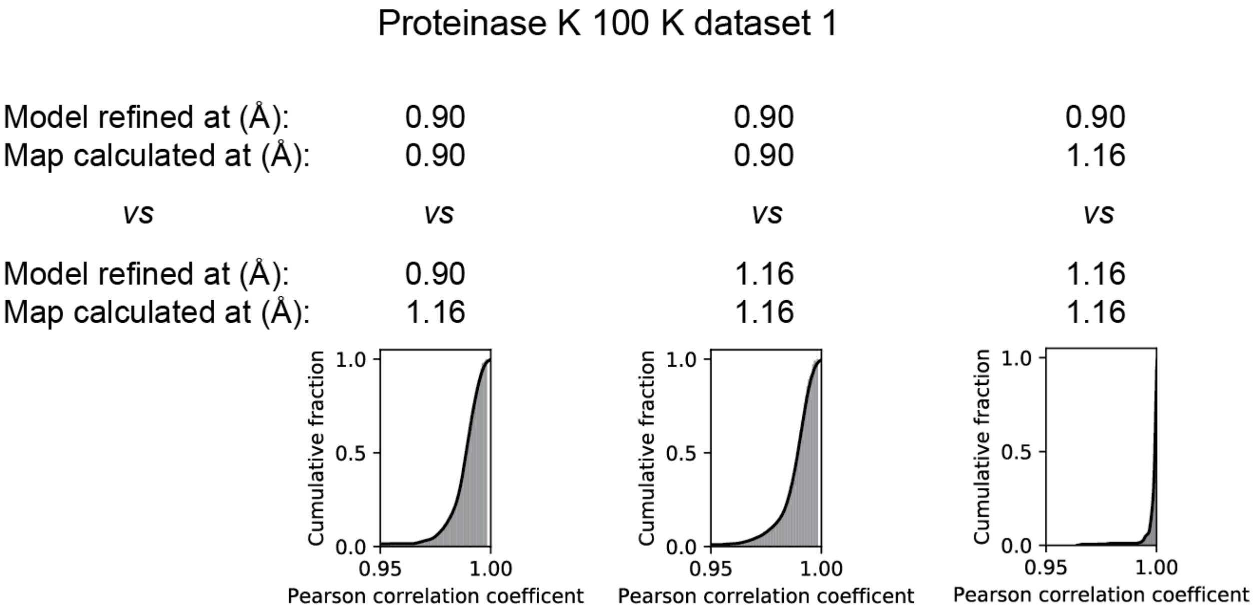

**Figure S9. Matching electron density map resolution is required for accurate *Ringer* analysis and Pearson correlation coefficients calculation.** The resolution of diffraction datasets inevitably decreases with increasing X-ray damage. Thus, for all protein crystals in this work, the least X-ray damaged and increasingly (most) damaged datasets are of different resolutions, with the least damaged dataset being of highest resolution. To evaluate the effect of electron density map resolution on calculating Pearson correlation coefficients ($P_{CC}$) from *Ringer* plots, we compared the *Ringer* profiles obtained from the least damaged proteinase K dataset which was either refined at the optimal resolution of the dataset (0.90 Å) or at a reduced resolution (1.16 Å) to match the resolution of the most damaged dataset and the electron density map used for obtaining *Ringer* plots was either left at the optimal resolution (0.90 Å) or reduced to match the resolution of the most damaged dataset (1.16 Å). Thus, *Ringer* profiles were obtained from three possible combinations of resolutions used for refinement and resolutions used for map calculations: 0.90 Å and 0.90 Å, 0.90 Å and 1.16 Å, and 1.16 Å and 1.16 Å, respectively. The fourth possible combination of 1.16 Å (refinement) and 0.90 Å (map calculation) was not included as this scenario is not relevant for the analysis. The analysis shows that for the same least damaged dataset, artifactually lower $P_{CC}$ values are obtained if the resolutions of the electron density maps used for calculating *Ringer* profiles do not match (left and middle plots of cumulative fraction $P_{CC}$). The plot of cumulative fraction $P_{CC}$ on the right shows that when the resolution of the electron density maps is matched (i.e. 1.16 Å) there is a near perfect agreement between the *Ringer* profiles ($P_{CC}$ is close to 1.0) with small differences originating from small coordinate changes due to the same model being refined against two different resolution datasets. Thus, all *Ringer* comparisons between increasingly damaged datasets have been performed by calculating *Ringer* profiles from electron density maps with matched resolutions – e.g., the for the comparison of the least and most damaged proeinase K cryo datasets, each structural model has been refined at the optimal dataset resolution (0.90 Å and 1.16 Å, respectively) but the *Ringer* profiles were calculated using electron density maps at 1.16 Å.

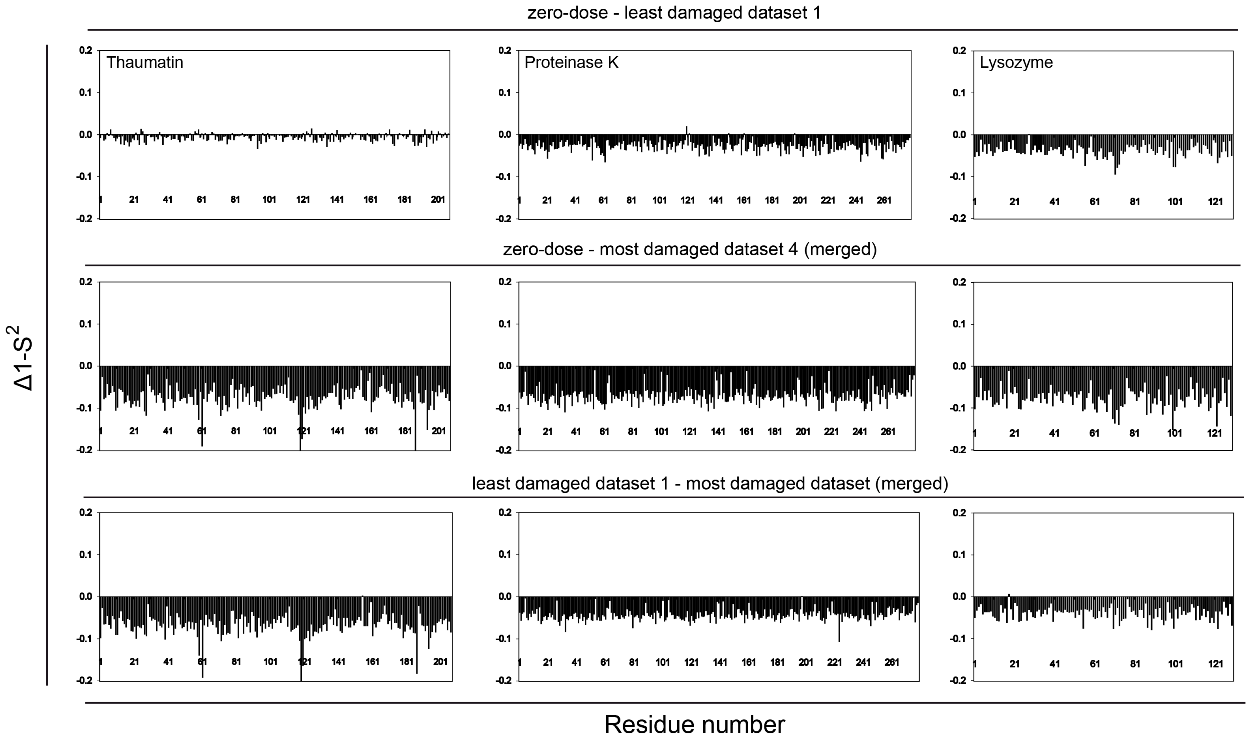

**Figure S10. Comparison of the (1-$S^2$) values obtained from the least damaged and most damaged (merged) room temperature datasets and the (1-$S^2$) values extrapolated to zero-dose.** The figure shows $\Delta$(1-$S^2$) values between extrapolated zero-dose (1-$S^2$) and least damaged (1-$S^2$) (top), extrapolated zero-dose (1-$S^2$) and most damaged (merged) (1-$S^2$) (middle), and least damaged (1-$S^2$) and most damaged (merged) (bottom). Overall, the lowest $\Delta$(1-$S^2$) values are between the zero-dose (1-$S^2$) and least damaged (1-$S^2$), providing confidence in using (1-$S^2$) values for functional analyses if zero-dose (1-$S^2$) values cannot be obtained.

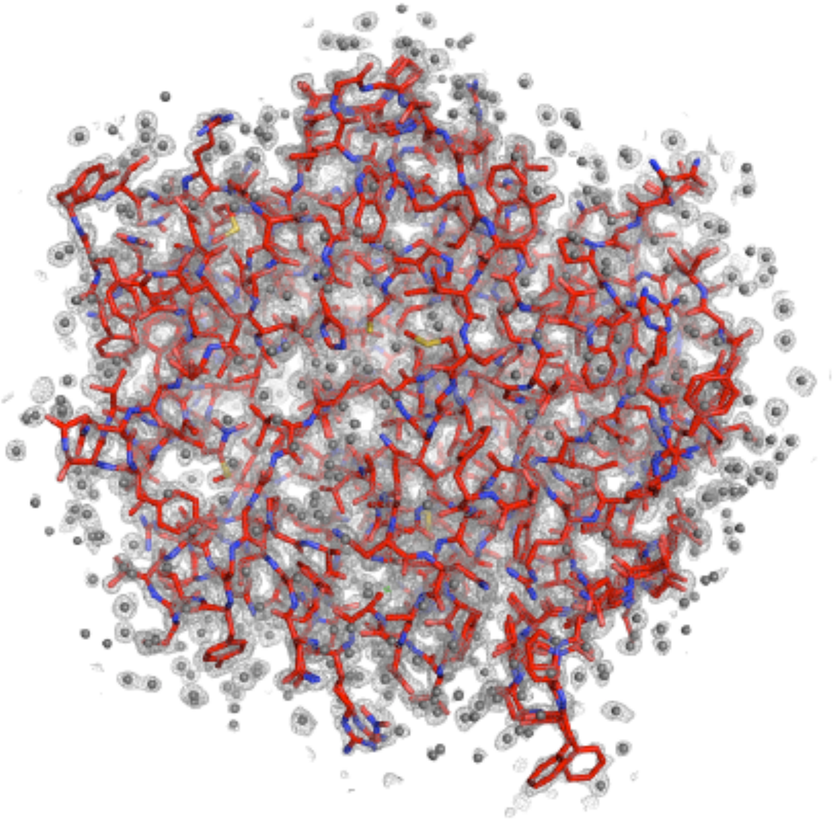

**Figure S11.** 100 K structural model of proteinase K (red sticks) and its electron density map (grey mesh, final 2Fo-Fc electron density map contoured at 1σ) refined using the least damaged dataset (dataset 1, see **Table S21**). Water molecules are shown as grey spheres

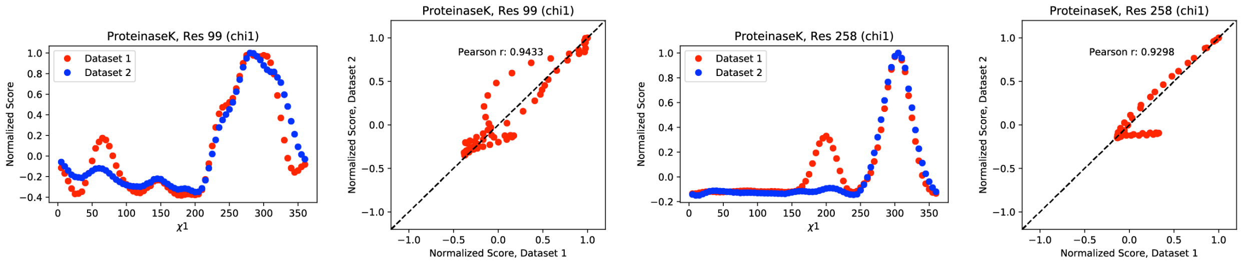

**Figure S12.** Normalized *Ringer* profiles with Pearson correlation coefficients ($P_{CC}$) ≤ 0.95 from proteinase K 100 K data for dataset 1 (red) and dataset 2 (blue). Also shown are the associated correlation plots and respective $P_{CC}$ values.

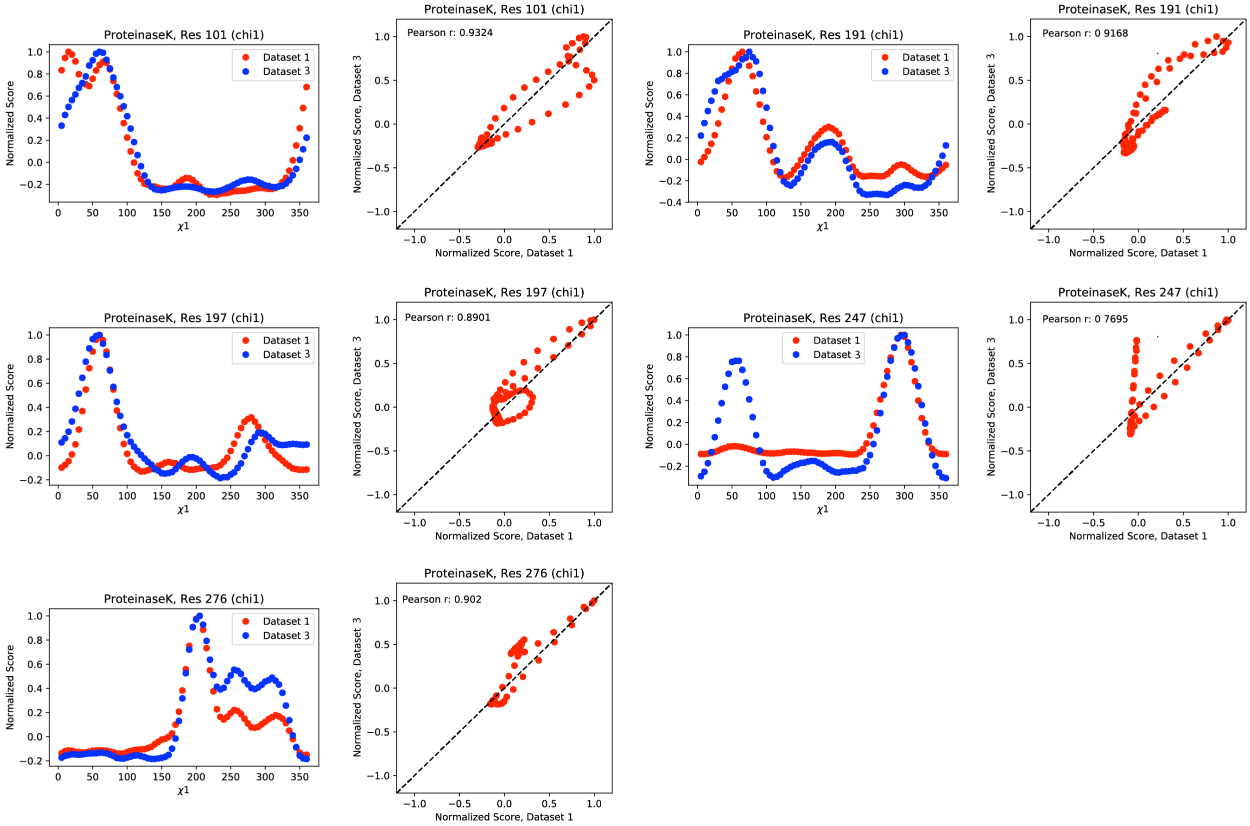

**Figure S13.** Normalized *Ringer* profiles with Pearson correlation coefficients ($P_{CC}$) ≤ 0.95 from proteinase K 100 K data for dataset 1 (red) and dataset 3 (blue). Also shown are the associated correlation plots and respective $P_{CC}$ values.

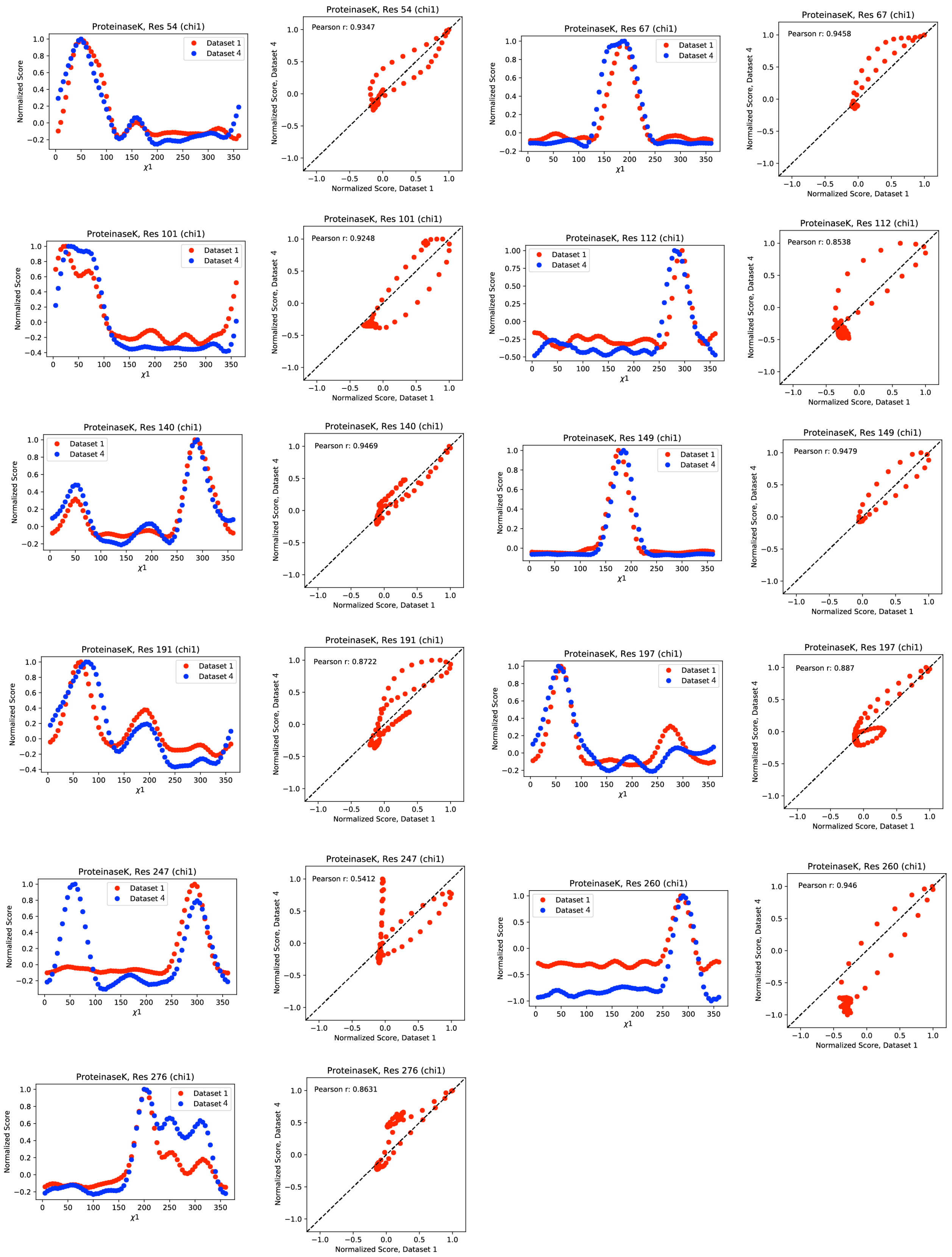

ProteinaseK, Res 54 (chi1)
Pearson r: 0.9347
ProteinaseK, Res 67 (chi1)
Pearson r: 0.9458
ProteinaseK, Res 101 (chi1)
Pearson r: 0.9248
ProteinaseK, Res 112 (chi1)
Pearson r: 0.8538
ProteinaseK, Res 140 (chi1)
Pearson r: 0.9469
ProteinaseK, Res 149 (chi1)
Pearson r: 0.9479
ProteinaseK, Res 191 (chi1)
Pearson r: 0.8722
ProteinaseK, Res 197 (chi1)
Pearson r: 0.887
ProteinaseK, Res 247 (chi1)
Pearson r: 0.5412
ProteinaseK, Res 260 (chi1)
Pearson r: 0.946
ProteinaseK, Res 276 (chi1)
Pearson r: 0.8631
Dataset 1
Dataset 4
Normalized Score
χ1
Normalized Score, Dataset 1
Normalized Score, Dataset 4

**Figure S14.** Normalized *Ringer* profiles with Pearson correlation coefficients ($P_{CC}$) ≤ 0.95 from proteinase K 100 K data for dataset 1 (red) and dataset 4 (blue). Also shown are the associated correlation plots and respective $P_{CC}$ values.

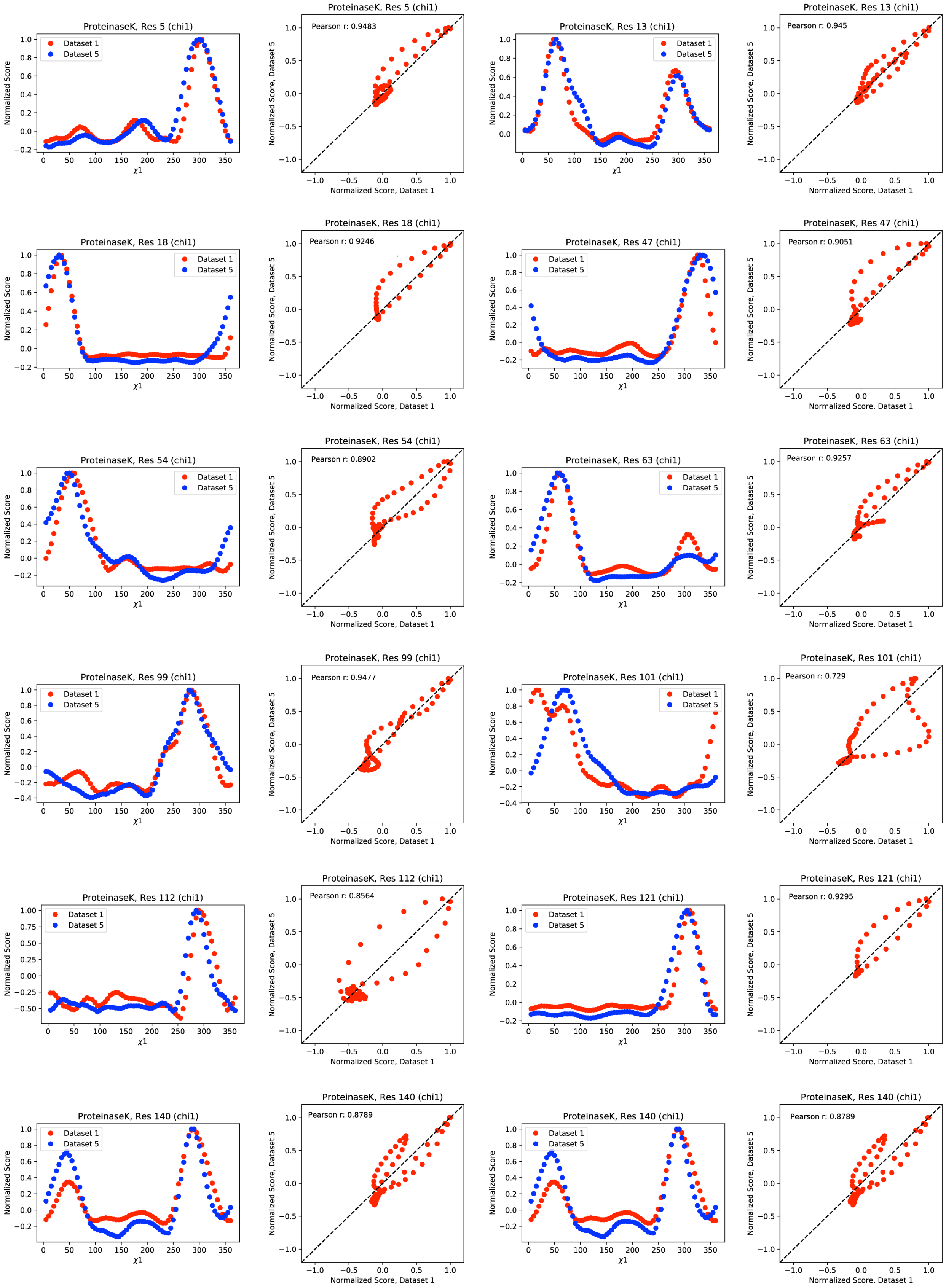
ProteinaseK, Res 5 (chi1)
Dataset 1
Dataset 5
Normalized Score
χ1
ProteinaseK, Res 5 (chi1)
Pearson r: 0.9483
Normalized Score, Dataset 5
Normalized Score, Dataset 1
ProteinaseK, Res 13 (chi1)
Dataset 1
Dataset 5
ProteinaseK, Res 13 (chi1)
Pearson r: 0.945
ProteinaseK, Res 18 (chi1)
Dataset 1
Dataset 5
ProteinaseK, Res 18 (chi1)
Pearson r: 0 9246
ProteinaseK, Res 47 (chi1)
Dataset 1
Dataset 5
ProteinaseK, Res 47 (chi1)
Pearson r: 0.9051
ProteinaseK, Res 54 (chi1)
Dataset 1
Dataset 5
ProteinaseK, Res 54 (chi1)
Pearson r: 0.8902
ProteinaseK, Res 63 (chi1)
Dataset 1
Dataset 5
ProteinaseK, Res 63 (chi1)
Pearson r: 0.9257
ProteinaseK, Res 99 (chi1)
Dataset 1
Dataset 5
ProteinaseK, Res 99 (chi1)
Pearson r: 0.9477
ProteinaseK, Res 101 (chi1)
Dataset 1
Dataset 5
ProteinaseK, Res 101 (chi1)
Pearson r: 0.729
ProteinaseK, Res 112 (chi1)
Dataset 1
Dataset 5
ProteinaseK, Res 112 (chi1)
Pearson r: 0.8564
ProteinaseK, Res 121 (chi1)
Dataset 1
Dataset 5
ProteinaseK, Res 121 (chi1)
Pearson r: 0.9295
ProteinaseK, Res 140 (chi1)
Dataset 1
Dataset 5
ProteinaseK, Res 140 (chi1)
Pearson r: 0.8789
ProteinaseK, Res 140 (chi1)
Dataset 1
Dataset 5
ProteinaseK, Res 140 (chi1)
Pearson r: 0.8789

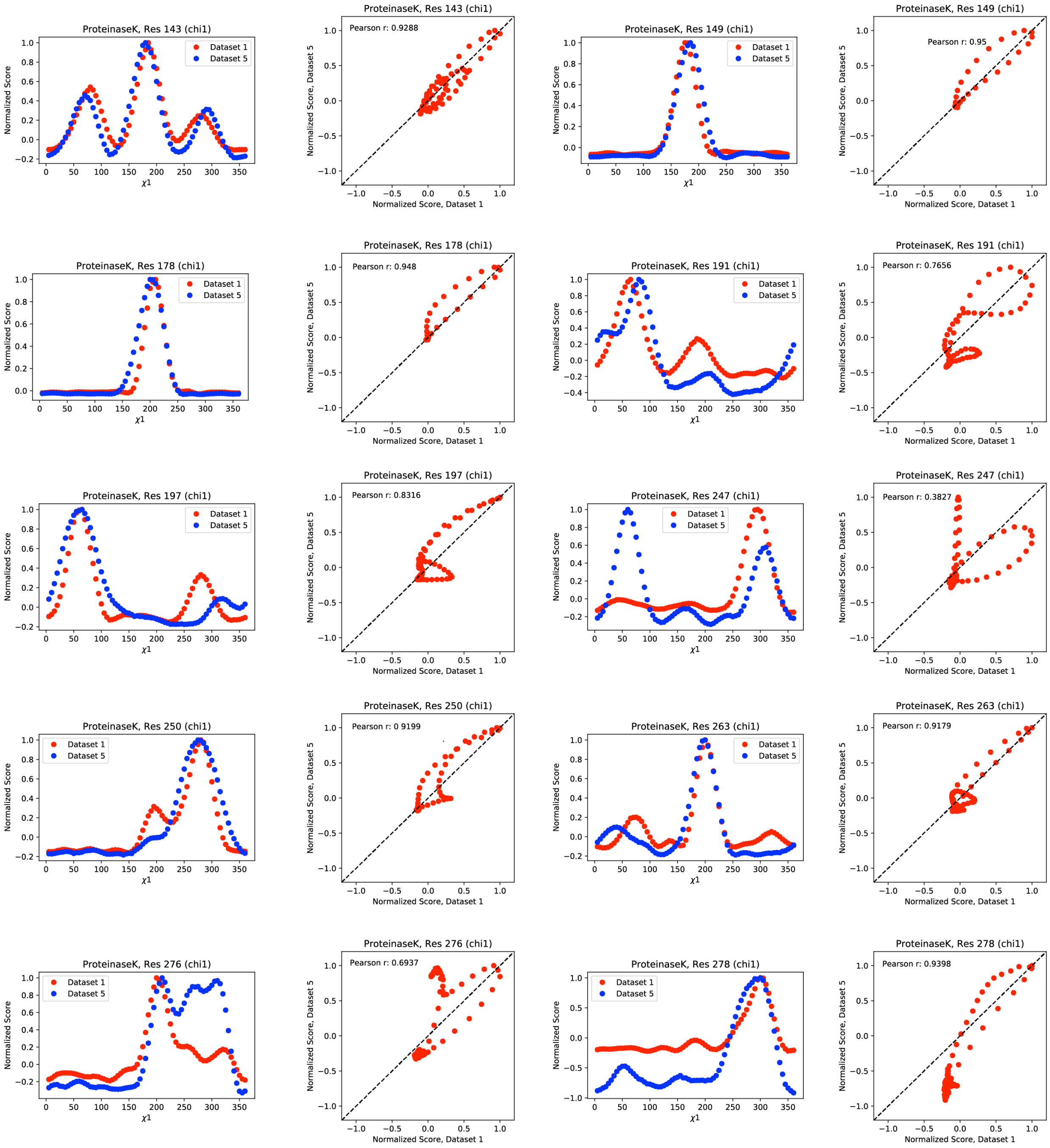

**Figure S15.** Normalized *Ringer* profiles with Pearson correlation coefficients ($P_{CC}$) ≤ 0.95 from proteinase K 100 K data for dataset 1 (red) and dataset 5 (blue). Also shown are the associated correlation plots and respective $P_{CC}$ values.

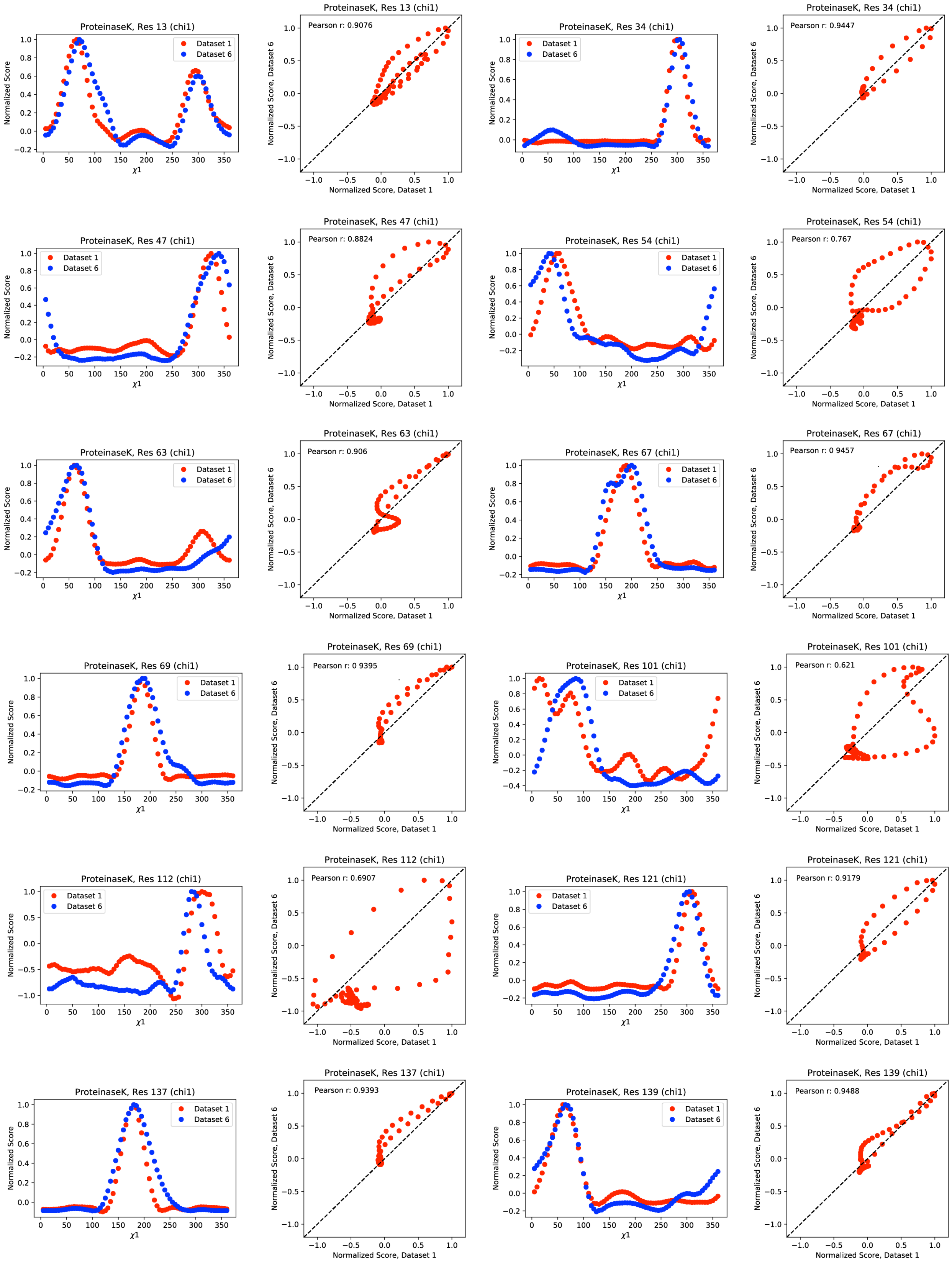
ProteinaseK, Res 13 (chi1)
Dataset 1
Dataset 6
Normalized Score
χ1
ProteinaseK, Res 13 (chi1)
Pearson r: 0.9076
Normalized Score, Dataset 6
Normalized Score, Dataset 1
ProteinaseK, Res 34 (chi1)
ProteinaseK, Res 34 (chi1)
Pearson r: 0.9447
ProteinaseK, Res 47 (chi1)
ProteinaseK, Res 47 (chi1)
Pearson r: 0.8824
ProteinaseK, Res 54 (chi1)
ProteinaseK, Res 54 (chi1)
Pearson r: 0.767
ProteinaseK, Res 63 (chi1)
ProteinaseK, Res 63 (chi1)
Pearson r: 0.906
ProteinaseK, Res 67 (chi1)
ProteinaseK, Res 67 (chi1)
Pearson r: 0 9457
ProteinaseK, Res 69 (chi1)
ProteinaseK, Res 69 (chi1)
Pearson r: 0 9395
ProteinaseK, Res 101 (chi1)
ProteinaseK, Res 101 (chi1)
Pearson r: 0.621
ProteinaseK, Res 112 (chi1)
ProteinaseK, Res 112 (chi1)
Pearson r: 0.6907
ProteinaseK, Res 121 (chi1)
ProteinaseK, Res 121 (chi1)
Pearson r: 0.9179
ProteinaseK, Res 137 (chi1)
ProteinaseK, Res 137 (chi1)
Pearson r: 0.9393
ProteinaseK, Res 139 (chi1)
ProteinaseK, Res 139 (chi1)
Pearson r: 0.9488

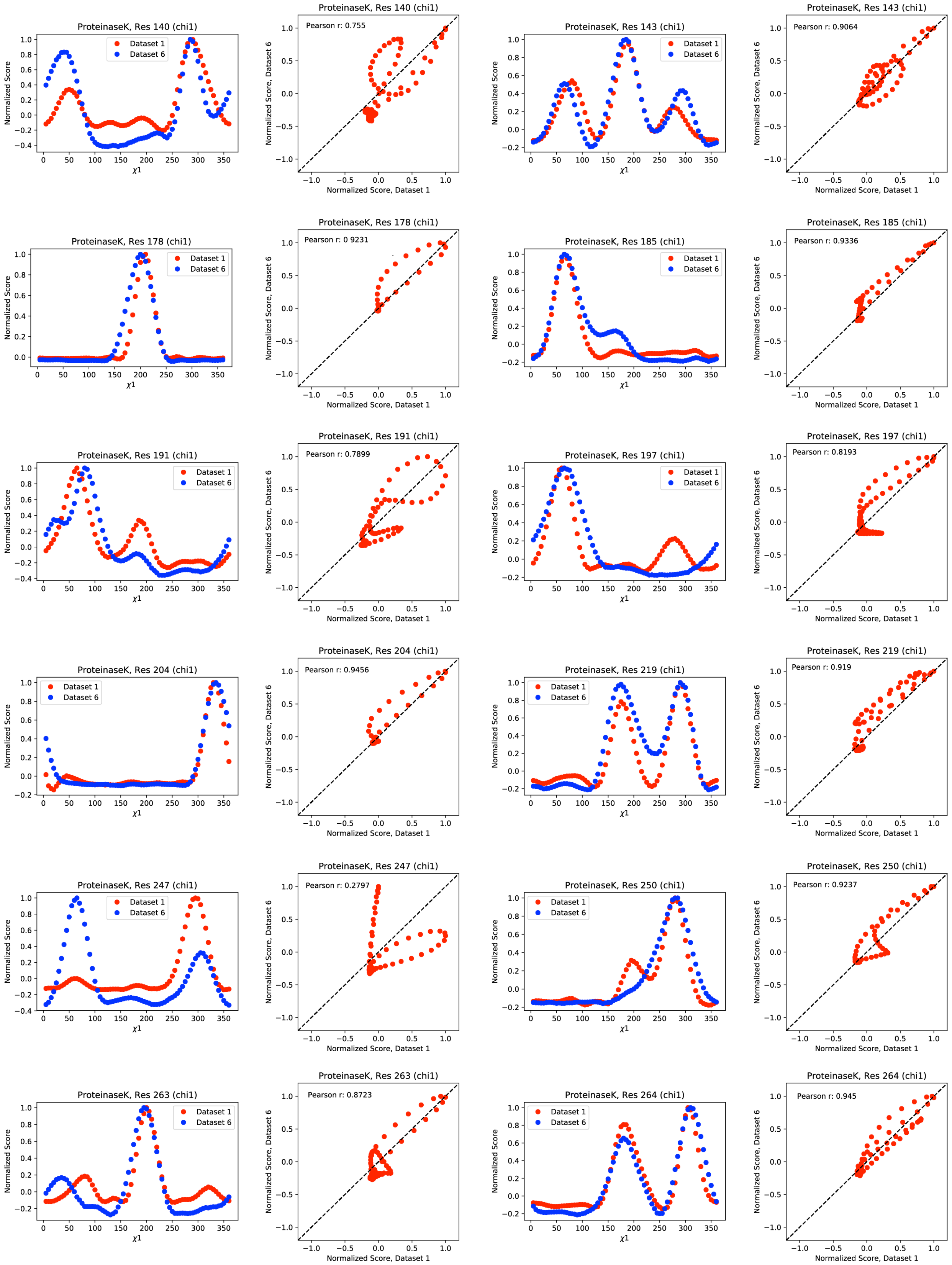

ProteinaseK, Res 140 (chi1)
Dataset 1
Dataset 6
Normalized Score
χ1
ProteinaseK, Res 140 (chi1)
Pearson r: 0.755
Normalized Score, Dataset 6
Normalized Score, Dataset 1
ProteinaseK, Res 143 (chi1)
Dataset 1
Dataset 6
Normalized Score
χ1
ProteinaseK, Res 143 (chi1)
Pearson r: 0.9064
Normalized Score, Dataset 6
Normalized Score, Dataset 1
ProteinaseK, Res 178 (chi1)
Dataset 1
Dataset 6
Normalized Score
χ1
ProteinaseK, Res 178 (chi1)
Pearson r: 0 9231
Normalized Score, Dataset 6
Normalized Score, Dataset 1
ProteinaseK, Res 185 (chi1)
Dataset 1
Dataset 6
Normalized Score
χ1
ProteinaseK, Res 185 (chi1)
Pearson r: 0.9336
Normalized Score, Dataset 6
Normalized Score, Dataset 1
ProteinaseK, Res 191 (chi1)
Dataset 1
Dataset 6
Normalized Score
χ1
ProteinaseK, Res 191 (chi1)
Pearson r: 0.7899
Normalized Score, Dataset 6
Normalized Score, Dataset 1
ProteinaseK, Res 197 (chi1)
Dataset 1
Dataset 6
Normalized Score
χ1
ProteinaseK, Res 197 (chi1)
Pearson r: 0.8193
Normalized Score, Dataset 6
Normalized Score, Dataset 1
ProteinaseK, Res 204 (chi1)
Dataset 1
Dataset 6
Normalized Score
χ1
ProteinaseK, Res 204 (chi1)
Pearson r: 0.9456
Normalized Score, Dataset 6
Normalized Score, Dataset 1
ProteinaseK, Res 219 (chi1)
Dataset 1
Dataset 6
Normalized Score
χ1
ProteinaseK, Res 219 (chi1)
Pearson r: 0.919
Normalized Score, Dataset 6
Normalized Score, Dataset 1
ProteinaseK, Res 247 (chi1)
Dataset 1
Dataset 6
Normalized Score
χ1
ProteinaseK, Res 247 (chi1)
Pearson r: 0.2797
Normalized Score, Dataset 6
Normalized Score, Dataset 1
ProteinaseK, Res 250 (chi1)
Dataset 1
Dataset 6
Normalized Score
χ1
ProteinaseK, Res 250 (chi1)
Pearson r: 0.9237
Normalized Score, Dataset 6
Normalized Score, Dataset 1
ProteinaseK, Res 263 (chi1)
Dataset 1
Dataset 6
Normalized Score
χ1
ProteinaseK, Res 263 (chi1)
Pearson r: 0.8723
Normalized Score, Dataset 6
Normalized Score, Dataset 1
ProteinaseK, Res 264 (chi1)
Dataset 1
Dataset 6
Normalized Score
χ1
ProteinaseK, Res 264 (chi1)
Pearson r: 0.945
Normalized Score, Dataset 6
Normalized Score, Dataset 1

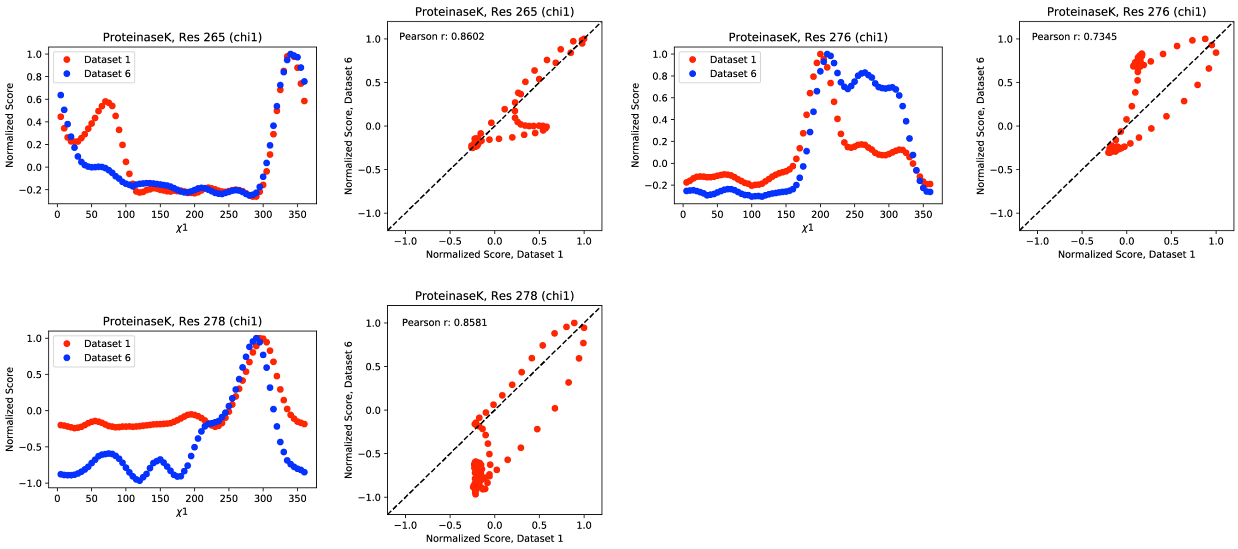

**Figure S16.** Normalized *Ringer* profiles with Pearson correlation coefficients ($P_{CC}$) ≤ 0.95 from proteinase K 100 K data for dataset 1 (red) and dataset 6 (blue). Also shown are the associated correlation plots and respective $P_{CC}$ values.

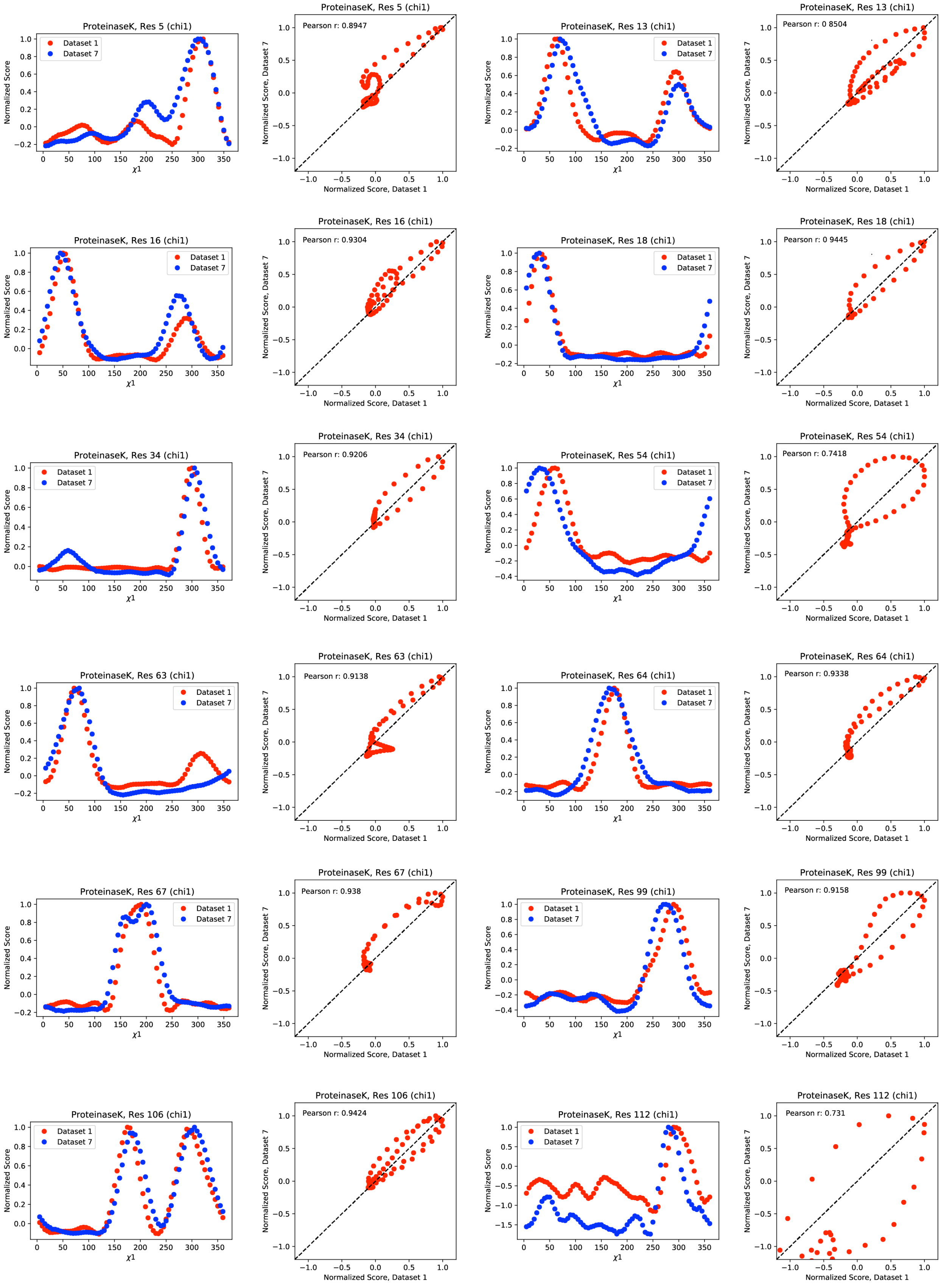
ProteinaseK, Res 5 (chi1)
Dataset 1
Dataset 7
Normalized Score
χ1
ProteinaseK, Res 5 (chi1)
Pearson r: 0.8947
Normalized Score, Dataset 7
Normalized Score, Dataset 1
ProteinaseK, Res 13 (chi1)
Dataset 1
Dataset 7
Normalized Score
χ1
ProteinaseK, Res 13 (chi1)
Pearson r: 0 8504
Normalized Score, Dataset 7
Normalized Score, Dataset 1
ProteinaseK, Res 16 (chi1)
Dataset 1
Dataset 7
Normalized Score
χ1
ProteinaseK, Res 16 (chi1)
Pearson r: 0.9304
Normalized Score, Dataset 7
Normalized Score, Dataset 1
ProteinaseK, Res 18 (chi1)
Dataset 1
Dataset 7
Normalized Score
χ1
ProteinaseK, Res 18 (chi1)
Pearson r: 0 9445
Normalized Score, Dataset 7
Normalized Score, Dataset 1
ProteinaseK, Res 34 (chi1)
Dataset 1
Dataset 7
Normalized Score
χ1
ProteinaseK, Res 34 (chi1)
Pearson r: 0.9206
Normalized Score, Dataset 7
Normalized Score, Dataset 1
ProteinaseK, Res 54 (chi1)
Dataset 1
Dataset 7
Normalized Score
χ1
ProteinaseK, Res 54 (chi1)
Pearson r: 0.7418
Normalized Score, Dataset 7
Normalized Score, Dataset 1
ProteinaseK, Res 63 (chi1)
Dataset 1
Dataset 7
Normalized Score
χ1
ProteinaseK, Res 63 (chi1)
Pearson r: 0.9138
Normalized Score, Dataset 7
Normalized Score, Dataset 1
ProteinaseK, Res 64 (chi1)
Dataset 1
Dataset 7
Normalized Score
χ1
ProteinaseK, Res 64 (chi1)
Pearson r: 0.9338
Normalized Score, Dataset 7
Normalized Score, Dataset 1
ProteinaseK, Res 67 (chi1)
Dataset 1
Dataset 7
Normalized Score
χ1
ProteinaseK, Res 67 (chi1)
Pearson r: 0.938
Normalized Score, Dataset 7
Normalized Score, Dataset 1
ProteinaseK, Res 99 (chi1)
Dataset 1
Dataset 7
Normalized Score
χ1
ProteinaseK, Res 99 (chi1)
Pearson r: 0.9158
Normalized Score, Dataset 7
Normalized Score, Dataset 1
ProteinaseK, Res 106 (chi1)
Dataset 1
Dataset 7
Normalized Score
χ1
ProteinaseK, Res 106 (chi1)
Pearson r: 0.9424
Normalized Score, Dataset 7
Normalized Score, Dataset 1
ProteinaseK, Res 112 (chi1)
Dataset 1
Dataset 7
Normalized Score
χ1
ProteinaseK, Res 112 (chi1)
Pearson r: 0.731
Normalized Score, Dataset 7
Normalized Score, Dataset 1

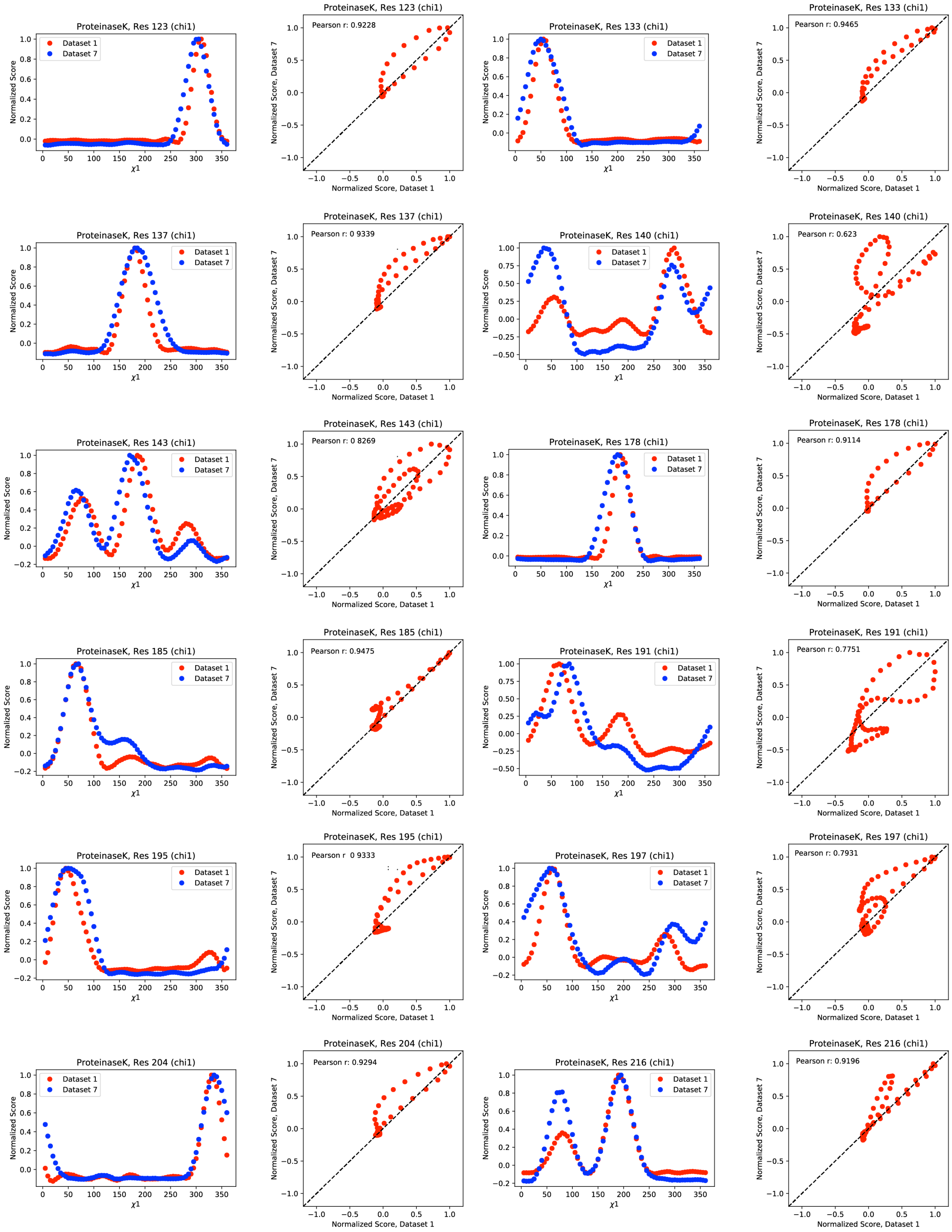
ProteinaseK, Res 123 (chi1)
Dataset 1
Dataset 7
Normalized Score
χ1
ProteinaseK, Res 123 (chi1)
Pearson r: 0.9228
Normalized Score, Dataset 7
Normalized Score, Dataset 1
ProteinaseK, Res 133 (chi1)
ProteinaseK, Res 133 (chi1)
Pearson r: 0.9465
ProteinaseK, Res 137 (chi1)
ProteinaseK, Res 137 (chi1)
Pearson r: 0 9339
ProteinaseK, Res 140 (chi1)
ProteinaseK, Res 140 (chi1)
Pearson r: 0.623
ProteinaseK, Res 143 (chi1)
ProteinaseK, Res 143 (chi1)
Pearson r: 0 8269
ProteinaseK, Res 178 (chi1)
ProteinaseK, Res 178 (chi1)
Pearson r: 0.9114
ProteinaseK, Res 185 (chi1)
ProteinaseK, Res 185 (chi1)
Pearson r: 0.9475
ProteinaseK, Res 191 (chi1)
ProteinaseK, Res 191 (chi1)
Pearson r: 0.7751
ProteinaseK, Res 195 (chi1)
ProteinaseK, Res 195 (chi1)
Pearson r 0 9333
ProteinaseK, Res 197 (chi1)
ProteinaseK, Res 197 (chi1)
Pearson r: 0.7931
ProteinaseK, Res 204 (chi1)
ProteinaseK, Res 204 (chi1)
Pearson r: 0.9294
ProteinaseK, Res 216 (chi1)
ProteinaseK, Res 216 (chi1)
Pearson r: 0.9196

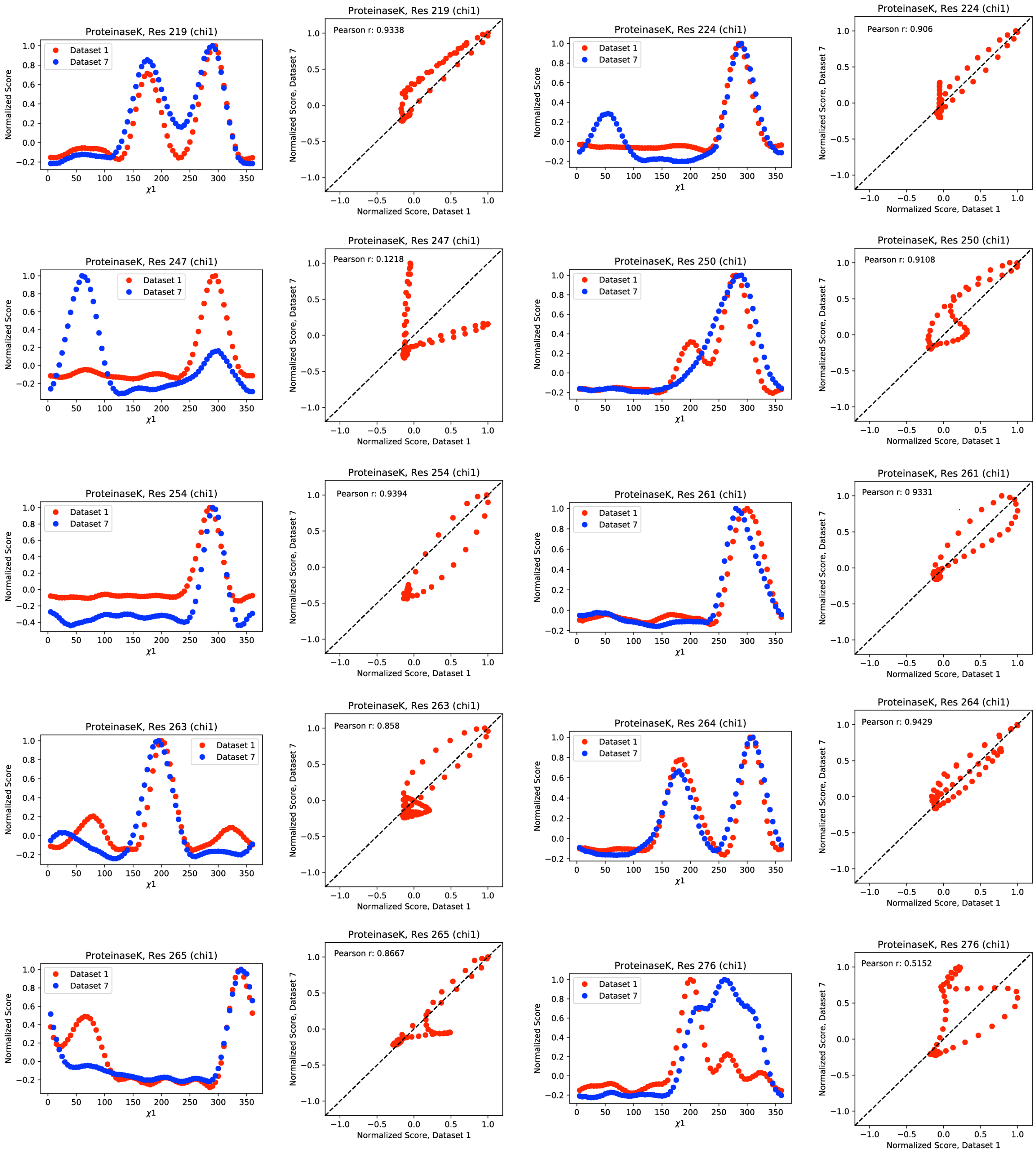

**Figure S17.** Normalized *Ringer* profiles with Pearson correlation coefficients ($P_{CC}$) ≤ 0.95 from proteinase K 100 K data for dataset 1 (red) and dataset 7 (blue). Also shown are the associated correlation plots and respective $P_{CC}$ values.

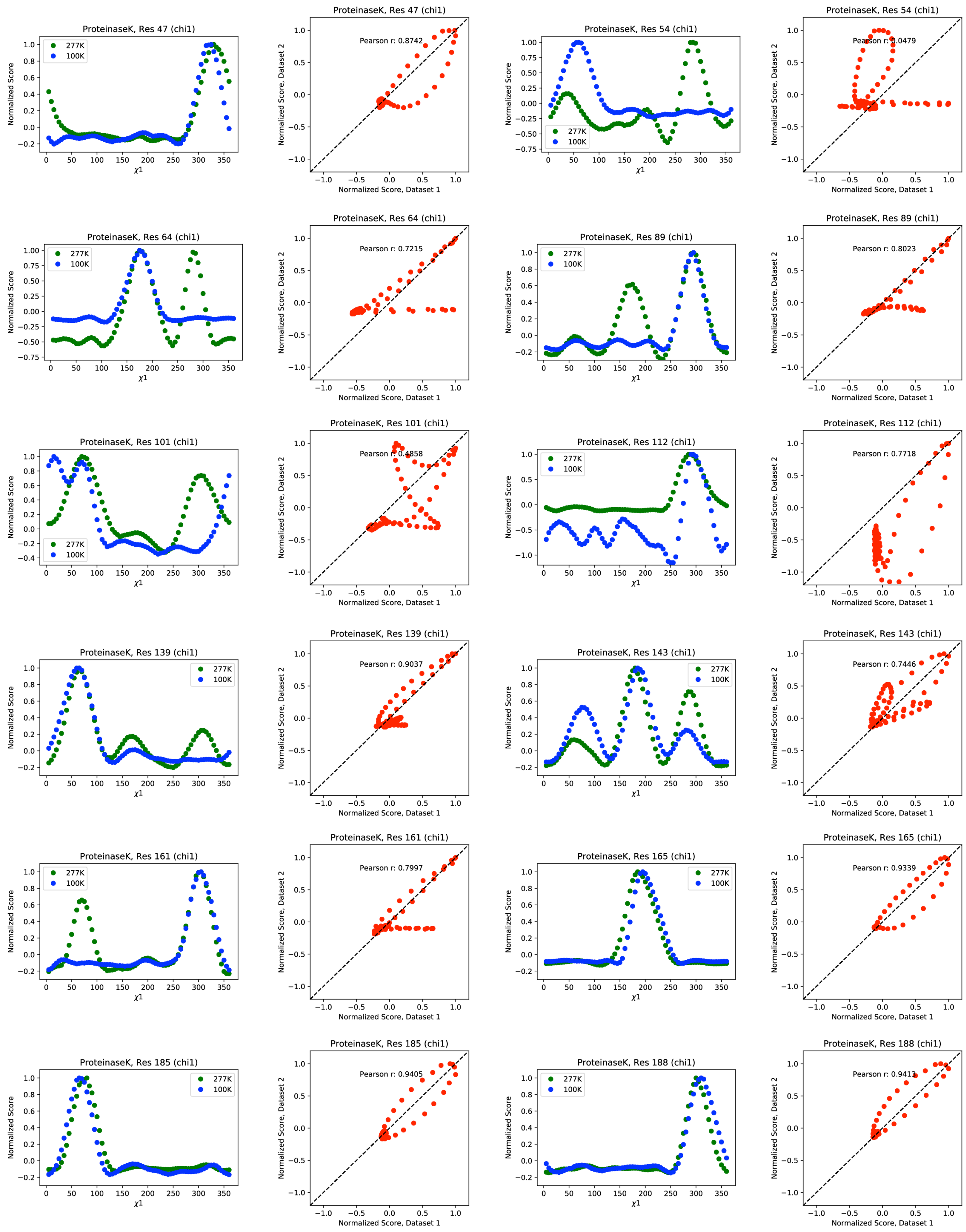
ProteinaseK, Res 47 (chi1)
277K
100K
Normalized Score
$\chi 1$
ProteinaseK, Res 47 (chi1)
Pearson r: 0.8742
Normalized Score, Dataset 1
Normalized Score, Dataset 2
ProteinaseK, Res 54 (chi1)
ProteinaseK, Res 54 (chi1)
Pearson r: 0.0479
ProteinaseK, Res 64 (chi1)
ProteinaseK, Res 64 (chi1)
Pearson r: 0.7215
ProteinaseK, Res 89 (chi1)
ProteinaseK, Res 89 (chi1)
Pearson r: 0.8023
ProteinaseK, Res 101 (chi1)
ProteinaseK, Res 101 (chi1)
Pearson r: 0.4858
ProteinaseK, Res 112 (chi1)
ProteinaseK, Res 112 (chi1)
Pearson r: 0.7718
ProteinaseK, Res 139 (chi1)
ProteinaseK, Res 139 (chi1)
Pearson r: 0.9037
ProteinaseK, Res 143 (chi1)
ProteinaseK, Res 143 (chi1)
Pearson r: 0.7446
ProteinaseK, Res 161 (chi1)
ProteinaseK, Res 161 (chi1)
Pearson r: 0.7997
ProteinaseK, Res 165 (chi1)
ProteinaseK, Res 165 (chi1)
Pearson r: 0.9339
ProteinaseK, Res 185 (chi1)
ProteinaseK, Res 185 (chi1)
Pearson r: 0.9405
ProteinaseK, Res 188 (chi1)
ProteinaseK, Res 188 (chi1)
Pearson r: 0.9413

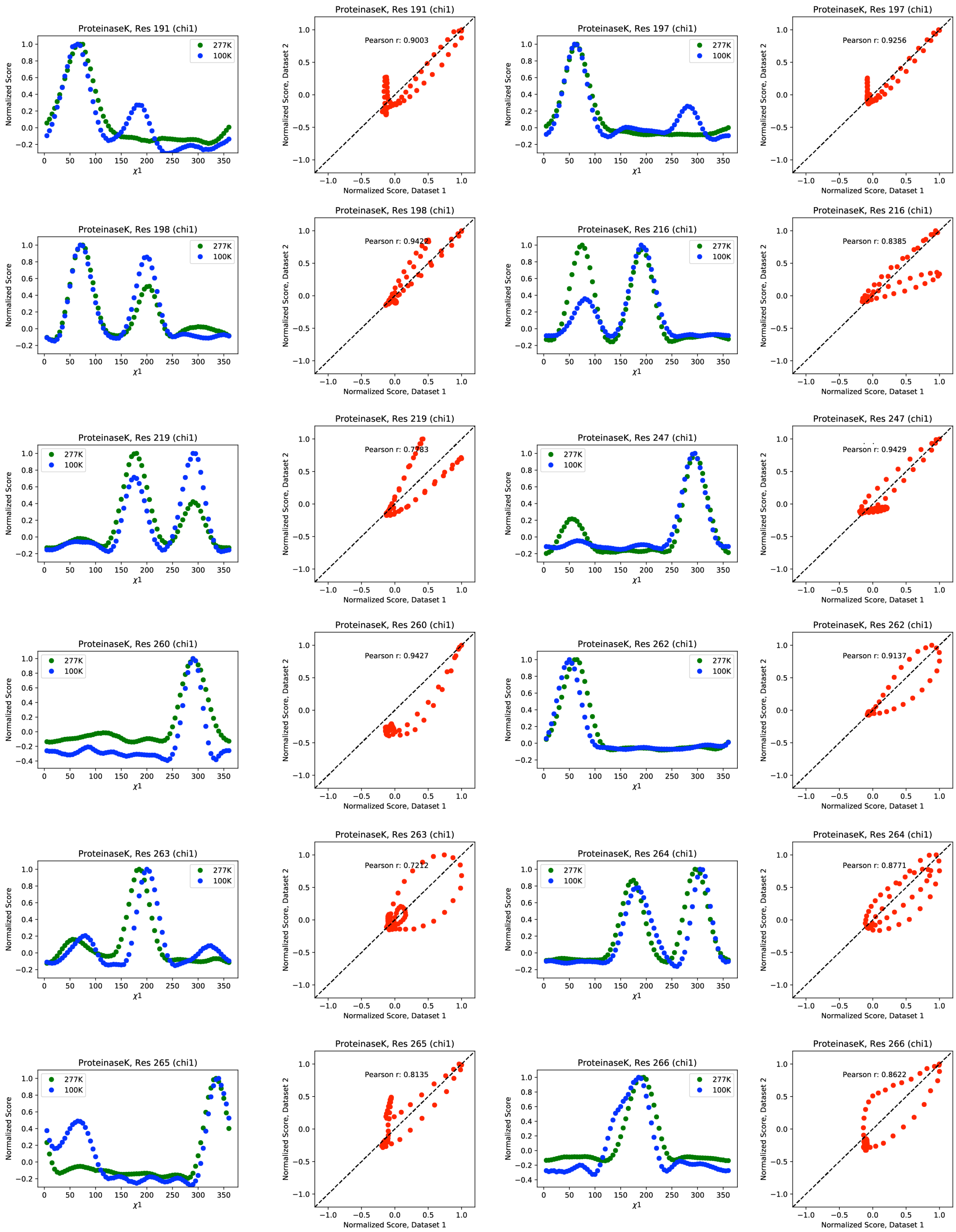
ProteinaseK, Res 191 (chi1)
ProteinaseK, Res 191 (chi1)
Pearson r: 0.9003
ProteinaseK, Res 197 (chi1)
ProteinaseK, Res 197 (chi1)
Pearson r: 0.9256
ProteinaseK, Res 198 (chi1)
ProteinaseK, Res 198 (chi1)
Pearson r: 0.9422
ProteinaseK, Res 216 (chi1)
ProteinaseK, Res 216 (chi1)
Pearson r: 0.8385
ProteinaseK, Res 219 (chi1)
ProteinaseK, Res 219 (chi1)
Pearson r: 0.7783
ProteinaseK, Res 247 (chi1)
ProteinaseK, Res 247 (chi1)
Pearson r: 0.9429
ProteinaseK, Res 260 (chi1)
ProteinaseK, Res 260 (chi1)
Pearson r: 0.9427
ProteinaseK, Res 262 (chi1)
ProteinaseK, Res 262 (chi1)
Pearson r: 0.9137
ProteinaseK, Res 263 (chi1)
ProteinaseK, Res 263 (chi1)
Pearson r: 0.7212
ProteinaseK, Res 264 (chi1)
ProteinaseK, Res 264 (chi1)
Pearson r: 0.8771
ProteinaseK, Res 265 (chi1)
ProteinaseK, Res 265 (chi1)
Pearson r: 0.8135
ProteinaseK, Res 266 (chi1)
ProteinaseK, Res 266 (chi1)
Pearson r: 0.8622
277K
100K
Normalized Score
χ1
Normalized Score, Dataset 1
Normalized Score, Dataset 2

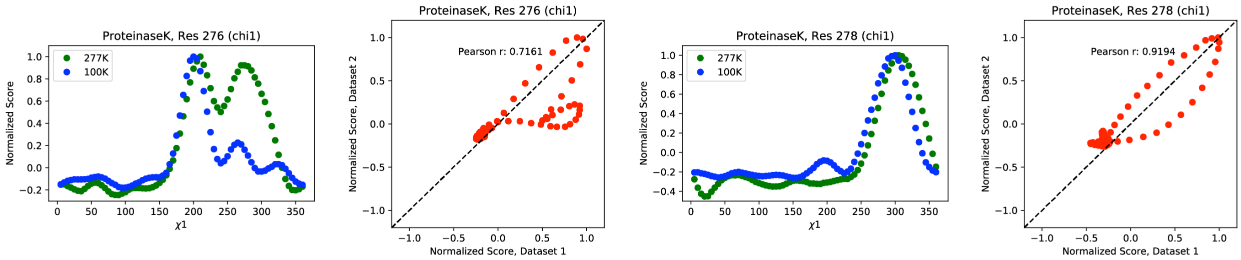

**Figure S18.** Normalized *Ringer* profiles for the proteinase K 277 K least damaged dataset 1 (green) and the 100 K least damaged dataset 1 (blue) with Pearson correlation coefficients ($P_{CC}$) ≤ 0.95. Also shown are the associated correlation plots and respective $P_{CC}$ values.

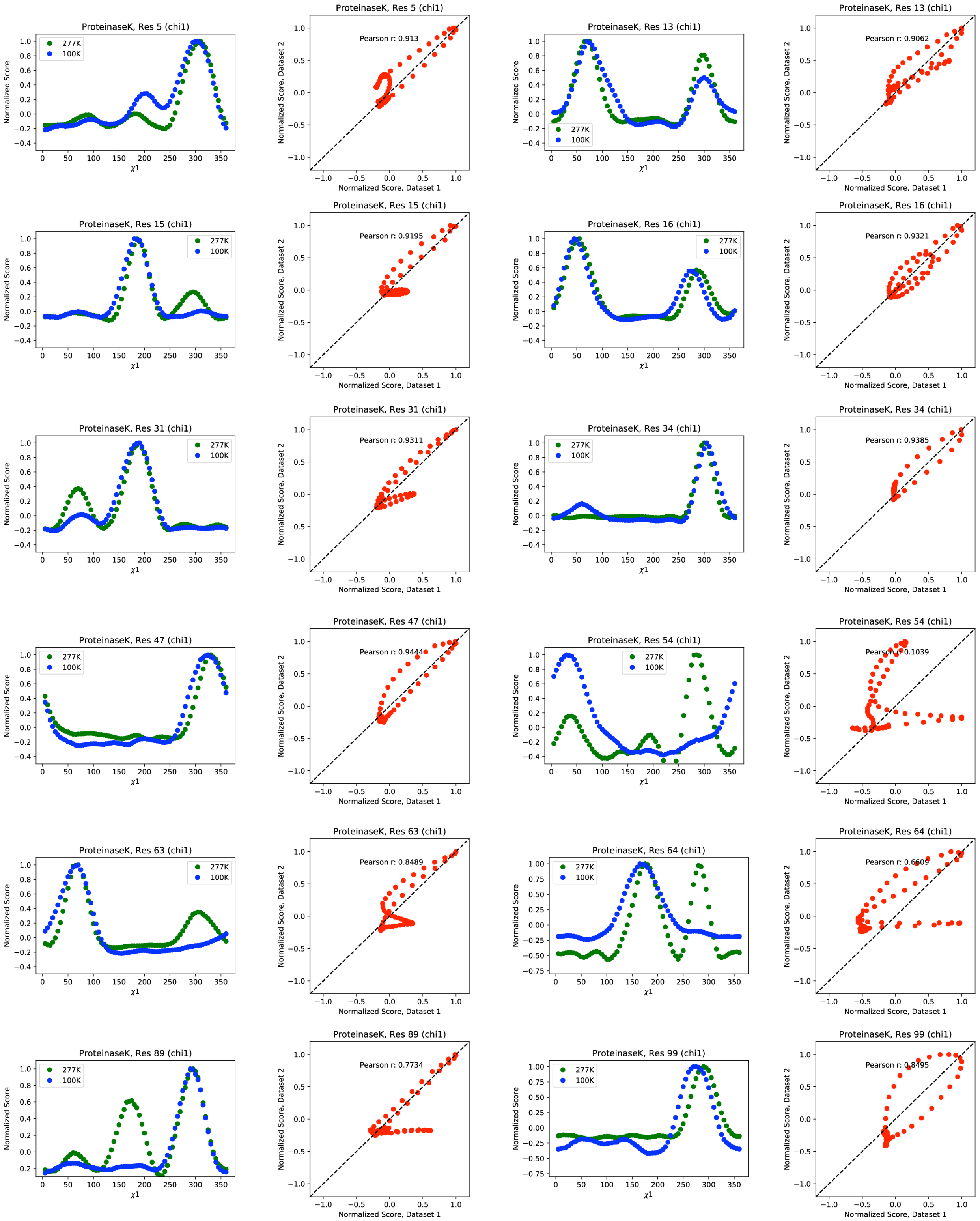

ProteinaseK, Res 5 (chi1)
ProteinaseK, Res 5 (chi1)
Pearson r: 0.913
ProteinaseK, Res 13 (chi1)
ProteinaseK, Res 13 (chi1)
Pearson r: 0.9062
ProteinaseK, Res 15 (chi1)
ProteinaseK, Res 15 (chi1)
Pearson r: 0.9195
ProteinaseK, Res 16 (chi1)
ProteinaseK, Res 16 (chi1)
Pearson r: 0.9321
ProteinaseK, Res 31 (chi1)
ProteinaseK, Res 31 (chi1)
Pearson r: 0.9311
ProteinaseK, Res 34 (chi1)
ProteinaseK, Res 34 (chi1)
Pearson r: 0.9385
ProteinaseK, Res 47 (chi1)
ProteinaseK, Res 47 (chi1)
Pearson r: 0.9444
ProteinaseK, Res 54 (chi1)
ProteinaseK, Res 54 (chi1)
Pearson r: 0.1039
ProteinaseK, Res 63 (chi1)
ProteinaseK, Res 63 (chi1)
Pearson r: 0.8489
ProteinaseK, Res 64 (chi1)
ProteinaseK, Res 64 (chi1)
Pearson r: 0.6609
ProteinaseK, Res 89 (chi1)
ProteinaseK, Res 89 (chi1)
Pearson r: 0.7734
ProteinaseK, Res 99 (chi1)
ProteinaseK, Res 99 (chi1)
Pearson r: 0.8495
277K
100K
Normalized Score
χ1
Normalized Score, Dataset 1
Normalized Score, Dataset 2

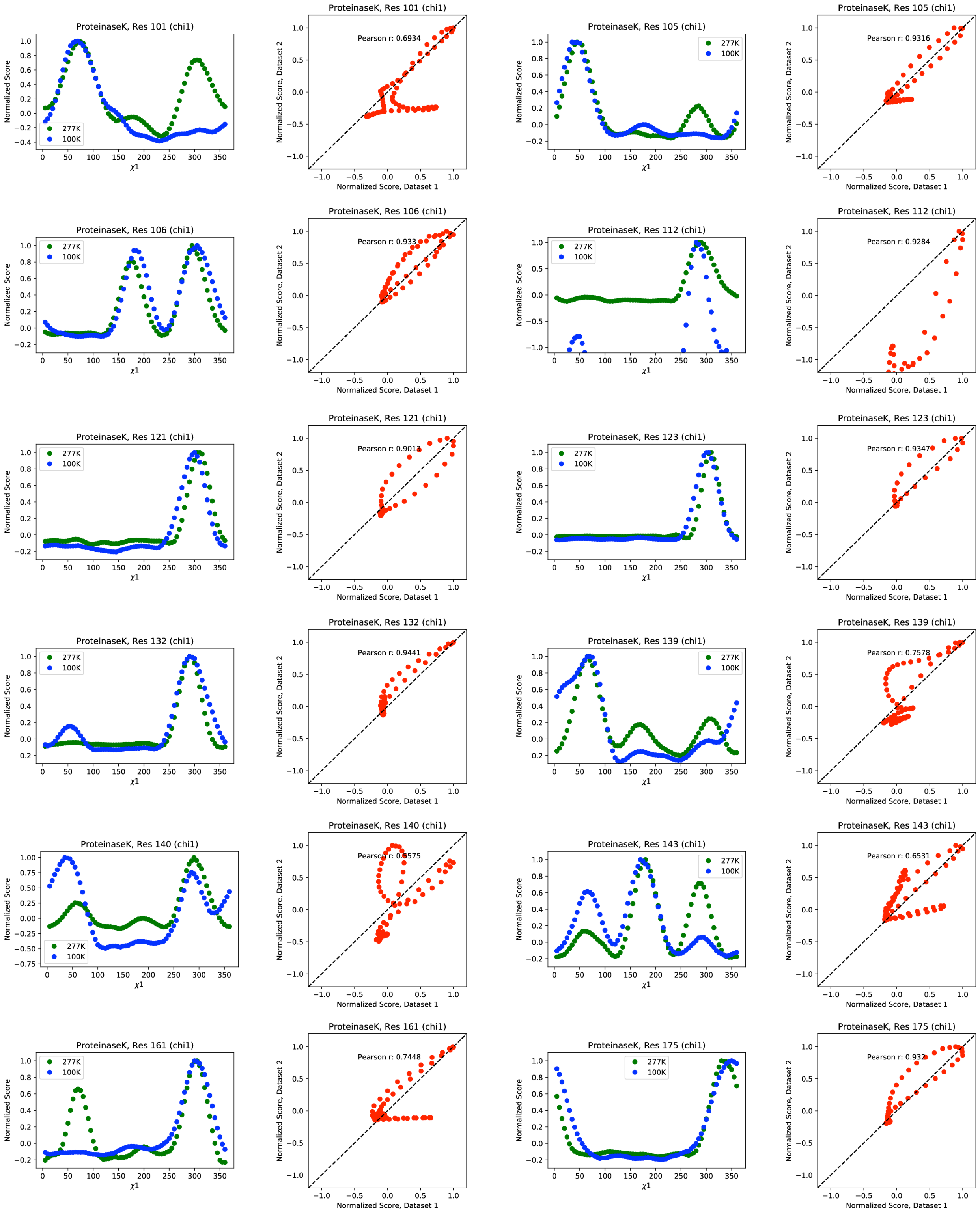
ProteinaseK, Res 101 (chi1)
Pearson r: 0.6934
ProteinaseK, Res 105 (chi1)
Pearson r: 0.9316
ProteinaseK, Res 106 (chi1)
Pearson r: 0.933
ProteinaseK, Res 112 (chi1)
Pearson r: 0.9284
ProteinaseK, Res 121 (chi1)
Pearson r: 0.9013
ProteinaseK, Res 123 (chi1)
Pearson r: 0.9347
ProteinaseK, Res 132 (chi1)
Pearson r: 0.9441
ProteinaseK, Res 139 (chi1)
Pearson r: 0.7578
ProteinaseK, Res 140 (chi1)
ProteinaseK, Res 143 (chi1)
Pearson r: 0.6531
ProteinaseK, Res 161 (chi1)
Pearson r: 0.7448
ProteinaseK, Res 175 (chi1)
277K
100K
Normalized Score
χ1
Normalized Score, Dataset 1
Normalized Score, Dataset 2

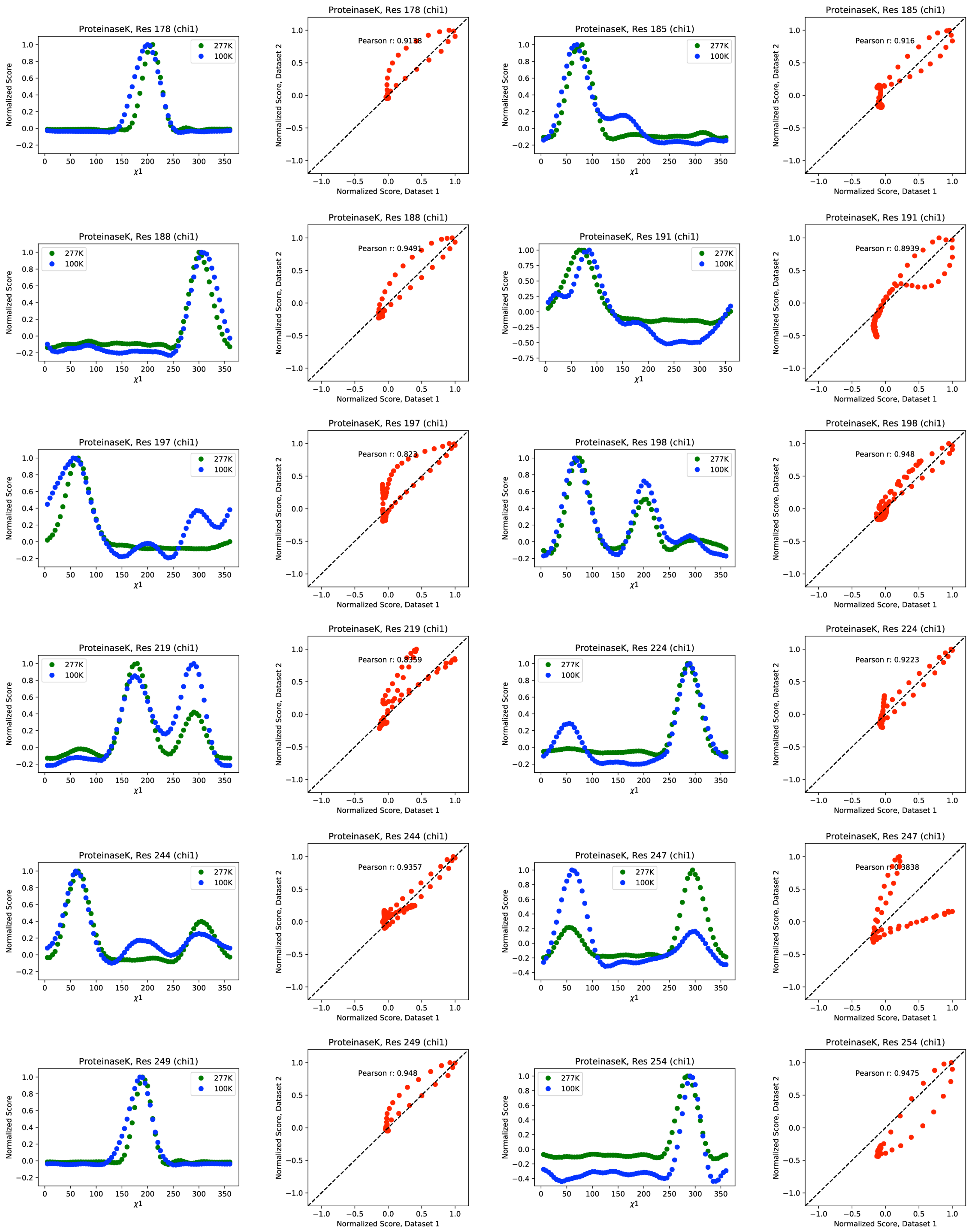
ProteinaseK, Res 178 (chi1)
277K
100K
Normalized Score
χ1
ProteinaseK, Res 178 (chi1)
Pearson r: 0.9138
Normalized Score, Dataset 2
Normalized Score, Dataset 1
ProteinaseK, Res 185 (chi1)
ProteinaseK, Res 185 (chi1)
Pearson r: 0.916
ProteinaseK, Res 188 (chi1)
ProteinaseK, Res 188 (chi1)
Pearson r: 0.9491
ProteinaseK, Res 191 (chi1)
ProteinaseK, Res 191 (chi1)
Pearson r: 0.8939
ProteinaseK, Res 197 (chi1)
ProteinaseK, Res 197 (chi1)
Pearson r: 0.823
ProteinaseK, Res 198 (chi1)
ProteinaseK, Res 198 (chi1)
Pearson r: 0.948
ProteinaseK, Res 219 (chi1)
ProteinaseK, Res 219 (chi1)
Pearson r: 0.8359
ProteinaseK, Res 224 (chi1)
ProteinaseK, Res 224 (chi1)
Pearson r: 0.9223
ProteinaseK, Res 244 (chi1)
ProteinaseK, Res 244 (chi1)
Pearson r: 0.9357
ProteinaseK, Res 247 (chi1)
ProteinaseK, Res 247 (chi1)
Pearson r: 0.3838
ProteinaseK, Res 249 (chi1)
ProteinaseK, Res 249 (chi1)
Pearson r: 0.948
ProteinaseK, Res 254 (chi1)
ProteinaseK, Res 254 (chi1)
Pearson r: 0.9475

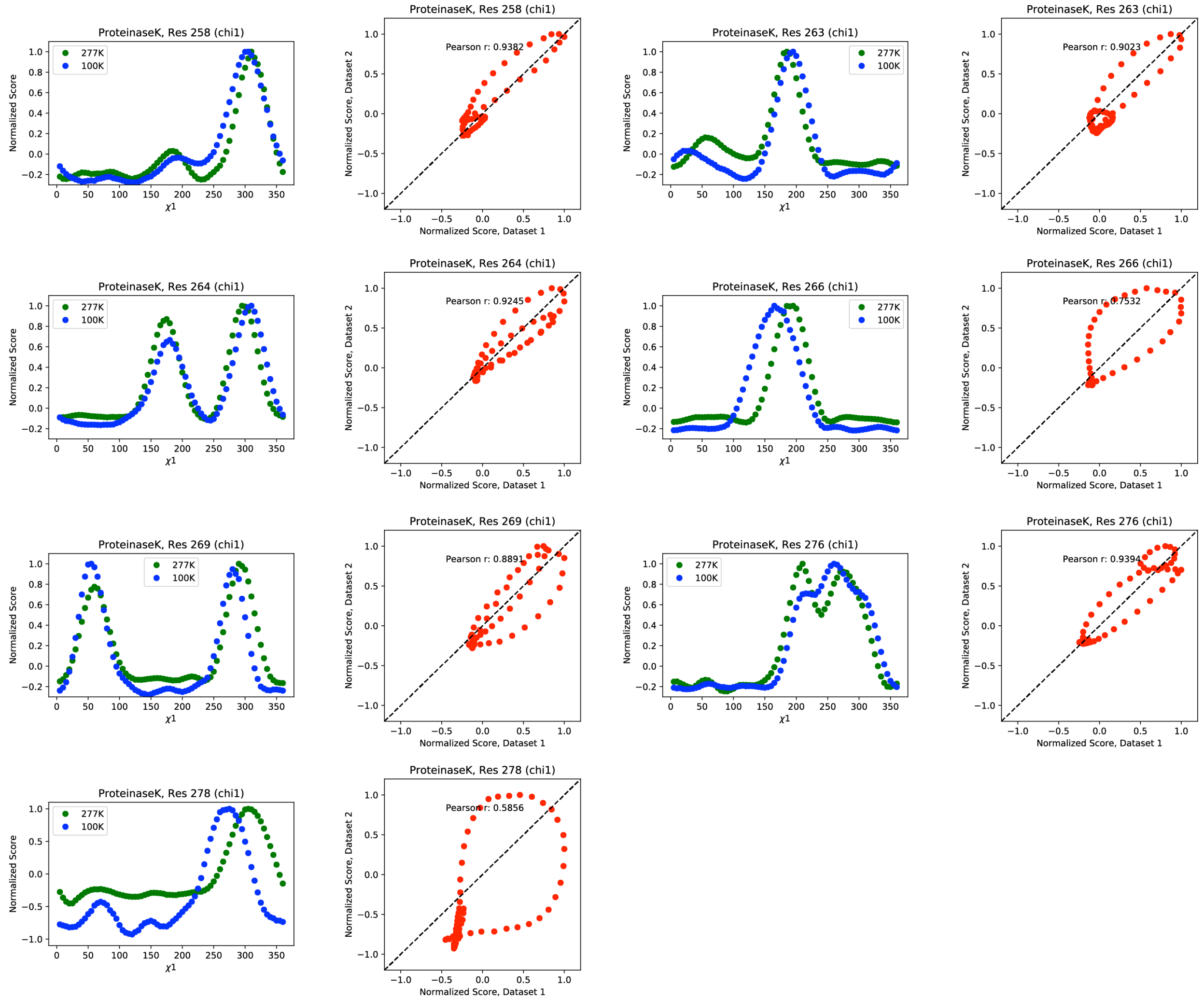

**Figure S19.** Normalized *Ringer* profiles for the proteinase K 277 K least damaged dataset 1 (green) and the 100 K most damaged dataset 7 (blue) with Pearson correlation coefficients ($P_{CC}$) ≤ 0.95. Also shown are the associated correlation plots and respective $P_{CC}$ values.

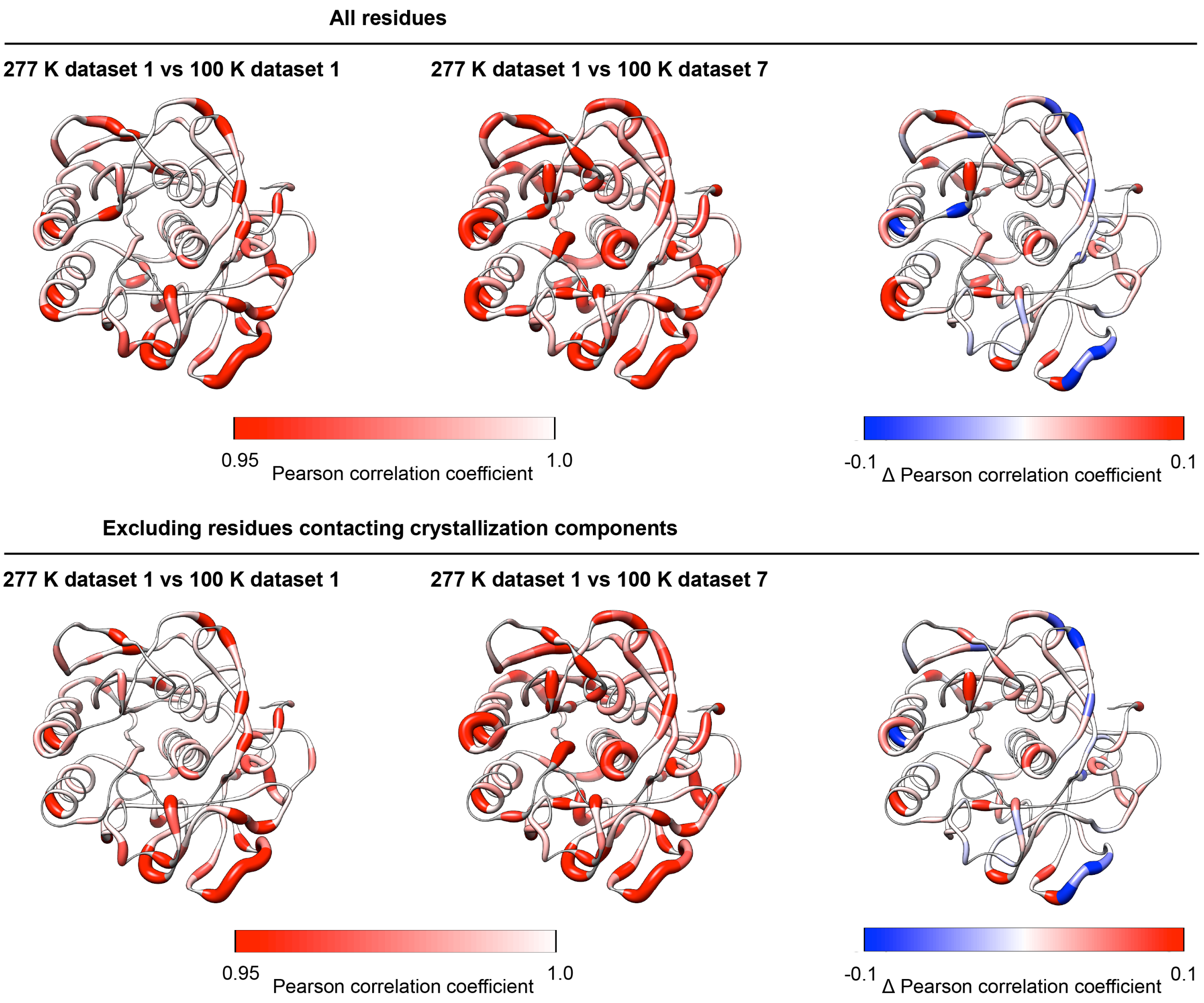

**Figure S20.** Comparison of $P_{CC}$ values (plotted on the proteinase K structure) between the *Ringer* profiles for residues in the 277 K least X-ray damaged dataset 1 and 100 K least X-ray damaged dataset 1 (left), the 277 K least X-ray damaged dataset 1 and 100 K most X-ray damaged dataset 7 (middle), and the difference ($\Delta$)$P_{CC}$ obtained by subtracting top middle from top left $P_{CC}$ values, plotted on the proteinase K structure (right). All residues with $\chi 1$ angles (top) and excluding residues in contact with crystallization components (bottom, see **Table S24**). Residues with no $\chi 1$ angles and excluded residues are colored in grey. The diameter of the worm representation is correlated with the magnitude of the $P_{CC}$.

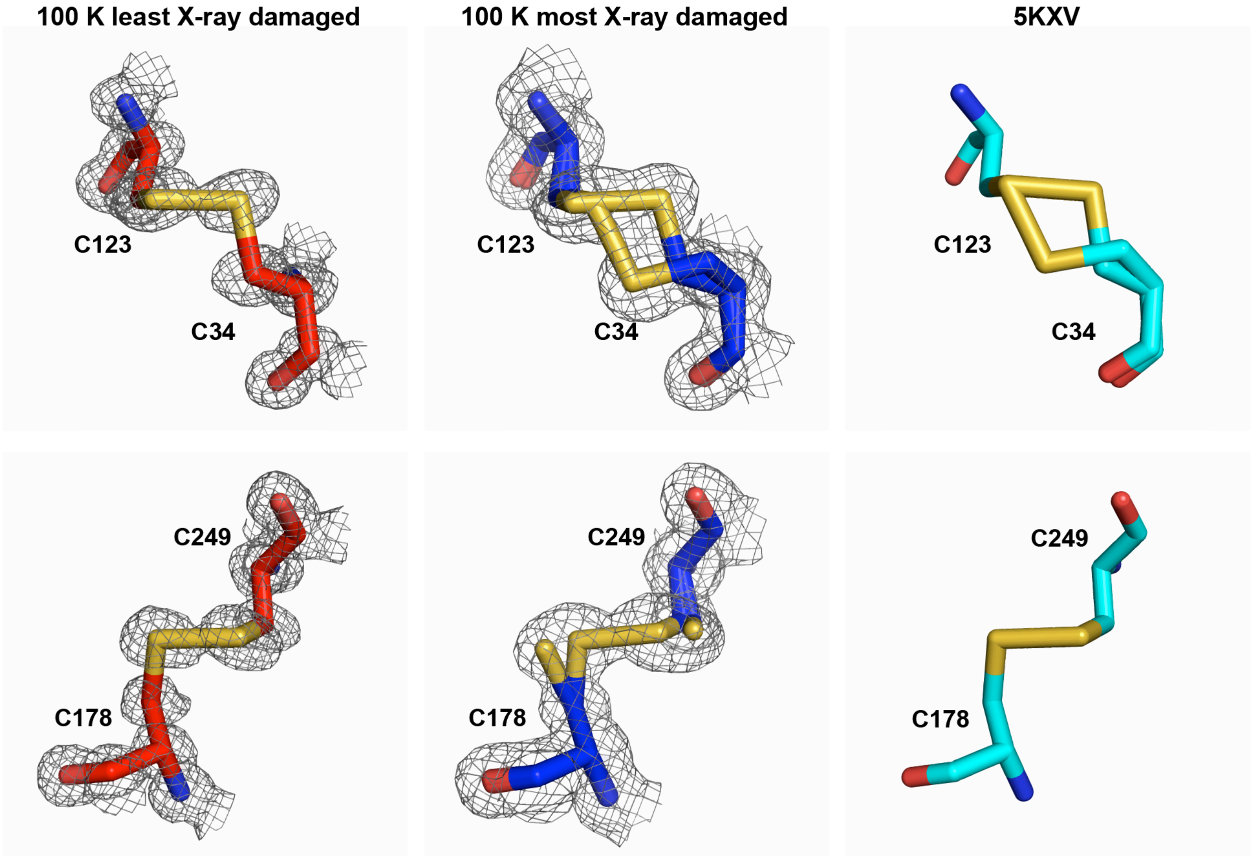

**Figure S21.** A proteinase K cryo structure from the PDB ((60) PDB 5KXV; 0.98 Å resolution) shows evidence for X-ray damage as judged by the deposited coordinates compared to the least and most X-ray damaged proteinase K cryo datasets obtained in this work. The disulfide bond C34-C123 from 5KXV is similar to the disulfide bond in our most X-ray damaged dataset (top). While the coordinates for the other disulfide, C178-C249, indicate lack of damage, Figure S4, B from (60) clearly indicated the presence of X-ray damage and electron density similar to our most damaged dataset.

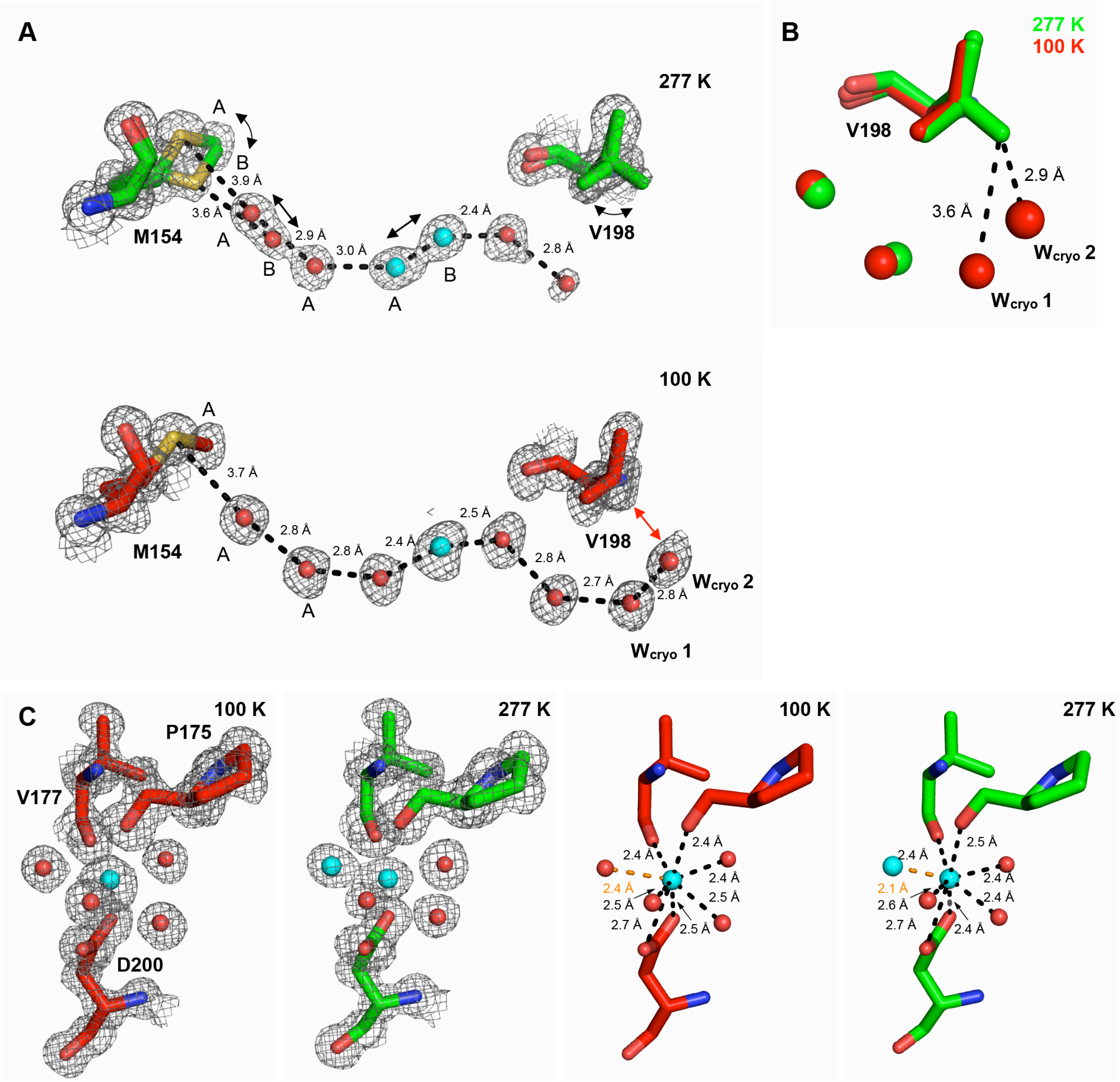

**Figure S22. Extended network of interactions couples motions of distant residues M154 and V198 in proteinase K. (A)** M154 and V198 motions are quenched at cryo (bottom) but not at room temperature (top). **(B)** Water molecules form an extended network upon freezing which quenches V198 motion – waters $W_{cryo}$ 1 and $W_{cryo}$ 2 appear too close to V198 to allow the existence of the alternative rotameric state observed at room temperature. The distance between $W_{cryo}$ 2 and the alternative V198 conformation observed at room temperature would be 2.9 Å, substantially shorter than the sum of van der Waals radii of ~3.6 Å between V198 $CH_3$ (about 2.0 Å) and oxygen (average of about 1.55 Å) (61, 62). These water molecules are not observed in the room temperature dataset. **(C)** The $Ca^{2+}$ ion bound to the proteinase K high affinity site occupies two distinct positions at room temperature but a single position at cryo temperature.

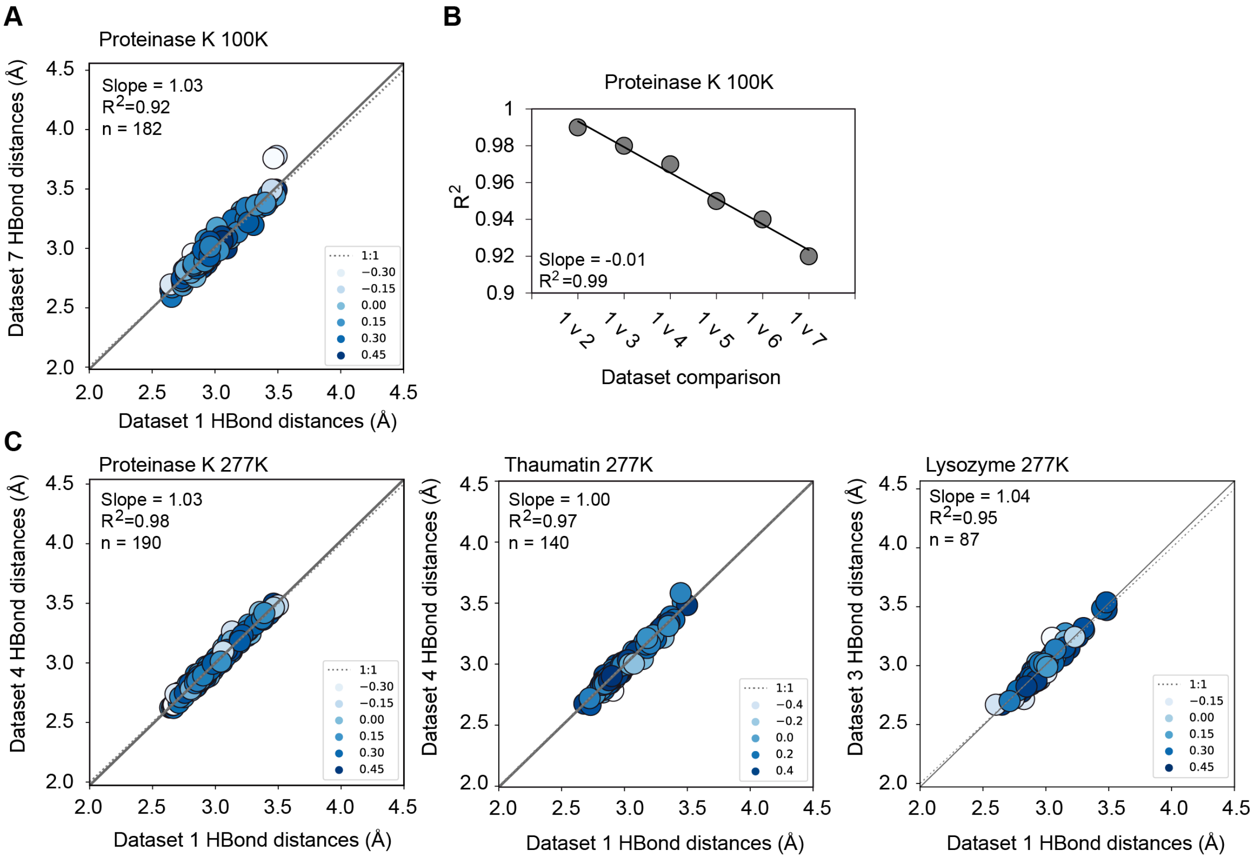

**Figure S23. X-ray damage impacts the determination of hydrogen bond lengths more at cryo temperature than at room temperature**. Correlation plots of hydrogen bond lengths obtained from the least and most damaged datasets at cryo temperature (**A**). Correlation points are colored according to the relative B-factor of the hydrogen bonding groups such that higher values (darker blue) and lower values (white) correspond to atoms with low- and high B-factors relative to the average, respectively (see Materials and Methods). (**B**) Correlation coefficients ($R^2$) obtained from correlation plots of hydrogen bond lengths from the least damaged ("1") and increasingly damaged proteinase K 100 K structures ("2–7") (see **Figure S24** for individual correlation plots). Differences between proteinase K structures are unlikely to result from differences in refinement strategy as all structures were refined using the same refinement parameters and increasingly X-ray damaged models were refined in a consistent manner (see Materials and Methods). (**C**) Correlation plots of hydrogen bond lengths obtained from the least and most damaged datasets at room temperature. Colors used as in (A). The analysis excluded all residues with more than one conformation present in the model (see Materials and Methods). Similar results were obtained with all residues included (**Figures S25**). See *Supplementary file 2* for hydrogen bond length numerical values.

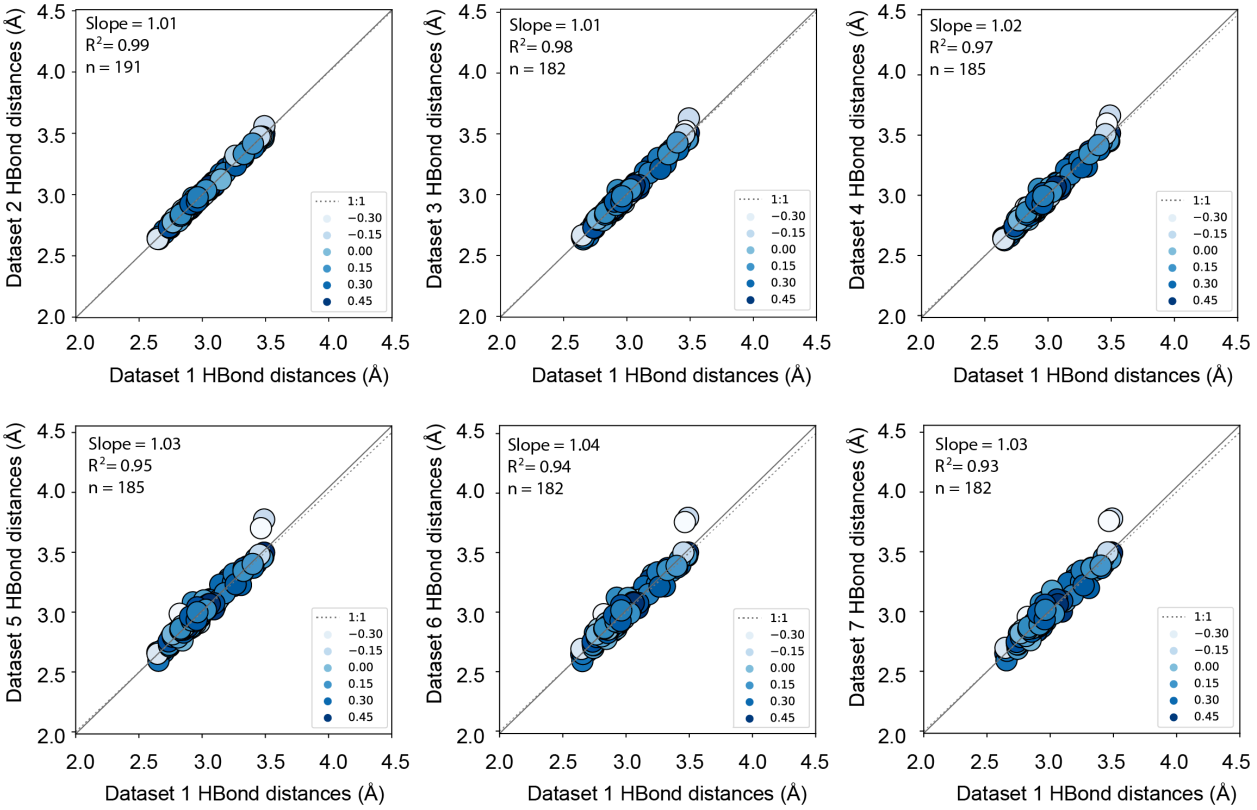

**Figure S24. Gradual deterioration of measured hydrogen bond lengths in proteinase K with increasing X-ray damage at 100 K**. Correlation plots of hydrogen bond lengths obtained from the least (dataset 1) and increasingly X-ray damaged (dataset 2-7) proteinase K datasets at 100 K. Correlation points are colored according to the relative B-factor of the hydrogen bonding groups such that higher values (darker blue) and lower values (white) correspond to atoms with low- and high B-factors relative to the average, respectively (see Materials and Methods). The individual correlation coefficients ($R^2$) as a function of dataset number are plotted in Figure S23B. Differences between proteinase K structures are unlikely to be caused by differences in refinement strategy as all structures have been refined using the same refinement parameters and increasingly damaged models have been refined in a highly consistent manner (see Materials and Methods). The analysis excluded all residues with more than one conformation present in the model (see Materials and Methods). Similar results were obtained with all residues included (**Figure S25**). See *Supplementary file 2* for hydrogen bond length numerical values.

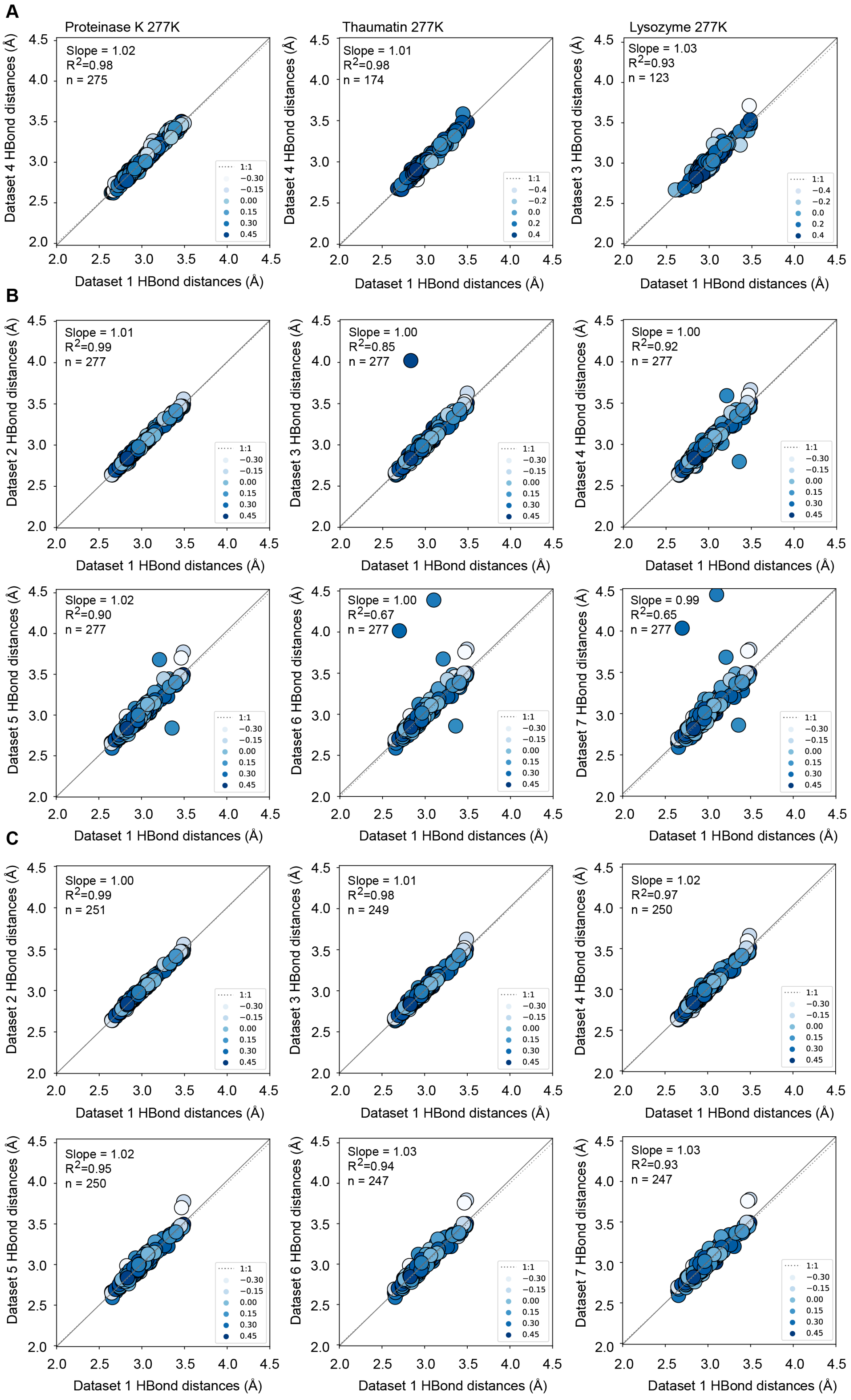

A
Proteinase K 277K
Slope = 1.02
$R^2$=0.98
n = 275
Dataset 4 HBond distances (Å)
Dataset 1 HBond distances (Å)
1:1
−0.30
−0.15
0.00
0.15
0.30
0.45
Thaumatin 277K
Slope = 1.01
$R^2$=0.98
n = 174
Dataset 4 HBond distances (Å)
Dataset 1 HBond distances (Å)
1:1
−0.4
−0.2
0.0
0.2
0.4
Lysozyme 277K
Slope = 1.03
$R^2$=0.93
n = 123
Dataset 3 HBond distances (Å)
Dataset 1 HBond distances (Å)
1:1
−0.4
−0.2
0.0
0.2
0.4
B
Slope = 1.01
$R^2$=0.99
n = 277
Dataset 2 HBond distances (Å)
Dataset 1 HBond distances (Å)
Slope = 1.00
$R^2$=0.85
n = 277
Dataset 3 HBond distances (Å)
Dataset 1 HBond distances (Å)
Slope = 1.00
$R^2$=0.92
n = 277
Dataset 4 HBond distances (Å)
Dataset 1 HBond distances (Å)
Slope = 1.02
$R^2$=0.90
n = 277
Dataset 5 HBond distances (Å)
Dataset 1 HBond distances (Å)
Slope = 1.00
$R^2$=0.67
n = 277
Dataset 6 HBond distances (Å)
Dataset 1 HBond distances (Å)
Slope = 0.99
$R^2$=0.65
n = 277
Dataset 7 HBond distances (Å)
Dataset 1 HBond distances (Å)
1:1
−0.30
−0.15
0.00
0.15
0.30
0.45
C
Slope = 1.00
$R^2$=0.99
n = 251
Dataset 2 HBond distances (Å)
Dataset 1 HBond distances (Å)
Slope = 1.01
$R^2$=0.98
n = 249
Dataset 3 HBond distances (Å)
Dataset 1 HBond distances (Å)
Slope = 1.02
$R^2$=0.97
n = 250
Dataset 4 HBond distances (Å)
Dataset 1 HBond distances (Å)
Slope = 1.02
$R^2$=0.95
n = 250
Dataset 5 HBond distances (Å)
Dataset 1 HBond distances (Å)
Slope = 1.03
$R^2$=0.94
n = 247
Dataset 6 HBond distances (Å)
Dataset 1 HBond distances (Å)
Slope = 1.03
$R^2$=0.93
n = 247
Dataset 7 HBond distances (Å)
Dataset 1 HBond distances (Å)
1:1
−0.30
−0.15
0.00
0.15
0.30
0.45

**Figure S25. X-ray damage impacts the determination of hydrogen bond lengths in proteinase K more at cryo temperature than at room temperature**. (**A**) Correlation plots of hydrogen bond lengths obtained from the least and most damaged datasets at room temperature. Correlation points are colored according to the relative B-factor of the hydrogen bonding groups such that higher values (darker blue) and lower values (white) correspond to atoms with low- and high B-factors relative to the average, respectively (see Materials and Methods). (**B**) Correlation plots of hydrogen bond lengths obtained from the least (dataset 1) and increasingly X-ray damaged (dataset 2-7) proteinase K datasets at 100 K. Colors used as in (A). (A) and (B) analyses included all residues and average values are plotted for residues with more than one conformation present in the model (see Materials and Methods). (**C**) Correlation plots of hydrogen bond lengths obtained from the least (dataset 1) and increasingly X-ray damaged (dataset 2-7) proteinase K datasets at 100 K. The analysis excluded all side chains with more than one conformation present in the model but included average values for backbone hydrogen bonding groups with more than one conformation (see Materials and Methods). Colors used as in (A). See *Supplementary file 2* for hydrogen bond length numerical values.

| Thaumatin 277 K diffraction data collection statistics | | | | |
|---|---|---|---|---|
| Batch | 1 | 2 | 3 | 4 |
| Wavelength (Å) | 0.88557 | | | |
| Resolution range (Å) | 38-26-1.22 (1.24-1.22) | 38.28-1.29 (1.31-1.29) | 38.29-1.38 (1.40-1.38) | 38.29-1.48 (1.51-1.48) |
| Average diffraction weighted dose (MGy) | 0.016 | 0.114 | 0.212 | 0.310 |
| Space group | $P4_12_12$ | | | |
| Unit cell | 58.81 58.81 151.17 90.00 90.00 90.00 | 58.84 58.84 151.21 90.00 90.00 90.00 | 58.85 58.85 151.26 90.00 90.00 90.00 | 58.83 58.83 151.27 90.00 90.00 90.00 |
| Unit cell volume ($Å^3$) | 522839 | 523511 | 523862.2 | 523540.8 |
| Total reflections | 686302 (30123) | 587692 (29088) | 480296 (22950) | 389623 (17213) |
| Multiplicity | 8.6 (7.9) | 8.7 (8.6) | 8.6 (8.6) | 8.6 (8.0) |
| Mosaicity (°) | 0.12 | 0.13 | 0.14 | 0.14 |
| Completeness (%) | 99.9 (98.6) | 99.9 (98.1) | 99.9 (98.0) | 99.9 (97.9) |
| Mean I/sigma(I) | 8.5 (0.5) | 8.9 (0.5) | 9.2 (0.5) | 9.1 (0.6) |
| Wilson B-factor | 20.9 | 23.7 | 26.6 | 28.7 |
| R-merge | 0.110 (4.479) | 0.118 (5.270) | 0.139 (7.530) | 0.186 (9.560) |
| R-pim | 0.040 (1.678) | 0.042 (1.868) | 0.050 (2.668) | 0.066 (3.509) |
| $CC_{1/2}$ | 0.999 (0.304) | 0.999 (0.302) | 0.999 (0.320) | 0.999 (0.335) |
| Isa | 17.3 | 18.0 | 17.7 | 18.0 |

| Thaumatin crystal structure refinement statistics | | | |
|---|---|---|---|
| Dataset | 1 | | 4 |
| PDB code | 7LFG | | 7LJV |
| Resolution range (Å) | 37.79 - 1.22 (1.26 - 1.22) | | 37.82 - 1.48 (1.53 - 1.48) |
| Reflections used in refinement | 78744 (6867) | | 43991 (3270) |
| Rwork | 0.135 (0.320) | | 0.138 (0.303) |
| Rfree | 0.150 (0.337) | | 0.169 (0.329) |
| No. on non-hydrogen atoms | 1957 | | 1911 |
| Protein | 1764 | | 1744 |
| Ligand/ion | 10 | | 10 |
| Water | 165 | | 143 |
| RMS (bonds) | 0.007 | | 0.007 |
| RMS (angles) | 0.94 | | 0.92 |
| Average B-factor | 21.77 | | 26.59 |
| Protein | 20.2 | | 25.34 |
| Ligand/ion | 17.85 | | 23.44 |
| Water | 37.17 | | 40.69 |
| Ramachandran (%) | | | |
| Favored | 98.1 | | 98.1 |
| Allowed | 1.9 | | 1.9 |
| Outliers | 0 | | 0 |

**Table S1**. **Thaumatin room temperature diffraction and traditional single conformation model refinement statistics.** Diffraction statistics for increasingly damaged thaumatin datasets obtained at 277 K from a single crystal. Diffraction statistics are reported for datasets of 120° total rotations obtained from the same crystal orientation.

Values in parenthesis are for the highest resolution shells. All statistics were obtained from Aimless (41), with the exception of $CC_{1/2}$, which was obtained from XSCALE (39). Average diffraction weighted doses (DWD) were estimated using the program RADDOSE 3D (35, 36). Refinement statistics were obtained from phenix (*phenix.table_one*) using the final refined models and reflections file.

| Proteinase K 277 K diffraction data collection statistics | | | | |
|---|---|---|---|---|
| Batch | 1 | 2 | 3 | 4 |
| Wavelength (Å) | 0.88557 | | | |
| Resolution range (Å) | 34.91-1.02<br>(1.04-1.02) | 34.84-1.10<br>(1.12-1.10) | 34.81-1.30<br>(1.32-1.30) | 34.85-1.43<br>(1.45-1.43) |
| Average diffraction weighted dose (MGy) | 0.006 | 0.023 | 0.041 | 0.058 |
| Space group | $P4_32_12$ | | | |
| Unit cell | 67.82 67.82<br>101.86 90.00<br>90.00 90.00 | 67.71 67.71<br>101.60 90.00<br>90.00 90.00 | 67.65 67.65<br>101.52 90.00<br>90.00 90.00 | 67.71 67.71<br>101.65 90.00<br>90.00 90.00 |
| Unit cell volume ($Å^3$) | 468510.4 | 465799.8 | 464608.6 | 466029.1 |
| Total reflections | 822825 (28269) | 695054 (23190) | 480869 (17003) | 379902 (16456) |
| Multiplicity | 6.8 (4.9) | 7.2 (5.0) | 8.2 (6.0) | 8.6 (7.7) |
| Mosaicity (°) | 0.08 | 0.12 | 0.22 | 0.30 |
| Completeness (%) | 99.9<br>(98.3) | 99.9 (98.7) | 99.9 (98.9) | 99.9 (98.8) |
| Mean I/sigma(I) | 8.5 (0.9) | 8.8 (0.8) | 12.1 (0.7) | 11.4 (0.6) |
| Wilson B-factor | 12.2 | 14.7 | 20.8 | 24.6 |
| R-merge | 0.098 (1.720) | 0.102 (1.946) | 0.107 (2.646) | 0.130 (3.551) |
| R-pim | 0.038 (0.862) | 0.039 (0.950) | 0.039 (1.138) | 0.047 (1.335) |
| $CC_{1/2}$ | 0.999 (0.359) | 0.999 (0.319) | 0.999 (0.306) | 0.999 (0.320) |
| Isa | 19.3 | 17.7 | 35.4 | 40.6 |

| Proteinase K crystal structure refinement statistics | | | |
|---|---|---|---|
| Dataset | 1 | | 4 |
| PDB code | 7LN7 | | 7LPT |
| Resolution range (Å) | 33.91 - 1.02<br>(1.06 - 1.02) | | 34.85 - 1.43<br>(1.48 - 1.43) |
| Reflections used in refinement | 120585 (11691) | | 44021 (4101) |
| Rwork | 0.123 (0.283) | | 0.125 (0.269) |
| Rfree | 0.138 (0.284) | | 0.162 (0.323) |
| No. on non-hydrogen atoms | 2817 | | 2735 |
| Protein | 2477 | | 2435 |
| Ligand/ion | 17 | | 17 |
| Water | 287 | | 283 |
| RMS (bonds) | 0.006 | | 0.007 |
| RMS (angles) | 0.92 | | 0.89 |
| Average B-factor | 11.81 | | 19.87 |
| Protein | 9.73 | | 17.87 |
| Ligand/ion | 37.12 | | 52.69 |
| Water | 26.46 | | 35.10 |
| Ramachandran (%) | | | |
| Favored | 97.1 | | 96.4 |
| Allowed | 2.9 | | 3.6 |
| Outliers | 0 | | 0 |

**Table S2. Proteinase K room temperature diffraction and traditional single conformation model refinement statistics.** Diffraction statistics for increasingly damaged proteinase K datasets obtained at 277 K from a single crystal. Diffraction statistics are reported for datasets of 120° total rotations obtained from the same crystal orientation. Values in parenthesis are for the highest resolution shells. All statistics were obtained from Aimless (41), with the exception of $CC_{1/2}$, which was obtained from XSCALE (39). Average diffraction weighted doses (DWD) were estimated using the program RADDOSE 3D (35, 36). Refinement statistics were obtained from phenix (*phenix.table_one*) using the final refined models and reflections file.

| Lysozyme 277 K diffraction data collection statistics | | | |
|---|---|---|---|
| Batch | 1 | 2 | 3 |
| Wavelength (Å) | 0.88557 | | |
| Resolution range (Å) | 38.63-1.13 (1.15-1.13) | 38.61-1.30 (1.32-1.30) | 38.70-1.52 (1.55-1.52) |
| Average diffraction weighted dose (MGy) | 0.017 | 0.069 | 0.121 |
| Space group | $P4_32_12$ | | |
| Unit cell | 77.26 77.26 37.31 90.00 90.00 90.00 | 77.22 77.22 37.19 90.00 90.00 90.00 | 77.40 77.40 37.21 90.00 90.00 90.00 |
| Unit cell volume ($Å^3$) | 222707.4 | 221761.3 | 222916.2 |
| Total reflections | 365238 (17155) | 241397 (11737) | 152376 (6153) |
| Multiplicity | 8.5 (8.4) | 8.6 (8.7) | 8.5 (7.2) |
| Mosaicity (°) | 0.14 | 0.25 | 0.37 |
| Completeness (%) | 99.9 (99.1) | 99.9 (98.2) | 99.9 (98.1) |
| Mean I/sigma(I) | 12.4 (0.9) | 14.3 (0.8) | 12.7 (0.6) |
| Wilson B-factor | 18.9 | 24.5 | 31.3 |
| R-merge | 0.072 (2.720) | 0.077 (3.766) | 0.102 (4.132) |
| R-pim | 0.026 (0.989) | 0.028 (1.356) | 0.037 (1.620) |
| $CC_{1/2}$ | 0.999 (0.326) | 0.999 (0.317) | 0.999 (0.335) |
| Isa | 20.1 | 25.7 | 28.9 |
| **Lysozyme crystal structure refinement statistics** | | | |
| Dataset | 1 | | 3 |
| PDB code | 7LLP | | 7LN8 |
| Resolution range (Å) | 33.6 - 1.13 (1.17 - 1.13) | | 33.53 - 1.52 (1.58 - 1.52) |
| Reflections used in refinement | 42596 (4064) | | 17663 (1565) |
| Rwork | 0.132 (0.292) | | 0.139 (0.261) |
| Rfree | 0.155 (0.313) | | 0.200 (0.375) |
| No. on non-hydrogen atoms | 1383 | | 1367 |
| Protein | 1247 | | 1247 |
| Ligand/ion | 3 | | 3 |
| Water | 116 | | 102 |
| RMS (bonds) | 0.008 | | 0.008 |
| RMS (angles) | 0.99 | | 0.91 |
| Average B-factor | 20.3 | | 26.99 |
| Protein | 19.1 | | 26.02 |
| Ligand/ion | 28.83 | | 39.18 |
| Water | 31.44 | | 37.04 |
| Ramachandran (%) | | | |
| Favored | 99.2 | | 100 |
| Allowed | 0.8 | | 0 |
| Outliers | 0 | | 0 |

**Table S3**. **Lysozyme room temperature diffraction and traditional single conformation model refinement statistics.** Diffraction statistics for increasingly damaged lysozyme datasets obtained at 277 K from a single crystal. Diffraction statistics are reported for datasets of 120° total rotations obtained from the same crystal orientation. Values in parenthesis are for the highest resolution shells. All statistics were obtained from Aimless (41), with the exception of $CC_{1/2}$, which was obtained from XSCALE (39). Average diffraction weighted doses (DWD) were estimated using the program RADDOSE 3D (35, 36). Refinement statistics were obtained from phenix (*phenix.table_one*) using the final refined models and reflections file.

| Lysozyme 277 K diffraction data collection statistics | | |
|---|---|---|
| Batch | Crystal 1 dataset 1 | Crystal 2 dataset 1 |
| Wavelength (Å) | 0.88557 | 0.88557 |
| Resolution range (Å) | 38.63-1.13 (1.15-1.13) | 38.70-1.10 (1.12-1.10 |
| Dose (MGy) [a] | 0.058 | 0.029 |
| Space group | $P4_32_12$ | $P4_32_12$ |
| Unit cell | 77.26 77.26 37.31 90.00 90.00 90.00 | 77.41 77.41 37.42 90.00 90.00 90.00 |
| Unit cell volume (Å$^3$) | 222707.4 | 224232.2 |
| Total reflections | 365238 (17155) | 399774 (18647) |
| Multiplicity | 8.5 (8.4) | 8.6 (8.4) |
| Mosaicity (°) | 0.14 | 0.08 |
| Completeness (%) | 99.9 (99.1) | 99.3 (98.3) |
| Mean I/sigma(I) | 12.4 (0.9) | 9.8 (0.7) |
| Wilson B-factor | 18.9 | 17.6 |
| R-merge | 0.072 (2.720) | 0.083 (2.940) |
| R-pim | 0.026 (0.989) | 0.029 (1.066) |
| $CC_{1/2}$ | 0.999 (0.326) | 0.999 (0.302) |
| Isa | 20.1 | 19.5 |
| **Lysozyme multi-conformer model refinement statistics** | | |
| Dataset | 1 | 1 |
| PDB code | 7LN9 | 7LPM |
| Resolution range (Å) | 33.6 - 1.13 (1.171 - 1.13) | 34.62 - 1.1 (1.14 - 1.1) |
| Reflections used in refinement | 42596 (4064) | 46101 (4434) |
| Rwork | 0.136 (0.292) | 0.136 (0.292) |
| Rfree | 0.156 (0.308) | 0.158 (0.301) |
| No. on non-hydrogen atoms | 3065 | 3125 |
| Protein | 2925 | 2984 |
| Ligand/ion | 4 | 4 |
| Water | 116 | 112 |
| RMS (bonds) | 0.006 | 0.006 |
| RMS (angles) | 0.84 | 0.81 |
| Average B-factor | 15.8 | 15.14 |
| Protein | 15.2 | 14.55 |
| Ligand/ion | 22.25 | 22.14 |
| Water | 28.48 | 27.8 |
| Ramachandran (%) | | |
| Favored | 96.1 | 96.1 |
| Allowed | 3.9 | 3.9 |
| Outliers | 0 | 0 |

**Table S4**. **Room temperature (277K) diffraction statistics and multi-conformer refinement statistics for two lysozyme crystals.** Diffraction statistics are reported for datasets of 120° total rotations. Values in parenthesis are for the highest resolution shells. All statistics were obtained from Aimless (41), with the exception of $CC_{1/2}$, which was obtained from XSCALE (39). Average diffraction weighted doses (DWD) were estimated using the program RADDOSE 3D (35, 36). Multi-conformer models were obtained as described in Materials and Methods. Refinement statistics were obtained from phenix (*phenix.table_one*) using the final refined models and reflections files. Diffraction statistics for crystal 1 are from **Table S3** dataset 1.

| Thaumatin | | | Proteinase K | | | Lysozyme | | |
|---|---|---|---|---|---|---|---|---|
| Residue | MSE | $P_{CC}$ | Residue | MSE | $P_{CC}$ | Residue | MSE | $P_{CC}$ |
| 2 | 0.0036 | 0.9789 | 3 | 0.0007 | 0.9980 | 1 | 0.0056 | 0.9785 |
| 3 | 0.0009 | 0.9976 | 4 | 0.0026 | 0.9878 | 2 | 0.0111 | 0.9664 |
| 4 | 0.0001 | 0.9997 | 5 | 0.0290 | 0.9279 | 3 | 0.0005 | 0.9987 |
| 5 | 0.0013 | 0.9972 | 7 | 0.0015 | 0.9938 | 5 | 0.0004 | 0.9992 |
| 6 | 0.0016 | 0.9968 | 8 | 0.0028 | 0.9931 | 6 | 0.0010 | 0.9956 |
| 7 | 0.0004 | 0.9982 | 10 | 0.0013 | 0.9965 | 7 | 0.0020 | 0.9952 |
| 8 | 0.0008 | 0.9967 | 12 | 0.0013 | 0.9963 | 8 | 0.0043 | 0.9844 |
| 9 | 0.0006 | 0.9969 | 13 | 0.0035 | 0.9910 | 12 | 0.0008 | 0.9968 |
| 10 | 0.0016 | 0.9950 | 14 | 0.0019 | 0.9922 | 13 | 0.0032 | 0.9899 |
| 11 | 0.0004 | 0.9993 | 15 | 0.0036 | 0.9824 | 14 | 0.0049 | 0.9870 |
| 12 | 0.0012 | 0.9977 | 16 | 0.0016 | 0.9947 | 15 | 0.0008 | 0.9968 |
| 13 | 0.0039 | 0.9877 | 17 | 0.0008 | 0.9970 | 17 | 0.0006 | 0.9991 |
| 14 | 0.0008 | 0.9967 | 18 | 0.0027 | 0.9879 | 18 | 0.0151 | 0.9473 |
| 18 | 0.0005 | 0.9983 | 20 | 0.0036 | 0.9883 | 19 | 0.0025 | 0.9921 |
| 19 | 0.0004 | 0.9989 | 21 | 0.0012 | 0.9971 | 20 | 0.0015 | 0.9952 |
| 21 | 0.0003 | 0.9989 | 22 | 0.0022 | 0.9903 | 21 | 0.0022 | 0.9935 |
| 24 | 0.0038 | 0.9838 | 23 | 0.0012 | 0.9959 | 23 | 0.0015 | 0.9958 |
| 25 | 0.0017 | 0.9971 | 24 | 0.0026 | 0.9921 | 24 | 0.0014 | 0.9955 |
| 29 | 0.0062 | 0.9722 | 25 | 0.0033 | 0.9927 | 25 | 0.0002 | 0.9994 |
| 30 | 0.0008 | 0.9969 | 26 | 0.0029 | 0.9937 | 27 | 0.0002 | 0.9996 |
| 31 | 0.0003 | 0.9993 | 27 | 0.0030 | 0.9899 | 28 | 0.0012 | 0.9972 |
| 32 | 0.0010 | 0.9978 | 28 | 0.0019 | 0.9930 | 29 | 0.0025 | 0.9931 |
| 33 | 0.0022 | 0.9899 | 31 | 0.0031 | 0.9907 | 30 | 0.0004 | 0.9984 |
| 35 | 0.0002 | 0.9993 | 33 | 0.0025 | 0.9890 | 33 | 0.0003 | 0.9988 |
| 36 | 0.0005 | 0.9978 | 34 | 0.0005 | 0.9973 | 34 | 0.0020 | 0.9943 |
| 37 | 0.0004 | 0.9986 | 35 | 0.0029 | 0.9949 | 35 | 0.0003 | 0.9990 |
| 38 | 0.0043 | 0.9871 | 36 | 0.0041 | 0.9898 | 36 | 0.0011 | 0.9952 |
| 39 | 0.0012 | 0.9959 | 37 | 0.0041 | 0.9901 | 37 | 0.0031 | 0.9942 |
| 40 | 0.0084 | 0.9746 | 38 | 0.0017 | 0.9953 | 38 | 0.0012 | 0.9966 |
| 41 | 0.0028 | 0.9916 | 39 | 0.0021 | 0.9941 | 39 | 0.0005 | 0.9980 |
| 42 | 0.1057 | 0.8173 | 40 | 0.0014 | 0.9957 | 40 | 0.0007 | 0.9974 |
| 43 | 0.0009 | 0.9981 | 42 | 0.0016 | 0.9945 | 41 | 0.0005 | 0.9980 |
| 45 | 0.0023 | 0.9967 | 43 | 0.0015 | 0.9948 | 43 | 0.0027 | 0.9902 |
| 46 | 0.0049 | 0.9900 | 45 | 0.0014 | 0.9964 | 44 | 0.0007 | 0.9982 |
| 49 | 0.0008 | 0.9980 | 46 | 0.0016 | 0.9942 | 45 | 0.0040 | 0.9860 |
| 50 | 0.0007 | 0.9981 | 47 | 0.0073 | 0.9826 | 46 | 0.0025 | 0.9941 |
| 51 | 0.0021 | 0.9952 | 48 | 0.0071 | 0.9768 | 47 | 0.0119 | 0.9602 |
| 53 | 0.0003 | 0.9988 | 49 | 0.0029 | 0.9913 | 48 | 0.0017 | 0.9966 |
| 54 | 0.0012 | 0.9965 | 50 | 0.0022 | 0.9929 | 50 | 0.0007 | 0.9969 |
| 55 | 0.0005 | 0.9987 | 52 | 0.0005 | 0.9980 | 51 | 0.0012 | 0.9953 |
| 56 | 0.0007 | 0.9969 | 54 | 0.0187 | 0.9574 | 52 | 0.0014 | 0.9962 |
| 57 | 0.0008 | 0.9977 | 55 | 0.0048 | 0.9805 | 53 | 0.0005 | 0.9982 |
| 58 | 0.0006 | 0.9979 | 56 | 0.0018 | 0.9934 | 55 | 0.0042 | 0.9894 |
| 59 | 0.0025 | 0.9907 | 57 | 0.0018 | 0.9946 | 56 | 0.0014 | 0.9971 |
| 60 | 0.0683 | 0.8947 | 58 | 0.0020 | 0.9960 | 57 | 0.0004 | 0.9987 |
| 61 | 0.0039 | 0.9920 | 59 | 0.0052 | 0.9804 | 58 | 0.0007 | 0.9981 |
| 63 | 0.0106 | 0.9700 | 60 | 0.0008 | 0.9979 | 59 | 0.0023 | 0.9912 |
| 65 | 0.0010 | 0.9972 | 61 | 0.0044 | 0.9890 | 60 | 0.0014 | 0.9953 |
| 66 | 0.0010 | 0.9958 | 62 | 0.0025 | 0.9899 | 61 | 0.0098 | 0.9752 |
| 67 | 0.0003 | 0.9991 | 63 | 0.0048 | 0.9850 | 62 | 0.0088 | 0.9851 |

| | | | | | | | | |
|---|---|---|---|---|---|---|---|---|
| 68 | 0.0006 | 0.9989 | 64 | 0.1133 | 0.9295 | 63 | 0.0008 | 0.9971 |
| 70 | 0.0009 | 0.9975 | 65 | 0.0015 | 0.9943 | 64 | 0.0009 | 0.9963 |
| 71 | 0.0014 | 0.9920 | 67 | 0.0040 | 0.9920 | 65 | 0.0024 | 0.9923 |
| 74 | 0.0038 | 0.9908 | 69 | 0.0018 | 0.9939 | 66 | 0.0013 | 0.9945 |
| 75 | 0.0022 | 0.9922 | 71 | 0.0027 | 0.9915 | 68 | 0.0016 | 0.9958 |
| 76 | 0.0032 | 0.9903 | 72 | 0.0015 | 0.9957 | 69 | 0.0015 | 0.9949 |
| 77 | 0.0004 | 0.9980 | 73 | 0.0004 | 0.9983 | 70 | 0.1498 | 0.9790 |
| 78 | 0.0690 | 0.9190 | 76 | 0.0027 | 0.9910 | 72 | 0.0376 | 0.9677 |
| 79 | 0.0071 | 0.9815 | 77 | 0.0021 | 0.9948 | 73 | 0.0035 | 0.9931 |
| 80 | 0.0026 | 0.9911 | 79 | 0.0046 | 0.9788 | 74 | 0.0018 | 0.9954 |
| 82 | 0.0033 | 0.9862 | 80 | 0.0050 | 0.9813 | 75 | 0.0096 | 0.9741 |
| 83 | 0.0020 | 0.9936 | 81 | 0.0024 | 0.9929 | 76 | 0.0006 | 0.9969 |
| 84 | 0.0006 | 0.9974 | 82 | 0.0034 | 0.9891 | 77 | 0.0160 | 0.9744 |
| 85 | 0.0038 | 0.9912 | 84 | 0.0034 | 0.9943 | 78 | 0.0057 | 0.9704 |
| 86 | 0.0016 | 0.9965 | 86 | 0.0003 | 0.9989 | 79 | 0.0035 | 0.9873 |
| 87 | 0.0014 | 0.9965 | 87 | 0.0008 | 0.9970 | 80 | 0.0008 | 0.9961 |
| 89 | 0.0013 | 0.9954 | 88 | 0.0052 | 0.9877 | 81 | 0.0045 | 0.9907 |
| 90 | 0.0008 | 0.9982 | 89 | 0.0321 | 0.9356 | 83 | 0.0012 | 0.9966 |
| 91 | 0.0007 | 0.9980 | 90 | 0.0049 | 0.9826 | 84 | 0.0020 | 0.9949 |
| 92 | 0.0015 | 0.9948 | 91 | 0.0031 | 0.9906 | 85 | 0.0011 | 0.9991 |
| 93 | 0.0008 | 0.9979 | 93 | 0.0066 | 0.9806 | 86 | 0.0052 | 0.9891 |
| 94 | 0.0042 | 0.9863 | 94 | 0.0007 | 0.9977 | 87 | 0.0018 | 0.9944 |
| 95 | 0.0010 | 0.9972 | 95 | 0.0009 | 0.9973 | 88 | 0.0031 | 0.9902 |
| 97 | 0.0023 | 0.9921 | 96 | 0.0053 | 0.9787 | 89 | 0.0053 | 0.9839 |
| 98 | 0.0014 | 0.9953 | 97 | 0.0005 | 0.9992 | 91 | 0.0009 | 0.9978 |
| 99 | 0.0002 | 0.9995 | 98 | 0.0011 | 0.9965 | 92 | 0.0065 | 0.9855 |
| 100 | 0.0034 | 0.9919 | 99 | 0.0051 | 0.9838 | 93 | 0.0011 | 0.9969 |
| 101 | 0.0009 | 0.9985 | 101 | 0.0585 | 0.9083 | 94 | 0.0001 | 0.9993 |
| 102 | 0.0028 | 0.9926 | 103 | 0.0123 | 0.9656 | 96 | 0.0006 | 0.9978 |
| 103 | 0.0011 | 0.9948 | 104 | 0.0023 | 0.9976 | 97 | 0.0128 | 0.9661 |
| 104 | 0.0008 | 0.9969 | 105 | 0.0065 | 0.9966 | 98 | 0.0040 | 0.9924 |
| 105 | 0.0002 | 0.9991 | 106 | 0.0013 | 0.9962 | 99 | 0.0020 | 0.9938 |
| 106 | 0.0004 | 0.9985 | 107 | 0.0019 | 0.9947 | 100 | 0.0003 | 0.9991 |
| 108 | 0.0012 | 0.9970 | 108 | 0.0028 | 0.9893 | 101 | 0.0506 | 0.9735 |
| 109 | 0.0005 | 0.9987 | 111 | 0.0012 | 0.9962 | 103 | 0.0158 | 0.9668 |
| 110 | 0.0026 | 0.9901 | 112 | 0.0052 | 0.9873 | 105 | 0.0030 | 0.9870 |
| 111 | 0.0009 | 0.9976 | 113 | 0.0007 | 0.9978 | 106 | 0.0008 | 0.9978 |
| 112 | 0.0009 | 0.9980 | 114 | 0.0052 | 0.9915 | 108 | 0.0016 | 0.9954 |
| 113 | 0.0012 | 0.9973 | 116 | 0.0009 | 0.9980 | 109 | 0.0048 | 0.9762 |
| 114 | 0.0008 | 0.9978 | 117 | 0.0016 | 0.9952 | 111 | 0.0040 | 0.9921 |
| 115 | 0.0006 | 0.9984 | 118 | 0.0048 | 0.9837 | 112 | 0.0113 | 0.9930 |
| 116 | 0.0012 | 0.9959 | 119 | 0.0019 | 0.9939 | 113 | 0.0012 | 0.9967 |
| 117 | 0.0007 | 0.9978 | 120 | 0.0015 | 0.9941 | 114 | 0.0027 | 0.9927 |
| 118 | 0.0009 | 0.9971 | 121 | 0.0006 | 0.9979 | 115 | 0.0006 | 0.9979 |
| 119 | 3.9550* | 0.8304 | 122 | 0.0065 | 0.9780 | 116 | 0.0025 | 0.9895 |
| 121 | 0.0003 | 0.9985 | 123 | 0.0007 | 0.9975 | 118 | 0.0020 | 0.9951 |
| 122 | 0.0082 | 0.9973 | 124 | 0.0024 | 0.9919 | 119 | 0.0021 | 0.9933 |
| 124 | 0.0039 | 0.9879 | 125 | 0.0015 | 0.9943 | 120 | 0.0018 | 0.9941 |
| 125 | 0.0005 | 0.9994 | 127 | 0.0018 | 0.9962 | 121 | 0.0156 | 0.9674 |
| 126 | 0.0006 | 0.9970 | 128 | 0.0013 | 0.9967 | 123 | 0.0012 | 0.9976 |
| 129 | 0.0004 | 0.9989 | 130 | 0.0013 | 0.9940 | 124 | 0.0024 | 0.9909 |
| 130 | 0.0012 | 0.9968 | 131 | 0.0050 | 0.9893 | 125 | 0.0183 | 0.9566 |
| 131 | 0.0015 | 0.9947 | 132 | 0.0007 | 0.9975 | 127 | 0.0008 | 0.9972 |

| | | | | | | | | |
|---|---|---|---|---|---|---|---|---|
| 133 | 0.0048 | 0.9879 | 133 | 0.0055 | 0.9845 | 128 | 0.0168 | 0.9711 |
| 134 | 0.0010 | 0.9963 | 137 | 0.0023 | 0.9928 | 129 | 0.0013 | 0.9971 |
| 135 | 0.0013 | 0.9964 | 138 | 0.0004 | 0.9990 | **Average** | 0.0055 | 0.9905 |
| 137 | 0.0011 | 0.9964 | 139 | 0.0117 | 0.9625 | **STDEV** | 0.0158 | 0.0106 |
| 138 | 0.0005 | 0.9987 | 140 | 0.0098 | 0.9700 | | | |
| 139 | 0.0041 | 0.9943 | 141 | 0.0022 | 0.9941 | | | |
| 141 | 0.0009 | 0.9973 | 142 | 0.0027 | 0.9893 | | | |
| 145 | 0.0006 | 0.9980 | 143 | 0.0093 | 0.9747 | | | |
| 146 | 0.0011 | 0.9967 | 147 | 0.0008 | 0.9978 | | | |
| 147 | 0.0008 | 0.9982 | 148 | 0.0030 | 0.9887 | | | |
| 149 | 0.0003 | 0.9987 | 149 | 0.0009 | 0.9967 | | | |
| 150 | 0.0008 | 0.9964 | 150 | 0.0058 | 0.9800 | | | |
| 151 | 0.0010 | 0.9979 | 151 | 0.0004 | 0.9986 | | | |
| 152 | 0.0010 | 0.9979 | 153 | 0.0016 | 0.9958 | | | |
| 153 | 0.0002 | 0.9996 | 154 | 0.0062 | 0.9788 | | | |
| 154 | 0.0009 | 0.9981 | 155 | 0.0016 | 0.9951 | | | |
| 155 | 0.0023 | 0.9870 | 157 | 0.0032 | 0.9935 | | | |
| 156 | 0.0008 | 0.9965 | 161 | 0.0034 | 0.9912 | | | |
| 157 | 0.0008 | 0.9975 | 162 | 0.0146 | 0.9757 | | | |
| 158 | 0.0002 | 0.9992 | 163 | 0.0037 | 0.9840 | | | |
| 159 | 0.0026 | 0.9910 | 165 | 0.0020 | 0.9962 | | | |
| 160 | 0.0004 | 0.9988 | 167 | 0.0126 | 0.9747 | | | |
| 161 | 0.0028 | 0.9875 | 168 | 0.0042 | 0.9833 | | | |
| 163 | 0.0053 | 0.9959 | 169 | 0.0012 | 0.9962 | | | |
| 164 | 0.0002 | 0.9991 | 170 | 0.0004 | 0.9987 | | | |
| 166 | 0.0017 | 0.9944 | 171 | 0.0013 | 0.9960 | | | |
| 167 | 0.0011 | 0.9977 | 173 | 0.0005 | 0.9989 | | | |
| 168 | 0.0016 | 0.9963 | 174 | 0.0025 | 0.9941 | | | |
| 169 | 0.0006 | 0.9983 | 175 | 0.0074 | 0.9804 | | | |
| 170 | 0.0003 | 0.9991 | 176 | 0.0023 | 0.9930 | | | |
| 171 | 0.0012 | 0.9948 | 177 | 0.0004 | 0.9989 | | | |
| 172 | 0.0015 | 0.9957 | 178 | 0.0008 | 0.9964 | | | |
| 173 | 0.0003 | 0.9988 | 179 | 0.0037 | 0.9903 | | | |
| 174 | 0.0016 | 0.9940 | 180 | 0.0015 | 0.9950 | | | |
| 175 | 0.0047 | 0.9825 | 183 | 0.0005 | 0.9979 | | | |
| 176 | 0.0002 | 0.9991 | 184 | 0.0015 | 0.9959 | | | |
| 177 | 0.0007 | 0.9965 | 185 | 0.0015 | 0.9942 | | | |
| 178 | 0.0004 | 0.9988 | 186 | 0.0027 | 0.9939 | | | |
| 179 | 0.0021 | 0.9910 | 187 | 0.0014 | 0.9952 | | | |
| 181 | 0.0012 | 0.9956 | 188 | 0.0039 | 0.9888 | | | |
| 182 | 0.0002 | 0.9994 | 189 | 0.0005 | 0.9986 | | | |
| 183 | 0.0006 | 0.9983 | 190 | 0.0007 | 0.9981 | | | |
| 184 | 0.0036 | 0.9886 | 191 | 0.0117 | 0.9671 | | | |
| 185 | 0.0010 | 0.9968 | 192 | 0.0009 | 0.9971 | | | |
| 186 | 0.0008 | 0.9981 | 193 | 0.0015 | 0.9939 | | | |
| 187 | 0.0094 | 0.9835 | 194 | 0.0017 | 0.9961 | | | |
| 188 | 0.0022 | 0.9932 | 195 | 0.0033 | 0.9905 | | | |
| 189 | 0.0006 | 0.9980 | 197 | 0.0041 | 0.9914 | | | |
| 190 | 0.0059 | 0.9814 | 198 | 0.0016 | 0.9947 | | | |
| 191 | 0.0040 | 0.9887 | 199 | 0.0031 | 0.9924 | | | |
| 192 | 0.0047 | 0.9818 | 200 | 0.0026 | 0.9934 | | | |
| 193 | 0.0004 | 0.9985 | 201 | 0.0016 | 0.9968 | | | |
| 194 | 0.0004 | 0.9987 | 202 | 0.0040 | 0.9879 | | | |

| | | | | | |
|---|---|---|---|---|---|
| 196 | 0.0019 | 0.9927 | 204 | 0.0026 | 0.9918 |
| 197 | 0.0012 | 0.9952 | 206 | 0.0024 | 0.9959 |
| 198 | 0.0006 | 0.9984 | 207 | 0.0083 | 0.9771 |
| 199 | 0.0009 | 0.9976 | 208 | 0.0077 | 0.9819 |
| 200 | 0.0010 | 0.9960 | 209 | 0.0018 | 0.9960 |
| 201 | 0.0023 | 0.9929 | 210 | 0.0012 | 0.9955 |
| 202 | 0.0004 | 0.9981 | 211 | 0.0015 | 0.9954 |
| 203 | 0.0019 | 0.9931 | 212 | 0.0017 | 0.9949 |
| 204 | 0.0019 | 0.9920 | 213 | 0.0005 | 0.9988 |
| 205 | 0.0023 | 0.9901 | 216 | 0.0080 | 0.9765 |
| 206 | 0.0017 | 0.9960 | 217 | 0.0015 | 0.9945 |
| **Average** | 0.0031 | 0.9918 | 218 | 0.0010 | 0.9962 |
| **STDEV** | 0.0110 | 0.0215 | 219 | 0.0044 | 0.9804 |
| | | | 220 | 0.0046 | 0.9860 |
| | | | 221 | 0.0013 | 0.9949 |
| | | | 223 | 0.0020 | 0.9951 |
| | | | 224 | 0.0007 | 0.9975 |
| | | | 225 | 0.0032 | 0.9897 |
| | | | 227 | 0.0019 | 0.9958 |
| | | | 228 | 0.0005 | 0.9980 |
| | | | 229 | 0.0017 | 0.9963 |
| | | | 230 | 0.0035 | 0.9920 |
| | | | 233 | 0.0023 | 0.9933 |
| | | | 236 | 0.0013 | 0.9958 |
| | | | 237 | 0.0088 | 0.9779 |
| | | | 238 | 0.0027 | 0.9980 |
| | | | 239 | 0.0017 | 0.9948 |
| | | | 240 | 0.0022 | 0.9952 |
| | | | 242 | 0.0015 | 0.9950 |
| | | | 243 | 0.0014 | 0.9948 |
| | | | 244 | 0.0034 | 0.9885 |
| | | | 247 | 0.0137 | 0.9809 |
| | | | 249 | 0.0008 | 0.9958 |
| | | | 250 | 0.0016 | 0.9949 |
| | | | 251 | 0.0014 | 0.9956 |
| | | | 252 | 0.0014 | 0.9955 |
| | | | 254 | 0.0029 | 0.9895 |
| | | | 255 | 0.0018 | 0.9949 |
| | | | 257 | 0.0038 | 0.9892 |
| | | | 258 | 0.0023 | 0.9918 |
| | | | 260 | 0.0044 | 0.9894 |
| | | | 261 | 0.0080 | 0.9760 |
| | | | 262 | 0.0012 | 0.9957 |
| | | | 263 | 0.0012 | 0.9950 |
| | | | 264 | 0.0031 | 0.9910 |
| | | | 265 | 0.0119 | 0.9949 |
| | | | 266 | 0.0107 | 0.9737 |
| | | | 268 | 0.0017 | 0.9926 |
| | | | 269 | 0.0015 | 0.9959 |
| | | | 270 | 0.0010 | 0.9956 |
| | | | 271 | 0.0015 | 0.9973 |
| | | | 272 | 0.0042 | 0.9848 |
| | | | 274 | 0.0007 | 0.9977 |

| | | | | | |
|---|---|---|---|---|---|
| | | | 275 | 0.0019 | 0.9957 |
| | | | 276 | 0.0132 | 0.9807 |
| | | | 277 | 0.0009 | 0.9973 |
| | | | 278 | 0.1030 | 0.7832 |
| | | | **Average** | 0.0046 | 0.9891 |
| | | | **STDEV** | 0.0115 | 0.0183 |

**Table S5**. $P_{CC}$ and MSE values for $\chi^1$ dihedral angles for all residues (excluding, Ala, Gly, and Pro) for thaumatin, proteinase K, and lysozyme.

| Cysteine residue # | MSE | $P_{CC}$ |
|---|---|---|
| **Thaumatin** | | |
| 9 | 0.0006 | 0.9969 |
| 56 | 0.0007 | 0.9969 |
| 66 | 0.0010 | 0.9958 |
| 121 | 0.0003 | 0.9985 |
| 126 | 0.0006 | 0.9970 |
| 134 | 0.0010 | 0.9963 |
| 145 | 0.0006 | 0.9980 |
| 149 | 0.0003 | 0.9987 |
| 158 | 0.0002 | 0.9992 |
| 159 | 0.0026 | 0.9910 |
| 164 | 0.0002 | 0.9991 |
| 177 | 0.0007 | 0.9965 |
| 193 | 0.0004 | 0.9985 |
| 204 | 0.0019 | 0.9920 |
| **Average** | 0.0008 | 0.9968 |
| **Standard deviation** | 0.0007 | 0.0024 |
| **Maximum** | 0.0026 | 0.9992 |
| **Minimum** | 0.0002 | 0.9910 |
| **Lysozyme** | | |
| 6 | 0.0010 | 0.9956 |
| 30 | 0.0004 | 0.9984 |
| 64 | 0.0009 | 0.9963 |
| 76 | 0.0006 | 0.9969 |
| 80 | 0.0008 | 0.9961 |
| 94 | 0.0001 | 0.9993 |
| 115 | 0.0006 | 0.9979 |
| 127 | 0.0008 | 0.9972 |
| **Average** | 0.0007 | 0.9972 |
| **Standard deviation** | 0.0003 | 0.0012 |
| **Maximum** | 0.0010 | 0.9993 |
| **Minimum** | 0.0001 | 0.9956 |
| **Proteinase K** | | |
| 34 | 0.0005 | 0.9973 |
| 123 | 0.0007 | 0.9975 |
| 178 | 0.0008 | 0.9964 |
| 249 | 0.0008 | 0.9958 |
| **Average** | 0.0007 | 0.9967 |
| **Standard deviation** | 0.0001 | 0.0007 |
| **Maximum** | 0.0008 | 0.9975 |
| **Minimum** | 0.0005 | 0.9958 |

**Table S6**. $P_{CC}$ and MSE values for disulfide forming cysteine residues $\chi^1$ dihedral angles (extracted from Table S5).

| **Residue # / dihedral angle** | **MSE** | **$P_{CC}$** |
|---|---|---|
| | **Lysozyme** | |
| 35 $\chi^1$ | 0.0003 | 0.9990 |
| 52 $\chi^1$ | 0.0014 | 0.9962 |
| 35 $\chi^2$ | 0.0019 | 0.9943 |
| 52 $\chi^2$ | 0.0063 | 0.9743 |
| | **Proteinase K** | |
| 39 $\chi^1$ | 0.0021 | 0.9941 |
| 69 $\chi^1$ | 0.0018 | 0.9939 |
| 161 $\chi^1$ | 0.0034 | 0.9912 |
| 224 $\chi^1$ | 0.0007 | 0.9975 |
| 39 $\chi^2$ | 0.0029 | 0.9977 |
| 69 $\chi^2$ | 0.0025 | 0.9932 |
| 161 $\chi^2$ | 0.0207 | 0.9567 |
| 224 $\chi^2$ | N/A | N/A |
| **Average** | 0.0040 | 0.9898 |
| **Standard deviation** | 0.0055 | 0.0123 |

**Table S7**. $P_{CC}$ and MSE values for functional active site residues in proteinase K and lysozyme $\chi^1$ and $\chi^2$ dihedral angles ($\chi^1$ extracted from Table S5).

| Thaumatin refinement statistics multi-conformer models | | | | |
|---|---|---|---|---|
| Dataset | 1 | 2 | 3 | 4 |
| PDB code | 7LJW | 7LJZ | 7LK5 | 7LK6 |
| Resolution range (Å) | 38.26 -1.22 (1.26 -1.22) | 37.8 - 1.29 (1.33 - 1.29) | 37.82 - 1.38 (1.43 - 1.38) | 37.82 - 1.48 (1.53 - 1.48) |
| Reflections used in refinement | 78731 (6867) | 66525 (5655) | 54141 (4338) | 43991 (3270) |
| Rwork | 0.142 (0.332) | 0.1444 (0.335) | 0.146 (0.348) | 0.146 (0.317) |
| Rfree | 0.156 (0.340) | 0.1600 (0.338) | 0.166 (0.387) | 0.169 (0.335) |
| No. on non-hydrogen atoms | 3771 | 3766 | 3766 | 3764 |
| Protein | 3578 | 3578 | 3578 | 3578 |
| Ligand/ion | 20 | 20 | 20 | 20 |
| Water | 151 | 148 | 148 | 143 |
| RMS (bonds) | 0.009 | 0.009 | 0.009 | 0.009 |
| RMS (angles) | 1.06 | 1.08 | 1.08 | 1.07 |
| Average B-factor | 18.18 | 19.64 | 21.62 | 22.49 |
| Protein | 17.48 | 18.95 | 20.93 | 21.84 |
| Ligand/ion | 16.13 | 16.95 | 18.02 | 19.51 |
| Water | 32.85 | 34.57 | 36.57 | 36.85 |
| Ramachandran (%) | | | | |
| Favored | 96.6 | 96.1 | 95.1 | 94.6 |
| Allowed | 3.4 | 3.9 | 4.9 | 5.4 |
| Outliers | 0 | 0 | 0 | 0 |

**Table S8**. **Refinement statistics for multi-conformer thaumatin models obtained from increasingly damaged datasets from a single crystal at room temperature (277 K).** Multi-conformer models were refined as discrined in Materials and Methods and using diffraction datasets from **Table S1**. Refinement statistics were obtained from phenix (*phenix.table_one*) using the final refined models and reflections file.

| Proteinase K refinement statistics multi-conformer models | | | | |
|---|---|---|---|---|
| Dataset | 1 | 2 | 3 | 4 |
| PDB code | 7LPU | 7LPV | 7LQ8 | 7LQ9 |
| Resolution range (Å) | 33.91 - 1.02 (1.06 - 1.02) | 32.12 - 1.1 (1.14 - 1.1) | 33.83 - 1.301 (1.35 - 1.301) | 34.85 - 1.43 (1.48 - 1.43) |
| Reflections used in refinement | 120585 (11691) | 95793 (9247) | 58149 (5476) | 44021 (4101) |
| Rwork | 0.119 (0.287) | 0.123 (0.285) | 0.121 (0.294) | 0.124 (0.272) |
| Rfree | 0.138 (0.301) | 0.146 (0.289) | 0.151 (0.322) | 0.162 (0.317) |
| No. on non-hydrogen atoms | 6724 | 6723 | 6676 | 6676 |
| Protein | 6345 | 6345 | 6312 | 6312 |
| Ligand/ion | 22 | 22 | 22 | 22 |
| Water | 282 | 281 | 265 | 263 |
| RMS (bonds) | 0.007 | 0.007 | 0.007 | 0.007 |
| RMS (angles) | 0.95 | 0.95 | 0.92 | 0.92 |
| Average B-factor | 8.97 | 10.96 | 13.61 | 16.33 |
| Protein | 8.1 | 10.05 | 12.76 | 15.46 |
| Ligand/ion | 22.93 | 25.63 | 30.83 | 35.33 |
| Water | 23.57 | 26.29 | 28.23 | 31.12 |
| Ramachandran (%) | | | | |
| Favored | 96.03 | 95.67 | 95.67 | 94.95 |
| Allowed | 3.97 | 4.33 | 4.33 | 5.05 |
| Outliers | 0 | 0 | 0 | 0 |

**Table S9**. **Refinement statistics for multi-conformer proteinase K models obtained from increasingly damaged datasets from a single crystal at room temperature (277 K).** Multi-conformer models were refined as discrined in Materials and Methods and using diffraction datasets from **Table S2**. Refinement statistics were obtained from phenix (*phenix.table_one*) using the final refined models and reflections file.

| Lysozyme refinement statistics multi-conformer models | | | |
|---|---|---|---|
| Dataset | 1 | 2 | 3 |
| PDB code | 7LN9 | 7LOQ | 7LOR |
| Resolution range (Å) | 33.6 - 1.13 (1.171 - 1.13) | 34.54 - 1.301 (1.348 - 1.301) | 33.53 - 1.522 (1.576 - 1.522) |
| Reflections used in refinement | 42596 (4064) | 27852 (2526) | 17663 (1565) |
| Rwork | 0.136 (0.292) | 0.138 (0.318) | 0.145 (0.290) |
| Rfree | 0.156 (0.308) | 0.167 (0.347) | 0.197 (0.396) |
| No. on non-hydrogen atoms | 3065 | 3065 | 3051 |
| Protein | 2925 | 2925 | 2925 |
| Ligand/ion | 4 | 4 | 4 |
| Water | 113 | 113 | 102 |
| RMS (bonds) | 0.006 | 0.006 | 0.005 |
| RMS (angles) | 0.84 | 0.82 | 0.75 |
| Average B-factor | 15.8 | 17.81 | 22.5 |
| Protein | 15.2 | 17.19 | 22.01 |
| Ligand/ion | 22.25 | 25.1 | 30.13 |
| Water | 28.48 | 30.79 | 34.02 |
| Ramachandran (%) | | | |
| Favored | 96.06 | 93.7 | 95.28 |
| Allowed | 3.94 | 6.3 | 4.72 |
| Outliers | 0 | 0 | 0 |

**Table S10**. **Refinement statistics for multi-conformer lysozyme models obtained from increasingly damaged datasets from a single crystal at room temperature (277 K).** Multi-conformer models were refined as discrined in Materials and Methods and using diffraction datasets from **Table S3**. Refinement statistics were obtained from phenix (*phenix.table_one*) using the final refined models and reflections file.

| Residue # | (1-$S^2$) lysozyme crystal 1 | (1-$S^2$) lysozyme crystal 2 | Δ(1-$S^2$) |
|---|---|---|---|
| 1 | 0.413 | 0.406 | 0.01 |
| 2 | 0.399 | 0.383 | 0.02 |
| 3 | 0.361 | 0.347 | 0.01 |
| 4 | 0.405 | 0.374 | 0.03 |
| 5 | 0.408 | 0.387 | 0.02 |
| 6 | 0.35 | 0.311 | 0.04 |
| 7 | 0.364 | 0.341 | 0.02 |
| 8 | 0.345 | 0.341 | 0.00 |
| 9 | 0.366 | 0.36 | 0.01 |
| 10 | 0.379 | 0.39 | -0.01 |
| 11 | 0.392 | 0.406 | -0.01 |
| 12 | 0.384 | 0.363 | 0.02 |
| 13 | 0.404 | 0.402 | 0.00 |
| 14 | 0.667 | 0.632 | 0.04 |
| 15 | 0.465 | 0.461 | 0.00 |
| 16 | 0.493 | 0.482 | 0.01 |
| 17 | 0.514 | 0.498 | 0.02 |
| 18 | 0.769 | 0.714 | 0.06 |
| 19 | 0.592 | 0.594 | 0.00 |
| 20 | 0.437 | 0.437 | 0.00 |
| 21 | 0.565 | 0.602 | -0.04 |
| 22 | 0.402 | 0.376 | 0.03 |
| 23 | 0.344 | 0.339 | 0.01 |
| 24 | 0.399 | 0.405 | -0.01 |
| 25 | 0.388 | 0.373 | 0.02 |
| 26 | 0.327 | 0.307 | 0.02 |
| 27 | 0.335 | 0.3 | 0.03 |
| 28 | 0.333 | 0.317 | 0.02 |
| 29 | 0.329 | 0.281 | 0.05 |
| 30 | 0.296 | 0.272 | 0.02 |
| 31 | 0.292 | 0.315 | -0.02 |
| 32 | 0.328 | 0.288 | 0.04 |
| 33 | 0.312 | 0.289 | 0.02 |
| 34 | 0.327 | 0.327 | 0.00 |
| 35 | 0.344 | 0.327 | 0.02 |
| 36 | 0.322 | 0.29 | 0.03 |
| 37 | 0.481 | 0.439 | 0.04 |
| 38 | 0.35 | 0.314 | 0.04 |
| 39 | 0.381 | 0.403 | -0.02 |
| 40 | 0.309 | 0.313 | 0.00 |
| 41 | 0.295 | 0.281 | 0.01 |
| 42 | 0.297 | 0.295 | 0.00 |
| 43 | 0.313 | 0.314 | 0.00 |
| 44 | 0.42 | 0.392 | 0.03 |
| 45 | 0.484 | 0.481 | 0.00 |
| 46 | 0.598 | 0.592 | 0.01 |
| 47 | 0.776 | 0.832 | -0.06 |
| 48 | 0.462 | 0.376 | 0.09 |
| 49 | 0.422 | 0.402 | 0.02 |
| 50 | 0.377 | 0.344 | 0.03 |
| 51 | 0.367 | 0.365 | 0.00 |
| 52 | 0.319 | 0.337 | -0.02 |

| | | | |
|---|---|---|---|
| 53 | 0.296 | 0.277 | 0.02 |
| 54 | 0.286 | 0.271 | 0.02 |
| 55 | 0.45 | 0.374 | 0.08 |
| 56 | 0.322 | 0.315 | 0.01 |
| 57 | 0.281 | 0.281 | 0.00 |
| 58 | 0.333 | 0.322 | 0.01 |
| 59 | 0.69 | 0.664 | 0.03 |
| 60 | 0.339 | 0.304 | 0.03 |
| 61 | 0.512 | 0.562 | -0.05 |
| 62 | 0.482 | 0.497 | -0.02 |
| 63 | 0.347 | 0.347 | 0.00 |
| 64 | 0.311 | 0.311 | 0.00 |
| 65 | 0.408 | 0.408 | 0.00 |
| 66 | 0.334 | 0.332 | 0.00 |
| 67 | 0.36 | 0.362 | 0.00 |
| 68 | 0.386 | 0.394 | -0.01 |
| 69 | 0.359 | 0.355 | 0.00 |
| 70 | 0.607 | 0.625 | -0.02 |
| 71 | 0.543 | 0.598 | -0.06 |
| 72 | 0.645 | 0.61 | 0.04 |
| 73 | 0.634 | 0.706 | -0.07 |
| 74 | 0.451 | 0.444 | 0.01 |
| 75 | 0.448 | 0.448 | 0.00 |
| 76 | 0.388 | 0.368 | 0.02 |
| 77 | 0.834 | 0.815 | 0.02 |
| 78 | 0.659 | 0.654 | 0.01 |
| 79 | 0.592 | 0.545 | 0.05 |
| 80 | 0.304 | 0.301 | 0.00 |
| 81 | 0.699 | 0.698 | 0.00 |
| 82 | 0.465 | 0.494 | -0.03 |
| 83 | 0.377 | 0.354 | 0.02 |
| 84 | 0.388 | 0.361 | 0.03 |
| 85 | 0.478 | 0.444 | 0.03 |
| 86 | 0.818 | 0.841 | -0.02 |
| 87 | 0.459 | 0.447 | 0.01 |
| 88 | 0.447 | 0.382 | 0.06 |
| 89 | 0.416 | 0.41 | 0.01 |
| 90 | 0.44 | 0.397 | 0.04 |
| 91 | 0.393 | 0.375 | 0.02 |
| 92 | 0.424 | 0.387 | 0.04 |
| 93 | 0.606 | 0.571 | 0.04 |
| 94 | 0.384 | 0.364 | 0.02 |
| 95 | 0.384 | 0.365 | 0.02 |
| 96 | 0.39 | 0.405 | -0.02 |
| 97 | 0.862 | 0.853 | 0.01 |
| 98 | 0.39 | 0.363 | 0.03 |
| 99 | 0.362 | 0.368 | -0.01 |
| 100 | 0.532 | 0.489 | 0.04 |
| 101 | 0.716 | 0.725 | -0.01 |
| 102 | 0.581 | 0.529 | 0.05 |
| 103 | 0.709 | 0.687 | 0.02 |
| 104 | 0.39 | 0.387 | 0.00 |
| 105 | 0.326 | 0.295 | 0.03 |

| | | | |
|---|---|---|---|
| 106 | 0.357 | 0.327 | 0.03 |
| 107 | 0.381 | 0.382 | 0.00 |
| 108 | 0.32 | 0.294 | 0.03 |
| 109 | 0.81 | 0.787 | 0.02 |
| 110 | 0.429 | 0.384 | 0.05 |
| 111 | 0.324 | 0.316 | 0.01 |
| 112 | 0.725 | 0.707 | 0.02 |
| 113 | 0.379 | 0.345 | 0.03 |
| 114 | 0.385 | 0.368 | 0.02 |
| 115 | 0.331 | 0.308 | 0.02 |
| 116 | 0.365 | 0.37 | -0.01 |
| 117 | 0.459 | 0.421 | 0.04 |
| 118 | 0.392 | 0.401 | -0.01 |
| 119 | 0.477 | 0.486 | -0.01 |
| 120 | 0.436 | 0.426 | 0.01 |
| 121 | 0.809 | 0.77 | 0.04 |
| 122 | 0.472 | 0.539 | -0.07 |
| 123 | 0.396 | 0.369 | 0.03 |
| 124 | 0.401 | 0.387 | 0.01 |
| 125 | 0.911 | 0.92 | -0.01 |
| 126 | 0.444 | 0.42 | 0.02 |
| 127 | 0.373 | 0.374 | 0.00 |
| 128 | 0.865 | 0.86 | 0.01 |
| 129 | 0.703 | 0.687 | 0.02 |
| **Average** | 0.451 | 0.439 | 0.01 |

**Table S11. (1-$S^2$) values obtained from two independent lysozyme crystals at room temperature are highly similar.** (1-$S^2$) values were calculated from multi-conformer models (lysozyme crystal 1 and crystal 2, **Table S4**) and as described in Materials and Methods.

| | $(1-S^2)$ | | | | Slope $(1-S^2)$/MGy | Intercept (zero-dose $1-S^2$) | $R^2$ | $\Delta(1-S^2)$ |
|---|---|---|---|---|---|---|---|---|
| **Residue #** | **Dataset 1** | **Dataset 2** | **Dataset 3** | **Dataset 4** | | | | **Datasets 1 - 4** |
| **Dose (MGy)** | 0.016 | 0.114 | 0.212 | 0.31 | | | | |
| 1 | 0.557 | 0.594 | 0.671 | 0.687 | 0.477 | 0.55 | 0.946 | -0.130 |
| 2 | 0.823 | 0.844 | 0.872 | 0.870 | 0.172 | 0.824 | 0.877 | -0.047 |
| 3 | 0.403 | 0.416 | 0.443 | 0.492 | 0.300 | 0.39 | 0.930 | -0.089 |
| 4 | 0.424 | 0.451 | 0.486 | 0.531 | 0.363 | 0.414 | 0.987 | -0.107 |
| 5 | 0.346 | 0.374 | 0.384 | 0.395 | 0.160 | 0.349 | 0.932 | -0.049 |
| 6 | 0.335 | 0.377 | 0.439 | 0.449 | 0.412 | 0.333 | 0.941 | -0.114 |
| 7 | 0.320 | 0.388 | 0.409 | 0.417 | 0.318 | 0.332 | 0.836 | -0.097 |
| 8 | 0.350 | 0.362 | 0.408 | 0.395 | 0.185 | 0.349 | 0.736 | -0.045 |
| 9 | 0.320 | 0.346 | 0.375 | 0.409 | 0.302 | 0.313 | 0.996 | -0.089 |
| 10 | 0.459 | 0.494 | 0.567 | 0.613 | 0.546 | 0.444 | 0.983 | -0.154 |
| 11 | 0.404 | 0.459 | 0.525 | 0.568 | 0.569 | 0.396 | 0.994 | -0.164 |
| 12 | 0.417 | 0.447 | 0.507 | 0.515 | 0.361 | 0.413 | 0.932 | -0.098 |
| 13 | 0.408 | 0.386 | 0.455 | 0.474 | 0.272 | 0.386 | 0.716 | -0.066 |
| 14 | 0.355 | 0.385 | 0.413 | 0.438 | 0.283 | 0.352 | 0.998 | -0.083 |
| 15 | 0.405 | 0.420 | 0.442 | 0.538 | 0.430 | 0.381 | 0.826 | -0.133 |
| 16 | 0.396 | 0.386 | 0.427 | 0.441 | 0.180 | 0.383 | 0.776 | -0.045 |
| 17 | 0.451 | 0.470 | 0.555 | 0.588 | 0.506 | 0.434 | 0.943 | -0.137 |
| 18 | 0.501 | 0.497 | 0.552 | 0.618 | 0.414 | 0.474 | 0.860 | -0.117 |
| 19 | 0.584 | 0.600 | 0.681 | 0.710 | 0.468 | 0.567 | 0.935 | -0.126 |
| 20 | 0.590 | 0.640 | 0.739 | 0.755 | 0.606 | 0.582 | 0.938 | -0.165 |
| 21 | 0.718 | 0.772 | 0.860 | 0.905 | 0.662 | 0.706 | 0.985 | -0.187 |
| 22 | 0.553 | 0.640 | 0.706 | 0.733 | 0.618 | 0.557 | 0.952 | -0.180 |
| 23 | 0.565 | 0.589 | 0.640 | 0.677 | 0.395 | 0.553 | 0.983 | -0.112 |
| 24 | 0.522 | 0.532 | 0.616 | 0.667 | 0.530 | 0.498 | 0.931 | -0.145 |
| 25 | 0.572 | 0.660 | 0.649 | 0.692 | 0.356 | 0.585 | 0.784 | -0.120 |
| 26 | 0.651 | 0.755 | 0.760 | 0.832 | 0.559 | 0.658 | 0.902 | -0.181 |
| 27 | 0.483 | 0.539 | 0.625 | 0.695 | 0.737 | 0.465 | 0.994 | -0.212 |
| 28 | 0.374 | 0.371 | 0.462 | 0.496 | 0.466 | 0.35 | 0.876 | -0.122 |
| 29 | 0.645 | 0.680 | 0.699 | 0.728 | 0.273 | 0.643 | 0.988 | -0.083 |
| 30 | 0.421 | 0.459 | 0.512 | 0.516 | 0.345 | 0.421 | 0.920 | -0.095 |
| 31 | 0.426 | 0.446 | 0.480 | 0.498 | 0.255 | 0.421 | 0.985 | -0.072 |
| 32 | 0.408 | 0.414 | 0.451 | 0.448 | 0.160 | 0.404 | 0.819 | -0.040 |
| 33 | 0.384 | 0.404 | 0.453 | 0.501 | 0.408 | 0.369 | 0.971 | -0.117 |
| 34 | 0.338 | 0.376 | 0.437 | 0.454 | 0.417 | 0.333 | 0.962 | -0.116 |
| 35 | 0.336 | 0.374 | 0.400 | 0.460 | 0.406 | 0.326 | 0.972 | -0.124 |
| 36 | 0.413 | 0.471 | 0.484 | 0.512 | 0.316 | 0.418 | 0.922 | -0.099 |
| 37 | 0.498 | 0.553 | 0.577 | 0.628 | 0.422 | 0.495 | 0.980 | -0.130 |
| 38 | 0.535 | 0.545 | 0.641 | 0.681 | 0.545 | 0.512 | 0.920 | -0.146 |
| 39 | 0.472 | 0.533 | 0.576 | 0.625 | 0.512 | 0.468 | 0.995 | -0.153 |
| 40 | 0.870 | 0.872 | 0.890 | 0.908 | 0.135 | 0.863 | 0.919 | -0.038 |
| 41 | 0.523 | 0.549 | 0.595 | 0.606 | 0.301 | 0.519 | 0.954 | -0.083 |
| 42 | 0.865 | 0.891 | 0.914 | 0.928 | 0.216 | 0.864 | 0.983 | -0.063 |
| 43 | 0.722 | 0.755 | 0.797 | 0.846 | 0.422 | 0.711 | 0.993 | -0.124 |
| 44 | 0.526 | 0.568 | 0.594 | 0.656 | 0.424 | 0.517 | 0.974 | -0.130 |
| 45 | 0.548 | 0.559 | 0.612 | 0.626 | 0.293 | 0.539 | 0.926 | -0.078 |
| 46 | 0.821 | 0.834 | 0.861 | 0.855 | 0.132 | 0.821 | 0.806 | -0.034 |
| 47 | 0.445 | 0.468 | 0.553 | 0.565 | 0.454 | 0.434 | 0.913 | -0.120 |
| 48 | 0.381 | 0.392 | 0.408 | 0.478 | 0.313 | 0.364 | 0.826 | -0.097 |
| 49 | 0.422 | 0.456 | 0.481 | 0.510 | 0.295 | 0.419 | 0.996 | -0.088 |
| 50 | 0.386 | 0.406 | 0.464 | 0.456 | 0.273 | 0.383 | 0.830 | -0.070 |

| | | | | | | | | |
|---|---|---|---|---|---|---|---|---|
| 51 | 0.341 | 0.356 | 0.389 | 0.433 | 0.315 | 0.328 | 0.957 | -0.092 |
| 52 | 0.377 | 0.391 | 0.412 | 0.440 | 0.214 | 0.37 | 0.978 | -0.063 |
| 53 | 0.335 | 0.365 | 0.415 | 0.441 | 0.376 | 0.328 | 0.985 | -0.106 |
| 54 | 0.380 | 0.420 | 0.463 | 0.466 | 0.307 | 0.382 | 0.912 | -0.086 |
| 55 | 0.526 | 0.555 | 0.617 | 0.658 | 0.467 | 0.513 | 0.983 | -0.132 |
| 56 | 0.415 | 0.451 | 0.494 | 0.557 | 0.479 | 0.401 | 0.983 | -0.142 |
| 57 | 0.496 | 0.563 | 0.628 | 0.626 | 0.464 | 0.503 | 0.881 | -0.130 |
| 58 | 0.555 | 0.624 | 0.710 | 0.712 | 0.568 | 0.558 | 0.905 | -0.157 |
| 59 | 0.697 | 0.798 | 0.899 | 0.887 | 0.685 | 0.709 | 0.855 | -0.190 |
| 60 | 1.013 | 1.025 | 1.068 | 1.074 | 0.231 | 1.007 | 0.914 | -0.061 |
| 61 | 0.989 | 1.095 | 1.235 | 1.242 | 0.917 | 0.991 | 0.913 | -0.253 |
| 62 | 0.729 | 0.778 | 0.904 | 0.910 | 0.683 | 0.719 | 0.903 | -0.181 |
| 63 | 0.883 | 0.918 | 0.929 | 0.946 | 0.204 | 0.886 | 0.941 | -0.063 |
| 64 | 0.493 | 0.520 | 0.570 | 0.619 | 0.437 | 0.479 | 0.984 | -0.126 |
| 65 | 0.433 | 0.457 | 0.513 | 0.550 | 0.415 | 0.421 | 0.980 | -0.117 |
| 66 | 0.408 | 0.451 | 0.481 | 0.499 | 0.309 | 0.409 | 0.967 | -0.091 |
| 67 | 0.423 | 0.488 | 0.499 | 0.532 | 0.345 | 0.429 | 0.913 | -0.109 |
| 68 | 0.452 | 0.482 | 0.521 | 0.517 | 0.239 | 0.454 | 0.866 | -0.065 |
| 69 | 0.320 | 0.335 | 0.408 | 0.420 | 0.381 | 0.309 | 0.907 | -0.100 |
| 70 | 0.379 | 0.429 | 0.464 | 0.503 | 0.415 | 0.376 | 0.994 | -0.124 |
| 71 | 0.432 | 0.434 | 0.477 | 0.512 | 0.289 | 0.417 | 0.911 | -0.080 |
| 72 | 0.524 | 0.559 | 0.645 | 0.675 | 0.550 | 0.511 | 0.962 | -0.151 |
| 73 | 0.496 | 0.533 | 0.620 | 0.652 | 0.566 | 0.483 | 0.965 | -0.156 |
| 74 | 0.580 | 0.603 | 0.652 | 0.659 | 0.292 | 0.576 | 0.933 | -0.079 |
| 75 | 0.486 | 0.535 | 0.602 | 0.620 | 0.479 | 0.483 | 0.959 | -0.134 |
| 76 | 0.457 | 0.514 | 0.591 | 0.687 | 0.783 | 0.435 | 0.987 | -0.230 |
| 77 | 0.481 | 0.535 | 0.612 | 0.656 | 0.614 | 0.471 | 0.990 | -0.175 |
| 78 | 0.872 | 0.894 | 0.944 | 0.978 | 0.376 | 0.861 | 0.981 | -0.106 |
| 79 | 0.849 | 0.871 | 0.912 | 0.896 | 0.186 | 0.852 | 0.718 | -0.047 |
| 80 | 0.461 | 0.494 | 0.547 | 0.594 | 0.461 | 0.449 | 0.992 | -0.133 |
| 81 | 0.463 | 0.496 | 0.520 | 0.548 | 0.285 | 0.46 | 0.996 | -0.085 |
| 82 | 0.442 | 0.500 | 0.552 | 0.585 | 0.491 | 0.44 | 0.986 | -0.143 |
| 83 | 0.417 | 0.439 | 0.462 | 0.482 | 0.222 | 0.414 | 0.999 | -0.065 |
| 84 | 0.413 | 0.440 | 0.445 | 0.473 | 0.189 | 0.412 | 0.944 | -0.060 |
| 85 | 0.329 | 0.352 | 0.387 | 0.373 | 0.170 | 0.332 | 0.725 | -0.044 |
| 86 | 0.333 | 0.350 | 0.379 | 0.394 | 0.216 | 0.329 | 0.985 | -0.061 |
| 87 | 0.349 | 0.363 | 0.378 | 0.389 | 0.138 | 0.347 | 0.996 | -0.040 |
| 88 | 0.366 | 0.381 | 0.434 | 0.423 | 0.229 | 0.364 | 0.784 | -0.057 |
| 89 | 0.384 | 0.430 | 0.473 | 0.524 | 0.472 | 0.376 | 0.999 | -0.140 |
| 90 | 0.438 | 0.451 | 0.498 | 0.535 | 0.345 | 0.424 | 0.960 | -0.097 |
| 91 | 0.502 | 0.539 | 0.603 | 0.605 | 0.381 | 0.5 | 0.908 | -0.103 |
| 92 | 0.516 | 0.563 | 0.621 | 0.636 | 0.427 | 0.514 | 0.956 | -0.120 |
| 93 | 0.453 | 0.498 | 0.530 | 0.556 | 0.348 | 0.453 | 0.984 | -0.103 |
| 94 | 0.519 | 0.514 | 0.618 | 0.675 | 0.584 | 0.486 | 0.883 | -0.156 |
| 95 | 0.531 | 0.571 | 0.680 | 0.705 | 0.644 | 0.517 | 0.942 | -0.174 |
| 96 | 0.536 | 0.592 | 0.585 | 0.731 | 0.590 | 0.515 | 0.793 | -0.195 |
| 97 | 0.611 | 0.683 | 0.759 | 0.768 | 0.558 | 0.614 | 0.923 | -0.157 |
| 98 | 0.425 | 0.485 | 0.528 | 0.566 | 0.476 | 0.423 | 0.988 | -0.141 |
| 99 | 0.508 | 0.528 | 0.561 | 0.626 | 0.395 | 0.491 | 0.935 | -0.118 |
| 100 | 0.391 | 0.430 | 0.478 | 0.472 | 0.297 | 0.394 | 0.857 | -0.081 |
| 101 | 0.396 | 0.427 | 0.469 | 0.510 | 0.392 | 0.387 | 0.996 | -0.114 |
| 102 | 0.347 | 0.368 | 0.414 | 0.410 | 0.240 | 0.346 | 0.863 | -0.063 |
| 103 | 0.362 | 0.424 | 0.451 | 0.498 | 0.444 | 0.361 | 0.979 | -0.136 |

| | | | | | | | | |
|---|---|---|---|---|---|---|---|---|
| 104 | 0.373 | 0.403 | 0.445 | 0.463 | 0.318 | 0.369 | 0.980 | -0.090 |
| 105 | 0.399 | 0.414 | 0.465 | 0.492 | 0.337 | 0.388 | 0.962 | -0.093 |
| 106 | 0.426 | 0.453 | 0.475 | 0.489 | 0.215 | 0.426 | 0.981 | -0.063 |
| 107 | 0.338 | 0.369 | 0.390 | 0.416 | 0.260 | 0.336 | 0.995 | -0.078 |
| 108 | 0.333 | 0.357 | 0.378 | 0.394 | 0.208 | 0.332 | 0.992 | -0.061 |
| 109 | 0.345 | 0.355 | 0.403 | 0.444 | 0.352 | 0.329 | 0.946 | -0.099 |
| 110 | 0.565 | 0.573 | 0.614 | 0.619 | 0.207 | 0.559 | 0.896 | -0.054 |
| 111 | 0.387 | 0.439 | 0.471 | 0.524 | 0.452 | 0.382 | 0.991 | -0.137 |
| 112 | 0.783 | 0.778 | 0.800 | 0.798 | 0.068 | 0.779 | 0.629 | -0.015 |
| 113 | 0.399 | 0.448 | 0.480 | 0.510 | 0.372 | 0.399 | 0.985 | -0.111 |
| 114 | 0.357 | 0.407 | 0.415 | 0.474 | 0.366 | 0.354 | 0.934 | -0.117 |
| 115 | 0.495 | 0.528 | 0.633 | 0.639 | 0.548 | 0.484 | 0.898 | -0.144 |
| 116 | 0.552 | 0.590 | 0.699 | 0.693 | 0.543 | 0.545 | 0.865 | -0.141 |
| 117 | 0.454 | 0.516 | 0.524 | 0.598 | 0.449 | 0.45 | 0.928 | -0.144 |
| 118 | 0.488 | 0.553 | 0.660 | 0.699 | 0.755 | 0.477 | 0.973 | -0.211 |
| 119 | 1.043 | 1.132 | 1.314 | 1.305 | 0.988 | 1.037 | 0.879 | -0.262 |
| 120 | 0.695 | 0.743 | 0.884 | 0.875 | 0.695 | 0.686 | 0.860 | -0.180 |
| 121 | 0.591 | 0.679 | 0.707 | 0.765 | 0.561 | 0.594 | 0.960 | -0.174 |
| 122 | 0.823 | 0.859 | 0.967 | 0.992 | 0.628 | 0.808 | 0.939 | -0.169 |
| 123 | 0.510 | 0.591 | 0.608 | 0.649 | 0.443 | 0.517 | 0.923 | -0.139 |
| 124 | 0.477 | 0.562 | 0.592 | 0.647 | 0.551 | 0.48 | 0.964 | -0.170 |
| 125 | 0.699 | 0.729 | 0.770 | 0.792 | 0.327 | 0.694 | 0.988 | -0.093 |
| 126 | 0.437 | 0.516 | 0.530 | 0.543 | 0.339 | 0.451 | 0.810 | -0.106 |
| 127 | 0.517 | 0.546 | 0.602 | 0.668 | 0.519 | 0.499 | 0.973 | -0.151 |
| 128 | 0.523 | 0.542 | 0.604 | 0.630 | 0.391 | 0.511 | 0.958 | -0.107 |
| 129 | 0.466 | 0.519 | 0.570 | 0.660 | 0.646 | 0.448 | 0.979 | -0.194 |
| 130 | 0.435 | 0.463 | 0.518 | 0.505 | 0.270 | 0.436 | 0.801 | -0.070 |
| 131 | 0.462 | 0.491 | 0.590 | 0.578 | 0.456 | 0.456 | 0.829 | -0.116 |
| 132 | 0.456 | 0.491 | 0.571 | 0.610 | 0.553 | 0.442 | 0.975 | -0.154 |
| 133 | 0.468 | 0.494 | 0.514 | 0.554 | 0.284 | 0.461 | 0.979 | -0.086 |
| 134 | 0.414 | 0.466 | 0.502 | 0.513 | 0.340 | 0.418 | 0.929 | -0.099 |
| 135 | 0.571 | 0.568 | 0.607 | 0.690 | 0.404 | 0.543 | 0.809 | -0.119 |
| 136 | 0.679 | 0.712 | 0.752 | 0.797 | 0.402 | 0.669 | 0.995 | -0.118 |
| 137 | 0.574 | 0.602 | 0.611 | 0.696 | 0.383 | 0.558 | 0.848 | -0.122 |
| 138 | 0.432 | 0.453 | 0.475 | 0.468 | 0.133 | 0.435 | 0.778 | -0.036 |
| 139 | 0.663 | 0.681 | 0.755 | 0.788 | 0.458 | 0.647 | 0.950 | -0.125 |
| 140 | 0.597 | 0.621 | 0.685 | 0.679 | 0.316 | 0.594 | 0.853 | -0.082 |
| 141 | 0.857 | 0.903 | 0.971 | 0.934 | 0.305 | 0.867 | 0.639 | -0.077 |
| 142 | 0.948 | 0.970 | 1.057 | 1.066 | 0.450 | 0.937 | 0.901 | -0.118 |
| 143 | 0.685 | 0.691 | 0.822 | 0.810 | 0.516 | 0.668 | 0.777 | -0.125 |
| 144 | 0.406 | 0.420 | 0.478 | 0.470 | 0.255 | 0.402 | 0.811 | -0.064 |
| 145 | 0.337 | 0.379 | 0.395 | 0.466 | 0.411 | 0.327 | 0.938 | -0.129 |
| 146 | 0.418 | 0.406 | 0.465 | 0.438 | 0.121 | 0.412 | 0.355 | -0.020 |
| 147 | 0.330 | 0.360 | 0.402 | 0.408 | 0.282 | 0.329 | 0.936 | -0.078 |
| 148 | 0.325 | 0.359 | 0.399 | 0.426 | 0.350 | 0.32 | 0.995 | -0.101 |
| 149 | 0.308 | 0.354 | 0.401 | 0.401 | 0.333 | 0.312 | 0.892 | -0.093 |
| 150 | 0.353 | 0.381 | 0.407 | 0.459 | 0.351 | 0.343 | 0.970 | -0.106 |
| 151 | 0.357 | 0.384 | 0.439 | 0.426 | 0.267 | 0.358 | 0.799 | -0.069 |
| 152 | 0.357 | 0.377 | 0.412 | 0.397 | 0.158 | 0.36 | 0.699 | -0.040 |
| 153 | 0.350 | 0.369 | 0.393 | 0.398 | 0.171 | 0.35 | 0.948 | -0.048 |
| 154 | 0.323 | 0.349 | 0.392 | 0.386 | 0.237 | 0.324 | 0.850 | -0.063 |

| | | | | | | | | |
|---|---|---|---|---|---|---|---|---|
| 155 | 0.877 | 0.865 | 0.889 | 0.907 | 0.116 | 0.866 | 0.675 | -0.030 |
| 156 | 0.349 | 0.388 | 0.423 | 0.456 | 0.363 | 0.345 | 0.999 | -0.107 |
| 157 | 0.325 | 0.335 | 0.374 | 0.401 | 0.272 | 0.314 | 0.958 | -0.076 |
| 158 | 0.361 | 0.381 | 0.403 | 0.459 | 0.322 | 0.348 | 0.930 | -0.098 |
| 159 | 0.795 | 0.811 | 0.850 | 0.830 | 0.147 | 0.798 | 0.611 | -0.035 |
| 160 | 0.838 | 0.855 | 0.859 | 0.887 | 0.154 | 0.835 | 0.920 | -0.049 |
| 161 | 0.524 | 0.568 | 0.666 | 0.711 | 0.672 | 0.508 | 0.974 | -0.187 |
| 162 | 0.557 | 0.595 | 0.685 | 0.746 | 0.670 | 0.536 | 0.979 | -0.189 |
| 163 | 0.709 | 0.762 | 0.755 | 0.849 | 0.421 | 0.7 | 0.832 | -0.140 |
| 164 | 0.444 | 0.482 | 0.513 | 0.581 | 0.451 | 0.431 | 0.968 | -0.137 |
| 165 | 0.431 | 0.472 | 0.503 | 0.583 | 0.497 | 0.416 | 0.955 | -0.152 |
| 166 | 0.595 | 0.618 | 0.650 | 0.644 | 0.183 | 0.597 | 0.833 | -0.049 |
| 167 | 0.350 | 0.370 | 0.414 | 0.429 | 0.287 | 0.344 | 0.964 | -0.079 |
| 168 | 0.852 | 0.867 | 0.885 | 0.888 | 0.129 | 0.852 | 0.938 | -0.036 |
| 169 | 0.360 | 0.375 | 0.400 | 0.447 | 0.292 | 0.348 | 0.940 | -0.087 |
| 170 | 0.374 | 0.409 | 0.438 | 0.441 | 0.235 | 0.377 | 0.906 | -0.067 |
| 171 | 0.334 | 0.374 | 0.401 | 0.428 | 0.315 | 0.333 | 0.989 | -0.094 |
| 172 | 0.391 | 0.443 | 0.476 | 0.458 | 0.239 | 0.403 | 0.682 | -0.067 |
| 173 | 0.377 | 0.395 | 0.419 | 0.433 | 0.196 | 0.374 | 0.991 | -0.056 |
| 174 | 0.391 | 0.393 | 0.449 | 0.500 | 0.391 | 0.37 | 0.905 | -0.109 |
| 175 | 0.478 | 0.480 | 0.537 | 0.605 | 0.447 | 0.452 | 0.890 | -0.127 |
| 176 | 0.457 | 0.481 | 0.528 | 0.523 | 0.250 | 0.457 | 0.859 | -0.066 |
| 177 | 0.371 | 0.412 | 0.453 | 0.507 | 0.458 | 0.361 | 0.995 | -0.136 |
| 178 | 0.689 | 0.725 | 0.741 | 0.753 | 0.212 | 0.692 | 0.932 | -0.064 |
| 179 | 0.658 | 0.734 | 0.768 | 0.821 | 0.534 | 0.658 | 0.977 | -0.163 |
| 180 | 0.439 | 0.478 | 0.507 | 0.535 | 0.323 | 0.437 | 0.993 | -0.096 |
| 181 | 0.397 | 0.423 | 0.456 | 0.465 | 0.242 | 0.396 | 0.959 | -0.068 |
| 182 | 0.373 | 0.396 | 0.430 | 0.468 | 0.326 | 0.364 | 0.989 | -0.095 |
| 183 | 0.397 | 0.434 | 0.476 | 0.505 | 0.373 | 0.392 | 0.995 | -0.108 |
| 184 | 0.506 | 0.565 | 0.646 | 0.606 | 0.389 | 0.517 | 0.676 | -0.100 |
| 185 | 0.500 | 0.558 | 0.581 | 0.622 | 0.397 | 0.501 | 0.973 | -0.122 |
| 186 | 0.454 | 0.464 | 0.519 | 0.557 | 0.371 | 0.438 | 0.945 | -0.103 |
| 187 | 0.684 | 0.663 | 0.823 | 0.802 | 0.524 | 0.658 | 0.668 | -0.118 |
| 188 | 0.852 | 0.881 | 0.933 | 0.931 | 0.295 | 0.851 | 0.886 | -0.079 |
| 189 | 0.597 | 0.648 | 0.698 | 0.816 | 0.721 | 0.572 | 0.948 | -0.219 |
| 190 | 0.649 | 0.702 | 0.785 | 0.854 | 0.712 | 0.631 | 0.993 | -0.205 |
| 191 | 0.549 | 0.587 | 0.677 | 0.732 | 0.652 | 0.53 | 0.978 | -0.183 |
| 192 | 0.547 | 0.587 | 0.657 | 0.672 | 0.454 | 0.542 | 0.950 | -0.125 |
| 193 | 0.485 | 0.578 | 0.660 | 0.653 | 0.598 | 0.497 | 0.860 | -0.168 |
| 194 | 0.618 | 0.682 | 0.768 | 0.889 | 0.917 | 0.59 | 0.980 | -0.271 |
| 195 | 0.480 | 0.501 | 0.597 | 0.583 | 0.413 | 0.473 | 0.803 | -0.103 |
| 196 | 0.549 | 0.576 | 0.667 | 0.674 | 0.476 | 0.539 | 0.901 | -0.125 |
| 197 | 0.499 | 0.557 | 0.583 | 0.587 | 0.296 | 0.508 | 0.851 | -0.088 |
| 198 | 0.484 | 0.519 | 0.600 | 0.661 | 0.624 | 0.464 | 0.980 | -0.177 |
| 199 | 0.433 | 0.476 | 0.507 | 0.530 | 0.329 | 0.433 | 0.981 | -0.097 |
| 200 | 0.465 | 0.489 | 0.509 | 0.572 | 0.348 | 0.452 | 0.922 | -0.107 |

| | | | | | | | | |
|---|---|---|---|---|---|---|---|---|
| 201 | 0.404 | 0.435 | 0.474 | 0.452 | 0.187 | 0.411 | 0.640 | -0.048 |
| 202 | 0.359 | 0.382 | 0.421 | 0.412 | 0.202 | 0.361 | 0.810 | -0.053 |
| 203 | 0.329 | 0.354 | 0.359 | 0.403 | 0.232 | 0.323 | 0.907 | -0.074 |
| 204 | 0.363 | 0.393 | 0.457 | 0.457 | 0.353 | 0.36 | 0.895 | -0.094 |
| 205 | 0.409 | 0.458 | 0.513 | 0.510 | 0.365 | 0.413 | 0.879 | -0.101 |
| 206 | 0.631 | 0.665 | 0.772 | 0.762 | 0.510 | 0.624 | 0.845 | -0.131 |
| 207 | 0.728 | 0.800 | 0.889 | 0.893 | 0.596 | 0.73 | 0.911 | -0.165 |
| **Average** | 0.501 | 0.535 | 0.585 | 0.611 | 0.390 | 0.494 | 0.912 | -0.111 |

**Table S12. (1-$S^2$) values obtained from increasingly X-ray damaged thaumatin room temperature (277 K) datasets.** (1-$S^2$) values were calculated from thaumatin multi-conformer models (**Table S8**) and as described in Materials and Methods. Slopes, correlation coefficients, and intercepts were obtained via linear regression.

| | $(1-S^2)$ | | | | Slope $(1-S^2)$/MGy | Intercept (zero-dose $1-S^2$) | $R^2$ | $\Delta(1-S^2)$ |
|---|---|---|---|---|---|---|---|---|
| **Residue #** | **Dataset 1** | **Dataset 2** | **Dataset 3** | **Dataset 4** | | | | **Datasets 1-4** |
| **Dose (MGy)** | 0.006 | 0.023 | 0.041 | 0.058 | | | | |
| 1 | 0.364 | 0.402 | 0.540 | 0.538 | 3.808 | 0.339 | 0.876 | -0.174 |
| 2 | 0.304 | 0.356 | 0.438 | 0.475 | 3.424 | 0.284 | 0.984 | -0.171 |
| 3 | 0.217 | 0.270 | 0.337 | 0.429 | 4.039 | 0.184 | 0.984 | -0.212 |
| 4 | 0.297 | 0.336 | 0.420 | 0.459 | 3.281 | 0.273 | 0.979 | -0.162 |
| 5 | 0.712 | 0.733 | 0.773 | 0.786 | 1.509 | 0.703 | 0.970 | -0.074 |
| 6 | 0.245 | 0.307 | 0.372 | 0.431 | 3.581 | 0.224 | 1.000 | -0.186 |
| 7 | 0.229 | 0.291 | 0.366 | 0.467 | 4.533 | 0.193 | 0.987 | -0.238 |
| 8 | 0.196 | 0.243 | 0.340 | 0.382 | 3.771 | 0.170 | 0.978 | -0.186 |
| 9 | 0.191 | 0.237 | 0.278 | 0.353 | 3.026 | 0.168 | 0.978 | -0.162 |
| 10 | 0.234 | 0.269 | 0.360 | 0.423 | 3.787 | 0.200 | 0.978 | -0.189 |
| 11 | 0.233 | 0.272 | 0.333 | 0.430 | 3.746 | 0.197 | 0.961 | -0.197 |
| 12 | 0.195 | 0.249 | 0.325 | 0.453 | 4.882 | 0.149 | 0.961 | -0.258 |
| 13 | 0.521 | 0.571 | 0.599 | 0.673 | 2.777 | 0.502 | 0.966 | -0.152 |
| 14 | 0.191 | 0.248 | 0.367 | 0.396 | 4.228 | 0.165 | 0.957 | -0.205 |
| 15 | 0.738 | 0.775 | 0.790 | 0.844 | 1.910 | 0.726 | 0.951 | -0.106 |
| 16 | 0.281 | 0.374 | 0.431 | 0.555 | 5.044 | 0.249 | 0.978 | -0.274 |
| 17 | 0.310 | 0.368 | 0.464 | 0.523 | 4.229 | 0.281 | 0.992 | -0.213 |
| 18 | 0.259 | 0.307 | 0.427 | 0.422 | 3.513 | 0.241 | 0.882 | -0.163 |
| 19 | 0.330 | 0.386 | 0.474 | 0.567 | 4.593 | 0.292 | 0.989 | -0.237 |
| 20 | 0.298 | 0.396 | 0.467 | 0.591 | 5.454 | 0.263 | 0.987 | -0.293 |
| 21 | 0.300 | 0.345 | 0.444 | 0.582 | 5.431 | 0.244 | 0.954 | -0.282 |
| 22 | 0.270 | 0.306 | 0.391 | 0.480 | 4.112 | 0.230 | 0.971 | -0.210 |
| 23 | 0.193 | 0.234 | 0.310 | 0.356 | 3.251 | 0.169 | 0.989 | -0.163 |
| 24 | 0.273 | 0.334 | 0.407 | 0.478 | 3.954 | 0.246 | 0.999 | -0.205 |
| 25 | 0.206 | 0.278 | 0.348 | 0.396 | 3.679 | 0.189 | 0.993 | -0.190 |
| 26 | 0.222 | 0.271 | 0.318 | 0.426 | 3.783 | 0.188 | 0.951 | -0.204 |
| 27 | 0.663 | 0.662 | 0.724 | 0.782 | 2.412 | 0.631 | 0.892 | -0.119 |
| 28 | 0.286 | 0.347 | 0.399 | 0.508 | 4.122 | 0.253 | 0.968 | -0.222 |
| 29 | 0.258 | 0.325 | 0.391 | 0.486 | 4.308 | 0.227 | 0.990 | -0.228 |
| 30 | 0.202 | 0.253 | 0.301 | 0.352 | 2.861 | 0.185 | 0.999 | -0.150 |
| 31 | 0.653 | 0.706 | 0.732 | 0.788 | 2.473 | 0.641 | 0.979 | -0.135 |
| 32 | 0.220 | 0.262 | 0.324 | 0.399 | 3.443 | 0.191 | 0.985 | -0.179 |
| 33 | 0.342 | 0.391 | 0.459 | 0.525 | 3.547 | 0.316 | 0.996 | -0.183 |
| 34 | 0.226 | 0.286 | 0.354 | 0.395 | 3.306 | 0.209 | 0.992 | -0.169 |
| 35 | 0.252 | 0.310 | 0.278 | 0.381 | 2.025 | 0.240 | 0.665 | -0.129 |
| 36 | 0.193 | 0.231 | 0.315 | 0.364 | 3.436 | 0.166 | 0.983 | -0.171 |
| 37 | 0.213 | 0.266 | 0.344 | 0.393 | 3.555 | 0.190 | 0.994 | -0.180 |
| 38 | 0.190 | 0.229 | 0.295 | 0.409 | 4.153 | 0.148 | 0.947 | -0.219 |
| 39 | 0.185 | 0.227 | 0.272 | 0.342 | 2.964 | 0.162 | 0.983 | -0.157 |
| 40 | 0.180 | 0.233 | 0.269 | 0.350 | 3.133 | 0.158 | 0.972 | -0.170 |
| 41 | 0.203 | 0.235 | 0.300 | 0.349 | 2.894 | 0.179 | 0.987 | -0.146 |
| 42 | 0.222 | 0.285 | 0.351 | 0.418 | 3.758 | 0.199 | 1.000 | -0.196 |
| 43 | 0.265 | 0.345 | 0.412 | 0.461 | 3.764 | 0.250 | 0.989 | -0.196 |
| 44 | 0.324 | 0.370 | 0.468 | 0.493 | 3.485 | 0.302 | 0.958 | -0.169 |
| 45 | 0.372 | 0.445 | 0.508 | 0.515 | 2.830 | 0.369 | 0.912 | -0.143 |
| 46 | 0.249 | 0.291 | 0.350 | 0.422 | 3.322 | 0.222 | 0.987 | -0.173 |
| 47 | 0.229 | 0.295 | 0.394 | 0.439 | 4.195 | 0.205 | 0.985 | -0.210 |
| 48 | 0.185 | 0.246 | 0.316 | 0.372 | 3.627 | 0.164 | 0.999 | -0.187 |

| | | | | | | | | |
|---|---|---|---|---|---|---|---|---|
| 49 | 0.213 | 0.248 | 0.346 | 0.410 | 3.966 | 0.177 | 0.975 | -0.197 |
| 50 | 0.292 | 0.322 | 0.357 | 0.445 | 2.836 | 0.263 | 0.925 | -0.153 |
| 51 | 0.262 | 0.308 | 0.395 | 0.450 | 3.746 | 0.234 | 0.989 | -0.188 |
| 52 | 0.165 | 0.243 | 0.284 | 0.331 | 3.094 | 0.157 | 0.976 | -0.166 |
| 53 | 0.246 | 0.273 | 0.353 | 0.509 | 4.992 | 0.186 | 0.900 | -0.263 |
| 54 | 0.882 | 0.883 | 0.898 | 0.902 | 0.433 | 0.877 | 0.900 | -0.020 |
| 55 | 0.252 | 0.307 | 0.355 | 0.412 | 3.033 | 0.234 | 0.998 | -0.160 |
| 56 | 0.243 | 0.297 | 0.390 | 0.454 | 4.176 | 0.212 | 0.992 | -0.211 |
| 57 | 0.242 | 0.310 | 0.372 | 0.455 | 4.026 | 0.216 | 0.995 | -0.213 |
| 58 | 0.198 | 0.256 | 0.345 | 0.416 | 4.273 | 0.167 | 0.996 | -0.218 |
| 59 | 0.234 | 0.261 | 0.375 | 0.460 | 4.559 | 0.187 | 0.957 | -0.226 |
| 60 | 0.237 | 0.294 | 0.384 | 0.494 | 4.948 | 0.194 | 0.981 | -0.257 |
| 61 | 0.205 | 0.257 | 0.362 | 0.478 | 5.312 | 0.156 | 0.975 | -0.273 |
| 62 | 0.309 | 0.326 | 0.490 | 0.586 | 5.732 | 0.244 | 0.932 | -0.277 |
| 63 | 0.755 | 0.783 | 0.832 | 0.854 | 1.991 | 0.742 | 0.983 | -0.099 |
| 64 | 0.788 | 0.798 | 0.825 | 0.850 | 1.225 | 0.776 | 0.970 | -0.062 |
| 65 | 0.207 | 0.262 | 0.302 | 0.389 | 3.363 | 0.182 | 0.972 | -0.182 |
| 66 | 0.213 | 0.273 | 0.348 | 0.443 | 4.396 | 0.179 | 0.989 | -0.230 |
| 67 | 0.204 | 0.237 | 0.318 | 0.376 | 3.435 | 0.174 | 0.980 | -0.172 |
| 68 | 0.181 | 0.233 | 0.294 | 0.384 | 3.849 | 0.150 | 0.983 | -0.203 |
| 69 | 0.223 | 0.270 | 0.338 | 0.401 | 3.461 | 0.197 | 0.995 | -0.178 |
| 70 | 0.167 | 0.237 | 0.285 | 0.372 | 3.806 | 0.143 | 0.986 | -0.205 |
| 71 | 0.199 | 0.243 | 0.289 | 0.365 | 3.124 | 0.174 | 0.979 | -0.166 |
| 72 | 0.180 | 0.236 | 0.276 | 0.340 | 2.985 | 0.162 | 0.991 | -0.160 |
| 73 | 0.173 | 0.215 | 0.284 | 0.370 | 3.793 | 0.139 | 0.978 | -0.197 |
| 74 | 0.171 | 0.224 | 0.320 | 0.348 | 3.610 | 0.150 | 0.966 | -0.177 |
| 75 | 0.158 | 0.198 | 0.255 | 0.282 | 2.468 | 0.144 | 0.986 | -0.124 |
| 76 | 0.172 | 0.206 | 0.285 | 0.362 | 3.732 | 0.137 | 0.975 | -0.190 |
| 77 | 0.178 | 0.221 | 0.272 | 0.381 | 3.789 | 0.142 | 0.945 | -0.203 |
| 78 | 0.226 | 0.271 | 0.319 | 0.388 | 3.067 | 0.203 | 0.988 | -0.162 |
| 79 | 0.170 | 0.209 | 0.306 | 0.329 | 3.307 | 0.148 | 0.951 | -0.159 |
| 80 | 0.225 | 0.303 | 0.365 | 0.443 | 4.112 | 0.202 | 0.997 | -0.218 |
| 81 | 0.203 | 0.292 | 0.324 | 0.416 | 3.848 | 0.186 | 0.966 | -0.213 |
| 82 | 0.211 | 0.255 | 0.335 | 0.391 | 3.567 | 0.184 | 0.991 | -0.180 |
| 83 | 0.175 | 0.223 | 0.274 | 0.315 | 2.707 | 0.160 | 0.999 | -0.140 |
| 84 | 0.183 | 0.229 | 0.295 | 0.362 | 3.466 | 0.156 | 0.994 | -0.179 |
| 85 | 0.187 | 0.239 | 0.314 | 0.417 | 4.396 | 0.149 | 0.978 | -0.230 |
| 86 | 0.211 | 0.273 | 0.344 | 0.379 | 3.307 | 0.196 | 0.985 | -0.168 |
| 87 | 0.229 | 0.270 | 0.317 | 0.402 | 3.250 | 0.200 | 0.966 | -0.173 |
| 88 | 0.213 | 0.255 | 0.315 | 0.431 | 4.100 | 0.172 | 0.945 | -0.218 |
| 89 | 0.718 | 0.741 | 0.747 | 0.772 | 0.963 | 0.714 | 0.951 | -0.054 |
| 90 | 0.219 | 0.272 | 0.328 | 0.384 | 3.166 | 0.199 | 1.000 | -0.165 |
| 91 | 0.217 | 0.271 | 0.344 | 0.361 | 2.907 | 0.205 | 0.957 | -0.144 |
| 92 | 0.172 | 0.236 | 0.269 | 0.376 | 3.699 | 0.145 | 0.950 | -0.204 |
| 93 | 0.204 | 0.251 | 0.298 | 0.413 | 3.869 | 0.168 | 0.940 | -0.209 |
| 94 | 0.167 | 0.248 | 0.299 | 0.356 | 3.549 | 0.154 | 0.988 | -0.189 |
| 95 | 0.209 | 0.264 | 0.334 | 0.386 | 3.456 | 0.188 | 0.998 | -0.177 |
| 96 | 0.225 | 0.284 | 0.417 | 0.446 | 4.586 | 0.196 | 0.951 | -0.221 |
| 97 | 0.303 | 0.329 | 0.477 | 0.511 | 4.452 | 0.263 | 0.920 | -0.208 |
| 98 | 0.321 | 0.342 | 0.436 | 0.455 | 2.860 | 0.297 | 0.924 | -0.134 |
| 99 | 0.353 | 0.427 | 0.583 | 0.637 | 5.804 | 0.314 | 0.970 | -0.284 |
| 100 | 0.314 | 0.378 | 0.435 | 0.490 | 3.361 | 0.297 | 0.998 | -0.176 |
| 101 | 0.840 | 0.872 | 0.909 | 0.912 | 1.456 | 0.837 | 0.921 | -0.072 |

| | | | | | | | | |
|---|---|---|---|---|---|---|---|---|
| 102 | 0.341 | 0.407 | 0.453 | 0.579 | 4.361 | 0.305 | 0.951 | -0.238 |
| 103 | 0.395 | 0.482 | 0.606 | 0.681 | 5.649 | 0.360 | 0.994 | -0.286 |
| 104 | 0.348 | 0.416 | 0.476 | 0.512 | 3.173 | 0.336 | 0.983 | -0.164 |
| 105 | 0.712 | 0.727 | 0.757 | 0.786 | 1.449 | 0.699 | 0.982 | -0.074 |
| 106 | 0.264 | 0.300 | 0.378 | 0.490 | 4.345 | 0.219 | 0.952 | -0.226 |
| 107 | 0.266 | 0.319 | 0.401 | 0.466 | 3.922 | 0.237 | 0.995 | -0.200 |
| 108 | 0.241 | 0.293 | 0.365 | 0.411 | 3.347 | 0.220 | 0.995 | -0.170 |
| 109 | 0.247 | 0.314 | 0.378 | 0.487 | 4.502 | 0.212 | 0.981 | -0.240 |
| 110 | 0.205 | 0.262 | 0.335 | 0.408 | 3.920 | 0.177 | 0.997 | -0.203 |
| 111 | 0.296 | 0.338 | 0.407 | 0.433 | 2.763 | 0.280 | 0.977 | -0.137 |
| 112 | 0.255 | 0.303 | 0.410 | 0.494 | 4.740 | 0.214 | 0.983 | -0.239 |
| 113 | 0.248 | 0.299 | 0.402 | 0.442 | 3.944 | 0.222 | 0.975 | -0.194 |
| 114 | 0.186 | 0.240 | 0.280 | 0.366 | 3.329 | 0.161 | 0.972 | -0.180 |
| 115 | 0.248 | 0.317 | 0.377 | 0.438 | 3.620 | 0.229 | 0.998 | -0.190 |
| 116 | 0.214 | 0.274 | 0.347 | 0.376 | 3.216 | 0.200 | 0.977 | -0.162 |
| 117 | 0.234 | 0.283 | 0.398 | 0.447 | 4.341 | 0.202 | 0.974 | -0.213 |
| 118 | 0.306 | 0.334 | 0.372 | 0.435 | 2.441 | 0.284 | 0.965 | -0.129 |
| 119 | 0.256 | 0.329 | 0.406 | 0.434 | 3.515 | 0.244 | 0.968 | -0.178 |
| 120 | 0.406 | 0.542 | 0.604 | 0.582 | 3.391 | 0.425 | 0.736 | -0.176 |
| 121 | 0.264 | 0.356 | 0.427 | 0.514 | 4.715 | 0.239 | 0.997 | -0.250 |
| 122 | 0.717 | 0.784 | 0.799 | 0.824 | 1.927 | 0.719 | 0.895 | -0.107 |
| 123 | 0.229 | 0.282 | 0.371 | 0.391 | 3.311 | 0.212 | 0.956 | -0.162 |
| 124 | 0.221 | 0.270 | 0.358 | 0.444 | 4.353 | 0.184 | 0.986 | -0.223 |
| 125 | 0.228 | 0.261 | 0.335 | 0.363 | 2.758 | 0.208 | 0.971 | -0.135 |
| 126 | 0.227 | 0.292 | 0.357 | 0.450 | 4.216 | 0.197 | 0.990 | -0.223 |
| 127 | 0.220 | 0.239 | 0.314 | 0.368 | 2.987 | 0.190 | 0.960 | -0.148 |
| 128 | 0.187 | 0.236 | 0.297 | 0.357 | 3.282 | 0.164 | 0.998 | -0.170 |
| 129 | 0.220 | 0.272 | 0.302 | 0.482 | 4.678 | 0.169 | 0.852 | -0.262 |
| 130 | 0.201 | 0.259 | 0.297 | 0.391 | 3.489 | 0.175 | 0.965 | -0.190 |
| 131 | 0.272 | 0.304 | 0.339 | 0.406 | 2.509 | 0.250 | 0.963 | -0.134 |
| 132 | 0.192 | 0.244 | 0.333 | 0.409 | 4.256 | 0.158 | 0.992 | -0.217 |
| 133 | 0.255 | 0.304 | 0.420 | 0.516 | 5.172 | 0.208 | 0.980 | -0.261 |
| 134 | 0.267 | 0.332 | 0.422 | 0.546 | 5.326 | 0.221 | 0.980 | -0.279 |
| 135 | 0.247 | 0.320 | 0.413 | 0.479 | 4.537 | 0.220 | 0.997 | -0.232 |
| 136 | 0.254 | 0.292 | 0.396 | 0.508 | 4.980 | 0.203 | 0.962 | -0.254 |
| 137 | 0.246 | 0.290 | 0.394 | 0.428 | 3.744 | 0.220 | 0.965 | -0.182 |
| 138 | 0.280 | 0.357 | 0.405 | 0.543 | 4.802 | 0.243 | 0.952 | -0.263 |
| 139 | 0.886 | 0.892 | 0.912 | 0.904 | 0.428 | 0.885 | 0.675 | -0.018 |
| 140 | 0.779 | 0.797 | 0.841 | 0.878 | 1.962 | 0.761 | 0.978 | -0.099 |
| 141 | 0.205 | 0.270 | 0.319 | 0.411 | 3.829 | 0.179 | 0.982 | -0.206 |
| 142 | 0.249 | 0.306 | 0.375 | 0.436 | 3.622 | 0.226 | 0.999 | -0.187 |
| 143 | 0.888 | 0.892 | 0.896 | 0.924 | 0.642 | 0.879 | 0.780 | -0.036 |
| 144 | 0.226 | 0.275 | 0.345 | 0.412 | 3.610 | 0.199 | 0.995 | -0.186 |
| 145 | 0.206 | 0.268 | 0.344 | 0.378 | 3.406 | 0.190 | 0.982 | -0.172 |
| 146 | 0.259 | 0.289 | 0.355 | 0.438 | 3.466 | 0.224 | 0.962 | -0.179 |
| 147 | 0.212 | 0.270 | 0.323 | 0.465 | 4.660 | 0.168 | 0.935 | -0.253 |
| 148 | 0.203 | 0.267 | 0.312 | 0.433 | 4.217 | 0.169 | 0.952 | -0.230 |
| 149 | 0.192 | 0.247 | 0.307 | 0.358 | 3.207 | 0.173 | 1.000 | -0.166 |
| 150 | 0.784 | 0.819 | 0.843 | 0.839 | 1.087 | 0.786 | 0.821 | -0.055 |
| 151 | 0.216 | 0.263 | 0.302 | 0.378 | 3.014 | 0.193 | 0.976 | -0.162 |
| 152 | 0.222 | 0.277 | 0.347 | 0.437 | 4.108 | 0.189 | 0.988 | -0.215 |
| 153 | 0.227 | 0.266 | 0.318 | 0.431 | 3.812 | 0.189 | 0.935 | -0.204 |
| 154 | 0.344 | 0.367 | 0.415 | 0.476 | 2.552 | 0.319 | 0.964 | -0.132 |

| | | | | | | | | |
|---|---|---|---|---|---|---|---|---|
| 155 | 0.247 | 0.288 | 0.353 | 0.409 | 3.168 | 0.223 | 0.994 | -0.162 |
| 156 | 0.188 | 0.254 | 0.297 | 0.388 | 3.690 | 0.164 | 0.978 | -0.200 |
| 157 | 0.571 | 0.586 | 0.637 | 0.676 | 2.106 | 0.550 | 0.966 | -0.105 |
| 158 | 0.188 | 0.274 | 0.331 | 0.370 | 3.464 | 0.180 | 0.970 | -0.182 |
| 159 | 0.205 | 0.241 | 0.335 | 0.444 | 4.663 | 0.157 | 0.960 | -0.239 |
| 160 | 0.193 | 0.257 | 0.302 | 0.345 | 2.878 | 0.182 | 0.989 | -0.152 |
| 161 | 0.720 | 0.747 | 0.767 | 0.761 | 0.823 | 0.722 | 0.781 | -0.041 |
| 162 | 0.515 | 0.525 | 0.593 | 0.673 | 3.118 | 0.477 | 0.919 | -0.158 |
| 163 | 0.240 | 0.275 | 0.368 | 0.447 | 4.108 | 0.201 | 0.974 | -0.207 |
| 164 | 0.289 | 0.362 | 0.392 | 0.526 | 4.248 | 0.256 | 0.927 | -0.237 |
| 165 | 0.225 | 0.278 | 0.322 | 0.378 | 2.889 | 0.208 | 0.997 | -0.153 |
| 166 | 0.215 | 0.253 | 0.305 | 0.351 | 2.645 | 0.196 | 0.997 | -0.136 |
| 167 | 0.608 | 0.639 | 0.666 | 0.675 | 1.311 | 0.605 | 0.953 | -0.067 |
| 168 | 0.238 | 0.306 | 0.391 | 0.496 | 4.936 | 0.200 | 0.991 | -0.258 |
| 169 | 0.250 | 0.326 | 0.402 | 0.467 | 4.178 | 0.228 | 0.999 | -0.217 |
| 170 | 0.224 | 0.252 | 0.324 | 0.452 | 4.343 | 0.174 | 0.919 | -0.228 |
| 171 | 0.192 | 0.223 | 0.299 | 0.355 | 3.251 | 0.163 | 0.980 | -0.163 |
| 172 | 0.217 | 0.244 | 0.301 | 0.436 | 4.100 | 0.168 | 0.892 | -0.219 |
| 173 | 0.192 | 0.252 | 0.343 | 0.404 | 4.182 | 0.164 | 0.995 | -0.212 |
| 174 | 0.224 | 0.265 | 0.325 | 0.394 | 3.276 | 0.197 | 0.988 | -0.170 |
| 175 | 0.271 | 0.343 | 0.342 | 0.461 | 3.257 | 0.250 | 0.863 | -0.190 |
| 176 | 0.369 | 0.410 | 0.437 | 0.535 | 3.011 | 0.341 | 0.918 | -0.166 |
| 177 | 0.184 | 0.228 | 0.335 | 0.357 | 3.607 | 0.161 | 0.947 | -0.173 |
| 178 | 0.202 | 0.253 | 0.302 | 0.425 | 4.121 | 0.164 | 0.940 | -0.223 |
| 179 | 0.161 | 0.224 | 0.294 | 0.352 | 3.696 | 0.139 | 0.999 | -0.191 |
| 180 | 0.165 | 0.228 | 0.282 | 0.348 | 3.464 | 0.145 | 0.998 | -0.183 |
| 181 | 0.166 | 0.200 | 0.276 | 0.331 | 3.285 | 0.138 | 0.984 | -0.165 |
| 182 | 0.236 | 0.261 | 0.325 | 0.352 | 2.373 | 0.218 | 0.971 | -0.116 |
| 183 | 0.172 | 0.235 | 0.270 | 0.336 | 3.024 | 0.156 | 0.985 | -0.164 |
| 184 | 0.209 | 0.246 | 0.330 | 0.363 | 3.144 | 0.186 | 0.972 | -0.154 |
| 185 | 0.269 | 0.321 | 0.413 | 0.478 | 4.136 | 0.238 | 0.992 | -0.209 |
| 186 | 0.294 | 0.366 | 0.441 | 0.511 | 4.172 | 0.269 | 1.000 | -0.217 |
| 187 | 0.236 | 0.260 | 0.358 | 0.464 | 4.498 | 0.186 | 0.943 | -0.228 |
| 188 | 0.228 | 0.273 | 0.333 | 0.372 | 2.830 | 0.211 | 0.995 | -0.144 |
| 189 | 0.238 | 0.277 | 0.326 | 0.382 | 2.764 | 0.217 | 0.994 | -0.144 |
| 190 | 0.209 | 0.230 | 0.339 | 0.388 | 3.722 | 0.172 | 0.947 | -0.179 |
| 191 | 0.302 | 0.356 | 0.447 | 0.561 | 4.989 | 0.257 | 0.977 | -0.259 |
| 192 | 0.228 | 0.281 | 0.347 | 0.420 | 3.690 | 0.201 | 0.995 | -0.192 |
| 193 | 0.190 | 0.221 | 0.299 | 0.410 | 4.242 | 0.144 | 0.944 | -0.220 |
| 194 | 0.196 | 0.229 | 0.292 | 0.318 | 2.470 | 0.180 | 0.978 | -0.122 |
| 195 | 0.254 | 0.298 | 0.383 | 0.431 | 3.545 | 0.228 | 0.987 | -0.177 |
| 196 | 0.210 | 0.288 | 0.414 | 0.450 | 4.871 | 0.185 | 0.966 | -0.240 |
| 197 | 0.668 | 0.723 | 0.736 | 0.754 | 1.554 | 0.671 | 0.887 | -0.086 |
| 198 | 0.603 | 0.567 | 0.659 | 0.670 | 1.697 | 0.570 | 0.621 | -0.067 |
| 199 | 0.286 | 0.323 | 0.332 | 0.459 | 3.024 | 0.253 | 0.813 | -0.173 |
| 200 | 0.212 | 0.271 | 0.309 | 0.389 | 3.266 | 0.191 | 0.979 | -0.177 |
| 201 | 0.193 | 0.234 | 0.353 | 0.412 | 4.468 | 0.155 | 0.971 | -0.219 |
| 202 | 0.188 | 0.236 | 0.311 | 0.369 | 3.554 | 0.162 | 0.995 | -0.181 |
| 203 | 0.182 | 0.217 | 0.311 | 0.432 | 4.852 | 0.130 | 0.950 | -0.250 |
| 204 | 0.188 | 0.226 | 0.278 | 0.361 | 3.280 | 0.158 | 0.968 | -0.173 |
| 205 | 0.199 | 0.245 | 0.299 | 0.373 | 3.309 | 0.173 | 0.987 | -0.174 |
| 206 | 0.199 | 0.223 | 0.304 | 0.364 | 3.315 | 0.166 | 0.966 | -0.165 |
| 207 | 0.735 | 0.762 | 0.789 | 0.807 | 1.397 | 0.729 | 0.993 | -0.072 |

| | | | | | | | | |
|---|---|---|---|---|---|---|---|---|
| 208 | 0.189 | 0.250 | 0.308 | 0.395 | 3.882 | 0.161 | 0.989 | -0.206 |
| 209 | 0.212 | 0.290 | 0.344 | 0.401 | 3.567 | 0.198 | 0.991 | -0.189 |
| 210 | 0.172 | 0.211 | 0.260 | 0.372 | 3.726 | 0.135 | 0.933 | -0.200 |
| 211 | 0.205 | 0.269 | 0.316 | 0.404 | 3.697 | 0.180 | 0.983 | -0.199 |
| 212 | 0.208 | 0.246 | 0.321 | 0.452 | 4.636 | 0.158 | 0.937 | -0.244 |
| 213 | 0.208 | 0.282 | 0.370 | 0.371 | 3.322 | 0.201 | 0.904 | -0.163 |
| 214 | 0.226 | 0.284 | 0.341 | 0.461 | 4.375 | 0.188 | 0.959 | -0.235 |
| 215 | 0.269 | 0.321 | 0.381 | 0.527 | 4.787 | 0.221 | 0.930 | -0.258 |
| 216 | 0.719 | 0.759 | 0.811 | 0.841 | 2.404 | 0.706 | 0.992 | -0.122 |
| 217 | 0.249 | 0.317 | 0.327 | 0.459 | 3.666 | 0.221 | 0.880 | -0.210 |
| 218 | 0.256 | 0.321 | 0.398 | 0.451 | 3.806 | 0.235 | 0.997 | -0.195 |
| 219 | 0.647 | 0.681 | 0.729 | 0.717 | 1.487 | 0.646 | 0.811 | -0.070 |
| 220 | 0.275 | 0.320 | 0.408 | 0.439 | 3.339 | 0.254 | 0.972 | -0.164 |
| 221 | 0.209 | 0.246 | 0.345 | 0.441 | 4.573 | 0.164 | 0.969 | -0.232 |
| 222 | 0.214 | 0.241 | 0.327 | 0.422 | 4.083 | 0.170 | 0.953 | -0.208 |
| 223 | 0.180 | 0.232 | 0.281 | 0.385 | 3.812 | 0.148 | 0.961 | -0.205 |
| 224 | 0.408 | 0.509 | 0.584 | 0.593 | 3.623 | 0.408 | 0.902 | -0.185 |
| 225 | 0.180 | 0.232 | 0.304 | 0.415 | 4.464 | 0.140 | 0.971 | -0.235 |
| 226 | 0.167 | 0.235 | 0.323 | 0.354 | 3.734 | 0.150 | 0.973 | -0.187 |
| 227 | 0.170 | 0.205 | 0.271 | 0.357 | 3.604 | 0.135 | 0.968 | -0.187 |
| 228 | 0.166 | 0.203 | 0.266 | 0.361 | 3.723 | 0.130 | 0.961 | -0.195 |
| 229 | 0.165 | 0.215 | 0.268 | 0.346 | 3.423 | 0.139 | 0.987 | -0.181 |
| 230 | 0.169 | 0.200 | 0.273 | 0.350 | 3.542 | 0.135 | 0.971 | -0.181 |
| 231 | 0.180 | 0.214 | 0.309 | 0.357 | 3.604 | 0.150 | 0.972 | -0.177 |
| 232 | 0.160 | 0.213 | 0.263 | 0.321 | 3.062 | 0.141 | 0.999 | -0.161 |
| 233 | 0.191 | 0.230 | 0.310 | 0.393 | 3.945 | 0.155 | 0.978 | -0.202 |
| 234 | 0.178 | 0.231 | 0.284 | 0.334 | 2.994 | 0.161 | 1.000 | -0.156 |
| 235 | 0.174 | 0.220 | 0.293 | 0.352 | 3.491 | 0.148 | 0.995 | -0.178 |
| 236 | 0.175 | 0.219 | 0.273 | 0.304 | 2.536 | 0.162 | 0.992 | -0.129 |
| 237 | 0.158 | 0.200 | 0.260 | 0.318 | 3.104 | 0.135 | 0.995 | -0.160 |
| 238 | 0.202 | 0.272 | 0.342 | 0.403 | 3.868 | 0.181 | 0.999 | -0.201 |
| 239 | 0.225 | 0.300 | 0.368 | 0.440 | 4.096 | 0.202 | 0.999 | -0.215 |
| 240 | 0.274 | 0.322 | 0.392 | 0.453 | 3.490 | 0.249 | 0.996 | -0.179 |
| 241 | 0.256 | 0.292 | 0.372 | 0.446 | 3.738 | 0.222 | 0.979 | -0.190 |
| 242 | 0.277 | 0.314 | 0.394 | 0.469 | 3.773 | 0.243 | 0.980 | -0.192 |
| 243 | 0.231 | 0.282 | 0.326 | 0.424 | 3.576 | 0.201 | 0.961 | -0.193 |
| 244 | 0.251 | 0.301 | 0.401 | 0.565 | 5.987 | 0.188 | 0.943 | -0.314 |
| 245 | 0.266 | 0.302 | 0.357 | 0.491 | 4.191 | 0.220 | 0.910 | -0.225 |
| 246 | 0.331 | 0.418 | 0.517 | 0.560 | 4.521 | 0.312 | 0.979 | -0.229 |
| 247 | 0.679 | 0.678 | 0.705 | 0.745 | 1.294 | 0.660 | 0.856 | -0.066 |
| 248 | 0.213 | 0.262 | 0.336 | 0.459 | 4.664 | 0.168 | 0.959 | -0.246 |
| 249 | 0.229 | 0.276 | 0.335 | 0.488 | 4.799 | 0.178 | 0.915 | -0.259 |
| 250 | 0.681 | 0.702 | 0.702 | 0.749 | 1.168 | 0.671 | 0.832 | -0.068 |
| 251 | 0.200 | 0.252 | 0.289 | 0.352 | 2.830 | 0.183 | 0.989 | -0.152 |
| 252 | 0.190 | 0.235 | 0.320 | 0.358 | 3.390 | 0.167 | 0.981 | -0.168 |
| 253 | 0.258 | 0.306 | 0.354 | 0.382 | 2.415 | 0.248 | 0.988 | -0.124 |
| 254 | 0.320 | 0.360 | 0.443 | 0.489 | 3.396 | 0.294 | 0.984 | -0.169 |
| 255 | 0.248 | 0.311 | 0.367 | 0.484 | 4.386 | 0.212 | 0.966 | -0.236 |
| 256 | 0.283 | 0.352 | 0.342 | 0.423 | 2.345 | 0.275 | 0.842 | -0.140 |
| 257 | 0.250 | 0.304 | 0.354 | 0.433 | 3.440 | 0.225 | 0.987 | -0.183 |
| 258 | 0.699 | 0.713 | 0.757 | 0.794 | 1.893 | 0.680 | 0.967 | -0.095 |
| 259 | 0.322 | 0.330 | 0.439 | 0.543 | 4.443 | 0.266 | 0.915 | -0.221 |
| 260 | 0.356 | 0.343 | 0.439 | 0.547 | 3.851 | 0.298 | 0.847 | -0.191 |

| | | | | | | | | |
|---|---|---|---|---|---|---|---|---|
| 261 | 0.246 | 0.280 | 0.344 | 0.390 | 2.853 | 0.224 | 0.990 | -0.144 |
| 262 | 0.287 | 0.324 | 0.373 | 0.495 | 3.863 | 0.246 | 0.917 | -0.208 |
| 263 | 0.255 | 0.302 | 0.349 | 0.398 | 2.735 | 0.238 | 1.000 | -0.143 |
| 264 | 0.305 | 0.325 | 0.410 | 0.464 | 3.236 | 0.272 | 0.958 | -0.159 |
| 265 | 0.537 | 0.590 | 0.597 | 0.638 | 1.776 | 0.534 | 0.925 | -0.101 |
| 266 | 0.617 | 0.692 | 0.728 | 0.795 | 3.271 | 0.603 | 0.981 | -0.178 |
| 267 | 0.401 | 0.393 | 0.411 | 0.521 | 2.168 | 0.362 | 0.656 | -0.120 |
| 268 | 0.235 | 0.263 | 0.287 | 0.366 | 2.392 | 0.211 | 0.910 | -0.131 |
| 269 | 0.282 | 0.314 | 0.421 | 0.499 | 4.363 | 0.239 | 0.967 | -0.217 |
| 270 | 0.226 | 0.266 | 0.332 | 0.381 | 3.054 | 0.204 | 0.994 | -0.155 |
| 271 | 0.242 | 0.283 | 0.319 | 0.398 | 2.893 | 0.218 | 0.962 | -0.156 |
| 272 | 0.248 | 0.282 | 0.354 | 0.415 | 3.296 | 0.219 | 0.984 | -0.167 |
| 273 | 0.206 | 0.240 | 0.292 | 0.386 | 3.400 | 0.172 | 0.949 | -0.180 |
| 274 | 0.221 | 0.253 | 0.312 | 0.341 | 2.411 | 0.205 | 0.985 | -0.120 |
| 275 | 0.242 | 0.271 | 0.320 | 0.395 | 2.919 | 0.214 | 0.960 | -0.153 |
| 276 | 0.803 | 0.800 | 0.851 | 0.870 | 1.454 | 0.784 | 0.873 | -0.067 |
| 277 | 0.272 | 0.336 | 0.419 | 0.434 | 3.275 | 0.260 | 0.946 | -0.162 |
| 278 | 0.789 | 0.832 | 0.874 | 0.910 | 2.328 | 0.777 | 0.999 | -0.121 |
| 279 | 0.826 | 0.857 | 0.883 | 0.906 | 1.528 | 0.819 | 0.995 | -0.080 |
| **Average** | 0.290 | 0.338 | 0.403 | 0.471 | 3.483 | 0.264 | 0.959 | -0.180 |

**Table S13. (1-$S^2$) values obtained from increasingly X-ray damaged proteinase K room temperature (277 K) datasets.** (1-$S^2$) values were calculated from proteinase K multi-conformer models (**Table S9**) and as described in Materials and Methods. Slopes, correlation coefficients, and intercepts were obtained via linear regression.

| | $(1-S^2)$ | | | Slope $(1-S^2)$/MGy | Intercept (zero-dose $1-S^2$) | $R^2$ | $\Delta(1-S^2)$ |
|---|---|---|---|---|---|---|---|
| **Residue #** | **Dataset 1** | **Dataset 2** | **Dataset 3** | | | | **Datasets 1-3** |
| **Dose (MGy)** | 0.017 | 0.069 | 0.121 | | | | |
| 1 | 0.413 | 0.471 | 0.629 | 2.077 | 0.361 | 0.933 | -0.216 |
| 2 | 0.399 | 0.434 | 0.557 | 1.519 | 0.359 | 0.906 | -0.158 |
| 3 | 0.361 | 0.377 | 0.530 | 1.625 | 0.311 | 0.820 | -0.169 |
| 4 | 0.405 | 0.426 | 0.458 | 0.510 | 0.395 | 0.986 | -0.053 |
| 5 | 0.408 | 0.494 | 0.615 | 1.990 | 0.368 | 0.991 | -0.207 |
| 6 | 0.350 | 0.435 | 0.503 | 1.471 | 0.328 | 0.996 | -0.153 |
| 7 | 0.364 | 0.425 | 0.561 | 1.894 | 0.319 | 0.954 | -0.197 |
| 8 | 0.345 | 0.393 | 0.457 | 1.077 | 0.324 | 0.993 | -0.112 |
| 9 | 0.366 | 0.424 | 0.541 | 1.683 | 0.328 | 0.963 | -0.175 |
| 10 | 0.379 | 0.437 | 0.589 | 2.019 | 0.329 | 0.937 | -0.210 |
| 11 | 0.392 | 0.436 | 0.563 | 1.644 | 0.350 | 0.927 | -0.171 |
| 12 | 0.384 | 0.424 | 0.512 | 1.231 | 0.355 | 0.955 | -0.128 |
| 13 | 0.404 | 0.498 | 0.601 | 1.894 | 0.370 | 0.999 | -0.197 |
| 14 | 0.667 | 0.701 | 0.748 | 0.779 | 0.652 | 0.991 | -0.081 |
| 15 | 0.465 | 0.555 | 0.677 | 2.038 | 0.425 | 0.992 | -0.212 |
| 16 | 0.493 | 0.505 | 0.620 | 1.221 | 0.455 | 0.820 | -0.127 |
| 17 | 0.514 | 0.549 | 0.665 | 1.452 | 0.476 | 0.912 | -0.151 |
| 18 | 0.769 | 0.766 | 0.816 | 0.452 | 0.752 | 0.702 | -0.047 |
| 19 | 0.592 | 0.649 | 0.750 | 1.519 | 0.559 | 0.975 | -0.158 |
| 20 | 0.437 | 0.501 | 0.538 | 0.971 | 0.425 | 0.977 | -0.101 |
| 21 | 0.565 | 0.591 | 0.697 | 1.269 | 0.530 | 0.891 | -0.132 |
| 22 | 0.402 | 0.444 | 0.573 | 1.644 | 0.360 | 0.921 | -0.171 |
| 23 | 0.344 | 0.405 | 0.534 | 1.827 | 0.302 | 0.959 | -0.190 |
| 24 | 0.399 | 0.476 | 0.615 | 2.077 | 0.353 | 0.973 | -0.216 |
| 25 | 0.388 | 0.417 | 0.547 | 1.529 | 0.345 | 0.881 | -0.159 |
| 26 | 0.327 | 0.357 | 0.495 | 1.615 | 0.282 | 0.879 | -0.168 |
| 27 | 0.335 | 0.373 | 0.436 | 0.971 | 0.314 | 0.980 | -0.101 |
| 28 | 0.333 | 0.324 | 0.320 | -0.125 | 0.334 | 0.953 | 0.013 |
| 29 | 0.329 | 0.372 | 0.514 | 1.779 | 0.282 | 0.913 | -0.185 |
| 30 | 0.296 | 0.328 | 0.436 | 1.346 | 0.260 | 0.911 | -0.140 |
| 31 | 0.292 | 0.352 | 0.428 | 1.308 | 0.267 | 0.995 | -0.136 |
| 32 | 0.328 | 0.348 | 0.473 | 1.394 | 0.287 | 0.851 | -0.145 |
| 33 | 0.312 | 0.353 | 0.460 | 1.423 | 0.277 | 0.938 | -0.148 |
| 34 | 0.327 | 0.364 | 0.488 | 1.548 | 0.286 | 0.911 | -0.161 |
| 35 | 0.344 | 0.389 | 0.459 | 1.106 | 0.321 | 0.984 | -0.115 |
| 36 | 0.322 | 0.359 | 0.504 | 1.750 | 0.274 | 0.895 | -0.182 |
| 37 | 0.481 | 0.541 | 0.685 | 1.962 | 0.434 | 0.947 | -0.204 |
| 38 | 0.350 | 0.413 | 0.529 | 1.721 | 0.312 | 0.972 | -0.179 |
| 39 | 0.381 | 0.395 | 0.482 | 0.971 | 0.352 | 0.852 | -0.101 |
| 40 | 0.309 | 0.378 | 0.488 | 1.721 | 0.273 | 0.983 | -0.179 |
| 41 | 0.295 | 0.334 | 0.432 | 1.317 | 0.263 | 0.942 | -0.137 |
| 42 | 0.297 | 0.329 | 0.432 | 1.298 | 0.263 | 0.916 | -0.135 |
| 43 | 0.313 | 0.350 | 0.435 | 1.173 | 0.285 | 0.951 | -0.122 |
| 44 | 0.420 | 0.477 | 0.616 | 1.885 | 0.374 | 0.945 | -0.196 |
| 45 | 0.484 | 0.559 | 0.711 | 2.183 | 0.434 | 0.963 | -0.227 |
| 46 | 0.598 | 0.611 | 0.720 | 1.173 | 0.562 | 0.829 | -0.122 |
| 47 | 0.776 | 0.867 | 0.942 | 1.596 | 0.752 | 0.997 | -0.166 |
| 48 | 0.462 | 0.539 | 0.630 | 1.615 | 0.432 | 0.998 | -0.168 |
| 49 | 0.422 | 0.514 | 0.623 | 1.933 | 0.386 | 0.998 | -0.201 |
| 50 | 0.377 | 0.432 | 0.602 | 2.163 | 0.321 | 0.920 | -0.225 |

| | | | | | | | |
|---|---|---|---|---|---|---|---|
| 51 | 0.367 | 0.397 | 0.432 | 0.625 | 0.356 | 0.998 | -0.065 |
| 52 | 0.319 | 0.356 | 0.485 | 1.596 | 0.277 | 0.907 | -0.166 |
| 53 | 0.296 | 0.341 | 0.416 | 1.154 | 0.271 | 0.980 | -0.120 |
| 54 | 0.286 | 0.300 | 0.387 | 0.971 | 0.257 | 0.852 | -0.101 |
| 55 | 0.450 | 0.532 | 0.629 | 1.721 | 0.418 | 0.998 | -0.179 |
| 56 | 0.322 | 0.356 | 0.579 | 2.471 | 0.248 | 0.847 | -0.257 |
| 57 | 0.281 | 0.349 | 0.486 | 1.971 | 0.236 | 0.964 | -0.205 |
| 58 | 0.333 | 0.356 | 0.446 | 1.087 | 0.303 | 0.895 | -0.113 |
| 59 | 0.690 | 0.741 | 0.751 | 0.587 | 0.687 | 0.869 | -0.061 |
| 60 | 0.339 | 0.362 | 0.484 | 1.394 | 0.299 | 0.866 | -0.145 |
| 61 | 0.512 | 0.573 | 0.727 | 2.067 | 0.461 | 0.941 | -0.215 |
| 62 | 0.482 | 0.559 | 0.741 | 2.490 | 0.422 | 0.948 | -0.259 |
| 63 | 0.347 | 0.406 | 0.524 | 1.702 | 0.308 | 0.964 | -0.177 |
| 64 | 0.311 | 0.366 | 0.462 | 1.452 | 0.279 | 0.976 | -0.151 |
| 65 | 0.408 | 0.437 | 0.610 | 1.942 | 0.351 | 0.855 | -0.202 |
| 66 | 0.334 | 0.386 | 0.546 | 2.038 | 0.281 | 0.920 | -0.212 |
| 67 | 0.360 | 0.429 | 0.609 | 2.394 | 0.301 | 0.938 | -0.249 |
| 68 | 0.386 | 0.408 | 0.532 | 1.404 | 0.345 | 0.860 | -0.146 |
| 69 | 0.359 | 0.428 | 0.549 | 1.827 | 0.319 | 0.976 | -0.190 |
| 70 | 0.607 | 0.665 | 0.817 | 2.019 | 0.557 | 0.937 | -0.210 |
| 71 | 0.543 | 0.673 | 0.960 | 4.010 | 0.449 | 0.955 | -0.417 |
| 72 | 0.645 | 0.680 | 0.916 | 2.606 | 0.567 | 0.845 | -0.271 |
| 73 | 0.634 | 0.796 | 1.010 | 3.615 | 0.564 | 0.994 | -0.376 |
| 74 | 0.451 | 0.502 | 0.615 | 1.577 | 0.414 | 0.955 | -0.164 |
| 75 | 0.448 | 0.490 | 0.626 | 1.712 | 0.403 | 0.915 | -0.178 |
| 76 | 0.388 | 0.461 | 0.581 | 1.856 | 0.349 | 0.981 | -0.193 |
| 77 | 0.834 | 0.863 | 0.932 | 0.942 | 0.811 | 0.947 | -0.098 |
| 78 | 0.659 | 0.714 | 0.802 | 1.375 | 0.630 | 0.983 | -0.143 |
| 79 | 0.592 | 0.650 | 0.725 | 1.279 | 0.567 | 0.995 | -0.133 |
| 80 | 0.304 | 0.339 | 0.423 | 1.144 | 0.276 | 0.947 | -0.119 |
| 81 | 0.699 | 0.724 | 0.797 | 0.942 | 0.675 | 0.926 | -0.098 |
| 82 | 0.465 | 0.509 | 0.616 | 1.452 | 0.430 | 0.945 | -0.151 |
| 83 | 0.377 | 0.396 | 0.525 | 1.423 | 0.334 | 0.844 | -0.148 |
| 84 | 0.388 | 0.414 | 0.498 | 1.058 | 0.360 | 0.915 | -0.110 |
| 85 | 0.478 | 0.582 | 0.619 | 1.356 | 0.466 | 0.930 | -0.141 |
| 86 | 0.818 | 0.854 | 0.925 | 1.029 | 0.795 | 0.966 | -0.107 |
| 87 | 0.459 | 0.494 | 0.624 | 1.587 | 0.416 | 0.900 | -0.165 |
| 88 | 0.447 | 0.457 | 0.523 | 0.731 | 0.425 | 0.847 | -0.076 |
| 89 | 0.416 | 0.467 | 0.566 | 1.442 | 0.383 | 0.967 | -0.150 |
| 90 | 0.440 | 0.542 | 0.656 | 2.077 | 0.403 | 0.999 | -0.216 |
| 91 | 0.393 | 0.411 | 0.556 | 1.567 | 0.345 | 0.832 | -0.163 |
| 92 | 0.424 | 0.460 | 0.590 | 1.596 | 0.381 | 0.903 | -0.166 |
| 93 | 0.606 | 0.624 | 0.711 | 1.010 | 0.577 | 0.874 | -0.105 |
| 94 | 0.384 | 0.475 | 0.562 | 1.712 | 0.356 | 1.000 | -0.178 |
| 95 | 0.384 | 0.423 | 0.568 | 1.769 | 0.336 | 0.900 | -0.184 |
| 96 | 0.390 | 0.472 | 0.588 | 1.904 | 0.352 | 0.990 | -0.198 |
| 97 | 0.862 | 0.870 | 0.904 | 0.404 | 0.851 | 0.887 | -0.042 |
| 98 | 0.390 | 0.429 | 0.596 | 1.981 | 0.335 | 0.886 | -0.206 |
| 99 | 0.362 | 0.439 | 0.489 | 1.221 | 0.346 | 0.985 | -0.127 |
| 100 | 0.532 | 0.625 | 0.857 | 3.125 | 0.456 | 0.943 | -0.325 |
| 101 | 0.716 | 0.787 | 1.020 | 2.923 | 0.639 | 0.914 | -0.304 |
| 102 | 0.581 | 0.649 | 0.787 | 1.981 | 0.536 | 0.963 | -0.206 |
| 103 | 0.709 | 0.774 | 0.870 | 1.548 | 0.678 | 0.988 | -0.161 |

| | | | | | | | |
|---|---|---|---|---|---|---|---|
| 104 | 0.390 | 0.400 | 0.512 | 1.173 | 0.353 | 0.811 | -0.122 |
| 105 | 0.326 | 0.377 | 0.529 | 1.952 | 0.276 | 0.924 | -0.203 |
| 106 | 0.357 | 0.392 | 0.561 | 1.962 | 0.301 | 0.874 | -0.204 |
| 107 | 0.381 | 0.444 | 0.554 | 1.663 | 0.345 | 0.976 | -0.173 |
| 108 | 0.320 | 0.368 | 0.477 | 1.510 | 0.284 | 0.952 | -0.157 |
| 109 | 0.810 | 0.821 | 0.847 | 0.356 | 0.801 | 0.948 | -0.037 |
| 110 | 0.429 | 0.477 | 0.565 | 1.308 | 0.400 | 0.972 | -0.136 |
| 111 | 0.324 | 0.336 | 0.412 | 0.846 | 0.299 | 0.850 | -0.088 |
| 112 | 0.725 | 0.746 | 0.800 | 0.721 | 0.707 | 0.939 | -0.075 |
| 113 | 0.379 | 0.418 | 0.545 | 1.596 | 0.337 | 0.914 | -0.166 |
| 114 | 0.385 | 0.441 | 0.577 | 1.846 | 0.340 | 0.945 | -0.192 |
| 115 | 0.331 | 0.365 | 0.438 | 1.029 | 0.307 | 0.958 | -0.107 |
| 116 | 0.365 | 0.425 | 0.502 | 1.317 | 0.340 | 0.995 | -0.137 |
| 117 | 0.459 | 0.501 | 0.602 | 1.375 | 0.426 | 0.946 | -0.143 |
| 118 | 0.392 | 0.465 | 0.600 | 2.000 | 0.348 | 0.971 | -0.208 |
| 119 | 0.477 | 0.578 | 0.730 | 2.433 | 0.427 | 0.987 | -0.253 |
| 120 | 0.436 | 0.489 | 0.527 | 0.875 | 0.424 | 0.991 | -0.091 |
| 121 | 0.809 | 0.837 | 0.892 | 0.798 | 0.791 | 0.966 | -0.083 |
| 122 | 0.472 | 0.508 | 0.712 | 2.308 | 0.405 | 0.860 | -0.240 |
| 123 | 0.396 | 0.419 | 0.582 | 1.788 | 0.342 | 0.841 | -0.186 |
| 124 | 0.401 | 0.473 | 0.597 | 1.885 | 0.360 | 0.977 | -0.196 |
| 125 | 0.911 | 0.938 | 0.987 | 0.731 | 0.895 | 0.973 | -0.076 |
| 126 | 0.444 | 0.533 | 0.651 | 1.990 | 0.405 | 0.994 | -0.207 |
| 127 | 0.373 | 0.442 | 0.602 | 2.202 | 0.320 | 0.950 | -0.229 |
| 128 | 0.865 | 0.893 | 0.937 | 0.692 | 0.851 | 0.984 | -0.072 |
| 129 | 0.703 | 0.790 | 0.942 | 2.298 | 0.653 | 0.976 | -0.239 |
| **Average** | 0.451 | 0.500 | 0.614 | 1.564 | 0.414 | 0.935 | -0.163 |

**Table S14. (1-$S^2$) values obtained from increasingly X-ray damaged lysozyme room temperature (277 K) datasets.** (1-$S^2$) values were calculated from lysozyme multi-conformer models (**Table S10**) and as described in Materials and Methods. Slopes, correlation coefficients, and intercepts were obtained via linear regression.

| Thaumatin 277 K diffraction data collection statistics | | | | |
|---|---|---|---|---|
| Dataset | 1 | 2 (merged) | 3 (merged) | 4 (merged) |
| Wavelength (Å) | 0.88557 | | | |
| Resolution range (Å) | 38-26-1.22 (1.24-1.22) | 38-27-1.22 (1.24-1.22) | 38-27-1.22 (1.24-1.22) | 38-27-1.22 (1.24-1.22) |
| DWD (MGy) | 0.016 | 0.114 | 0.212 | 0.310 |
| Space group | $P4_12_12$ | | | |
| Unit cell | 58.81 58.81 151.17 90.00 90.00 90.00 | 58.82 58.82 151.15 90.00 90.00 90.00 | 58.82 58.82 151.16 90.00 90.00 90.00 | 58.82 58.82 151.17 90.00 90.00 90.00 |
| Unit cell volume ($Å^3$) | 522839.0 | 522947.6 | 522982.2 | 523016.8 |
| Total reflections | 686302 (30123) | 4818222 (209908) | 8948567 (403085) | 13072934 (568854) |
| Multiplicity | 8.6 (7.9) | 60.0 (54.3) | 112.0 (100.5) | 163.6 (146.5) |
| Mosaicity (°) | 0.12 | 0.12 | 0.12 | 0.13 |
| Completeness (%) | 99.9 (98.6) | 100 (99.5) | 100 (99.5) | 100 (99.8) |
| Mean I/sigma(I) | 8.5 (0.5) | 21.5 (1.1) | 26.6 (1.2) | 28.5 (1.1) |
| Wilson B-factor | 20.9 | 21.8 | 22.6 | 23.0 |
| R-merge | 0.110 (4.479) | 0.135 (6.955) | 0.206 (16.249) | 0.459 (58.722) |
| R-pim | 0.040 (1.678) | 0.018 (0.938) | 0.019 (1.605) | 0.036 (4.793) |
| $CC_{1/2}$ | 0.999 (0.304) | 1.000 (0.65.7) | 1.000 (0.666) | 1.000 (0.587) |
| Isa | 17.3 | 16.1 | 15.5 | 15.2 |
| **Thaumatin refinement statistics multi-conformer models** | | | | |
| Dataset | 1 | 2 (merged) | 3 (merged) | 4 (merged) |
| PDB code | 7LJW | 7LNB | 7LNC | 7LND |
| Resolution range (Å) | 38.26 -1.22 (1.26 -1.22) | 37.79 - 1.22 (1.26 - 1.22) | 38.27 - 1.22 (1.26 - 1.22) | 36.41 - 1.22 (1.26 - 1.22) |
| Reflections used in refinement | 78731 (6867) | 79672 (7713) | 79668 (7721) | 79570 (7758) |
| Rwork | 0.142 (0.332) | 0.133 (0.274) | 0.136 (0.269) | 0.136 (0.300) |
| Rfree | 0.156 (0.340) | 0.145 (0.284) | 0.147 (0.272) | 0.148 (0.309) |
| No. on non-hydrogen atoms | 3771 | 3771 | 3771 | 3771 |
| Protein | 3578 | 3578 | 3578 | 3578 |
| Ligand/ion | 20 | 20 | 20 | 20 |
| Water | 151 | 151 | 151 | 151 |
| RMS (bonds) | 0.009 | 0.009 | 0.006 | 0.009 |
| RMS (angles) | 1.06 | 1.06 | 0.91 | 1.07 |
| Average B-factor | 18.18 | 18.14 | 18.48 | 20.89 |
| Protein | 17.48 | 17.45 | 17.79 | 20.19 |
| Ligand/ion | 16.13 | 15.56 | 15.63 | 17.97 |
| Water | 32.85 | 32.81 | 33.14 | 35.52 |
| Ramachandran (%) | | | | |
| Favored | 96.6 | 96.1 | 96.1 | 96.1 |
| Allowed | 3.4 | 3.9 | 3.9 | 3.9 |
| Outliers | 0 | 0 | 0 | 0 |

**Table S15**. **Room temperature (277 K) diffraction statistics and multi-conformer refinement statistics for increasingly damaged thaumatin datasets.** Diffraction statistics are reported for increasingly damaged datasets in which an increasing amount of increasingly damaged data have been merged together –i.e. dataset 1 is of 120°total rotation, dataset 2 is of 480° total rotation and so forth. Values in parenthesis are for the highest resolution shells. All diffraction statistics were obtained from Aimless (41), with the exception of $CC_{1/2}$, which was obtained from XSCALE (39). Average diffraction weighted doses (DWD) were estimated using the program RADDOSE 3D (35, 36). Multi-conformer models were obtained as described in Materials and Methods. Refinement statistics were obtained from phenix (*phenix.table_one*) using the final refined models and reflections files. Diffraction and refinement statistics for dataset 1 and the respective multi-conformer model are from **Table S1** and **Table S8**, respectively.

| Proteinase K 277 K diffraction data collection statistics | | | | |
|---|---|---|---|---|
| Dataset | 1 | 2 (merged) | 3 (merged) | 4 (merged) |
| Wavelength (Å) | 0.88557 | | | |
| Resolution range (Å) | 34.91-1.02 (1.04-1.02) | 34.89-1.02 (1.04-1.02) | 34.88-1.02 (1.04-1.02) | 34.89-1.02 (1.04-1.02) |
| DWD (MGy) | 0.006 | 0.023 | 0.041 | 0.058 |
| Space group | $P4_32_12$ | | | |
| Unit cell | 67.82 67.82 101.86 90.00 90.00 90.00 | 67.78 67.78 101.76 90.00 90.00 90.00 | 67.77 67.77 101.76 90.00 90.00 90.00 | 67.78 67.78 101.78 90.00 90.00 90.00 |
| Unit cell volume ($Å^3$) | 468510.4 | 467498.5 | 467360.6 | 467590.4 |
| Total reflections | 822825 (28269) | 3282233 (111354) | 5727309 (192083) | 8153634 (270403) |
| Multiplicity | 6.8 (4.9) | 27.2 (19.0) | 47.5 (32.7) | 67.6 (46.1) |
| Mosaicity (°) | 0.08 | 0.09 | 0.13 | 0.18 |
| Completeness (%) | 99.9 (98.3) | 100 (100) | 100 (100) | 100 (100) |
| Mean I/sigma(I) | 8.5 (0.9) | 14.4 (1.2) | 16.7 (1.2) | 17.7 (1.2) |
| Wilson B-factor | 12.2 | 13.0 | 13.7 | 14.2 |
| R-merge | 0.098 (1.720) | 0.133 (3.410) | 0.248 (12.902) | 0.625 (55.751) |
| R-pim | 0.038 (0.862) | 0.025 (0.792) | 0.037 (2.232) | 0.082 (8.158) |
| $CC_{1/2}$ | 0.999 (0.359) | 1.000 (0.544) | 1.000 (0.477) | 1.000 (0.442) |
| Isa | 19.3 | 14.8 | 14.9 | 15.2 |
| **Proteinase K refinement statistics multi-conformer models** | | | | |
| Dataset | 1 | 2 (merged) | 3 (merged) | 4 (merged) |
| PDB code | 7LPU | 7LQA | 7LQB | 7LQC |
| Resolution range (Å) | 33.91 - 1.02 (1.06 - 1.02) | 33.89 - 1.02 (1.06 - 1.02) | 33.89 - 1.02 (1.06 - 1.02) | 33.92 - 1.021 (1.06 - 1.021) |
| Reflections used in refinement | 120585 (11691) | 120536 (11840) | 120501 (11846) | 120548 (11846) |
| Rwork | 0.119 (0.287) | 0.110 (0.241) | 0.111 (0.243) | 0.111 (0.277) |
| Rfree | 0.138 (0.301) | 0.127 (0.257) | 0.128 (0.261) | 0.128 (0.285) |
| No. on non-hydrogen atoms | 6724 | 6724 | 6724 | 6724 |
| Protein | 6345 | 6345 | 6345 | 6345 |
| Ligand/ion | 22 | 22 | 22 | 22 |
| Water | 282 | 282 | 281 | 281 |
| RMS (bonds) | 0.007 | 0.007 | 0.007 | 0.007 |
| RMS (angles) | 0.95 | 0.96 | 0.96 | 0.96 |
| Average B-factor | 8.97 | 8.9 | 8.83 | 10.71 |
| Protein | 8.1 | 8.04 | 7.96 | 9.83 |
| Ligand/ion | 22.93 | 22.43 | 22.12 | 24.26 |
| Water | 23.57 | 23.52 | 23.45 | 25.5 |
| Ramachandran (%) | | | | |
| Favored | 96.03 | 95.67 | 95.67 | 96.75 |
| Allowed | 3.97 | 4.33 | 4.33 | 3.25 |
| Outliers | 0 | 0 | 0 | 0 |

**Table S16**. **Room temperature (277K) diffraction statistics and multi-conformer refinement statistics for increasingly damaged proteinase K datasets.** Diffraction statistics are reported for increasingly damaged datasets in which an increasing amount of increasingly damaged data have been merged together –i.e. dataset 1 is of 120°total rotation, dataset 2 is of 480° total rotation and so forth. Values in parenthesis are for the highest resolution shells. All diffraction statistics were obtained from Aimless (41), with the exception of $CC_{1/2}$, which was obtained from XSCALE (39). Average diffraction weighted doses (DWD) were estimated using the program RADDOSE 3D (35, 36). Multi-conformer models were obtained as described in Materials and Methods. Refinement statistics were obtained from phenix (*phenix.table_one*) using the final refined models and reflections files. Diffraction and refinement statistics for dataset 1 and the respective multi-conformer model are from **Table S2** and **Table S9**, respectively.

| Lysozyme 277 K diffraction data collection statistics | | | |
|---|---|---|---|
| Dataset | 1 | 2 (merged) | 3 (merged) |
| Wavelength (Å) | 0.88557 | | |
| Resolution range (Å) | 38.63-1.13 (1.15-1.13) | 38.63-1.13 (1.15-1.13) | 38.60-1.14 (1.16-1.14) |
| DWD (MGy) | 0.017 | 0.069 | 0.121 |
| Space group | $P4_32_12$ | | |
| Unit cell | 77.26 77.26 37.31 90.00 90.00 90.00 | 77.25 77.25 37.27 90.00 90.00 90.00 | 77.20 77.20 37.28 90.00 90.00 90.00 |
| Unit cell volume (Å$^3$) | 222707.4 | 222411.0 | 222182.8 |
| Total reflections | 365238 (17155) | 1455624 (67426) | 2468785 (105094) |
| Multiplicity | 8.5 (8.4) | 34.0 (32.7) | 59.2 (51.7) |
| Mosaicity (°) | 0.14 | 0.2 | 0.26 |
| Completeness (%) | 99.9 (99.1) | 100.0 (99.9) | 100.0 (100.0) |
| Mean I/sigma(I) | 12.4 (0.9) | 20.1 (1.1) | 19.3 (1.1) |
| Wilson B-factor | 18.9 | 20.1 | 21.0 |
| R-merge | 0.072 (2.720) | 0.119 (7.926) | 0.411 (56.454) |
| R-pim | 0.026 (0.989) | 0.021 (1.382) | 0.054 (7.853) |
| $CC_{1/2}$ | 0.999 (0.326) | 1.000 (0.419) | 1.000 (0.388) |
| Isa | 20.1 | 19.9 | 17.7 |
| **Lysozyme refinement statistics multi-conformer models** | | | |
| Dataset | 1 | 2 (merged) | 3 (merged) |
| PDB code | 7LN9 | 7LP6 | 7LPL |
| Resolution range (Å) | 33.6 - 1.13 (1.171 - 1.13) | 38.63 - 1.13 (1.17 - 1.13) | 33.56 - 1.14 (1.18 - 1.14) |
| Reflections used in refinement | 42596 (4064) | 42724 (4176) | 41595 (4012) |
| Rwork | 0.136 (0.292) | 0.136 (0.269) | 0.137 (0.304) |
| Rfree | 0.156 (0.308) | 0.156 (0.273) | 0.158 (0.301) |
| No. on non-hydrogen atoms | 3065 | 3065 | 3065 |
| Protein | 2925 | 2925 | 2925 |
| Ligand/ion | 4 | 4 | 4 |
| Water | 113 | 113 | 113 |
| RMS (bonds) | 0.006 | 0.007 | 0.006 |
| RMS (angles) | 0.84 | 0.87 | 0.85 |
| Average B-factor | 15.8 | 15.73 | 17.33 |
| Protein | 15.2 | 15.14 | 16.72 |
| Ligand/ion | 22.25 | 22.37 | 24.02 |
| Water | 28.48 | 28.31 | 30.12 |
| Ramachandran (%) | | | |
| Favored | 96.06 | 96.06 | 95.28 |
| Allowed | 3.94 | 3.94 | 4.72 |
| Outliers | 0 | 0 | 0 |

**Table S17**. **Room temperature (277K) diffraction statistics and multi-conformer refinement statistics for increasingly damaged lysozyme datasets.** Diffraction statistics are reported for increasingly damaged datasets in which an increasing amount of increasingly damaged data have been merged together –i.e. dataset 1 is of 120° total rotation, dataset 2 is of 480° total rotation and so forth. Values in parenthesis are for the highest resolution shells. All diffraction statistics were obtained from Aimless (41), with the exception of $CC_{1/2}$, which was obtained from XSCALE (39). Average diffraction weighted doses (DWD) were estimated using the program RADDOSE 3D (35, 36). Multi-conformer models were obtained as described in Materials and Methods. Refinement statistics were obtained from phenix (*phenix.table_one*) using the final refined models and reflections files. Diffraction and refinement statistics for dataset 1 and the respective multi-conformer model are from **Table S3** and **Table S10**, respectively.

| | $(1-S^2)$ | | | |
|---|---|---|---|---|
| **Residue #** | **Dataset 1** | **Dataset 2 (merged)** | **Dataset 3 (merged)** | **Dataset 4 (merged)** |
| **Dose (MGy)** | 0.016 | 0.114 | 0.212 | 0.31 |
| 1 | 0.557 | 0.576 | 0.589 | 0.655 |
| 2 | 0.823 | 0.817 | 0.817 | 0.849 |
| 3 | 0.403 | 0.405 | 0.407 | 0.468 |
| 4 | 0.424 | 0.422 | 0.422 | 0.488 |
| 5 | 0.346 | 0.33 | 0.334 | 0.391 |
| 6 | 0.335 | 0.328 | 0.331 | 0.401 |
| 7 | 0.320 | 0.329 | 0.325 | 0.395 |
| 8 | 0.350 | 0.335 | 0.337 | 0.395 |
| 9 | 0.320 | 0.312 | 0.313 | 0.373 |
| 10 | 0.459 | 0.474 | 0.487 | 0.549 |
| 11 | 0.404 | 0.418 | 0.429 | 0.494 |
| 12 | 0.417 | 0.41 | 0.408 | 0.466 |
| 13 | 0.408 | 0.374 | 0.379 | 0.442 |
| 14 | 0.355 | 0.349 | 0.355 | 0.412 |
| 15 | 0.405 | 0.401 | 0.398 | 0.469 |
| 16 | 0.396 | 0.393 | 0.395 | 0.468 |
| 17 | 0.451 | 0.428 | 0.438 | 0.503 |
| 18 | 0.501 | 0.483 | 0.483 | 0.555 |
| 19 | 0.584 | 0.582 | 0.6 | 0.665 |
| 20 | 0.590 | 0.625 | 0.623 | 0.672 |
| 21 | 0.718 | 0.696 | 0.714 | 0.788 |
| 22 | 0.553 | 0.58 | 0.594 | 0.652 |
| 23 | 0.565 | 0.577 | 0.58 | 0.634 |
| 24 | 0.522 | 0.51 | 0.517 | 0.594 |
| 25 | 0.572 | 0.58 | 0.586 | 0.646 |
| 26 | 0.651 | 0.637 | 0.657 | 0.717 |
| 27 | 0.483 | 0.489 | 0.497 | 0.572 |
| 28 | 0.374 | 0.387 | 0.396 | 0.467 |
| 29 | 0.645 | 0.621 | 0.624 | 0.662 |
| 30 | 0.421 | 0.408 | 0.415 | 0.468 |
| 31 | 0.426 | 0.42 | 0.415 | 0.482 |
| 32 | 0.408 | 0.41 | 0.41 | 0.459 |
| 33 | 0.384 | 0.371 | 0.377 | 0.438 |
| 34 | 0.338 | 0.348 | 0.352 | 0.419 |
| 35 | 0.336 | 0.327 | 0.327 | 0.394 |
| 36 | 0.413 | 0.422 | 0.426 | 0.481 |
| 37 | 0.498 | 0.5 | 0.499 | 0.57 |
| 38 | 0.535 | 0.535 | 0.543 | 0.598 |
| 39 | 0.472 | 0.491 | 0.497 | 0.573 |
| 40 | 0.870 | 0.877 | 0.879 | 0.892 |
| 41 | 0.523 | 0.521 | 0.527 | 0.596 |
| 42 | 0.865 | 0.877 | 0.883 | 0.909 |
| 43 | 0.722 | 0.715 | 0.736 | 0.789 |
| 44 | 0.526 | 0.534 | 0.55 | 0.616 |
| 45 | 0.548 | 0.564 | 0.567 | 0.623 |
| 46 | 0.821 | 0.821 | 0.831 | 0.859 |
| 47 | 0.445 | 0.437 | 0.439 | 0.493 |
| 48 | 0.381 | 0.377 | 0.39 | 0.446 |
| 49 | 0.422 | 0.428 | 0.426 | 0.496 |

| | | | | |
|---|---|---|---|---|
| 50 | 0.386 | 0.393 | 0.394 | 0.457 |
| 51 | 0.341 | 0.334 | 0.338 | 0.403 |
| 52 | 0.377 | 0.379 | 0.392 | 0.458 |
| 53 | 0.335 | 0.335 | 0.338 | 0.393 |
| 54 | 0.380 | 0.378 | 0.386 | 0.455 |
| 55 | 0.526 | 0.513 | 0.518 | 0.586 |
| 56 | 0.415 | 0.415 | 0.421 | 0.492 |
| 57 | 0.496 | 0.494 | 0.497 | 0.557 |
| 58 | 0.555 | 0.566 | 0.575 | 0.65 |
| 59 | 0.697 | 0.709 | 0.751 | 0.836 |
| 60 | 1.013 | 1.011 | 1.054 | 1.067 |
| 61 | 0.989 | 0.982 | 1.074 | 1.181 |
| 62 | 0.729 | 0.728 | 0.727 | 0.796 |
| 63 | 0.883 | 0.891 | 0.9 | 0.925 |
| 64 | 0.493 | 0.477 | 0.49 | 0.563 |
| 65 | 0.433 | 0.424 | 0.428 | 0.486 |
| 66 | 0.408 | 0.403 | 0.415 | 0.466 |
| 67 | 0.423 | 0.411 | 0.414 | 0.489 |
| 68 | 0.452 | 0.427 | 0.433 | 0.492 |
| 69 | 0.320 | 0.321 | 0.328 | 0.392 |
| 70 | 0.379 | 0.389 | 0.394 | 0.467 |
| 71 | 0.432 | 0.422 | 0.423 | 0.489 |
| 72 | 0.524 | 0.541 | 0.557 | 0.629 |
| 73 | 0.496 | 0.507 | 0.516 | 0.586 |
| 74 | 0.580 | 0.575 | 0.587 | 0.646 |
| 75 | 0.486 | 0.491 | 0.512 | 0.575 |
| 76 | 0.457 | 0.461 | 0.467 | 0.542 |
| 77 | 0.481 | 0.482 | 0.489 | 0.568 |
| 78 | 0.872 | 0.855 | 0.863 | 0.905 |
| 79 | 0.849 | 0.848 | 0.856 | 0.881 |
| 80 | 0.461 | 0.463 | 0.464 | 0.524 |
| 81 | 0.463 | 0.445 | 0.445 | 0.515 |
| 82 | 0.442 | 0.474 | 0.472 | 0.54 |
| 83 | 0.417 | 0.404 | 0.407 | 0.469 |
| 84 | 0.413 | 0.402 | 0.406 | 0.454 |
| 85 | 0.329 | 0.337 | 0.346 | 0.416 |
| 86 | 0.333 | 0.314 | 0.322 | 0.379 |
| 87 | 0.349 | 0.352 | 0.355 | 0.419 |
| 88 | 0.366 | 0.363 | 0.373 | 0.428 |
| 89 | 0.384 | 0.391 | 0.397 | 0.468 |
| 90 | 0.438 | 0.408 | 0.414 | 0.473 |
| 91 | 0.502 | 0.48 | 0.496 | 0.547 |
| 92 | 0.516 | 0.535 | 0.534 | 0.598 |
| 93 | 0.453 | 0.477 | 0.488 | 0.549 |
| 94 | 0.519 | 0.508 | 0.509 | 0.568 |
| 95 | 0.531 | 0.527 | 0.539 | 0.6 |
| 96 | 0.536 | 0.545 | 0.554 | 0.62 |
| 97 | 0.611 | 0.599 | 0.612 | 0.674 |
| 98 | 0.425 | 0.434 | 0.448 | 0.5 |
| 99 | 0.508 | 0.494 | 0.5 | 0.564 |
| 100 | 0.391 | 0.388 | 0.395 | 0.462 |
| 101 | 0.396 | 0.393 | 0.401 | 0.459 |
| 102 | 0.347 | 0.353 | 0.365 | 0.422 |

| | | | | |
|---|---|---|---|---|
| 103 | 0.362 | 0.348 | 0.36 | 0.414 |
| 104 | 0.373 | 0.367 | 0.369 | 0.435 |
| 105 | 0.399 | 0.392 | 0.4 | 0.453 |
| 106 | 0.426 | 0.406 | 0.407 | 0.471 |
| 107 | 0.338 | 0.337 | 0.334 | 0.399 |
| 108 | 0.333 | 0.324 | 0.327 | 0.379 |
| 109 | 0.345 | 0.34 | 0.338 | 0.394 |
| 110 | 0.565 | 0.553 | 0.57 | 0.602 |
| 111 | 0.387 | 0.383 | 0.388 | 0.457 |
| 112 | 0.783 | 0.781 | 0.783 | 0.805 |
| 113 | 0.399 | 0.408 | 0.423 | 0.481 |
| 114 | 0.357 | 0.368 | 0.369 | 0.435 |
| 115 | 0.495 | 0.494 | 0.498 | 0.571 |
| 116 | 0.552 | 0.548 | 0.556 | 0.625 |
| 117 | 0.454 | 0.455 | 0.455 | 0.521 |
| 118 | 0.488 | 0.499 | 0.508 | 0.592 |
| 119 | 1.043 | 1.053 | 1.154 | 1.298 |
| 120 | 0.695 | 0.727 | 0.746 | 0.859 |
| 121 | 0.591 | 0.596 | 0.615 | 0.691 |
| 122 | 0.823 | 0.822 | 0.848 | 0.92 |
| 123 | 0.510 | 0.512 | 0.523 | 0.59 |
| 124 | 0.477 | 0.49 | 0.496 | 0.582 |
| 125 | 0.699 | 0.717 | 0.728 | 0.764 |
| 126 | 0.437 | 0.453 | 0.458 | 0.53 |
| 127 | 0.517 | 0.517 | 0.529 | 0.599 |
| 128 | 0.523 | 0.532 | 0.533 | 0.606 |
| 129 | 0.466 | 0.481 | 0.492 | 0.552 |
| 130 | 0.435 | 0.438 | 0.447 | 0.509 |
| 131 | 0.462 | 0.466 | 0.471 | 0.544 |
| 132 | 0.456 | 0.452 | 0.46 | 0.522 |
| 133 | 0.468 | 0.475 | 0.474 | 0.532 |
| 134 | 0.414 | 0.418 | 0.434 | 0.494 |
| 135 | 0.571 | 0.563 | 0.562 | 0.616 |
| 136 | 0.679 | 0.662 | 0.667 | 0.732 |
| 137 | 0.574 | 0.556 | 0.566 | 0.621 |
| 138 | 0.432 | 0.429 | 0.436 | 0.498 |
| 139 | 0.663 | 0.645 | 0.645 | 0.692 |
| 140 | 0.597 | 0.598 | 0.599 | 0.666 |
| 141 | 0.857 | 0.851 | 0.871 | 0.907 |
| 142 | 0.948 | 0.913 | 0.953 | 1.029 |
| 143 | 0.685 | 0.705 | 0.712 | 0.77 |
| 144 | 0.406 | 0.391 | 0.394 | 0.455 |
| 145 | 0.337 | 0.347 | 0.349 | 0.406 |
| 146 | 0.418 | 0.413 | 0.419 | 0.467 |
| 147 | 0.330 | 0.335 | 0.339 | 0.402 |
| 148 | 0.325 | 0.325 | 0.334 | 0.397 |
| 149 | 0.308 | 0.302 | 0.306 | 0.364 |
| 150 | 0.353 | 0.348 | 0.355 | 0.42 |
| 151 | 0.357 | 0.364 | 0.368 | 0.418 |
| 152 | 0.357 | 0.344 | 0.348 | 0.411 |
| 153 | 0.350 | 0.342 | 0.344 | 0.397 |

| | | | | |
|---|---|---|---|---|
| 154 | 0.323 | 0.313 | 0.318 | 0.379 |
| 155 | 0.877 | 0.857 | 0.861 | 0.875 |
| 156 | 0.349 | 0.345 | 0.348 | 0.418 |
| 157 | 0.325 | 0.327 | 0.329 | 0.394 |
| 158 | 0.361 | 0.362 | 0.366 | 0.43 |
| 159 | 0.795 | 0.797 | 0.806 | 0.832 |
| 160 | 0.838 | 0.831 | 0.812 | 0.85 |
| 161 | 0.524 | 0.536 | 0.544 | 0.617 |
| 162 | 0.557 | 0.547 | 0.557 | 0.621 |
| 163 | 0.709 | 0.707 | 0.721 | 0.781 |
| 164 | 0.444 | 0.44 | 0.439 | 0.503 |
| 165 | 0.431 | 0.42 | 0.428 | 0.49 |
| 166 | 0.595 | 0.608 | 0.603 | 0.646 |
| 167 | 0.350 | 0.335 | 0.337 | 0.39 |
| 168 | 0.852 | 0.851 | 0.855 | 0.872 |
| 169 | 0.360 | 0.344 | 0.345 | 0.401 |
| 170 | 0.374 | 0.364 | 0.369 | 0.438 |
| 171 | 0.334 | 0.325 | 0.325 | 0.391 |
| 172 | 0.391 | 0.404 | 0.407 | 0.468 |
| 173 | 0.377 | 0.359 | 0.368 | 0.415 |
| 174 | 0.391 | 0.382 | 0.388 | 0.452 |
| 175 | 0.478 | 0.469 | 0.485 | 0.54 |
| 176 | 0.457 | 0.455 | 0.465 | 0.53 |
| 177 | 0.371 | 0.382 | 0.384 | 0.446 |
| 178 | 0.689 | 0.673 | 0.678 | 0.731 |
| 179 | 0.658 | 0.68 | 0.679 | 0.756 |
| 180 | 0.439 | 0.449 | 0.454 | 0.521 |
| 181 | 0.397 | 0.394 | 0.399 | 0.462 |
| 182 | 0.373 | 0.375 | 0.373 | 0.441 |
| 183 | 0.397 | 0.399 | 0.41 | 0.474 |
| 184 | 0.506 | 0.5 | 0.502 | 0.578 |
| 185 | 0.500 | 0.51 | 0.514 | 0.585 |
| 186 | 0.454 | 0.467 | 0.473 | 0.54 |
| 187 | 0.684 | 0.65 | 0.735 | 0.866 |
| 188 | 0.852 | 0.831 | 0.856 | 0.873 |
| 189 | 0.597 | 0.574 | 0.569 | 0.636 |
| 190 | 0.649 | 0.652 | 0.651 | 0.716 |
| 191 | 0.549 | 0.546 | 0.553 | 0.618 |
| 192 | 0.547 | 0.537 | 0.552 | 0.628 |
| 193 | 0.485 | 0.481 | 0.484 | 0.553 |
| 194 | 0.618 | 0.666 | 0.662 | 0.741 |
| 195 | 0.480 | 0.499 | 0.511 | 0.577 |
| 196 | 0.549 | 0.54 | 0.545 | 0.632 |
| 197 | 0.499 | 0.491 | 0.497 | 0.566 |
| 198 | 0.484 | 0.497 | 0.5 | 0.568 |
| 199 | 0.433 | 0.429 | 0.435 | 0.494 |

| | | | | |
|---|---|---|---|---|
| 200 | 0.465 | 0.447 | 0.45 | 0.523 |
| 201 | 0.404 | 0.395 | 0.394 | 0.456 |
| 202 | 0.359 | 0.353 | 0.357 | 0.423 |
| 203 | 0.329 | 0.326 | 0.324 | 0.384 |
| 204 | 0.363 | 0.342 | 0.344 | 0.414 |
| 205 | 0.409 | 0.419 | 0.422 | 0.488 |
| 206 | 0.631 | 0.627 | 0.631 | 0.689 |
| 207 | 0.728 | 0.737 | 0.744 | 0.812 |
| **Average** | 0.501 | 0.499 | 0.506 | 0.568 |

**Table S18. (1-$S^2$) values obtained from increasingly damaged (merged) thaumatin room temperature (277K) datasets.** (1-$S^2$) values were calculated from thaumatin multi-conformer models and as described in Materials and Methods. Dataset 1 (1-$S^2$) values are from **Table S11**.

| | $(1-S^2)$ | | | |
|---|---|---|---|---|
| **Residue #** | **Dataset 1** | **Dataset 2 (merged)** | **Dataset 3 (merged)** | **Dataset 4 (merged)** |
| **Dose (MGy)** | 0.006 | 0.023 | 0.041 | 0.058 |
| 1 | 0.364 | 0.36 | 0.353 | 0.401 |
| 2 | 0.304 | 0.305 | 0.31 | 0.36 |
| 3 | 0.217 | 0.206 | 0.204 | 0.256 |
| 4 | 0.297 | 0.284 | 0.287 | 0.333 |
| 5 | 0.712 | 0.708 | 0.696 | 0.715 |
| 6 | 0.245 | 0.245 | 0.241 | 0.294 |
| 7 | 0.229 | 0.228 | 0.231 | 0.284 |
| 8 | 0.196 | 0.192 | 0.188 | 0.235 |
| 9 | 0.191 | 0.192 | 0.187 | 0.238 |
| 10 | 0.234 | 0.222 | 0.218 | 0.266 |
| 11 | 0.233 | 0.223 | 0.221 | 0.272 |
| 12 | 0.195 | 0.187 | 0.183 | 0.234 |
| 13 | 0.521 | 0.512 | 0.513 | 0.549 |
| 14 | 0.191 | 0.192 | 0.193 | 0.247 |
| 15 | 0.738 | 0.749 | 0.752 | 0.762 |
| 16 | 0.281 | 0.293 | 0.295 | 0.345 |
| 17 | 0.310 | 0.32 | 0.317 | 0.367 |
| 18 | 0.259 | 0.253 | 0.246 | 0.306 |
| 19 | 0.330 | 0.33 | 0.326 | 0.374 |
| 20 | 0.298 | 0.302 | 0.298 | 0.352 |
| 21 | 0.300 | 0.305 | 0.299 | 0.347 |
| 22 | 0.270 | 0.268 | 0.262 | 0.308 |
| 23 | 0.193 | 0.189 | 0.186 | 0.241 |
| 24 | 0.273 | 0.27 | 0.264 | 0.315 |
| 25 | 0.206 | 0.203 | 0.202 | 0.259 |
| 26 | 0.222 | 0.222 | 0.223 | 0.276 |
| 27 | 0.663 | 0.635 | 0.638 | 0.671 |
| 28 | 0.286 | 0.29 | 0.291 | 0.345 |
| 29 | 0.258 | 0.277 | 0.274 | 0.326 |
| 30 | 0.202 | 0.21 | 0.207 | 0.258 |
| 31 | 0.653 | 0.663 | 0.659 | 0.698 |
| 32 | 0.220 | 0.216 | 0.213 | 0.261 |
| 33 | 0.342 | 0.371 | 0.374 | 0.425 |
| 34 | 0.226 | 0.225 | 0.227 | 0.278 |
| 35 | 0.252 | 0.261 | 0.258 | 0.307 |
| 36 | 0.193 | 0.185 | 0.183 | 0.239 |
| 37 | 0.213 | 0.206 | 0.209 | 0.258 |
| 38 | 0.190 | 0.193 | 0.191 | 0.242 |
| 39 | 0.185 | 0.176 | 0.173 | 0.224 |
| 40 | 0.180 | 0.166 | 0.163 | 0.217 |
| 41 | 0.203 | 0.201 | 0.198 | 0.25 |
| 42 | 0.222 | 0.237 | 0.235 | 0.284 |
| 43 | 0.265 | 0.271 | 0.269 | 0.316 |
| 44 | 0.324 | 0.313 | 0.318 | 0.363 |
| 45 | 0.372 | 0.387 | 0.384 | 0.429 |
| 46 | 0.249 | 0.245 | 0.241 | 0.296 |
| 47 | 0.229 | 0.241 | 0.243 | 0.295 |
| 48 | 0.185 | 0.194 | 0.189 | 0.238 |
| 49 | 0.213 | 0.216 | 0.214 | 0.262 |
| 50 | 0.292 | 0.289 | 0.285 | 0.33 |

| | | | | |
|---|---|---|---|---|
| 51 | 0.262 | 0.275 | 0.273 | 0.317 |
| 52 | 0.165 | 0.173 | 0.172 | 0.222 |
| 53 | 0.246 | 0.229 | 0.23 | 0.287 |
| 54 | 0.882 | 0.878 | 0.88 | 0.886 |
| 55 | 0.252 | 0.255 | 0.256 | 0.307 |
| 56 | 0.243 | 0.242 | 0.24 | 0.291 |
| 57 | 0.242 | 0.246 | 0.241 | 0.298 |
| 58 | 0.198 | 0.209 | 0.204 | 0.254 |
| 59 | 0.234 | 0.223 | 0.221 | 0.277 |
| 60 | 0.237 | 0.235 | 0.229 | 0.283 |
| 61 | 0.205 | 0.203 | 0.203 | 0.258 |
| 62 | 0.309 | 0.29 | 0.284 | 0.334 |
| 63 | 0.755 | 0.758 | 0.756 | 0.782 |
| 64 | 0.788 | 0.78 | 0.772 | 0.797 |
| 65 | 0.207 | 0.198 | 0.196 | 0.244 |
| 66 | 0.213 | 0.222 | 0.22 | 0.266 |
| 67 | 0.204 | 0.193 | 0.191 | 0.242 |
| 68 | 0.181 | 0.18 | 0.177 | 0.227 |
| 69 | 0.223 | 0.218 | 0.214 | 0.268 |
| 70 | 0.167 | 0.169 | 0.165 | 0.219 |
| 71 | 0.199 | 0.203 | 0.204 | 0.251 |
| 72 | 0.180 | 0.176 | 0.17 | 0.223 |
| 73 | 0.173 | 0.173 | 0.173 | 0.221 |
| 74 | 0.171 | 0.168 | 0.17 | 0.222 |
| 75 | 0.158 | 0.155 | 0.154 | 0.203 |
| 76 | 0.172 | 0.159 | 0.156 | 0.201 |
| 77 | 0.178 | 0.173 | 0.171 | 0.224 |
| 78 | 0.226 | 0.218 | 0.216 | 0.259 |
| 79 | 0.170 | 0.165 | 0.162 | 0.212 |
| 80 | 0.225 | 0.248 | 0.248 | 0.299 |
| 81 | 0.203 | 0.202 | 0.201 | 0.251 |
| 82 | 0.211 | 0.203 | 0.204 | 0.257 |
| 83 | 0.175 | 0.173 | 0.166 | 0.218 |
| 84 | 0.183 | 0.181 | 0.177 | 0.232 |
| 85 | 0.187 | 0.193 | 0.193 | 0.247 |
| 86 | 0.211 | 0.214 | 0.21 | 0.262 |
| 87 | 0.229 | 0.223 | 0.223 | 0.272 |
| 88 | 0.213 | 0.215 | 0.21 | 0.257 |
| 89 | 0.718 | 0.715 | 0.706 | 0.731 |
| 90 | 0.219 | 0.217 | 0.214 | 0.257 |
| 91 | 0.217 | 0.215 | 0.214 | 0.261 |
| 92 | 0.172 | 0.173 | 0.167 | 0.219 |
| 93 | 0.204 | 0.204 | 0.201 | 0.246 |
| 94 | 0.167 | 0.168 | 0.165 | 0.219 |
| 95 | 0.209 | 0.198 | 0.194 | 0.246 |
| 96 | 0.225 | 0.231 | 0.233 | 0.289 |
| 97 | 0.303 | 0.277 | 0.275 | 0.335 |
| 98 | 0.321 | 0.314 | 0.313 | 0.358 |
| 99 | 0.353 | 0.353 | 0.346 | 0.403 |
| 100 | 0.314 | 0.316 | 0.306 | 0.358 |
| 101 | 0.840 | 0.843 | 0.843 | 0.849 |
| 102 | 0.341 | 0.342 | 0.342 | 0.399 |
| 103 | 0.395 | 0.392 | 0.397 | 0.45 |

| | | | | |
|---|---|---|---|---|
| 104 | 0.348 | 0.348 | 0.35 | 0.396 |
| 105 | 0.712 | 0.698 | 0.702 | 0.719 |
| 106 | 0.264 | 0.264 | 0.257 | 0.3 |
| 107 | 0.266 | 0.261 | 0.256 | 0.309 |
| 108 | 0.241 | 0.233 | 0.228 | 0.283 |
| 109 | 0.247 | 0.243 | 0.243 | 0.299 |
| 110 | 0.205 | 0.204 | 0.206 | 0.256 |
| 111 | 0.296 | 0.298 | 0.29 | 0.341 |
| 112 | 0.255 | 0.247 | 0.249 | 0.301 |
| 113 | 0.248 | 0.25 | 0.249 | 0.299 |
| 114 | 0.186 | 0.184 | 0.184 | 0.241 |
| 115 | 0.248 | 0.249 | 0.249 | 0.301 |
| 116 | 0.214 | 0.217 | 0.218 | 0.271 |
| 117 | 0.234 | 0.233 | 0.23 | 0.282 |
| 118 | 0.306 | 0.294 | 0.289 | 0.339 |
| 119 | 0.256 | 0.266 | 0.265 | 0.314 |
| 120 | 0.406 | 0.424 | 0.431 | 0.474 |
| 121 | 0.264 | 0.268 | 0.261 | 0.309 |
| 122 | 0.717 | 0.741 | 0.754 | 0.773 |
| 123 | 0.229 | 0.235 | 0.234 | 0.29 |
| 124 | 0.221 | 0.222 | 0.223 | 0.273 |
| 125 | 0.228 | 0.211 | 0.21 | 0.264 |
| 126 | 0.227 | 0.226 | 0.222 | 0.277 |
| 127 | 0.220 | 0.21 | 0.208 | 0.256 |
| 128 | 0.187 | 0.179 | 0.18 | 0.233 |
| 129 | 0.220 | 0.221 | 0.219 | 0.267 |
| 130 | 0.201 | 0.193 | 0.191 | 0.247 |
| 131 | 0.272 | 0.261 | 0.256 | 0.305 |
| 132 | 0.192 | 0.184 | 0.184 | 0.235 |
| 133 | 0.255 | 0.257 | 0.255 | 0.297 |
| 134 | 0.267 | 0.27 | 0.266 | 0.328 |
| 135 | 0.247 | 0.236 | 0.234 | 0.286 |
| 136 | 0.254 | 0.243 | 0.24 | 0.289 |
| 137 | 0.246 | 0.254 | 0.249 | 0.293 |
| 138 | 0.280 | 0.285 | 0.286 | 0.343 |
| 139 | 0.886 | 0.89 | 0.892 | 0.897 |
| 140 | 0.779 | 0.765 | 0.767 | 0.794 |
| 141 | 0.205 | 0.212 | 0.208 | 0.256 |
| 142 | 0.249 | 0.249 | 0.245 | 0.294 |
| 143 | 0.888 | 0.882 | 0.884 | 0.898 |
| 144 | 0.226 | 0.226 | 0.221 | 0.277 |
| 145 | 0.206 | 0.208 | 0.204 | 0.253 |
| 146 | 0.259 | 0.252 | 0.253 | 0.308 |
| 147 | 0.212 | 0.206 | 0.202 | 0.253 |
| 148 | 0.203 | 0.202 | 0.199 | 0.252 |
| 149 | 0.192 | 0.191 | 0.192 | 0.245 |
| 150 | 0.784 | 0.792 | 0.794 | 0.8 |
| 151 | 0.216 | 0.215 | 0.212 | 0.266 |
| 152 | 0.222 | 0.222 | 0.217 | 0.27 |
| 153 | 0.227 | 0.211 | 0.207 | 0.256 |
| 154 | 0.344 | 0.332 | 0.33 | 0.372 |
| 155 | 0.247 | 0.245 | 0.239 | 0.285 |
| 156 | 0.188 | 0.187 | 0.183 | 0.231 |

| | | | | |
|---|---|---|---|---|
| 157 | 0.571 | 0.571 | 0.559 | 0.603 |
| 158 | 0.188 | 0.198 | 0.196 | 0.25 |
| 159 | 0.205 | 0.197 | 0.193 | 0.247 |
| 160 | 0.193 | 0.195 | 0.193 | 0.244 |
| 161 | 0.720 | 0.715 | 0.709 | 0.733 |
| 162 | 0.515 | 0.503 | 0.49 | 0.547 |
| 163 | 0.240 | 0.228 | 0.228 | 0.283 |
| 164 | 0.289 | 0.295 | 0.293 | 0.343 |
| 165 | 0.225 | 0.232 | 0.231 | 0.275 |
| 166 | 0.215 | 0.202 | 0.198 | 0.249 |
| 167 | 0.608 | 0.613 | 0.608 | 0.647 |
| 168 | 0.238 | 0.234 | 0.234 | 0.291 |
| 169 | 0.250 | 0.245 | 0.243 | 0.299 |
| 170 | 0.224 | 0.223 | 0.219 | 0.273 |
| 171 | 0.192 | 0.194 | 0.189 | 0.236 |
| 172 | 0.217 | 0.199 | 0.199 | 0.246 |
| 173 | 0.192 | 0.192 | 0.189 | 0.243 |
| 174 | 0.224 | 0.22 | 0.218 | 0.271 |
| 175 | 0.271 | 0.268 | 0.267 | 0.317 |
| 176 | 0.369 | 0.354 | 0.354 | 0.398 |
| 177 | 0.184 | 0.185 | 0.182 | 0.234 |
| 178 | 0.202 | 0.199 | 0.196 | 0.245 |
| 179 | 0.161 | 0.168 | 0.168 | 0.219 |
| 180 | 0.165 | 0.169 | 0.164 | 0.219 |
| 181 | 0.166 | 0.158 | 0.158 | 0.209 |
| 182 | 0.236 | 0.224 | 0.22 | 0.274 |
| 183 | 0.172 | 0.183 | 0.179 | 0.233 |
| 184 | 0.209 | 0.213 | 0.21 | 0.261 |
| 185 | 0.269 | 0.257 | 0.26 | 0.314 |
| 186 | 0.294 | 0.3 | 0.301 | 0.343 |
| 187 | 0.236 | 0.228 | 0.23 | 0.283 |
| 188 | 0.228 | 0.225 | 0.219 | 0.27 |
| 189 | 0.238 | 0.235 | 0.233 | 0.284 |
| 190 | 0.209 | 0.197 | 0.193 | 0.238 |
| 191 | 0.302 | 0.277 | 0.276 | 0.33 |
| 192 | 0.228 | 0.232 | 0.231 | 0.282 |
| 193 | 0.190 | 0.177 | 0.173 | 0.225 |
| 194 | 0.196 | 0.187 | 0.184 | 0.235 |
| 195 | 0.254 | 0.249 | 0.251 | 0.295 |
| 196 | 0.210 | 0.223 | 0.218 | 0.268 |
| 197 | 0.668 | 0.678 | 0.683 | 0.697 |
| 198 | 0.603 | 0.578 | 0.579 | 0.602 |
| 199 | 0.286 | 0.282 | 0.274 | 0.325 |
| 200 | 0.212 | 0.211 | 0.206 | 0.257 |
| 201 | 0.193 | 0.182 | 0.18 | 0.229 |
| 202 | 0.188 | 0.19 | 0.189 | 0.241 |
| 203 | 0.182 | 0.173 | 0.171 | 0.225 |
| 204 | 0.188 | 0.176 | 0.174 | 0.222 |
| 205 | 0.199 | 0.2 | 0.2 | 0.245 |
| 206 | 0.199 | 0.186 | 0.183 | 0.233 |
| 207 | 0.735 | 0.729 | 0.723 | 0.752 |
| 208 | 0.189 | 0.186 | 0.183 | 0.234 |
| 209 | 0.212 | 0.201 | 0.203 | 0.25 |

| | | | | |
|---|---|---|---|---|
| 210 | 0.172 | 0.151 | 0.149 | 0.205 |
| 211 | 0.205 | 0.205 | 0.203 | 0.259 |
| 212 | 0.208 | 0.213 | 0.212 | 0.264 |
| 213 | 0.208 | 0.206 | 0.207 | 0.257 |
| 214 | 0.226 | 0.235 | 0.232 | 0.289 |
| 215 | 0.269 | 0.274 | 0.27 | 0.321 |
| 216 | 0.719 | 0.728 | 0.727 | 0.744 |
| 217 | 0.249 | 0.243 | 0.233 | 0.283 |
| 218 | 0.256 | 0.247 | 0.245 | 0.299 |
| 219 | 0.647 | 0.653 | 0.649 | 0.657 |
| 220 | 0.275 | 0.265 | 0.263 | 0.313 |
| 221 | 0.209 | 0.2 | 0.2 | 0.25 |
| 222 | 0.214 | 0.207 | 0.204 | 0.257 |
| 223 | 0.180 | 0.167 | 0.166 | 0.219 |
| 224 | 0.408 | 0.414 | 0.426 | 0.514 |
| 225 | 0.180 | 0.173 | 0.17 | 0.221 |
| 226 | 0.167 | 0.173 | 0.172 | 0.226 |
| 227 | 0.170 | 0.163 | 0.159 | 0.208 |
| 228 | 0.166 | 0.166 | 0.161 | 0.212 |
| 229 | 0.165 | 0.158 | 0.155 | 0.202 |
| 230 | 0.169 | 0.175 | 0.169 | 0.216 |
| 231 | 0.180 | 0.183 | 0.178 | 0.23 |
| 232 | 0.160 | 0.15 | 0.148 | 0.2 |
| 233 | 0.191 | 0.186 | 0.187 | 0.236 |
| 234 | 0.178 | 0.181 | 0.175 | 0.228 |
| 235 | 0.174 | 0.176 | 0.17 | 0.22 |
| 236 | 0.175 | 0.172 | 0.168 | 0.219 |
| 237 | 0.158 | 0.162 | 0.162 | 0.216 |
| 238 | 0.202 | 0.214 | 0.209 | 0.257 |
| 239 | 0.225 | 0.228 | 0.224 | 0.275 |
| 240 | 0.274 | 0.26 | 0.258 | 0.303 |
| 241 | 0.256 | 0.245 | 0.246 | 0.299 |
| 242 | 0.277 | 0.282 | 0.276 | 0.329 |
| 243 | 0.231 | 0.237 | 0.237 | 0.284 |
| 244 | 0.251 | 0.241 | 0.24 | 0.294 |
| 245 | 0.266 | 0.276 | 0.271 | 0.321 |
| 246 | 0.331 | 0.346 | 0.353 | 0.4 |
| 247 | 0.679 | 0.682 | 0.673 | 0.69 |
| 248 | 0.213 | 0.203 | 0.202 | 0.252 |
| 249 | 0.229 | 0.239 | 0.235 | 0.284 |
| 250 | 0.681 | 0.682 | 0.67 | 0.693 |
| 251 | 0.200 | 0.201 | 0.199 | 0.248 |
| 252 | 0.190 | 0.184 | 0.183 | 0.233 |
| 253 | 0.258 | 0.251 | 0.248 | 0.295 |
| 254 | 0.320 | 0.317 | 0.317 | 0.362 |
| 255 | 0.248 | 0.243 | 0.244 | 0.302 |
| 256 | 0.283 | 0.298 | 0.289 | 0.33 |
| 257 | 0.250 | 0.244 | 0.244 | 0.297 |
| 258 | 0.699 | 0.711 | 0.696 | 0.724 |
| 259 | 0.322 | 0.3 | 0.294 | 0.344 |
| 260 | 0.356 | 0.344 | 0.341 | 0.399 |
| 261 | 0.246 | 0.24 | 0.238 | 0.285 |
| 262 | 0.287 | 0.266 | 0.268 | 0.313 |

| | | | | |
|---|---|---|---|---|
| 263 | 0.255 | 0.242 | 0.238 | 0.29 |
| 264 | 0.305 | 0.296 | 0.287 | 0.335 |
| 265 | 0.537 | 0.526 | 0.524 | 0.564 |
| 266 | 0.617 | 0.629 | 0.641 | 0.672 |
| 267 | 0.401 | 0.383 | 0.38 | 0.439 |
| 268 | 0.235 | 0.226 | 0.222 | 0.273 |
| 269 | 0.282 | 0.265 | 0.258 | 0.303 |
| 270 | 0.226 | 0.217 | 0.213 | 0.271 |
| 271 | 0.242 | 0.234 | 0.231 | 0.283 |
| 272 | 0.248 | 0.235 | 0.234 | 0.291 |
| 273 | 0.206 | 0.191 | 0.191 | 0.244 |
| 274 | 0.221 | 0.208 | 0.207 | 0.26 |
| 275 | 0.242 | 0.232 | 0.227 | 0.275 |
| 276 | 0.803 | 0.794 | 0.799 | 0.804 |
| 277 | 0.272 | 0.278 | 0.276 | 0.332 |
| 278 | 0.789 | 0.792 | 0.795 | 0.808 |
| 279 | 0.826 | 0.83 | 0.827 | 0.84 |
| **Average** | 0.290 | 0.288 | 0.286 | 0.333 |

**Table S19. (1-$S^2$) values obtained from increasingly damaged (merged) proteinase K room temperature (277K) datasets.** (1-$S^2$) values were calculated from thaumatin multi-conformer models and as described in Materials and Methods. Dataset 1 (1-$S^2$) values are from **Table S12**.

| | $(1-S^2)$ | | |
|---|---|---|---|
| **Residue #** | **Dataset 1 (merged)** | **Dataset 2 (merged)** | **Dataset 3 (merged)** |
| **Dose (MGy)** | 0.017 | 0.069 | 0.121 |
| 1 | 0.413 | 0.421 | 0.463 |
| 2 | 0.399 | 0.399 | 0.431 |
| 3 | 0.361 | 0.359 | 0.384 |
| 4 | 0.405 | 0.396 | 0.422 |
| 5 | 0.408 | 0.403 | 0.449 |
| 6 | 0.350 | 0.344 | 0.386 |
| 7 | 0.364 | 0.357 | 0.4 |
| 8 | 0.345 | 0.353 | 0.381 |
| 9 | 0.366 | 0.359 | 0.4 |
| 10 | 0.379 | 0.375 | 0.425 |
| 11 | 0.392 | 0.396 | 0.441 |
| 12 | 0.384 | 0.403 | 0.441 |
| 13 | 0.404 | 0.404 | 0.462 |
| 14 | 0.667 | 0.661 | 0.686 |
| 15 | 0.465 | 0.466 | 0.502 |
| 16 | 0.493 | 0.482 | 0.52 |
| 17 | 0.514 | 0.531 | 0.576 |
| 18 | 0.769 | 0.748 | 0.763 |
| 19 | 0.592 | 0.582 | 0.621 |
| 20 | 0.437 | 0.447 | 0.491 |
| 21 | 0.565 | 0.55 | 0.579 |
| 22 | 0.402 | 0.389 | 0.423 |
| 23 | 0.344 | 0.354 | 0.403 |
| 24 | 0.399 | 0.406 | 0.456 |
| 25 | 0.388 | 0.383 | 0.425 |
| 26 | 0.327 | 0.326 | 0.363 |
| 27 | 0.335 | 0.333 | 0.391 |
| 28 | 0.333 | 0.338 | 0.364 |
| 29 | 0.329 | 0.321 | 0.365 |
| 30 | 0.296 | 0.29 | 0.344 |
| 31 | 0.292 | 0.289 | 0.342 |
| 32 | 0.328 | 0.316 | 0.362 |
| 33 | 0.312 | 0.312 | 0.342 |
| 34 | 0.327 | 0.324 | 0.373 |
| 35 | 0.344 | 0.355 | 0.384 |
| 36 | 0.322 | 0.309 | 0.352 |
| 37 | 0.481 | 0.495 | 0.522 |
| 38 | 0.350 | 0.337 | 0.375 |
| 39 | 0.381 | 0.367 | 0.396 |
| 40 | 0.309 | 0.318 | 0.368 |
| 41 | 0.295 | 0.289 | 0.327 |
| 42 | 0.297 | 0.304 | 0.333 |
| 43 | 0.313 | 0.314 | 0.345 |
| 44 | 0.420 | 0.42 | 0.456 |
| 45 | 0.484 | 0.488 | 0.521 |
| 46 | 0.598 | 0.594 | 0.628 |
| 47 | 0.776 | 0.779 | 0.811 |
| 48 | 0.462 | 0.458 | 0.499 |
| 49 | 0.422 | 0.412 | 0.458 |

| | | | |
|---|---|---|---|
| 50 | 0.377 | 0.362 | 0.413 |
| 51 | 0.367 | 0.374 | 0.416 |
| 52 | 0.319 | 0.31 | 0.351 |
| 53 | 0.296 | 0.297 | 0.338 |
| 54 | 0.286 | 0.277 | 0.316 |
| 55 | 0.450 | 0.462 | 0.525 |
| 56 | 0.322 | 0.307 | 0.351 |
| 57 | 0.281 | 0.275 | 0.311 |
| 58 | 0.333 | 0.334 | 0.367 |
| 59 | 0.690 | 0.698 | 0.72 |
| 60 | 0.339 | 0.344 | 0.387 |
| 61 | 0.512 | 0.528 | 0.564 |
| 62 | 0.482 | 0.493 | 0.532 |
| 63 | 0.347 | 0.343 | 0.38 |
| 64 | 0.311 | 0.309 | 0.359 |
| 65 | 0.408 | 0.396 | 0.432 |
| 66 | 0.334 | 0.332 | 0.369 |
| 67 | 0.360 | 0.361 | 0.409 |
| 68 | 0.386 | 0.38 | 0.412 |
| 69 | 0.359 | 0.335 | 0.374 |
| 70 | 0.607 | 0.628 | 0.683 |
| 71 | 0.543 | 0.524 | 0.585 |
| 72 | 0.645 | 0.631 | 0.671 |
| 73 | 0.634 | 0.657 | 0.703 |
| 74 | 0.451 | 0.455 | 0.508 |
| 75 | 0.448 | 0.457 | 0.5 |
| 76 | 0.388 | 0.395 | 0.439 |
| 77 | 0.834 | 0.835 | 0.839 |
| 78 | 0.659 | 0.656 | 0.682 |
| 79 | 0.592 | 0.583 | 0.627 |
| 80 | 0.304 | 0.297 | 0.336 |
| 81 | 0.699 | 0.713 | 0.741 |
| 82 | 0.465 | 0.459 | 0.512 |
| 83 | 0.377 | 0.377 | 0.408 |
| 84 | 0.388 | 0.373 | 0.419 |
| 85 | 0.478 | 0.476 | 0.528 |
| 86 | 0.818 | 0.817 | 0.833 |
| 87 | 0.459 | 0.475 | 0.532 |
| 88 | 0.447 | 0.448 | 0.49 |
| 89 | 0.416 | 0.44 | 0.495 |
| 90 | 0.440 | 0.432 | 0.491 |
| 91 | 0.393 | 0.393 | 0.421 |
| 92 | 0.424 | 0.41 | 0.472 |
| 93 | 0.606 | 0.57 | 0.632 |
| 94 | 0.384 | 0.39 | 0.438 |
| 95 | 0.384 | 0.392 | 0.451 |
| 96 | 0.390 | 0.397 | 0.446 |
| 97 | 0.862 | 0.859 | 0.874 |
| 98 | 0.390 | 0.384 | 0.427 |
| 99 | 0.362 | 0.362 | 0.413 |
| 100 | 0.532 | 0.554 | 0.607 |
| 101 | 0.716 | 0.71 | 0.745 |
| 102 | 0.581 | 0.605 | 0.632 |

| | | | |
|---|---|---|---|
| 103 | 0.709 | 0.708 | 0.732 |
| 104 | 0.390 | 0.383 | 0.432 |
| 105 | 0.326 | 0.317 | 0.359 |
| 106 | 0.357 | 0.359 | 0.399 |
| 107 | 0.381 | 0.378 | 0.418 |
| 108 | 0.320 | 0.32 | 0.356 |
| 109 | 0.810 | 0.809 | 0.827 |
| 110 | 0.429 | 0.414 | 0.454 |
| 111 | 0.324 | 0.334 | 0.379 |
| 112 | 0.725 | 0.725 | 0.746 |
| 113 | 0.379 | 0.387 | 0.433 |
| 114 | 0.385 | 0.384 | 0.429 |
| 115 | 0.331 | 0.322 | 0.359 |
| 116 | 0.365 | 0.37 | 0.413 |
| 117 | 0.459 | 0.444 | 0.492 |
| 118 | 0.392 | 0.401 | 0.454 |
| 119 | 0.477 | 0.474 | 0.526 |
| 120 | 0.436 | 0.44 | 0.497 |
| 121 | 0.809 | 0.812 | 0.843 |
| 122 | 0.472 | 0.503 | 0.548 |
| 123 | 0.396 | 0.39 | 0.43 |
| 124 | 0.401 | 0.397 | 0.442 |
| 125 | 0.911 | 0.911 | 0.922 |
| 126 | 0.444 | 0.433 | 0.469 |
| 127 | 0.373 | 0.372 | 0.417 |
| 128 | 0.865 | 0.861 | 0.882 |
| 129 | 0.703 | 0.724 | 0.771 |
| **Average** | 0.451 | 0.450 | 0.491 |

**Table S20. (1-$S^2$) values obtained from increasingly damaged (merged) lysozyme room temperature (277K) datasets.** (1-$S^2$) values were calculated from thaumatin multi-conformer models and as described in Materials and Methods. Dataset 1 (1-$S^2$) values are from **Table S13**.

| Proteinase K 100 K diffraction data collection statistics | | | | | | | |
|---|---|---|---|---|---|---|---|
| Dataset | 1 | 2 | 3 | 4 | 5 | 6 | 7 |
| Wavelength (Å) | 0.88557 | | | | | | |
| Resolution range (Å) | 34.82-0.90 (0.92-0.90) | 34.82-0.91 (0.93-0.91) | 34.84-0.95 (0.97-0.95) | 34.84-1.01 (1.03-1.01) | 34.86-1.06 (1.08-1.06) | 34.86-1.11 (1.13-1.11) | 34.86-1.16 (1.18-1.16) |
| Dose (MGy) | 0.3 | 2.1 | 3.9 | 5.7 | 7.5 | 9.3 | 11.0 |
| Space group | $P4_32_12$ | | | | | | |
| Unit cell | 67.72 67.72 101.42 90.00 90.00 90.00 | 67.74 67.74 101.52 90.00 90.00 90.00 | 67.75 67.75 101.55 90.00 90.00 90.00 | 67.75 67.75 101.62 90.00 90.00 90.00 | 67.75 67.75 101.65 90.00 90.00 90.00 | 67.76 67.76 101.70 90.00 90.00 90.00 | 67.73 67.73 101.66 90.00 90.00 90.00 |
| Unit cell volume (Å$^3$) | 465112 | 465845.6 | 466120.8 | 466442.2 | 466579.9 | 466947.2 | 466350.3 |
| Total reflections | 1196300 (17467) | 1184687 (19176) | 1113598 (31210) | 993533 (29085) | 895131 (32249) | 799556 (33561) | 702050 (32297) |
| Multiplicity | 6.9 (2.2) | 7.1 (2.9) | 7.5 (4.4) | 8.0 (5.1) | 8.3 (6.2) | 8.5 (7.3) | 8.5 (8.2) |
| Mosaicity (°) | 0.18 | 0.19 | 0.19 | 0.19 | 0.19 | 0.19 | 0.19 |
| Completeness (%) | 99.6 (93.0) | 99.9 (81.1) | 99.9 (98.5) | 99.8 (95.7) | 100.0 (99.5) | 100.0 (99.9) | 99.9 (98.0) |
| Mean I/sigma(I) | 10.3 (1.0) | 10.4 (0.5) | 11.2 (0.6) | 11.8 (0.6) | 10.5 (0.7) | 12.7 (0.7) | 12.7 (0.7) |
| Wilson B-factor | 9.8 | 11.5 | 12.7 | 14.7 | 16.6 | 18.8 | 21.3 |
| R-merge | 0.076 (0.716) | 0.073 (1.436) | 0.073 (2.014) | 0.062 (1.965) | 0.067 (2.426) | 0.061 (2.412) | 0.061 (2.815) |
| R-pim | 0.029 (0.519) | 0.028 (0.922) | 0.027 (1.055) | 0.023 (0.922) | 0.024 (1.035) | 0.022 (0.941) | 0.022 (1.033) |
| $CC_{1/2}$ | 1.000 (0.604) | 1.000 (0.364) | 1.000 (0.306) | 1.000 (0.324) | 1.000 (0.340) | 1.000 (0.346) | 1.000 (0.312) |
| Isa | 20.7 | 21.1 | 24.0 | 25.4 | 26.4 | 28.7 | 31.0 |
| **Proteinase K crystal structure refinement statistics** | | | | | | | |
| PDB code | 7LTD | 7LTI | 7LTV | 7LU0 | 7LU1 | 7LQC | 7LU3 |
| Resolution range (Å) | 32.12 - 0.9 (0.93 - 0.90) | 32.12 - 0.91 (0.94 - 0.91) | 32.13 - 0.95 (0.98 - 0.95) | 33.88 - 1.01 (1.05 - 1.01) | 33.88 - 1.06 (1.10 - 1.06) | 32.14 - 1.11 (1.15 - 1.11) | 33.87 - 1.16 (1.20 - 1.16) |
| Reflections used in refinement | 173034 (16402) | 165151 (13733) | 147039 (13382) | 122587 (10937) | 106888 (10055) | 93318 (8735) | 81806 (7794) |
| Rwork | 0.157 (0.298) | 0.153 (0.315) | 0.150 (0.308) | 0.146 (0.321) | 0.144 (0.3003) | 0.145 (0.291) | 0.141 (0.305) |
| Rfree | 0.172 (0.310) | 0.170 (0.314) | 0.170 (0.314) | 0.168 (0.323) | 0.169 (0.313) | 0.173 (0.301) | 0.172 (0.311) |
| No. on non-hydrogen atoms | 3078 | 3070 | 3049 | 3004 | 2946 | 2850 | 2786 |
| Protein | 2492 | 2524 | 2555 | 2541 | 2533 | 2495 | 2473 |
| Ligand/ion | 21 | 21 | 21 | 21 | 17 | 13 | 13 |
| Water | 461 | 428 | 395 | 376 | 334 | 292 | 267 |
| RMS (bonds) | 0.005 | 0.006 | 0.006 | 0.006 | 0.006 | 0.007 | 0.007 |

| | | | | | | | |
|---|---|---|---|---|---|---|---|
| RMS (angles) | 0.9 | 0.9 | 0.89 | 0.89 | 0.89 | 0.89 | 0.89 |
| Average B-factor | 9.99 | 12.07 | 13.53 | 15.2 | 17.72 | 19.65 | 21.92 |
| Protein | 7.47 | 9.58 | 11.01 | 12.87 | 15.32 | 17.58 | 19.67 |
| Ligand/ion | 20.34 | 27.69 | 36.59 | 35.5 | 52.25 | 49.43 | 46.69 |
| Water | 20.73 | 23.42 | 26.16 | 27.63 | 31.61 | 33.59 | 39.39 |
| Ramachandran (%) | | | | | | | |
| Favored | 97.47 | 97.47 | 97.47 | 97.47 | 97.11 | 97.11 | 97.11 |
| Allowed | 2.53 | 2.53 | 2.17 | 2.17 | 2.53 | 2.53 | 2.53 |
| Outliers | 0 | 0 | 0.36 | 0.36 | 0.36 | 0.36 | 0.36 |

**Table S21**. **Diffraction data collection and refinement statistics for increasingly X-ray damaged proteinase K datasets obtained from a single cryo-cooled (100 K) crystal.** Diffraction statistics are reported for increasingly damaged dataset, each of 120° total rotation. Values in parenthesis are for the highest resolution shells. All diffraction statistics were obtained from Aimless (41), with the exception of $CC_{1/2}$, which was obtained from XSCALE (39). Average diffraction weighted doses (DWD) were estimated using the program RADDOSE 3D (35, 36). Structural models were obtained in a highly consistent manner and as described in Materials and Methods. Refinement statistics were obtained from phenix (*phenix.table_one*) using the final refined models and reflections files.

| | Pearson correlation coefficients ($P_{CC}$) | | | | | |
|---|---|---|---|---|---|---|
| **Residue #** | **Datasets 1v2** | **Datasets 1v3** | **Datasets 1v4** | **Datasets 1v5** | **Datasets 1v6** | **Datasets 1v7** |
| 3 | 0.999 | 0.996 | 0.995 | 0.987 | 0.983 | 0.986 |
| 4 | 0.997 | 0.993 | 0.987 | 0.979 | 0.975 | 0.969 |
| 5 | 0.991 | 0.970 | 0.983 | 0.948 | 0.952 | 0.895 |
| 7 | 0.990 | 0.991 | 0.986 | 0.982 | 0.970 | 0.970 |
| 8 | 0.993 | 0.984 | 0.981 | 0.980 | 0.954 | 0.951 |
| 10 | 0.999 | 0.994 | 0.990 | 0.985 | 0.977 | 0.962 |
| 12 | 0.996 | 0.993 | 0.994 | 0.996 | 0.992 | 0.990 |
| 13 | 0.995 | 0.980 | 0.961 | 0.945 | 0.908 | 0.850 |
| 14 | 0.999 | 0.999 | 0.998 | 0.994 | 0.993 | 0.967 |
| 15 | 0.998 | 0.986 | 0.986 | 0.985 | 0.970 | 0.989 |
| 16 | 0.996 | 0.993 | 0.991 | 0.980 | 0.963 | 0.930 |
| 17 | 0.997 | 0.984 | 0.991 | 0.982 | 0.980 | 0.973 |
| 18 | 0.990 | 0.979 | 0.968 | 0.925 | 0.969 | 0.944 |
| 20 | 0.998 | 0.996 | 0.996 | 0.993 | 0.978 | 0.967 |
| 21 | 0.993 | 0.987 | 0.984 | 0.986 | 0.987 | 0.990 |
| 22 | 0.997 | 0.996 | 0.992 | 0.995 | 0.992 | 0.985 |
| 23 | 0.999 | 0.999 | 0.998 | 0.994 | 0.997 | 0.992 |
| 24 | 0.997 | 0.996 | 0.994 | 0.992 | 0.991 | 0.981 |
| 25 | 0.997 | 0.995 | 0.988 | 0.986 | 0.982 | 0.977 |
| 26 | 0.994 | 0.993 | 0.984 | 0.982 | 0.978 | 0.972 |
| 27 | 0.994 | 0.998 | 0.996 | 0.988 | 0.983 | 0.977 |
| 28 | 1.000 | 0.997 | 0.995 | 0.993 | 0.988 | 0.984 |
| 31 | 0.987 | 0.980 | 0.977 | 0.966 | 0.975 | 0.962 |
| 33 | 0.999 | 0.997 | 0.996 | 0.996 | 0.992 | 0.996 |
| 34 | 0.998 | 0.993 | 0.986 | 0.956 | 0.945 | 0.921 |
| 35 | 0.997 | 0.993 | 0.992 | 0.986 | 0.986 | 0.982 |
| 36 | 0.997 | 0.997 | 0.995 | 0.983 | 0.980 | 0.982 |
| 37 | 0.998 | 0.995 | 0.987 | 0.984 | 0.989 | 0.982 |
| 38 | 0.998 | 0.994 | 0.993 | 0.991 | 0.997 | 0.990 |
| 39 | 0.992 | 0.987 | 0.969 | 0.974 | 0.978 | 0.966 |
| 40 | 0.999 | 0.995 | 0.994 | 0.987 | 0.984 | 0.982 |
| 42 | 0.998 | 0.992 | 0.991 | 0.984 | 0.979 | 0.981 |
| 43 | 0.998 | 0.996 | 0.997 | 0.998 | 0.994 | 0.990 |
| 45 | 0.996 | 0.994 | 0.985 | 0.983 | 0.964 | 0.953 |
| 46 | 0.999 | 0.998 | 0.999 | 0.996 | 0.999 | 0.984 |
| 47 | 0.991 | 0.970 | 0.959 | 0.905 | 0.882 | 0.916 |
| 48 | 0.996 | 0.997 | 0.995 | 0.990 | 0.980 | 0.967 |
| 49 | 0.998 | 0.990 | 0.985 | 0.980 | 0.973 | 0.966 |
| 50 | 0.993 | 0.994 | 0.992 | 0.986 | 0.973 | 0.972 |
| 52 | 0.999 | 0.994 | 0.989 | 0.990 | 0.990 | 0.994 |
| 54 | 0.996 | 0.986 | 0.935 | 0.890 | 0.767 | 0.742 |
| 55 | 0.997 | 0.991 | 0.987 | 0.984 | 0.975 | 0.957 |
| 56 | 0.990 | 0.985 | 0.975 | 0.976 | 0.981 | 0.969 |
| 57 | 0.998 | 0.997 | 0.991 | 0.988 | 0.980 | 0.957 |
| 58 | 0.998 | 0.998 | 0.997 | 0.995 | 0.994 | 0.993 |
| 59 | 0.998 | 0.996 | 0.992 | 0.990 | 0.990 | 0.973 |
| 60 | 0.996 | 0.990 | 0.983 | 0.968 | 0.971 | 0.963 |
| 61 | 0.996 | 0.988 | 0.989 | 0.950 | 0.962 | 0.965 |
| 62 | 0.997 | 0.981 | 0.978 | 0.965 | 0.990 | 0.987 |
| 63 | 0.982 | 0.980 | 0.978 | 0.926 | 0.906 | 0.914 |
| 64 | 0.999 | 0.995 | 0.988 | 0.951 | 0.970 | 0.934 |

| | | | | | | |
|---|---|---|---|---|---|---|
| 65 | 0.997 | 0.995 | 0.992 | 0.997 | 0.994 | 0.993 |
| 67 | 0.992 | 0.984 | 0.946 | 0.958 | 0.946 | 0.938 |
| 69 | 0.990 | 0.968 | 0.957 | 0.958 | 0.939 | 0.950 |
| 71 | 0.995 | 0.994 | 0.984 | 0.986 | 0.979 | 0.986 |
| 72 | 0.999 | 0.995 | 0.994 | 0.986 | 0.984 | 0.979 |
| 73 | 0.998 | 0.996 | 0.990 | 0.987 | 0.983 | 0.976 |
| 76 | 0.999 | 0.996 | 0.995 | 0.992 | 0.990 | 0.991 |
| 77 | 0.999 | 0.998 | 0.997 | 0.994 | 0.992 | 0.990 |
| 79 | 0.999 | 0.999 | 0.993 | 0.992 | 0.985 | 0.985 |
| 80 | 0.994 | 0.994 | 0.993 | 0.988 | 0.980 | 0.977 |
| 81 | 0.997 | 0.991 | 0.983 | 0.979 | 0.973 | 0.972 |
| 82 | 0.995 | 0.993 | 0.987 | 0.981 | 0.978 | 0.980 |
| 84 | 0.997 | 0.996 | 0.996 | 0.991 | 0.987 | 0.978 |
| 86 | 0.997 | 0.993 | 0.988 | 0.990 | 0.988 | 0.990 |
| 87 | 0.997 | 0.999 | 0.998 | 0.991 | 0.995 | 0.986 |
| 88 | 0.997 | 0.990 | 0.984 | 0.978 | 0.970 | 0.972 |
| 89 | 0.997 | 0.989 | 0.977 | 0.970 | 0.974 | 0.987 |
| 90 | 0.999 | 0.996 | 0.989 | 0.995 | 0.985 | 0.981 |
| 91 | 0.997 | 0.996 | 0.993 | 0.994 | 0.992 | 0.984 |
| 93 | 0.997 | 0.995 | 0.989 | 0.990 | 0.990 | 0.984 |
| 94 | 0.997 | 0.997 | 0.993 | 0.992 | 0.983 | 0.983 |
| 95 | 0.998 | 0.994 | 0.988 | 0.980 | 0.986 | 0.987 |
| 96 | 0.999 | 0.998 | 0.995 | 0.992 | 0.993 | 0.972 |
| 97 | 0.999 | 0.998 | 0.995 | 0.983 | 0.987 | 0.989 |
| 98 | 0.997 | 0.997 | 0.997 | 0.990 | 0.989 | 0.983 |
| 99 | 0.943 | 0.952 | 0.972 | 0.948 | 0.976 | 0.916 |
| 101 | 0.984 | 0.932 | 0.925 | 0.729 | 0.621 | 0.694 |
| 103 | 0.994 | 0.991 | 0.989 | 0.979 | 0.984 | 0.967 |
| 104 | 0.997 | 0.988 | 0.990 | 0.986 | 0.977 | 0.997 |
| 105 | 0.993 | 0.983 | 0.972 | 0.967 | 0.963 | 0.973 |
| 106 | 0.995 | 0.989 | 0.980 | 0.967 | 0.955 | 0.942 |
| 107 | 0.998 | 0.994 | 0.993 | 0.987 | 0.986 | 0.977 |
| 108 | 0.997 | 0.994 | 0.989 | 0.980 | 0.972 | 0.970 |
| 111 | 0.994 | 0.991 | 0.991 | 0.986 | 0.990 | 0.986 |
| 112 | 0.986 | 0.973 | 0.854 | 0.856 | 0.691 | 0.731 |
| 113 | 0.994 | 0.989 | 0.982 | 0.984 | 0.980 | 0.975 |
| 114 | 0.997 | 0.994 | 0.991 | 0.991 | 0.991 | 0.979 |
| 116 | 0.995 | 0.992 | 0.989 | 0.986 | 0.984 | 0.989 |
| 117 | 0.995 | 0.991 | 0.987 | 0.983 | 0.980 | 0.975 |
| 118 | 0.996 | 0.995 | 0.994 | 0.989 | 0.981 | 0.984 |
| 119 | 0.993 | 0.992 | 0.987 | 0.972 | 0.961 | 0.965 |
| 120 | 0.995 | 0.997 | 0.998 | 0.995 | 0.971 | 0.969 |
| 121 | 0.992 | 0.974 | 0.956 | 0.930 | 0.918 | 0.871 |
| 122 | 0.996 | 0.996 | 0.992 | 0.982 | 0.973 | 0.984 |
| 123 | 0.973 | 0.967 | 0.953 | 0.957 | 0.951 | 0.923 |
| 124 | 0.998 | 0.992 | 0.986 | 0.984 | 0.978 | 0.971 |
| 125 | 0.997 | 0.990 | 0.988 | 0.977 | 0.974 | 0.978 |
| 127 | 0.996 | 0.992 | 0.987 | 0.977 | 0.964 | 0.969 |
| 128 | 0.997 | 0.994 | 0.991 | 0.991 | 0.989 | 0.989 |
| 130 | 0.999 | 0.996 | 0.989 | 0.996 | 0.990 | 0.988 |
| 131 | 0.994 | 0.982 | 0.967 | 0.967 | 0.972 | 0.952 |
| 132 | 0.995 | 0.988 | 0.971 | 0.963 | 0.953 | 0.956 |
| 133 | 0.992 | 0.982 | 0.967 | 0.956 | 0.954 | 0.946 |

| | | | | | | |
|---|---|---|---|---|---|---|
| 137 | 0.994 | 0.981 | 0.968 | 0.951 | 0.939 | 0.934 |
| 138 | 0.997 | 0.996 | 0.997 | 0.997 | 0.997 | 0.994 |
| 139 | 0.999 | 0.999 | 0.992 | 0.976 | 0.949 | 0.906 |
| 140 | 0.993 | 0.974 | 0.947 | 0.879 | 0.755 | 0.623 |
| 141 | 0.998 | 0.996 | 0.992 | 0.991 | 0.987 | 0.976 |
| 142 | 0.998 | 0.999 | 0.999 | 0.997 | 0.996 | 0.993 |
| 143 | 0.983 | 0.975 | 0.953 | 0.929 | 0.906 | 0.827 |
| 147 | 0.998 | 0.998 | 0.995 | 0.993 | 0.984 | 0.981 |
| 148 | 0.999 | 0.998 | 0.998 | 0.992 | 0.994 | 0.992 |
| 149 | 0.977 | 0.966 | 0.948 | 0.950 | 0.959 | 0.983 |
| 150 | 0.999 | 0.995 | 0.988 | 0.976 | 0.960 | 0.980 |
| 151 | 0.991 | 0.986 | 0.982 | 0.979 | 0.983 | 0.968 |
| 153 | 0.998 | 0.996 | 0.995 | 0.993 | 0.994 | 0.986 |
| 154 | 0.996 | 0.991 | 0.987 | 0.974 | 0.958 | 0.951 |
| 155 | 0.995 | 0.993 | 0.993 | 0.991 | 0.993 | 0.992 |
| 157 | 0.998 | 0.992 | 0.988 | 0.979 | 0.977 | 0.987 |
| 161 | 0.996 | 0.990 | 0.989 | 0.985 | 0.985 | 0.982 |
| 162 | 0.996 | 0.995 | 0.986 | 0.966 | 0.970 | 0.950 |
| 163 | 0.992 | 0.993 | 0.993 | 0.992 | 0.992 | 0.982 |
| 165 | 0.998 | 0.997 | 0.992 | 0.982 | 0.990 | 0.975 |
| 167 | 0.999 | 0.989 | 0.990 | 0.981 | 0.970 | 0.997 |
| 168 | 0.991 | 0.987 | 0.977 | 0.968 | 0.957 | 0.955 |
| 169 | 0.997 | 0.996 | 0.993 | 0.978 | 0.974 | 0.976 |
| 170 | 0.998 | 0.995 | 0.996 | 0.996 | 0.996 | 0.996 |
| 171 | 0.998 | 0.996 | 0.995 | 0.969 | 0.953 | 0.973 |
| 173 | 0.999 | 0.997 | 0.997 | 0.997 | 0.993 | 0.994 |
| 174 | 0.997 | 0.994 | 0.989 | 0.975 | 0.970 | 0.983 |
| 175 | 0.993 | 0.991 | 0.989 | 0.991 | 0.986 | 0.976 |
| 176 | 0.995 | 0.990 | 0.986 | 0.972 | 0.963 | 0.960 |
| 177 | 0.998 | 0.996 | 0.995 | 0.988 | 0.971 | 0.983 |
| 178 | 0.986 | 0.976 | 0.969 | 0.948 | 0.923 | 0.911 |
| 179 | 0.997 | 0.997 | 0.996 | 0.994 | 0.994 | 0.992 |
| 180 | 0.999 | 0.998 | 0.997 | 0.995 | 0.987 | 0.987 |
| 183 | 0.999 | 0.999 | 0.997 | 0.994 | 0.992 | 0.996 |
| 184 | 0.999 | 0.995 | 0.997 | 0.997 | 0.988 | 0.998 |
| 185 | 0.995 | 0.990 | 0.978 | 0.953 | 0.934 | 0.948 |
| 186 | 0.996 | 0.985 | 0.973 | 0.972 | 0.972 | 0.964 |
| 187 | 0.999 | 0.997 | 0.996 | 0.996 | 0.994 | 0.981 |
| 188 | 0.999 | 0.998 | 0.993 | 0.992 | 0.988 | 0.995 |
| 189 | 0.995 | 0.995 | 0.988 | 0.993 | 0.989 | 0.987 |
| 190 | 0.999 | 0.995 | 0.994 | 0.973 | 0.983 | 0.977 |
| 191 | 0.967 | 0.917 | 0.872 | 0.766 | 0.790 | 0.775 |
| 192 | 0.999 | 0.991 | 0.987 | 0.988 | 0.967 | 0.987 |
| 193 | 0.998 | 0.996 | 0.998 | 0.995 | 0.993 | 0.993 |
| 194 | 0.999 | 0.998 | 0.995 | 0.995 | 0.997 | 0.988 |
| 195 | 0.993 | 0.990 | 0.986 | 0.980 | 0.971 | 0.933 |
| 197 | 0.967 | 0.890 | 0.887 | 0.832 | 0.819 | 0.793 |
| 198 | 0.996 | 0.994 | 0.984 | 0.961 | 0.972 | 0.955 |
| 199 | 0.999 | 0.994 | 0.993 | 0.992 | 0.989 | 0.983 |
| 200 | 0.994 | 0.991 | 0.989 | 0.985 | 0.965 | 0.984 |
| 201 | 0.999 | 0.997 | 0.994 | 0.982 | 0.965 | 0.972 |
| 202 | 0.999 | 0.997 | 0.996 | 0.996 | 0.990 | 0.983 |
| 204 | 0.997 | 0.996 | 0.984 | 0.981 | 0.946 | 0.929 |

| | | | | | | |
|---|---|---|---|---|---|---|
| 206 | 0.998 | 0.995 | 0.991 | 0.988 | 0.985 | 0.986 |
| 207 | 0.996 | 0.993 | 0.991 | 0.985 | 0.978 | 0.950 |
| 208 | 0.996 | 0.997 | 0.995 | 0.993 | 0.990 | 0.986 |
| 209 | 0.992 | 0.986 | 0.972 | 0.973 | 0.976 | 0.960 |
| 210 | 0.999 | 0.997 | 0.996 | 0.992 | 0.987 | 0.985 |
| 211 | 0.999 | 0.997 | 0.993 | 0.988 | 0.986 | 0.979 |
| 212 | 0.997 | 0.995 | 0.989 | 0.985 | 0.982 | 0.974 |
| 213 | 0.997 | 0.992 | 0.989 | 0.982 | 0.980 | 0.989 |
| 216 | 0.997 | 0.994 | 0.980 | 0.979 | 0.960 | 0.920 |
| 217 | 0.996 | 0.992 | 0.983 | 0.980 | 0.960 | 0.958 |
| 218 | 0.999 | 0.999 | 0.997 | 0.999 | 0.997 | 0.997 |
| 219 | 0.994 | 0.985 | 0.968 | 0.956 | 0.919 | 0.934 |
| 220 | 0.994 | 0.996 | 0.982 | 0.974 | 0.961 | 0.961 |
| 221 | 0.998 | 0.996 | 0.998 | 0.992 | 0.980 | 0.984 |
| 223 | 0.998 | 0.994 | 0.993 | 0.993 | 0.987 | 0.992 |
| 224 | 0.999 | 0.999 | 0.993 | 0.986 | 0.952 | 0.906 |
| 225 | 0.996 | 0.996 | 0.996 | 0.992 | 0.984 | 0.975 |
| 227 | 0.998 | 0.997 | 0.995 | 0.989 | 0.985 | 0.979 |
| 228 | 0.998 | 0.995 | 0.985 | 0.975 | 0.956 | 0.965 |
| 229 | 0.995 | 0.983 | 0.983 | 0.981 | 0.973 | 0.989 |
| 230 | 0.997 | 0.994 | 0.994 | 0.992 | 0.995 | 0.988 |
| 233 | 0.999 | 0.998 | 0.996 | 0.995 | 0.991 | 0.984 |
| 236 | 0.996 | 0.994 | 0.988 | 0.981 | 0.973 | 0.979 |
| 237 | 0.998 | 0.997 | 0.996 | 0.994 | 0.992 | 0.986 |
| 238 | 0.994 | 0.998 | 0.999 | 0.999 | 0.996 | 0.986 |
| 239 | 0.996 | 0.994 | 0.994 | 0.984 | 0.977 | 0.979 |
| 240 | 0.993 | 0.989 | 0.986 | 0.985 | 0.979 | 0.964 |
| 242 | 0.996 | 0.981 | 0.981 | 0.989 | 0.997 | 0.991 |
| 243 | 0.996 | 0.988 | 0.979 | 0.966 | 0.964 | 0.951 |
| 244 | 0.996 | 0.990 | 0.991 | 0.963 | 0.965 | 0.956 |
| 247 | 0.986 | 0.770 | 0.541 | 0.383 | 0.280 | 0.122 |
| 249 | 0.991 | 0.984 | 0.981 | 0.977 | 0.963 | 0.957 |
| 250 | 0.986 | 0.972 | 0.955 | 0.920 | 0.924 | 0.911 |
| 251 | 0.998 | 0.994 | 0.991 | 0.982 | 0.969 | 0.971 |
| 252 | 0.998 | 0.995 | 0.996 | 0.995 | 0.993 | 0.988 |
| 254 | 0.999 | 0.999 | 0.998 | 0.982 | 0.960 | 0.939 |
| 255 | 0.998 | 0.995 | 0.993 | 0.988 | 0.978 | 0.991 |
| 257 | 0.995 | 0.995 | 0.993 | 0.990 | 0.981 | 0.978 |
| 258 | 0.930 | 0.975 | 0.954 | 0.951 | 0.950 | 0.969 |
| 260 | 0.990 | 0.987 | 0.946 | 0.987 | 0.962 | 0.991 |
| 261 | 0.991 | 0.981 | 0.975 | 0.969 | 0.951 | 0.933 |
| 262 | 0.994 | 0.974 | 0.967 | 0.955 | 0.963 | 0.971 |
| 263 | 0.988 | 0.977 | 0.967 | 0.918 | 0.872 | 0.858 |
| 264 | 0.989 | 0.986 | 0.984 | 0.977 | 0.945 | 0.943 |
| 265 | 0.985 | 0.991 | 0.980 | 0.957 | 0.860 | 0.867 |
| 266 | 0.993 | 0.990 | 0.979 | 0.956 | 0.972 | 0.961 |
| 268 | 0.997 | 0.991 | 0.992 | 0.990 | 0.983 | 0.983 |
| 269 | 0.992 | 0.976 | 0.967 | 0.976 | 0.966 | 0.964 |
| 270 | 0.996 | 0.996 | 0.992 | 0.996 | 0.996 | 0.991 |
| 271 | 0.998 | 0.993 | 0.993 | 0.995 | 0.989 | 0.986 |
| 272 | 0.997 | 0.994 | 0.977 | 0.964 | 0.951 | 0.962 |
| 274 | 0.996 | 0.991 | 0.989 | 0.985 | 0.981 | 0.977 |
| 275 | 0.998 | 0.997 | 0.995 | 0.989 | 0.987 | 0.990 |

| | | | | | | |
|---|---|---|---|---|---|---|
| 276 | 0.982 | 0.902 | 0.863 | 0.694 | 0.734 | 0.515 |
| 277 | 0.997 | 0.990 | 0.987 | 0.976 | 0.973 | 0.977 |
| 278 | 0.992 | 0.981 | 0.961 | 0.940 | 0.858 | 0.773 |
| Average | 0.995 | 0.989 | 0.981 | 0.971 | 0.962 | 0.955 |
| Standard deviation | 0.007 | 0.020 | 0.036 | 0.055 | 0.068 | 0.082 |

**Table S22**. **Pearson correlation coefficients ($P_{CC}$) between *Ringer* profiles obtained from the least and increasingly X-ray damaged proteinase K datasets obtained from a single cryo-cooled (100 K) crystal.** Structural models (**Table S18**) were obtained in a highly consistent manner and as described in Materials and Methods. For each compared pair, *Ringer* profiles were calculated using models refined at the optimal resolution of the corresponding dataset (Table S21) and using electron density maps with matched resolutions such that the resolution of the electron density map from the least damaged dataset was matched to the resolution of the electron density map from the increasingly damaged dataset (see Materials and Methods and **Figure S9**).

| | Pearson correlation coefficients ($P_{CC}$) | |
|---|---|---|
| Residue # | 277 K dataset 1 vs 100 K dataset 1 | 277 K dataset 1 vs 100 K dataset 7 |
| 3 | 0.997 | 0.994 |
| 4 | 0.984 | 0.991 |
| 5 | 0.984 | 0.913 |
| 7 | 0.957 | 0.959 |
| 8 | 0.997 | 0.964 |
| 10 | 0.971 | 0.991 |
| 12 | 0.991 | 0.985 |
| 13 | 0.962 | 0.906 |
| 14 | 0.986 | 0.995 |
| 15 | 0.953 | 0.920 |
| 16 | 0.977 | 0.932 |
| 17 | 0.982 | 0.981 |
| 18 | 0.997 | 0.958 |
| 20 | 0.976 | 0.981 |
| 21 | 0.998 | 0.994 |
| 22 | 0.996 | 0.988 |
| 23 | 0.995 | 0.994 |
| 24 | 0.990 | 0.994 |
| 25 | 0.995 | 0.987 |
| 26 | 0.994 | 0.967 |
| 27 | 0.993 | 0.966 |
| 28 | 0.994 | 0.982 |
| 31 | 0.978 | 0.931 |
| 33 | 0.996 | 0.999 |
| 34 | 0.997 | 0.939 |
| 35 | 0.991 | 0.992 |
| 36 | 0.997 | 0.989 |
| 37 | 0.998 | 0.988 |
| 38 | 0.992 | 0.990 |
| 39 | 0.987 | 0.951 |
| 40 | 0.999 | 0.986 |
| 42 | 0.997 | 0.986 |
| 43 | 0.999 | 0.988 |
| 45 | 0.998 | 0.957 |
| 46 | 0.992 | 0.969 |
| 47 | 0.874 | 0.944 |
| 48 | 0.998 | 0.974 |
| 49 | 0.997 | 0.974 |
| 50 | 0.985 | 0.982 |
| 52 | 0.994 | 0.987 |
| 54 | 0.048 | 0.104 |
| 55 | 0.997 | 0.962 |
| 56 | 0.978 | 0.970 |
| 57 | 0.997 | 0.974 |
| 58 | 0.996 | 0.988 |
| 59 | 0.994 | 0.974 |
| 60 | 0.978 | 0.992 |
| 61 | 0.997 | 0.961 |
| 62 | 0.997 | 0.991 |
| 63 | 0.966 | 0.849 |
| 64 | 0.721 | 0.661 |

| | | |
|---|---|---|
| 65 | 0.999 | 0.996 |
| 67 | 0.998 | 0.951 |
| 69 | 0.998 | 0.960 |
| 71 | 0.991 | 0.989 |
| 72 | 0.997 | 0.977 |
| 73 | 0.998 | 0.979 |
| 76 | 0.994 | 0.993 |
| 77 | 0.997 | 0.992 |
| 79 | 0.998 | 0.976 |
| 80 | 0.993 | 0.989 |
| 81 | 0.991 | 0.983 |
| 82 | 0.996 | 0.983 |
| 84 | 0.997 | 0.983 |
| 86 | 0.994 | 0.995 |
| 87 | 0.981 | 0.984 |
| 88 | 0.985 | 0.991 |
| 89 | 0.802 | 0.773 |
| 90 | 0.997 | 0.987 |
| 91 | 0.998 | 0.979 |
| 93 | 0.996 | 0.985 |
| 94 | 0.993 | 0.990 |
| 95 | 0.992 | 0.991 |
| 96 | 0.995 | 0.988 |
| 97 | 0.993 | 0.988 |
| 98 | 0.993 | 0.994 |
| 99 | 0.974 | 0.850 |
| 101 | 0.486 | 0.693 |
| 103 | 0.982 | 0.968 |
| 104 | 0.999 | 0.994 |
| 105 | 0.981 | 0.932 |
| 106 | 0.987 | 0.933 |
| 107 | 0.996 | 0.983 |
| 108 | 0.990 | 0.991 |
| 111 | 0.999 | 0.985 |
| 112 | 0.772 | 0.928 |
| 113 | 0.998 | 0.983 |
| 114 | 0.987 | 0.991 |
| 116 | 0.995 | 0.996 |
| 117 | 0.994 | 0.979 |
| 118 | 0.981 | 0.985 |
| 119 | 0.990 | 0.989 |
| 120 | 0.984 | 0.982 |
| 121 | 0.997 | 0.901 |
| 122 | 0.964 | 0.972 |
| 123 | 0.999 | 0.935 |
| 124 | 0.992 | 0.989 |
| 125 | 0.994 | 0.991 |
| 127 | 0.996 | 0.976 |
| 128 | 0.995 | 0.989 |
| 130 | 0.998 | 0.995 |
| 131 | 0.994 | 0.970 |
| 132 | 0.993 | 0.944 |
| 133 | 0.987 | 0.951 |

| | | |
|---|---|---|
| 137 | 0.983 | 0.959 |
| 138 | 0.996 | 0.998 |
| 139 | 0.904 | 0.758 |
| 140 | 0.990 | 0.558 |
| 141 | 0.996 | 0.980 |
| 142 | 0.998 | 0.996 |
| 143 | 0.745 | 0.653 |
| 147 | 0.991 | 0.989 |
| 148 | 0.989 | 0.997 |
| 149 | 0.996 | 0.983 |
| 150 | 0.985 | 0.984 |
| 151 | 0.995 | 0.987 |
| 153 | 0.996 | 0.995 |
| 154 | 0.992 | 0.975 |
| 155 | 0.992 | 0.985 |
| 157 | 0.973 | 0.984 |
| 161 | 0.800 | 0.745 |
| 162 | 0.967 | 0.994 |
| 163 | 0.996 | 0.975 |
| 165 | 0.934 | 0.953 |
| 167 | 0.968 | 0.976 |
| 168 | 0.966 | 0.987 |
| 169 | 0.992 | 0.987 |
| 170 | 1.000 | 0.997 |
| 171 | 0.986 | 0.995 |
| 173 | 0.994 | 0.979 |
| 174 | 0.999 | 0.986 |
| 175 | 0.955 | 0.932 |
| 176 | 0.989 | 0.985 |
| 177 | 0.997 | 0.987 |
| 178 | 0.999 | 0.914 |
| 179 | 0.998 | 0.994 |
| 180 | 0.994 | 0.991 |
| 183 | 0.999 | 0.996 |
| 184 | 0.989 | 0.987 |
| 185 | 0.941 | 0.916 |
| 186 | 0.995 | 0.973 |
| 187 | 0.996 | 0.983 |
| 188 | 0.941 | 0.949 |
| 189 | 0.971 | 0.984 |
| 190 | 0.987 | 0.994 |
| 191 | 0.900 | 0.894 |
| 192 | 0.988 | 0.986 |
| 193 | 0.998 | 0.996 |
| 194 | 0.995 | 0.989 |
| 195 | 0.974 | 0.959 |
| 197 | 0.926 | 0.823 |
| 198 | 0.942 | 0.948 |
| 199 | 0.987 | 0.998 |
| 200 | 0.987 | 0.992 |
| 201 | 0.997 | 0.980 |
| 202 | 0.997 | 0.987 |
| 204 | 0.984 | 0.979 |

| | | |
|---|---|---|
| 206 | 0.994 | 0.989 |
| 207 | 0.952 | 0.974 |
| 208 | 0.987 | 0.987 |
| 209 | 0.998 | 0.971 |
| 210 | 0.999 | 0.989 |
| 211 | 0.994 | 0.988 |
| 212 | 0.997 | 0.972 |
| 213 | 0.993 | 0.988 |
| 216 | 0.839 | 0.971 |
| 217 | 0.985 | 0.959 |
| 218 | 0.999 | 0.993 |
| 219 | 0.778 | 0.836 |
| 220 | 0.997 | 0.972 |
| 221 | 0.990 | 0.997 |
| 223 | 0.967 | 0.957 |
| 224 | 0.998 | 0.922 |
| 225 | 0.991 | 0.963 |
| 227 | 0.993 | 0.986 |
| 228 | 0.986 | 0.980 |
| 229 | 0.993 | 0.988 |
| 230 | 0.995 | 0.995 |
| 233 | 0.997 | 0.994 |
| 236 | 0.999 | 0.978 |
| 237 | 0.990 | 0.982 |
| 238 | 0.981 | 0.992 |
| 239 | 0.987 | 0.990 |
| 240 | 0.989 | 0.962 |
| 242 | 0.960 | 0.949 |
| 243 | 0.996 | 0.961 |
| 244 | 0.993 | 0.936 |
| 247 | 0.943 | 0.384 |
| 249 | 0.997 | 0.948 |
| 250 | 0.950 | 0.957 |
| 251 | 0.992 | 0.979 |
| 252 | 0.996 | 0.995 |
| 254 | 0.998 | 0.948 |
| 255 | 0.996 | 0.990 |
| 257 | 0.997 | 0.980 |
| 258 | 0.951 | 0.938 |
| 260 | 0.943 | 0.952 |
| 261 | 0.976 | 0.978 |
| 262 | 0.914 | 0.967 |
| 263 | 0.721 | 0.902 |
| 264 | 0.877 | 0.925 |
| 265 | 0.814 | 0.952 |
| 266 | 0.862 | 0.753 |
| 268 | 0.988 | 0.982 |
| 269 | 0.966 | 0.889 |
| 270 | 0.984 | 0.978 |
| 271 | 0.998 | 0.993 |
| 272 | 0.987 | 0.989 |
| 274 | 0.991 | 0.992 |
| 275 | 0.990 | 0.997 |

| | | |
|---|---|---|
| 276 | 0.716 | 0.939 |
| 277 | 0.993 | 0.991 |
| 278 | 0.919 | 0.586 |

**Table S23**. **Pearson correlation coefficients ($P_{CC}$) between *Ringer* profiles obtained from the least X-ray damaged 277 K dataset 1 and the least (dataset 1) and most (dataset 7) X-ray damaged 100 K proteinase K datasets.** For each compared pair, *Ringer* profiles were calculated using models refined at the optimal resolution of the corresponding dataset and using electron density maps with matched resolutions at 1.16 Å (see Materials and Methods and **Figure S7**).

| **277 K crystal** | **100 K crystal** |
|---|---|
| T4 | S138 |
| T22 | S139 |
| S61 | S140 |
| S63 | R185 |
| D97 | T244 |
| S101 | S247 |
| R167 | |

**Table S24**. Proteinase K residues the side chains of which are in contact with crystallization components within the crystal.
